# Why we age: the four process model

**DOI:** 10.64898/2026.01.30.701154

**Authors:** J. Wordsworth, P. Yde Nielsen, E. Fielder, S. Chandrasegaran, D. Shanley

**Affiliations:** Biosciences Institute (NUBI), Newcastle University, Newcastle upon Tyne, Tyne & Wear, UK. NE4 5PL; Department of Applied Mathematics and Computer Science, Technical University of Denmark, 2800, Kongens Lyngby, Denmark

**Keywords:** Ageing, evolution, insulin resistance, inflammation, metabolism, homeostasis

## Abstract

**Highlights:**

- DNA damage-induced selection is the underlying cause of ageing.
- Faster metabolising cells naturally spread unless prevented by a deliberate process designed to prevent hyperfunctional diseases like cancer and fibrosis.
- As a result, slow metabolising mutants reduce the metabolic rate of faster neighbours epigenetically, shifting energy homeostasis to induce insulin resistance, weight gain, inflammation, metabolic syndrome and age-related disease.
- In post-mitotic tissues, mitochondria are undergoing similar selection and counterselection, contributing to the same metabolic slowdown.
- The four process model defined by these selection processes explains all major anti-ageing therapies and connects the hallmarks into a mechanistic framework.

Although ageing can be understood in terms of associated hallmarks and biomarkers, the processes which connect and cause these phenotypes are ill-defined. Here we suggest a unifying model of ageing as four distinct processes which connect the major observations and evidence into a single framework. It explains, from a single initial cause to the ultimate outcomes and diseases, why we age and die.

We suggest that although DNA damage is crucial to shift homeostasis, ageing itself is not caused by simple DNA damage accumulation. Instead, only specific sites of damage are relevant when they affect selection and the resulting ageing processes. For clarity, each process is given a name.

The first process, ‘celerisis’, results from the natural course of tissue-level selection for cells with elevated metabolic and proliferative rate. If the damaged DNA site gives the cell a selective advantage, it can spread within the tissue causing hyperfunctional diseases including cancer and fibrosis. However, many ageing phenotypes are more associated with *hypofunction*. Therefore, we suggest that our tissues have a mechanism to prevent the spread of hyperfunctional cells.

In proliferative tissues, a second process, ‘intrinsic ageing’, is the result of this defence mechanism induced through cell communication via Notch. Slower metabolising mutants induce epigenetic changes in faster cells, and then epigenetically slowed cells slow other cells, causing gradual metabolic slowdown across tissues.

The third process, ‘extrinsic ageing’, could then result directly from metabolic slowdown as body cells use less ATP. Mitochondria reduce catabolism, restoring ATP levels by burning less glucose and lipid. Build-up of these fuels in the cytoplasm reduces import, restoring equilibrium but inducing insulin resistance (IR), while the excess fuel is diverted to the adipose, causing weight gain, chronic inflammation, and metabolic syndrome. These outcomes could then combine with intrinsic ageing to induce age-related disease.

The final process of mitochondrial selection induces intrinsic ageing of single celled life as well as post-mitotic tissues and organisms. Together, the four processes produce a detailed mechanistic map that explains the evolutionary significance of ageing, removing old paradoxes, and connecting the hallmarks into a causal framework that furthers our understanding.

Graphical abstract

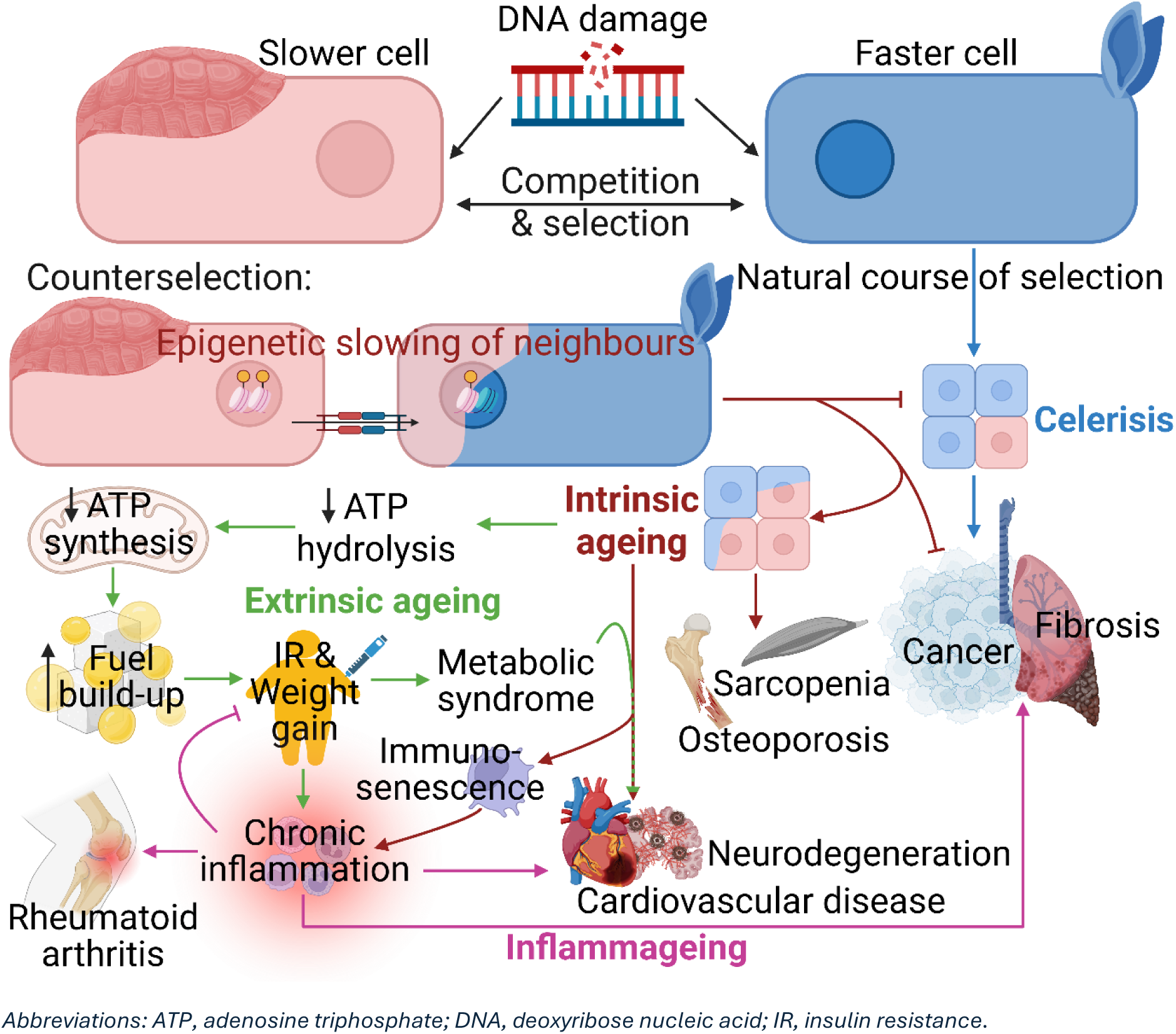

## How to read (and skip sections of) this document

The primary goal of this document is to produce a comprehensive, coherent biological mechanism for the ageing process. From the initial inducer, it is designed to indicate the basic pathway of causes and effects all the way to the ultimate inducers of disease and death. As such, it is a long document, breaking ageing into a collection of four main processes. However, gerontology is a multidisciplinary subject, and some parts can be skipped by readers mainly interested in certain aspects.

The secondary goal of this document is to demonstrate that the four process model fits with evolutionary theory and is thus evolutionarily plausible. Paragraphs in blue discuss the evolutionary implications which can be skipped without missing mechanistic insights or loss of coherence.

The tertiary goal of this document is to demonstrate the problems of the mainstream evolutionary and mechanistic theories in light of existing evidence, indicating how the four process model could solve some of the associated paradoxes. Parts 1 and 2.1 can be skipped by readers wishing to go straight to the four process model.

The document is split into nine parts each discussing a major aspect or process of ageing. Each part ends with a summary for readers in a hurry. **Clinical and translational research scientists** mainly interested in the implications for disease could read the summary for Part 3 then Parts 4-6 only.

Part 9 is for those interested in a mathematical representation of the four process model or further biological detail of the chain of metabolic slowdown described in Part 4.

### Part summary

**Part 1:** Assesses **older models** and the complexities they produce. The major issue discussed is **homeostasis**.

**Part 2:** Suggests the **‘longevity module’** can connect the hallmarks as an alternative way of looking at ageing. Secondly it suggests a **two opposing process model of ageing** based on correlations of age-related disease.

**Part 3:** Suggests how **cell competition, communication, selection and counterselection** might **induce generalised metabolic slowdown** as well as areas of localised **hyperfunction** within the body. It is largely based on our previous model^1^, but with additional clarifications.

**Part 4:** The **chain of causation from metabolic slowdown to metabolic syndrome**. Sections 4.1-4.4 were covered in a previous modelling paper^2^. It concludes with a **three process model of ageing**.

**Part 5:** Demonstrates how **metabolic slowdown, metabolic syndrome, and hyperfunction** combine to **induce age-related disease** and mortality.

**Part 6:** Discusses the implications of the three process model for **anti-ageing therapies**.

**Part 7:** Describes the **fourth process of** ageing based on **mitochondrial selection** is occurring in unicellular organisms as well as **post-mitotic organisms and tissues**.

**Part 8:** Discusses how differences in **lifespan and ageing** might **evolve**, limitations and conclusions of the four process model. It also briefly describes translation of the ideas into mathematical models, discussed further in the supplementary.

**Part 9:** Mathematical model of metabolic slowdown.

### Glossary

## Introduction

That ‘ageing is a complex, multifactorial process’ is an almost ubiquitously employed introduction in current geroscience literature. Yet the most informative aspect of this statement is the lack of mechanistic understanding it denotes. At the Global Healthspan Summit 2024, scientists expressed the view that ‘an understanding of aging in terms of basic principles is currently so unfeasible as to be unrealistic to pursue’^3^. As a result, the field has re-focussed on interventions aimed at age-associated pathologies without mechanistic connection to the ultimate cause. However, this pessimism was not always the case. In the 1970s it was widely believed that the mitochondrial free radical theory (MFRT) and the disposable soma theory (DST) would provide the mechanistic and evolutionary explanation for ageing, which could be slowed, stopped, or even reversed by the prevention and removal of DNA damage. The idea that ageing would reflect an accumulation of errors in the DNA was intuitive, quickly capturing the interest and imagination of scientists. It was over the ensuing decades, as the evidence proved inconsistent with these ideas, that scientists were forced to introduce significant mechanistic complexity to stay consistent with these hypotheses.

The mechanism described here suggests refinements of the damage theory designed to combine its crucial aspects with those of other major theories into a relatively simple sequence of events that leads to organismal decline and death. Using this unifying model, ageing once again becomes a disease we can understand, assess, and potentially cure.

We do not ask readers to accept every detail. The mechanisms span multiple disciplines, necessitating oversimplifications. However, we suggest that even with these inevitable imperfections, the four process model and its causal framework provides a superior null hypothesis to the undefined ‘complex multi-factorial process’ which currently dominates.

## Part 1: Redefining ageing

### 1.1 Problems with the damage accumulation model of ageing

Earlier ideas that ageing was inevitable ‘wear-and-tear’ were rejected because living organisms are open systems, capable of repairing themselves by extracting energy from their environment, thus lacking a thermodynamic imperative for ageing^4^. The identification of reactive oxygen species (ROS) produced by mitochondrial respiration suggested that the body’s own metabolism could produce unavoidable damage as a byproduct, but maintenance and repair could maintain functionality and thus life indefinitely.

Disposable soma theory predicted that organisms might evolve to accumulate damage because investing fewer resources in somatic maintenance would allow them to invest those resources in processes more essential to propagating their genetic line, such as growth and reproduction^5,6^. As both ageing and negligibly senescent organisms would still die from extrinsic causes (independent of intrinsic decline eg. cold, disease, and predation), the energy saved from ageing would provide a fitness advantage in the form of more (or more robust) offspring.

However, in the following decades, many studies showed that ROS were not unavoidable byproducts of cellular metabolism but crucial signalling molecules^7–9^. Human trials of antioxidant supplementation also showed either no benefit or increased mortality^10^. In the mitochondrial DNA, where oxidative damage should be higher than nuclear, several studies suggested that robust mechanisms existed to remove oxidative lesions^11,12^, and mutations arising from oxidative damage did not consistently increase with age^13^. The mounting counterevidence culminated in multiple papers suggesting the MFRT was no longer supported^14,15^. Many advocates of ageing via damage accumulation turned to alternative damaging agents, summarised in Figure 1A. In their proposed ‘Green theory’, Gems and McElwee^16^ discussed xenobiotics (e.g. toxins, drugs, carcinogens) among other potential sources, but others suggested such exogenous compounds would be unlikely to produce the same levels of damage as intrinsic sources such as our own metabolism^17,18^.

**Figure 1.**
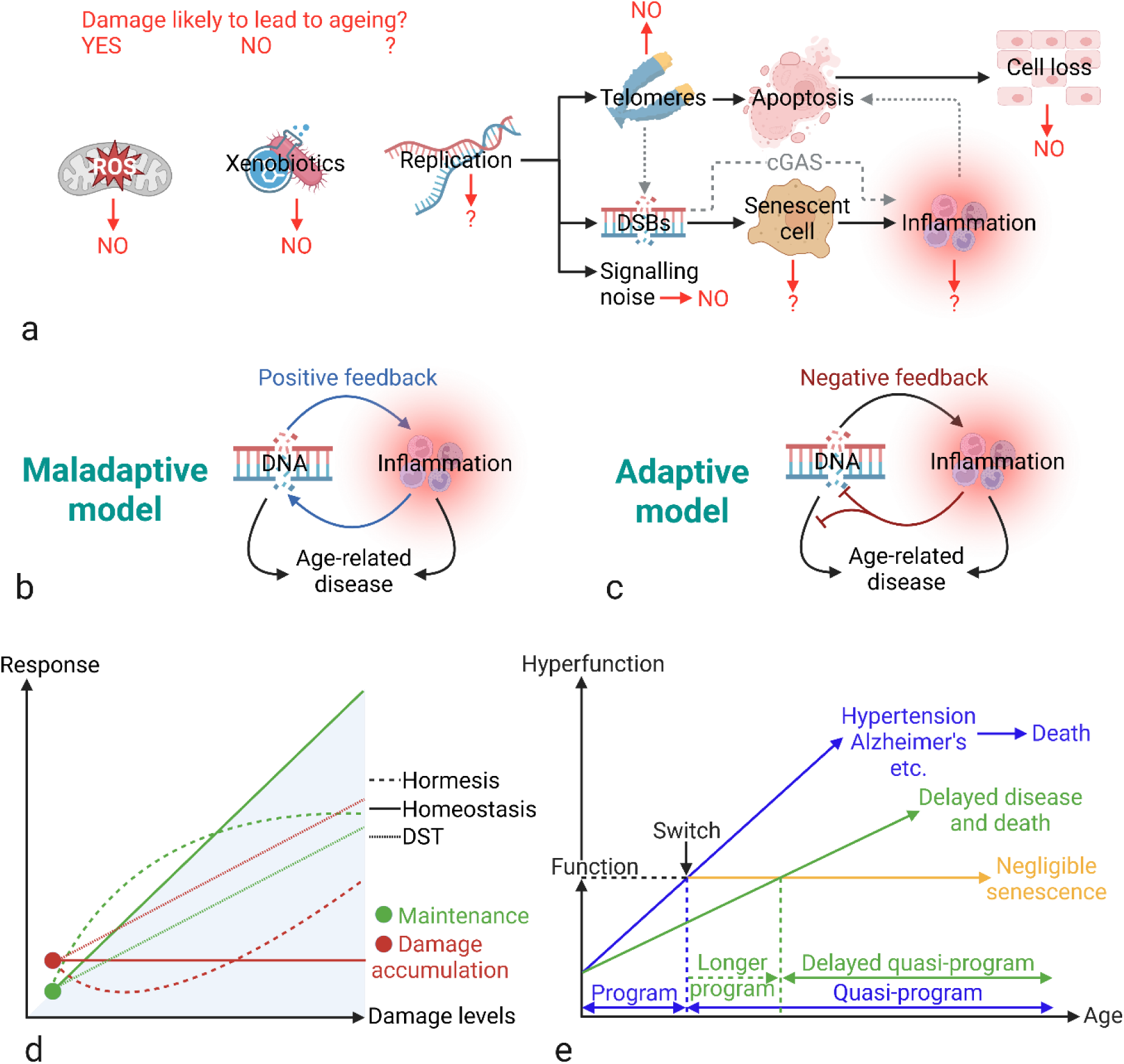
Hypotheses and models of ageing. (A) Different types of cellular and molecular damage hypothesised in ageing. In red, ‘YES’ reflects significant evidence for a causal role in ageing; ‘NO’ reflects significant counterevidence against a role in ageing; and ‘?’ reflects a plausible role with good correlative evidence and some causal evidence of contribution to lifespan or healthspan. (B) The standard maladaptive model of inflammageing where inflammation is an unavoidable and non-functional byproduct of damage, responding to damage by both increasing damage and inducing disease. (C) The adaptive model of inflammageing where inflammation reduces the damage that stimulated it, but also contributes to disease in other ways. (D) Predicted response in maintenance upregulation during exposure to a damaging agent and the resultant level of damage accumulation. In homeostasis, increases in damage are matched by increases in maintenance, so damage accumulation remains constant; in hormesis an over-response in maintenance results in reduced damage accumulation when damage induction increases, while DST predicts that increased damage induction should result in a less than homeostatic response and additional accumulation as some of the energy needed to block the extra accumulation is still needed for other processes. (E) TOR-centric quasi-program. In blue, the functional developmental program becomes a detrimental quasi-program after development ceases. As shown in green, it can be slowed, taking longer to induce age-related disease at the cost of longer development (consistent with antagonistic pleiotropy), but in yellow it can also be switched off regardless of speed producing negligible senescence (more consistent with mutation accumulation theory). Image created with BioRender.com. Abbreviations: DNA, deoxyribose nucleic acid; DSB, double strand break; DST, disposable soma theory.

If oxidative damage was not a significant inducer of ageing, then damage from DNA replication was the most plausible^13^. The initial hopes for telomeres reflected Olovnikov’s observation in 1971 that the known enzymes of DNA replication could not replicate the chromosome ends, thus causing an ‘end-replication problem’ that would cause the gradual shortening of the DNA with each cell division^19^. The subsequent discovery of telomerase could have largely removed their significance, except for the joint observations that the enzyme was deactivated in some tissues in larger organisms^20^, and cells had a finite replicative lifespan in culture^21^, which was due to telomere attrition^22^.

Subsequent studies, reviews, and meta-analyses suggested a weak correlation between telomere length and human ageing^23–25^; however, studies in mice also suggested that telomerase remained active in most tissues and telomeres did not significantly shorten with age^26^, while in the Leach’s storm petrel older birds had both longer telomeres and less variation in telomere length than in hatchlings^27^. Most inconsistently, when telomerase was knocked out in mice, it took at least three generations (G3) before mice began to show segmental progeroid syndromes, with G3 mice still demonstrating normal lifespan and G6 mice still viable. In no generation did the mice show ‘generalized premature ageing’, with cancer and alopecia and delayed wound healing reflecting the major disease increases^28,29^. Thus, many rejected telomere attrition as a significant mechanism in ageing^30^.

Notably, telomere attrition was just one kind of replication damage, and stalled replication forks, aberrant recombination, and even mechanical factors could all induce strand breaks, point mutations and chromosomal rearrangements capable of negatively impacting cell function. The discovery that telomere attrition led to cell senescence, a state of permanent arrest in cell culture^22,31^, was followed by the observation that high levels of damage at other DNA sites could also overwhelm the repair machinery and induce the same phenotype^32^.

Initial results showed senescent cells accumulated with age in animals^33^, and their removal could extend median, if not maximum, lifespan, reducing age-related disease^34^. Subsequent studies produced an inconsistent picture of the therapeutic potential of senolytics, from no lifespan benefits at all^35^ to significant maximum lifespan extension^36^, albeit with suggestions of additional toxicity^37^. The complex picture suggested that senescence had some significance in age-related decline (see Part 7.2), but most likely as one factor among many^38,39^. If ageing was caused by DNA damage during replication, then it was inducing effects in addition to, and independent of, cell senescence.

However, the mechanism by which DNA damage accumulation might induce ageing was problematic. The noise hypothesis suggested that damage could reduce the functionality of cells in random and multi-directional ways, increasing entropy within tissues^40^. While noise does increase with age, and is likely detrimental to tissue function^41,42^, there are good reasons to believe it may not cause ageing or age-related disease. Firstly, within and even between species, most morbidity and mortality occurs through a narrow range of age-related diseases more consistent with homeostatic shifts than random effects determined by whichever genes are damaged. Secondly, as many gene knockouts have recessive effects masked by another functional gene, modelling studies demonstrated that the mutation rate required to cause ageing by hitting two copies of the same gene in a significant number of cells was unrealistically high^43^. Equally inconsistent, haploid and diploid wasps exposed to X-irradiation showed no difference in rates of ageing, despite the haploids lacking the protection of another unmutated allele^44^. This left proponents of noise with the weaker argument that ageing could reflect the accumulation of mainly dominant mutations^45,46^.

However, noise is not the only effect of accumulating DNA damage. Regardless of what gene was damaged or what function obscured, damage would activate the DNA damage response (DDR). The DDR, in turn, could activate a series of pro-inflammatory pathways, such as cGAS-STING, as well as inducing cell senescence and the senescence-associated secretory phenotype (SASP). As stated by Giunta, et al.^47^, “reactive oxygen species (ROS) cause amplification of cytokine release, fuelling a self-perpetuating positive feedback loop. The end result of this cycle is a chronic and systemic pro-inflammatory state”. This is the inflammageing hypothesis^48^, reflecting that ageing and age-related disease are both so consistently associated with inflammation^49^. The resulting vicious cycle of damage, cell death, and inflammation could therefore induce a directional shift in homeostasis toward chronic inflammation and hypocellularity, independent of increasing noise^50,51^.

However, while cell loss occurs in some post-mitotic tissues, most mitotic tissues show no overt cell loss, matching the rate of replication with cell death^52^, some conversely showing significant cell expansion^53,54^. The ageing haematopoietic stem cell niche shows increasing cellularity in humans and mice, and higher proliferative potential upon transplant^55^. Indeed, more recent reports suggest that ageing cells can be more resistant to apoptosis^56–58^, and post-mitotic organisms like C elegans appear to retain their cellularity during ageing^59^.

Additionally, the positive feedback loop between damage and inflammation created several complexities. Studies showed juxtacrine and paracrine effects of senescent cells, which within the high concentrations of in vitro cultures, could spread damage and senescence to other cells^60–62^, consistent with the maladaptive model of positive feedback shown in Figure 1B. However, the same mechanism occurring at the much lower in vivo concentrations would lie fundamentally at odds with homeostasis. Damage response pathways that caused age-related disease by spreading damage and senescence would be the evolutionary equivalent of repeatedly hiring the same ‘cleaners’ despite their consistent tendency to make more mess.

Conversely, there is an adaptive model (Figure 1C), where damage and senescent cells could accumulate *despite* inflammation either because we do not invest sufficiently in maintenance or as a result of other trade-offs in the process of identifying and removing senescent cells^63^. Using this model, inflammation rises as a necessary negative feedback pathway to remove the increased damage and senescent cells, with other effects that induce disease. However, even this model would predict that significantly increasing DNA damage should shorten lifespan.

Contrarily, studies showed that antioxidant mutations in C elegans increased oxidative damage levels with positive or no effect on lifespan^64,65^, as was true for chronic γ and X-irradiation in mice and Guinea pigs^44,66^.

The paradox is conventionally explained by the hormesis hypothesis, whereby low levels of damage over-stimulate the stress response, reducing the total level of damage. Thus, whenever inducing damage is paradoxically beneficial, it reflects the increased capacity of the body to withstand further damage.

In the absence of studies showing that net DNA damage is actually reduced, the main evidence for hormesis is the role of the longevity regulator DAF-16/FOXO in upregulating the stress response, including multiple antioxidant enzymes. After initial experiments demonstrating that the longevity gained by daf-2 mutants was dependent on daf-16^67^, a small percentage (10-20%) of the extended daf-2 mutant lifespan was found to be lost by the additional knockdown of stress response genes, including antioxidant enzymes mediated by DAF-16/FOXO^68^.

According to the accumulation-based model, damage was activating FOXO which prevented additional damage, hence hormesis. However, multiple studies suggest that stress resistance is neither sufficient nor necessary for longevity or FOXO-induced longevity specifically. For example, glutathione peroxidase 4 (Gpx4) heterozygous knockout mice have increased lifespan, despite increased markers of oxidative damage, because their cells have increased sensitivity to stress-induced apoptosis^69^. Other antioxidant mutants also show increased oxidative stress and even oxidative DNA and protein damage, without affecting lifespan, further uncoupling the association^70^. One study looking specifically at long-lived daf-2 mutants with additional knockouts of superoxide dismutase (sod) 1-4 genes showed the latter had no effect on the daf-2 mediated lifespan increase, except loss of sod-4 which further extended it. In all cases, except sod-4, the antioxidant knockouts significantly increased stress sensitivity to either paraquat or hyperoxia or both^71^.

Notably, the over-stimulation of the stress response by damage fits uneasily with DST, which is the damage accumulation model’s main evolutionary underpinning. The central predictions of DST, that organisms age because they save energy by reducing investment in maintenance, is contradicted if the body hyperactivates the stress response when exposed to low-level damage. Even small increases in stress might be predicted to result in additional damage and reduced lifespan (Figure 1D) if there are more important processes requiring that energy. Most discussion of the consistency of hormesis and DST concerns calorie restriction (CR), where Shanley and Kirkwood^72^ argued persuasively that the reduced availability of resources would shift the priority of organisms from offspring production to survival. As offspring have fewer reserves, they are less likely to survive periods of starvation, cold, and disease which would likely accompany a calorie restricted environment. An organism that could pivot investment away from offspring to ensuring its own survival would be less likely to waste its resources, gaining a fitness advantage. While Speakman^73^ argued against this idea, the most notable inconsistency was argued by Mitteldorf^74^, that only mid and low-calorie diets would necessitate resource allocation – mid-calorie diets allocating to offspring and low calorie to survival – while high-calorie diets should provide enough energy for both maximal reproduction and longevity, thus increasing lifespan. A U-shaped curve for investment in maintenance would be the optimal evolution^74^, but resource excess is still associated with reduced lifespan.

Even if DST can explain CR, there are other examples of hormesis which are less consistent. People living in areas of elevated background radiation have shown reduced incidences of cancers consistent with hormesis^75^, but such conditions are not associated with the same shift in selective advantage from offspring to maintenance as during CR. Yet higher levels of damage still promote an over-response. Thus, while hormesis explains the paradox of damage sources failing to affect lifespan, it creates a new paradox in its place. Without the energy saving measures proposed by DST, damage lacks an evolutionary reason to accumulate in open systems. It was yet further complexity to understand.

### 1.2 Problems with other major models of ageing

Those unwilling to accept the additional complexities associated with damage accumulation returned to the older idea that ageing is programmed, the dominant hypothesis being Mikhail Blagosklonny’s TOR-centric quasi-programme (TCQP)^76^, that ageing is a **beneficial developmental programme which is left on after organisms reach adulthood despite losing its utility once development is complete, thus becoming a detrimental quasi-programme**.

Previously, Kirkwood had argued persuasively that cheaters would outcompete any organisms adopting self-sacrificial ageing programmes^6^, and although group selection theories do have advocates^77,78^, they have largely been discarded by mainstream evolutionary theory^79^. However, the TCQP circumvented the issues of cheaters because its purpose was a developmental programme that was beneficial to the individual rather than the group. Blagosklonny described the programme with the analogy of a car with no breaks – the faster an organism could reach maturity, and the bigger and stronger it was during early adulthood, the more competitive it would be early in life, but also the faster its quasi-programme would drive accelerated ageing (Figure 1E, blue and green lines) and the speeding car would crash^80^. Blagosklonny suggested that the TCQP was an example of antagonistic pleiotropy (AP)^81^ because it provided an early life advantage and a pleiotropic late-life cost^82^.

However, we would argue that while the rate of the developmental programme (and thus subsequent quasi-programme) might be antagonistically pleiotropic, this is only true if evolution is incapable of turning the programme off. Just as a car with functional brakes can accelerate faster and stop just as quickly by applying the breaks more strongly, organisms that could accelerate development to maturity, then turn the programme off would gain the benefits without the costs (see Figure 1E). To put it another way: if the off switch does not evolve, then this reflects genetic drift as organisms gain no advantage for not turning the programme off once its utility is complete. The TCQP and any developmental theory are therefore more consistent with mutation accumulation theory (MAT) than AP.

MAT suggested that mutations could arise inducing late-acting detrimental effects like ageing that could not be removed from the gene pool by selection because few organisms survived long enough to experience the detrimental effects^83^: most were killed, before the manifestations of ageing, due to extrinsic factors independent of their rate of intrinsic decline. In addition to quasi-programmes, damage accumulation could evolve and spread by drift if it occurred at a rate that did not affect wild animals; however, we now know that many organisms experience significant age-related decline in the wild, suggesting **ageing should offer a fitness advantage**^84^. A ‘selection shadow’, reflecting that detrimental effects happening later will have smaller fitness costs, has been proposed to produce ageing in the wild^85^, as many animals die of independent extrinsic factors which ‘shadow’ or reduce the selective pressures to reduce the rate of ageing. However, as long as significant numbers are experiencing age-related decline in the wild, then the selection shadow will not reduce costs entirely^86^. To explain ageing, it would therefore either still require some advantage even in the quasi-programmatic stage, or significant complexity in turning the programme off that makes selection more difficult. We suggest that a selection shadow is not sufficient, in itself, to explain why a developmental program fails to turn off despite its fitness detriment.

Indeed, a primary prediction of MAT or the selection shadow is that ageing should be relatively implastic, but in social insects worker lifespans can differ from those of their queens by two orders of magnitude^87^. Therefore, we suggest that the two dominant theories, DST and TCQP, which combine an evolutionary foundation with a predicted mechanism are unlikely to provide full explanations for ageing.

Aside from a focus around mTOR, there is little evidence as to the mechanism by which an mTOR-centric developmental programme might lead to our decline. Since Blagosklonny’s initial suggestions of ‘hyperfunction’ and over-stimulated growth, which are consistent with some age-related diseases and phenotypes, there has been suggestion that others more resemble the opposite – a hypofunctional programme^88^ – with little description of the evident complexities that would allow both hyperfunction and hypofunction to result from a single programme.

### 1.3 The significance of homeostasis

Another problem for the TCQP and any quasi-programme theory is the central importance of negative feedback in ensuring that programmes like the TOR-centric programme do not outlive their utility and become quasi-programmes. The default setting of any cell, tissue, and organ system is to retain metabolic homeostasis. By this logic, unless there is some functional purpose, or inducer of dysfunction, a developmental program is as likely to be left on indefinitely as, for example, the pancreas is to continue producing insulin once blood glucose has returned to equilibrium. The size, activity, and growth signals within our organs must be under strict regulation, as deviations result in disease and death. The body’s molecular and cellular programs are goal-directed enforcers of this homeostasis, even if some, like inflammation, shift tissues away from equilibrium to restore it. Development is designed to grow our organs and activate new pathways at precise timings and to exact amounts. While homeostasis is not infallible, it should contain built-in negative feedback systems to turn off the mTOR-centric program automatically when development is complete.

Homeostasis creates a similar problem for DST because an organism reducing investment in a maintenance pathway such as the antioxidant defence cannot simply divert that energy to reproduction. If the additional ROS that result from reduced detoxification cause damage to the DNA or proteins, then the cell will naturally upregulate repair mechanisms in response. Even if cells allowed the damage to accumulate, it would produce dysfunctional cells with misfolded proteins doing their job less efficiently than before. At each point that a cell attempts to redirect resources away from maintenance, it is creating increasing inefficiency and requiring more energy to achieve the same result. If the damage is severe, the cell must apoptose and be replaced, which would reflect a considerable loss of energy (Figure 2). While some proponents of damage accumulation suggest that cell loss is an important part of ageing, this is mainly thought to reflect cellular exhaustion from telomeric attrition or overwhelming DNA damage, not because tissues are diverting resources away from cell proliferation to other processes. Thus, even in the short-term, it is plausible that diverting energy from cellular maintenance will not save any energy for investment in other processes.

**Figure 2.**
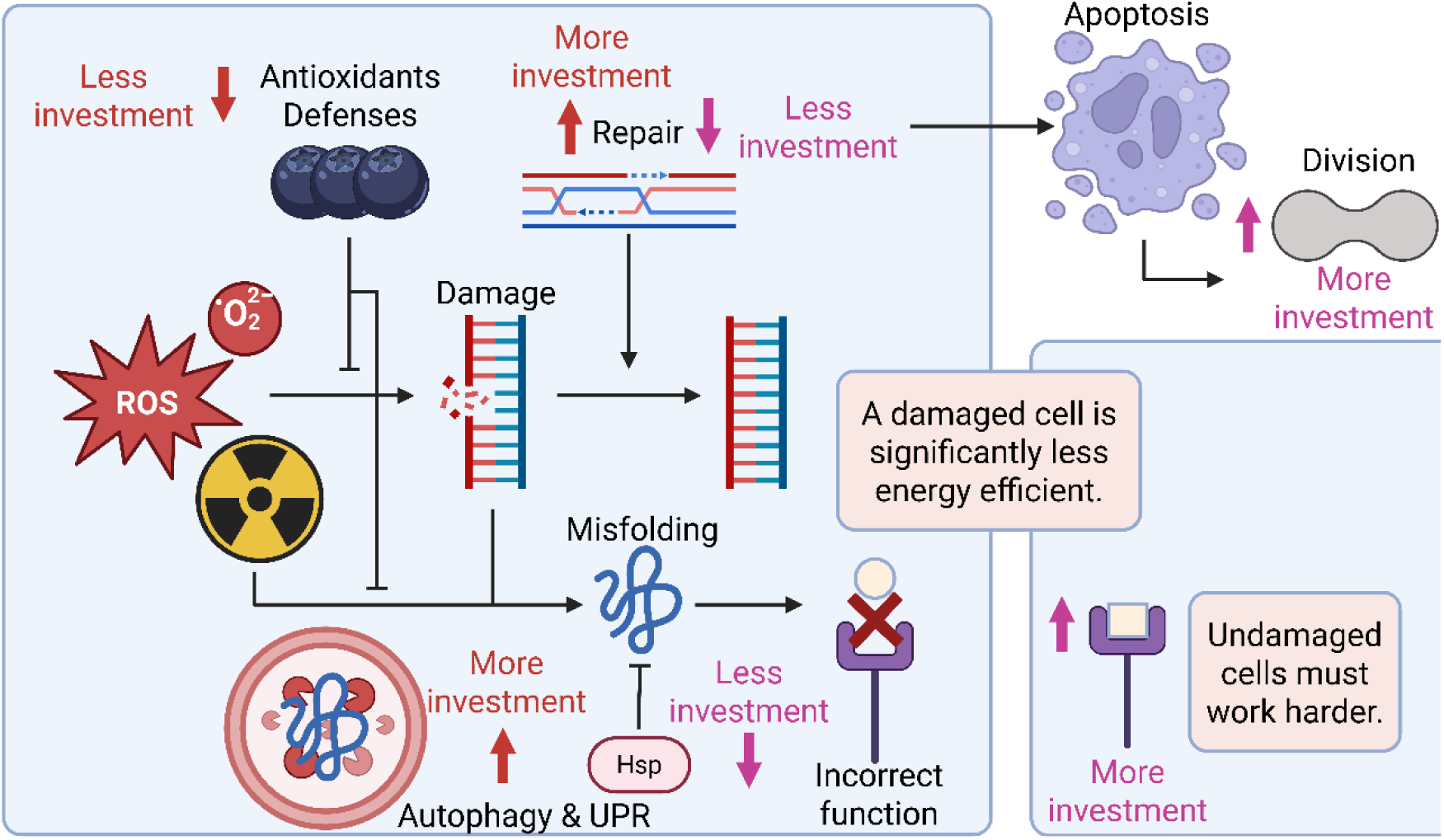
The homeostatic problem with lowering maintenance to save energy. Reducing antioxidant defence results in more DNA and protein damage, requiring more energy to correct the damage by DNA repair and protein degradation/replacement. If insufficient energy is invested in these processes, cells function incorrectly, and thus less efficiently, requiring other functional cells to do additional work to do their job, wasting energy as opposed to saving it. If the avoidable damage is severe, it will require apoptosis and replacement by a new functioning cell, also requiring more energy. At each point, if damage is not corrected, efficiency and energy are wasted, requiring a more energy intensive process to belatedly put the damage right. The final step is not subject to under-investment as undamaged cells will do the work of the damaged cell to maintain homeostasis. Image created with BioRender.com. Abbreviations: Hsp, heat shock protein; ROS, reactive oxygen species; UPR, unfolded protein response.

### 1.4 From mechanisms to hallmarks

While ageing could still result from other antagonistically pleiotropic processes, including those centred around damage, the mechanisms connecting damage to ageing phenotypes and diseases are still a subject of controversy, with some asking whether damage accumulation theory is still supported by the literature^89,90^. A recent review of the potential for computational models of ageing highlights this issue by not identifying a single model that demonstrates a connection between initial cause and homeostatic shifts that cause ageing and disease^91^.

Another review referred to “ageing-themed models”^92^, detailing many excellent models looking at particular aspects within gerontology, such as telomere shortening or loss of protein homeostasis, but without demonstrating any models that looked at the mechanisms of ageing itself. Indeed, the models of ageing as a process tend to assess the proposed inducers of ageing, such as in the recent senescence-based models by Kowald and Kirkwood^38^ and Karin, et al.^93^, where the inducer is both initial cause and ultimate effector of mortality, ie. mortality is directly proportionate to the accumulation of senescent cells. Although the senescence associated secretory phenotype (SASP) is implicated in chronic inflammation, this is far from a comprehensive mechanism of homeostatic changes describing how senescent cells lead to death. Indeed, fitting the mechanistic details for ageing into a workable model faces many unknowns and inconsistencies that create a picture of prohibitive complexity.

The result was that in 2013, Lopez-Otin et al. produced the hallmarks of ageing – a list of features associated with ageing, if not necessarily causal^39^. The previous publication of the hallmarks of cancer^94^ had been of significant utility to oncology as each hallmark reflected part of a causal pathway explaining the processes of transformation and metastasis. However, the hallmarks of ageing offered no comparable causal framework^95,96^. Some, such as telomeres, appeared only tangentially related to the ageing process, while the others had little obvious connection between each other, an initial cause (or causes), or age-related diseases as the ultimate effectors of mortality.

Mechanistic study then focussed on connecting the hallmarks to age-related disease^97,98^ rather than connecting DNA damage as the initial cause to the hallmarks of ageing. Indeed, ‘genomic instability’ is listed as a single hallmark among many, which notably associated less well with age-related diseases than other hallmarks^98^. In our opinion, the reliance upon the hallmarks without attempting to understand the mechanisms underlying them has generated only further complexity. The null hypothesis is that ageing occurs by a process too complex to fully understand, and interpreting new results in this light can only reinforce that the hypothesis is correct. No additional outcome could undo all the existing complexity and reject it.

It is therefore the purpose of this paper to outline an alternative null hypothesis which connects the hallmarks. We provide here a comprehensive mechanism of ageing from initial cause all the way to the main age-related diseases, including evolutionary foundations that demonstrate why organisms might evolve to age via this mechanism. Although multiple aspects are still theoretical, we suggest it explains many major geroscience observations and provides a causal scaffold beneath the hallmarks to compare new observations against. Of course, not every proposed mechanism will be correct, and those that are not can be replaced as new empirical data demonstrates requirement, but the use of this model should at least allow us to grow our understanding of ageing without simply furthering the complexity with which we understand the ageing process to involve.

## Part 1: Summary

1.1 | Ageing was initially viewed as simple. Damage accumulation was the underlying cause, driven primarily by mitochondrial ROS.

Damage would accumulate to allow organisms to spend less of their limited resources on somatic maintenance and more on reproduction, making them more evolutionarily successful.

However, the ensuing evidence suggested that ROS did not cause ageing. The only type of damage that could feasibly accumulate enough mutations to cause ageing was through replication, though less likely through telomere attrition.

Three main outcomes by which replicative damage might cause ageing are:

- Noise
- Cell loss
- Inflammation

Noise is unlikely to cause ageing. Cell loss may contribute to ageing in some post-mitotic tissues but most proliferative tissues show little or no cell loss, leaving inflammation and the SASP as the major plausible mechanisms by which replicative damage may promote ageing.

Even if replicative damage is the major natural source of ageing damage, this does not explain why increasing other types of damage either has no effect or extends longevity.

This is explained by hormesis; however, FOXO can still induce longevity in animals more sensitive to damage.

1.2 | Many scientists now suggest that ageing is a developmental program which remains active (as a non-functional quasi-program) even when it becomes detrimental.

However, such programs are examples of genetic drift as they offer no evolutionary advantage. Even in the presence of a selection shadow, they are unlikely to explain ageing in wild populations where it induces fitness costs.

Development is a hyperfunctional process – inducing growth – and can explain hyperfunctional diseases, but struggles to explain the many ageing phenotypes which are attributable to hypofunction.

1.3 | Neither developmental programs nor reducing investment in maintenance are wholly compatible with homeostasis as causes of ageing.

1.4 | The many results incompatible with ageing via damage accumulation or developmental programs have created a picture of great complexity to be understood as a collection of hallmarks rather than mechanisms.

## Part 2: From hallmarks back to mechanistic process

2.1 | The longevity module

When empirical study disconnects from the damage accumulation hypothesis, it repeatedly suggests that ageing and longevity can be impacted through interventions and behaviours that affect a central network of closely linked pathways, shown in Figure 3A (green box). López-Otín, et al.^99^ include its components nutrient sensing, mitochondrial dynamics, proteostasis, and autophagy as separate hallmarks, but when put together they form the energy/metabolism regulation network. The age-1, daf-2, and daf-16 mutant C. elegans, which stimulated the search for longevity genes, all form part of the insulin-like growth factor (IGF) pathway^100^.

**Figure 3.**
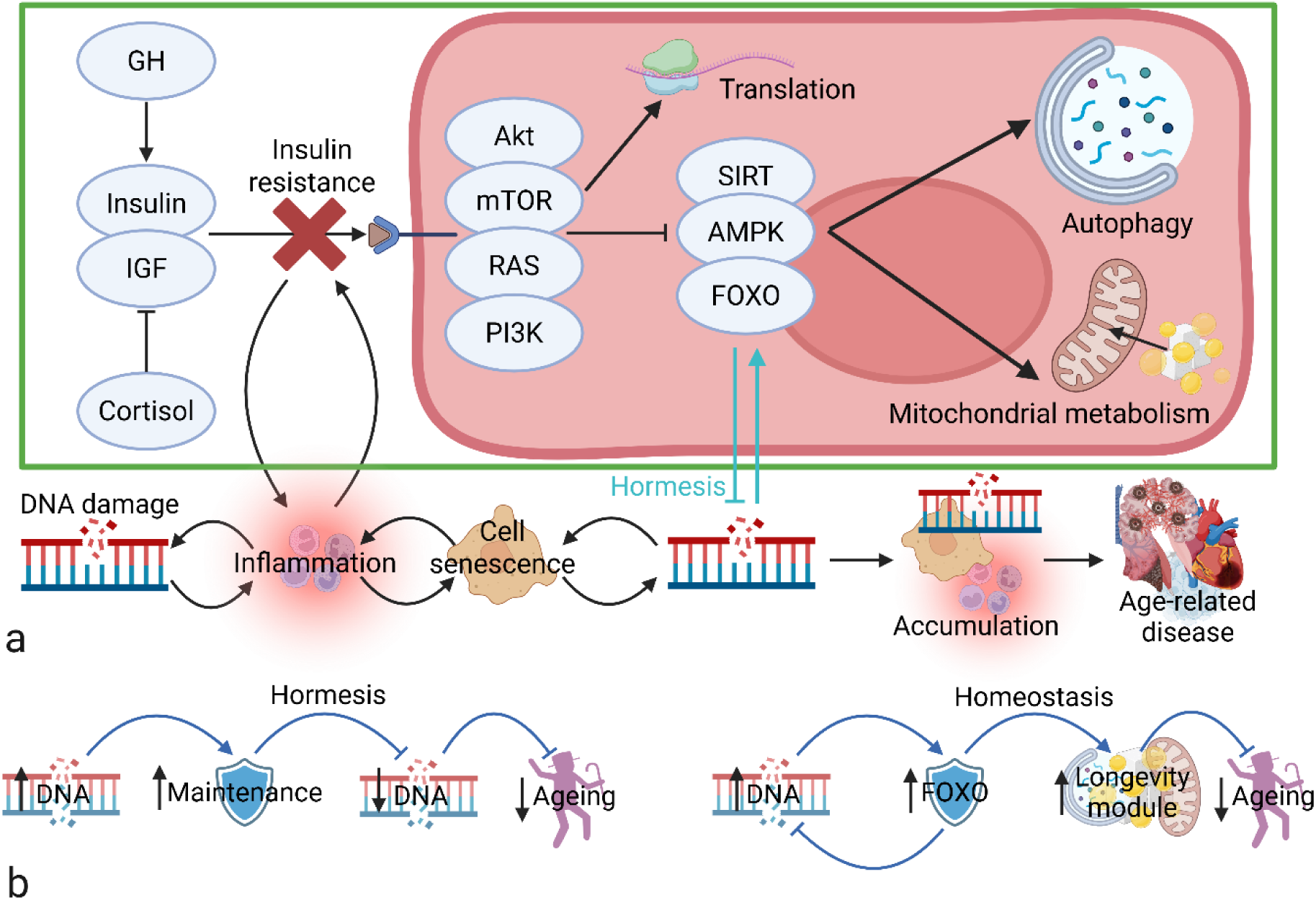
The longevity module and connections with damage and inflammation. (A) The longevity module in the green box reflects the major molecules and pathways known to be associated with ageing and lifespan modulation, all of which are involved in energy regulation. Turquoise arrows suggest why DNA damage can increase lifespan and lead to the phenomenon of hormesis. The sequence outside the green box demonstrates the inflammageing hypothesis whereby damage leads to inflammation, which both causes disease and interacts with the longevity module through insulin resistance. (B) DNA damage slows ageing either through preventing further damage in the hormesis hypothesis (left), or by shifting the homeostasis of the longevity module (right). Image created with BioRender.com. Abbreviations: AMPK, AMP kinase; DNA, deoxyribose nucleic acid; FOXO, forkhead box, sub-group O; GH, growth hormone; IGF, insulin-like growth factor; mTOR, mammalian target of rapamycin; SIRT, sirtuin 1.

Calorie restriction, rapamycin, and metformin all impact mTOR activity, which acts downstream of IGF to induce its pro-growth effects^101^. Long-lived Ames and Snell dwarf mice had mutations in growth hormone (GH), the primary inducer of IGF production, while the long-lived FIRKO mice are insulin receptor knockouts, reducing the insulin and IGF signal through mTOR^100^. Autophagy, mitochondrial biogenesis, and ATP production are inextricably linked with longevity^102^, with associated mutants impacting maximum lifespan and age-related diseases, and are all regulated by FOXO and mTOR^103^. While this network is well known, it could be described as the ‘**longevity module**’.

Modularity is the concept that systems can be broken into subsystems (or modules) defined by relative independence from each other; there is high interconnectivity within modules, but low interconnectivity between the components of different modules. In biology, specific processes frequently have their own modules to maintain evolutionary plasticity. Without modularity, a beneficial mutation in one system would be more likely to have detrimental effects in others, and less likely to undergo positive selection^104^. Evolution would be slower, and non-modular systems less competitive in changing environments. The existence of a longevity module should thus allow organisms to better adapt their lifespan to environmental changes such as shifting extrinsic mortality.

A longevity module poses challenges for theories like MAT and AP, at least in their initial forms which described ageing as the result specifically of accumulated gene mutations in multiple pathways and processes, each with small effects. Recently Gems and Kern^105^ suggested ageing could reflect multiple antagonistically pleiotropic quasi-programmes including developmental and late-acting ones. However, ageing as a multi-process, multi-pathway phenomenon is difficult to reconcile with both modularity and plasticity. Indeed, social insects which demonstrate plasticity in orders of magnitude in their rates of ageing between related individuals do appear to regulate these deviations through insulin/IGF and mTOR signalling^87^.

While many mutations and processes may have minor impacts on ageing and mortality, the major regulators appear to lie within the longevity module, controlling the energy and nutrient network – rather than DNA maintenance as predicted by ageing via damage accumulation.

The most obvious connection between DNA maintenance and the longevity module is through FOXO (see Figure 3A, turquoise), which is activated by DNA damage and essential for several stress response pathways^106,107^. Yet, by this connection, DNA damage should increase lifespan by upregulating the longevity module, contrary to damage accumulation predictions, but explaining both the long-lived antioxidant knockout mutants^64,71^ as well as hormesis.

Notably, despite the significance of Occam’s Razor to the DNA damage accumulation hypothesis, and its intuitive advantage based on our everyday experience of wear-and-tear, hormesis is arguably not the simplest explanation for the effects of FOXO. While the FOXO family are stress response genes, they are also regulators of energy homeostasis, upregulating autophagy and mitochondrial biogenesis and modulating Kreb’s cycle and oxidative phosphorylation^108,109^. A simpler explanation for the beneficial effects of DNA damage than an over-response which prevents greater levels of damage is a homeostatic response which activates longevity-inducing shifts in energy regulation (see Figure 3B).

By this mechanism, FOXO will also activate systems to reduce the stress that activated it^68,107^, preventing further damage as in standard negative feedback, but the lifespan extension will be independent of whether total damage is increased or decreased, modulated by the effects of FOXO on energy homeostasis and the longevity module.

Although there may be many contributing factors for why the hallmarks have dominated over the longevity module, the most critical reason may be the difficulty in connecting the module with DNA damage accumulation. Outside FOXO and hormesis, which reverse the predicted effect, the most widely accepted link is that inflammation is an inducer of insulin resistance (IR), such as via JNK-mediated inhibition of the insulin receptor substrates^110^, which connects to the IGF and mTOR pathways, and thus the longevity module. IR then induces more inflammation, such as through mTORC2 inhibition resulting in increased polarisation to inflammatory M1 macrophages through MCP-1^111^. This creates a positive feedback loop driving further inflammation^112^. A mechanistic assessment of this relationship would look something like Figure 3A (total).

However, the inflammageing model still limits the entire longevity module to a positive feedback loop with IR, with inflammation as the central inducer of age-related disease. It strongly predicts that the inflammation and maintenance should form the central longevity module. Additionally calorie restricted mice are more sensitive to radiation exposure rather than less, despite increasing insulin sensitivity^113^. While we will return to inflammageing in Part 5, significant complexity is required to overcome these inconsistencies.

2.2 | The two opposing process model of ageing

Perhaps rather than a complex multifactorial process, ageing is not a single process at all. The hallmarks and age-related diseases that we have assumed have a single underlying cause could instead reflect a series of different processes with varying degrees of interdependency. The age-related diseases could be distinct diseases resulting from the unique dysfunctions of their specific organs and organ systems. If true, we would predict that the age-related diseases would lack common molecular markers of decline and have little correlation with each other. However, the evidence largely contradicts this.

Multimorbidity studies suggest that the occurrence of one age-related disease is strongly associated with increased risk of others, while factors such as smoking and depression increase risk of developing multi-morbid conditions^114^, suggesting at least that if ageing is multiple different processes then they are highly connected and interdependent.

Without an underlying hypothesis it is difficult to tell which factors are significant and early drivers of disease mechanisms, and which are late actors that could be otherwise avoidable. This paper hypothesises that when all the minor factors and proximate causes are stripped away, ageing is two distinct and opposing processes which produce different types of age-related diseases. While we respect that other factors and interactions could be used to promote very different ideas, here we describe evidence consistent with the two opposing process model.

In genome wide association studies (GWAS), the most significant predictor of longevity is usually from variations in the apolipoprotein E (APOE) gene, with the APOE ε4 allele associated with the shortest lifespan and APOE ε2 the longest lifespan, with APOE ε3 producing intermediate longevity^115^. Interestingly, APOE ε4 carriers are also the most prone to developing Alzheimer’s disease and cardiovascular disease^116^. Another gene variant associated with longevity is the FOXO3a “G” allele, which is mainly a predictor of reduced cardiovascular mortality^117^. However, studies also indicate a protective role of FOXO3a and the “G” allele against incidence of Alzheimer’s^118^. Thus, if APOE and FOXO3a are connected to the mechanism of both Alzheimer’s and cardiovascular disease as well as longevity, they plausibly share a common underlying process, likely ageing. APOE is a distributor of triglycerides to tissues from very low density lipoprotein (VLDL) in the blood^119^, and FOXO3a is key modulator of energy homeostasis^120^, both linking to energy regulation and the longevity module^121^ and suggesting the common mechanism is indeed ageing.

Interestingly, the FOXO3a ‘G’ allele shows little relationship with cancer mortality^117^, while the APOE ε4 allele, which is associated with the highest risk of cardiovascular disease and Alzheimer’s disease, and the shortest longevity, is associated with the *lowest* overall cancer risk^122^. Relatedly, high LDL levels, which are associated with increased risk of cardiovascular disease^123^ are also associated with reduced cancer risk^124^. Conversely, the hallmark of ‘genomic instability’ associated primarily with cancers, while showing little association to other age-related diseases^97,98^.

Diseases which primarily associate with cancer include idiopathic pulmonary fibrosis (IPF) and chronic liver disease (CLD), which is also a fibrotic disease. Prevalence of lung cancer in IPF patients ranges from 4.4-13%, with rates as high as 48% in autopsy studies^125^. Even individuals who received lung transplants, which is the only known cure for IPF, still had a twenty times higher probability of developing lung cancer than the background rate^125^. Similarly, a study looking at individuals with diagnosed lung cancer found that one in three also had fibrotic scarring, and the scarring was associated with the same lobe as the cancer^126^. In the liver, underlying cirrhosis (end-stage fibrosis) is the most important risk factor predicting development of hepatocellular carcinoma^127^. Indeed, cirrhosis-inducing conditions such as infection with hepatitis viruses can induce multi-centric cancers (ie. independently formed) within the same liver^128^.

Thus, ageing could be split into two major processes, as shown in Figure 4. While these are plausibly unnecessary neologisms to be replaced with existing terminology, to avoid confusion we have called one ‘**intrinsic ageing**’, reflecting the underlying process responsible for cardiovascular disease and Alzheimer’s disease, and the second ‘**celerisis**’, from ‘celeritas’, the Latin for speed. This reflects that it is responsible for cancer and fibrosis, which could be described as overactivity disorders or states of hyperfunction, over-producing cells and extracellular matrix, respectively.

**Figure 4.**
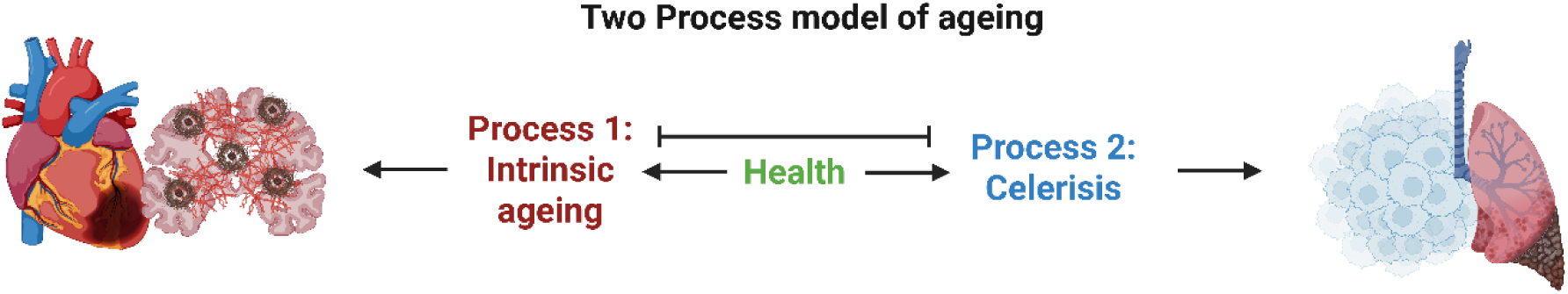
The two opposing process model of ageing which splits one complex process into two separate processes that induce different diseases. Intrinsic ageing leads to Alzheimer’s and cardiovascular disease, while celerisis leads to fibrosis and cancer. Image created with BioRender.com.

In Part 3, we summarise the observations from our previous paper, defining how a slight alteration to DNA damage accumulation theory could explain why ageing might be split into two opposing processes and why these processes evolved, potentially providing the ultimate cause of ageing^1^.

## Part 2: Summary

2.1 | Almost every gene, behaviour, and therapy which affect healthspan and lifespan are connected to the energy regulation network, which could be referred to as the **longevity module**.

Damage and maintenance have little connection to the longevity module. The major connection is via the activation of FOXO by which damage extends longevity – opposite to the predictions of damage accumulation as the cause of ageing, but explaining hormesis.

An alternative to the hormesis hypothesis, that damage extends lifespan by reducing damage, is that damage activates the longevity module through FOXO, affecting energy regulation. It shifts energy homeostasis.

Inflammageing reduces the entire longevity module to a system of positive feedback between IR and inflammation.

2.2 | Some lines of evidence suggest that age-related diseases could be split into two groups; Alzheimer’s and cardiovascular disease might result from one ageing process, while cancer and fibrosis might result from a separate one. We called these **intrinsic ageing** and **celerisis**, respectively. Together they form the **two opposing process model** of ageing.

## Part 3: Cell selection causes intrinsic ageing and celerisis

### 3.1 Mitotic tissues are sites of communication, competition, and selection

The DNA damage accumulation model has focussed overwhelmingly on mutations that do not impact selection within tissues, suggesting that ageing results in increasing noise, cell loss, and chronic basal inflammation. However, some mutations affect not only cellular function, but provide cells with traits that are advantageous compared to other cells within the same tissue. Such mutations give cells a selective advantage over wildtype cells causing them to spread within the tissue. The significance of the mutation on cell function is therefore amplified by the cell’s potential to spread. The result is not increasing noise but a shift in homeostasis toward the metabolic output of the mutant and its daughter cells. Cancer cells are a widely accepted example of selection occurring within tissues, but the importance of selection in the spread of non-cancerous mutants is also emerging^129^.

Notably, the threat of advantageous mutation may be underestimated under the assumption that cells can autonomously prevent their own replication if their DNA is damaged, thus preventing their spread. Canonical Ras genes are mutated in around 30% of human tumours^130^, but in healthy cells Ras mutation induces a brief period of hyper-proliferation before the cell either dies or permanently arrests, called oncogene-induced senescence^131^. However, autonomous mechanisms are only safe when the mutation induces a state of metabolism that is beyond physiological norms^132^. If instead of producing hyperproliferation, a ‘minor’ RAS mutation produced a smaller increase in division rate that normal cells could also reach when environmental stimuli were highly ‘pro-growth’, then having cells die or arrest when they reached that level of growth could be dangerous to the tissue: flooding the system with enough IGF or GH would cause tissues to degenerate from mass cell death (Figure 5A).

**Figure 5.**
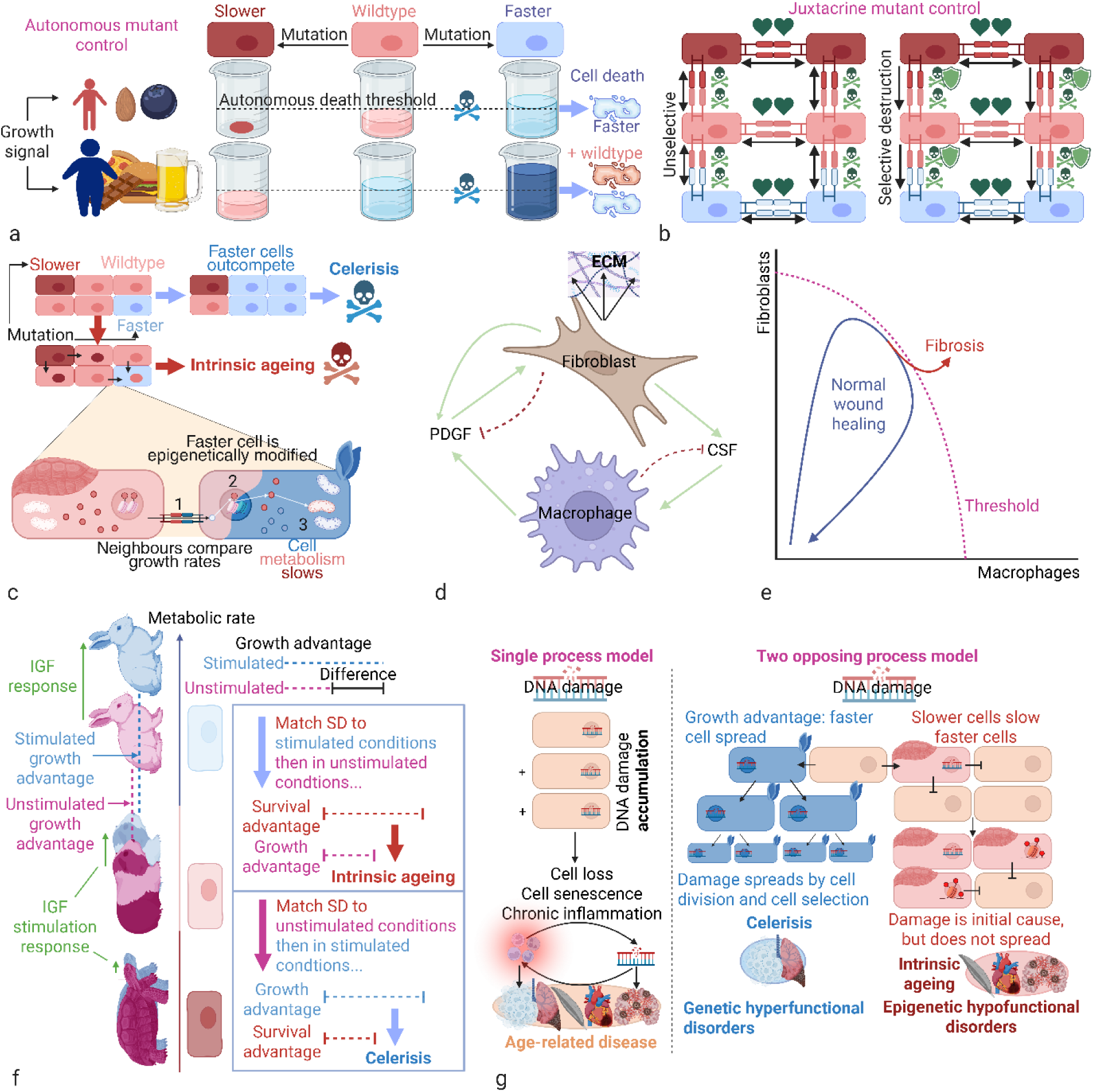
Effects of mutation and defence mechanisms. (A) Autonomous defence mechanisms against mutation cannot remove minor growth mutants because their change in growth rate cannot be distinguished from the effects of a shifting growth signal. While a low growth signal might allow the removal of only fast mutants, if the signal should increase too much, autonomous cell removal would result in apoptosis of wildtype cells responding normally to an aberrant signal. (B) Juxtacrine control of mutants by cell comparison and communication. Left, is the unselective control mechanism attempting to remove all cells which appear aberrant to their neighbours. The hearts represent a survival signal offered by similar cells, the skulls a death signal induced by cells that recognise differences in metabolic rate. Right, is a mutant control mechanism of more selective destruction, where slower cells are less likely to be killed in situations where they are identified as different, as indicated by the shields. (C) Top, shows how fast mutants will outcompete wildtype cells by their growth advantage, inducing celerisis. Bottom, slower cells reduce the metabolic rate of faster cells with semi-permanent epigenetic marks, preventing celerisis at the cost of intrinsic ageing. (D) Interaction between macrophages and fibroblasts consisting of positive and negative feedback. (E) The wound healing process as shown by Adler, et al.^133^ where fibroblasts and macrophages activate each other, closing the wound and clearing it of debris and pathogens, then deactivating once the signal dies. Or if they reach a threshold where the fibroblasts have sufficient population density to sustain the autonomous positive feedback shown in (D), the fibroblasts continue proliferating, causing fibrosis. (F) Growth mutants can have different sensitivity to growth signals. Slower mutants may be relatively insensitive to changing metabolic rate as growth signals change, whereas faster mutants may demonstrate larger changes in growth rates than wildtype cells to the same signal. If the survival advantage of slower cells from selective destruction is matched to the growth advantage of faster cells under low-growth conditions, then under high-growth conditions the increased growth advantage of faster cells will allow them to outcompete wildtype cells, producing celerisis. Conversely, if the survival advantage of slower cells from selective destruction is matched to the growth advantage of faster cells under high-growth conditions, then under low-growth conditions the increased survival advantage of slower cells will allow them to outcompete wildtype cells, producing intrinsic ageing. (G) Comparison of single process model with two opposing process model, demonstrating that in the former, random damage is accumulating without affecting selection. In the latter, the induced damage is still random, but the subset of damaged cells with increased metabolic rate will gain a growth advantage allowing them to outcompete wildtype cells, spreading throughout the tissue so that specific types of damage accumulate according to their induced advantage. However, damage that causes metabolic rate to slow will not spread, but instead the slower mutants will reduce the metabolic rate of surrounding cells by cell communication and epigenetic control. Image created with BioRender.com. Abbreviations: CSF, colony-stimulating factor; DNA, deoxyribose nucleic acid; ECM, extracellular matrix; IGF, insulin-like growth factor; PDGF, platelet-derived growth factor; SD, selective destruction.

This is what happens in type II diabetes. The pancreatic β-cells have an evolved response (called glucotoxicity) whereby prolonged exposure to high levels of glucose induce apoptosis. However, this is not designed to kill β-cells exposed to high glucose levels, but more likely to kill what Karin and Alon^132^ have described as ‘hyper-secreting’ mutant β-cells which aberrantly produce insulin during low glucose states. These cells are a significant threat to homeostasis because β-cells proliferate and produce insulin when blood glucose is high, but apoptose when glucose is low^134^. Thus, without glucotoxicity, hyper-secreting mutants could proliferate and induce wildtype cell apoptosis as they progressively drop blood glucose^132^. Unfortunately, glucotoxicity evolved in hunter-gatherers for whom refined sugar was not as continuously available as in the current Western diet. What was a reliably supra-physiological boundary that killed only mutant β-cells in our ancestors can now be achieved by a diet replete in refined sugar. In these conditions, glucotoxicity has produced a diabetes ‘pandemic’^135^ by killing normal β-cells.

Autonomous mechanisms thus have serious limitations, and additional mechanisms are *required* to control minor mutants. Consistently, Chan, et al.^136^ recently demonstrated that using different strengths of RAS mutation, the strongest RAS mutations reliably induced senescence, while weaker RAS mutations allowed cells to bypass arrest and apoptosis, inducing tumorigenesis, consistent with limitations of autonomous control on minor mutants.

Fortunately, autonomous pathways are not the only mechanisms of control. In mitotic tissues, cells are in constant competition, with various markers and receptors to indicate when cells change their behaviour relative to their neighbours^137–139^. Thus, when a cell obtains even a minor mutation affecting its metabolic outputs, juxtacrine comparison with the neighbouring cells indicates that the shift does not reflect a change in external stimulus – because the surrounding cells show no such equivalent change. Therefore, both the minor mutant and the surrounding cells can induce it to die.

In our previous paper, we suggested Notch signalling as a likely candidate for this juxtacrine communication. While our understanding of Notch signalling is incomplete, many aspects of its mechanism and the effects of associated mutations appear consistent with juxtacrine control of minor mutants. It is a potent anti-cancer pathway with frequent knockouts and activations in various cancers^140^, its juxtacrine signal can induce cell senescence^62^, and Notch mutations in the surrounding non-cancer tissue can aid tumour development^141^. While further study is needed to clarify the details of the mechanism, the results suggest a system by which Notch conveys a message from one cell to another – which cell is inducer and which is recipient being determined by the differences in expression in the Notch receptor and its ligands.

Fascinatingly, after we predicted the significance of Notch in the suppression of minor mutants^1^, Chan, et al.^136^ found one of the two main classes of minor RAS mutants forming cancer was defined by mutations in the Notch pathway, consistent with a knockout of juxtacrine control. Importantly, their results suggest that neither autonomous nor juxtacrine mechanisms of mutant control are completely sufficient to prevent cell selection and spread.

### 3.2 Tissue-level selection leads to celerisis, cancer, and fibrosis

Nelson and Masel^142^ ran a series of computational models of multicellular mitotic tissues suggesting that DNA damage would lead to ‘uncooperative’ mutants of the kind discussed above where selective advantage would potentiate spread. Under all conditions they tested, when competition was between numerically superior wildtype cells and uncooperative cells with a selective advantage, the outcome was the same. Uncooperative cells inevitably ended the simulations dominating the tissue because numeric superiority was not a sufficient counterweight to selective advantage. Being greater in number meant that the wildtype cells could initially identify and remove most of the uncooperative mutants, but unless this control was complete, as the mutant numbers gradually grew, the advantage of numeric superiority waned allowing the uncooperative cells to spread more easily.

In our agent-based models, instead of using general ‘uncooperative cells’, we focused on mutants with accelerated growth, metabolism, and replication (hereon ‘**faster**’ cells)^1^. We allowed neighbouring cells to compare metabolic rates. Similar cells protected each other with survival signals, but if a cell was deemed to be an outlier, then the group of similar cells would send a kill signal with a strength dependent on how different the mutant was. Only the degree of difference to the neighbours was relevant, not whether the mutant was faster or slower (Figure 5B, left), as we argued that greater difference would be both easier to recognise and represent more of a threat to tissue homeostasis. We used a range of likelihoods that mutants would be killed by these interactions, but even at the very highest levels of mutant control, our results replicated those of Nelson and Masel^142^. Faster cells consistently dominated.

We concluded that the **natural course of tissue-level selection** was toward faster cells, just as Nelson and Masel^142^ had concluded that the shift in homeostasis would make ageing inevitable. However, if faster mutant spread caused ageing, then the ageing phenotype should reflect the phenotype of these cells, including a chronic increase in metabolic rate. Ageing tissues should be defined by overgrowth (or hyperfunction) of cells and extracellular matrix (ECM), potentially resulting in a cachexic death as the body depleted its energy reserves.

While this does not accurately describe most of the ageing phenotype, there are age-related diseases which are similar. The most obvious are cancer and fibrosis – the diseases that we suggested could result from celerisis in Part 2.2 as a distinct ageing process. Cancers are well described to secrete metabolites that cause the breakdown of muscle and other tissues to fuel their own growth, producing bodily wasting called ‘cancer cachexia’^143^. Similarly, fibrosis is the consequence of hyperactive fibroblasts producing excessive ECM. These are diseases of hyperfunction, and if they share one underlying cause, a good candidate is the natural course of selection toward faster cells. Thus, we propose **celerisis** as the **process of selection causing the spread of faster cells** (Figure 5C, top).

However, a cancer cell is not simply a fast cell. The natural course of selection in tissues simply causes changes in the homeostatic equilibrium, while tissue size and function are maintained. Cancer cells reflect additional changes both in the loss of response to control stimuli and loss of functional activity, or transformation. Thus, transformation is not the natural or ultimate product of celerisis.

Notably, minor growth mutations will increase mutation rate making faster cells more likely to transform; however, the main reason celerisis produces cancer could reflect that transformed cells escape detection. In tissues where neighbouring cells are constantly comparing, competing, and controlling each other, as cells go through the process of transformation, including genomic instability, crisis, and uncontrolled cell division, such cells will appear less aberrant within a tissue that has undergone significant celerisis. Highly active or rapidly proliferating cells may fail to recognise transformed cells as metabolically distinct and therefore not induce the required kill signals.

Cancers are highly associated with tissues undergoing accelerated growth. One such condition is chronic wound healing, where high levels of cell death necessitate rapid proliferation and ECM production to maintain tissue architecture and cellularity. These conditions are frequently the sites of tumour formation^144^. In the liver, chronic hepatitis B or C infection creates a chronic wound healing response that is a potent inducer of hepatocellular carcinoma (HCC)^128^. Lung cancers too are associated with conditions of chronic cell death from smoking or other particulate exposure followed by compensatory proliferation^145^. Both chronic inflammation and DNA damage are likely crucial contributors to cancers under these conditions^144^, but we suggest that the continually proliferative state is also key as it increases the growth of advantage of faster mutants. Indeed, the signal for compensatory proliferation after cell death has been described to stimulate tumour cell growth^146^.

Consistently, as the Western diet is increasing cases of metabolic dysfunction-associated steatotic liver disease (MASLD), a significant subset are developing HCC in the absence of cirrhosis^147^. Although speculative, these patients might indicate that a strong growth signal does not require the same rates of cell death needed to induce HCC as were previously observed for hepatitis infection or alcohol-related liver disease (ArLD).

In developmental conditions not associated with inflammation or cell death, cancers are still associated with rapid growth. Neuroblastoma is the most common cancer in individuals less than one year of age^148^ likely because this period is associated with rapid development of the neural crest and proliferation of the precursor cells. Similarly, testicular cancer has peak incidence in younger adults^149^, which could (among other factors) reflect the period of peak germ cell activity, suggesting metabolic and proliferative rates could contribute to cancer induction.

Notably, in addition to cancer, hepatitis B and C infections are also highly associated with liver fibrosis^150^, just as smoking and particulate exposures induce both IPF and lung cancer^151^. At its core, the term ‘fibrosis’ refers to a disease state of overactive fibroblasts and their activated form, myofibroblasts, producing excessive ECM^152^. This is distinct from ‘adaptive fibrosis’, where the extra ECM has important tissue function. In IPF, the encroaching collagen mesh prevents alveoli from expanding to intake gasses, leading to hypoxemic respiratory failure^153^. In the liver, early-stage fibrosis can still be reversible, but in the lungs IPF is a chronic and irreversible condition where fibroblasts continue to activate but do not deactivate via the usual pathway.

In a computational study by Adler, et al.^133^, adaptive fibrosis was modelled as a circuit between immune cells and fibroblasts, where a wound would attract and activate macrophages, which would secrete platelet-derived growth factor (PDGF) to activate fibroblasts forming myofibroblasts. Myofibroblasts in turn would then secrete colony-stimulating factor (CSF) to activate macrophages, and PDGF to activate other fibroblasts (Figure 5D). Thus, under most conditions, fibrosis remained adaptive, maintained by immune cells and inflammation. When the wound closed and the immune cells deactivated, the myofibroblasts would lose their activation signal and either apoptose or senesce, helping to clear the ECM. However, if the wound remained open too long, or was repeatedly reopened before the myofibroblasts could deactivate, then the persistent inflammation signal would continue to increase the myofibroblast population to a threshold. Beyond this point, enough myofibroblasts would be secreting self-activating PDGF that no macrophages were required. At that point, the myofibroblasts became self-sustaining; wound closure and the exfiltration of immune cells no longer causing the population to decline (Figure 5E). Adaptive fibrosis would become fibrosis and wound healing would become disease.

A major determinant of whether the myofibroblasts reach their self-sustaining threshold is thus likely to be their metabolic rate. Faster fibroblasts with minor growth mutations will secrete more CSF and PDGF and respond more robustly to the PDGF signal; they may also proliferate faster and be slower to deactivate when the signal dies. Indeed, IPF fibroblasts have demonstrated increased proliferative rate due to repressed FOXO3a activity and resultant upregulated AKT signalling^154^, potentially requiring shorter and weaker signals to reach the threshold. In effect, celerisis should promote fibrosis.

Notably, there are multiple connections between fibrosis and cancer, including chronic inflammation^152^ (see Part 5.3). However, as we suggest here, celerisis itself could reflect a major underlying process that causes risk of both cancer and fibrosis to increase with age.

### 3.3 Selective destruction prevents celerisis but causes intrinsic ageing through metabolic slowdown

As described by Nelson and Masel^142^, the dominance of uncooperative cells with a selective advantage was inevitable, making ageing a necessary result. Equally, our agent-based models where wildtype cells could kill neighbouring fast cells with high efficiency also failed to prevent the spread of faster cells^1^. The implication of these results is that ageing reflects an unavoidable shift that is the natural course of tissue-level selection: sites of repeated wound healing produce cancer and fibrosis first because they are the tissues where the growth advantage is greatest, but all tissues should demonstrate increasing metabolic rate over time. However, as discussed, this is largely unrepresentative of the ageing phenotype.

Cancer and fibrosis aside, ageing is not defined by hyperfunctional growth, but by a state of hypofunction. Instead of increasing metabolic rate, there is strong evidence that most cells, tissues, and organisms undergo a progressive metabolic slowdown^155,156^. In humans, basal metabolic rate declines with age independent of the decline in fat-free mass^157–159^. It is also evident in mice^160^, rats^161–163^ and dogs^164^. It is well described that in humans, the rate of cell division declines with age^165^, while a meta-analysis of transcriptomic changes with age across model organisms and humans found that downregulation of mitochondrial genes and protein synthesis machinery, particularly ribosomal proteins and biogenesis factors, were the most consistent age-related changes^166^, highly consistent with a **metabolic slowdown**.

Our tissue models showed that, contrary to the results of Nelson and Masel^142^, a single change in the way neighbouring cells compare and control each other could prevent the inevitable dominance of faster cells. Using the same model where neighbouring cells compared metabolic rates and identified whether cells were likely to be mutant, we made a single change. In the second version, slower cells would be more likely to escape identification and killing by faster neighbours (Figure 5B, right). This time faster cells could be controlled, allowing our simulated tissues to retain homeostasis throughout. This reflected that slower cells now had their own selective advantage – a survival advantage – that counteracted the growth advantage of the faster cells. The natural course of tissue-level selection could thus be balanced by counterselection.

If the growth advantage was stronger than the survival advantage, then faster cells still spread, but by balancing the growth and survival advantage, wildtype cells could retain dominance throughout the simulations, and a stronger survival advantage could even cause the spread of slower cells, against the natural course of tissue-level selection, so that while wildtype cells outcompeted faster cells and slower cells eventually outcompeted wildtype cells. To maintain homeostasis in a tissue with minor mutants required balancing the natural course of selection with an equal force of counterselection.

We called this program of mutant identification which advantaged slower cells ‘**selective destruction**’, as its purpose was targeting mutant control toward killing faster cells rather than unselectively attempting to kill any cell presumed to be mutant (either slower or faster). It is worth noting that the survival advantage of slower cells refers specifically to the likelihood of being removed by selective destruction. As growth and survival signals are frequently linked and positively co-regulated, faster cells would also have a natural survival advantage (see Part 7.2 for more detail).

However, we also suggested that faster cells need not be killed by selective destruction. Unlike cancer cells, which have lost both cell function and response to control stimuli, faster cells still retain both these crucial characteristics; they have merely shifted their metabolic equilibrium. Plausibly, minor shifts in metabolic rate could be controlled epigenetically with chromatin modifications reversing the presumed genetic changes that caused the growth acceleration^167^ (Figure 5C). As these marks are semi-permanent, inherited by the daughter cells, they could induce long-term effects on metabolic rate that represent a more efficient mechanism to control minor mutants without exacerbating the cycle of cell death and proliferation that aids celerisis. We further demonstrated that epigenetic control was a more effective mutant control mechanism than simple killing^1^. Thus, we hypothesise **selective destruction is a mechanism of mutant control by juxtacrine induction of epigenetic alterations that advantages cells which are disadvantaged by the natural course of tissue-level selection** (in this case slower cells) by less frequently identifying them as mutant and either reprogramming or killing them.

As noted above, our models of selective destruction could maintain tissue homeostasis. The natural course of selection and counterselection could be balanced. However, our models were missing a key aspect of competition. Faster mutants with increased response to growth stimuli will not have a constant selective advantage. The difference between their growth rate and that of wildtype cells will change in accordance with the growth stimulus, increasing their selective advantage as growth stimuli increase (Figure 5F).

Assuming selective destruction lacks the same plasticity for how much of a counter-advantage is given to slower cells, then to prevent celerisis under any conditions, the process must be sufficiently selective to produce a survival advantage that equals the growth advantage of faster cells under the strongest growth conditions. If it is less selective, better identifying both faster and slower mutants, then stronger growth conditions will shift the net advantage to faster cells and cause celerisis. Preventing celerisis may thus require selective destruction to allow slower mutants to escape control under some conditions. Local spread of these mutants could then begin the gradual epigenetic reprogramming of the surrounding tissue toward their own slower metabolism. If true, intrinsic ageing evolved as an epigenetic mutant control programme that deliberately avoided the reprogramming (or killing) of slower mutants. Put another way, **intrinsic ageing may reflect metabolic slowdown induced by an overly selective form of selective destruction**. Essentially, our tissues become slower to avoid becoming faster.

This may explain why cancers and fibrosis frequently occur at sites of chronic wound healing where rapid growth conditions are sufficient to overcome selective destruction. Notably, under these conditions, selective destruction may even be suppressed if the short-term risks of open wounds and reduced cellularity outweigh the long-term risks of shifting tissue homeostasis.

Crucially, the existence of selective destruction as a program designed to stop celerisis could provide an evolutionary advantage for ageing organisms over negligibly senescent ones. While gerontologists frequently question why ageing might evolve, few directly question the evolution of cancer. This reflects partly that mainstream theory includes cancer as part of the same ageing process, but when separated as they are here, the evolution of cancer still requires no explanation. Cancer is evolution at the tissue-level. It reflects the natural course of tissue-level selection in the absence of control. Thus, cancer does not need to produce advantages at the organismal level because it reflects advantages gained by cells at the tissue-level. Indeed, in extreme cases, this natural course of tissue-level selection even becomes evolution, producing new organisms in the form of infectious cancers which spread from host to host^168^.

Intrinsic ageing could therefore evolve to combat the natural course of selection in tissues. Its advantage to the organism would be reduced risk of early death from celerisis and the resulting hyperfunctional disorders, and the cost would be gradual metabolic slowdown over time. Thus, evolution of intrinsic ageing would be antagonistically pleiotropic, preventing earlier or more sporadic mortality from the natural course of tissue-level selection.

It is worth noting that, contrary to our previous claims^1^, the underlying cause of ageing is still DNA damage. In the absence of damage, organisms with an unaltering DNA code could likely retain continuous molecular and cellular equilibrium with relative ease. There would be no selection and no need for counterselection by selective destruction. However, where ageing via random DNA damage accumulation has produced inconsistencies and complexities (Figure 5G, left), with few useful predictions or robust mechanistic models, ageing as a product of damage-induced selection and counterselection is consistent with the two opposing process model of ageing from Part 2.2. **Celerisis is a process of genetic change**. **DNA damage induces mutation and accumulates as selection causes the damaged cell to spread** (Figure 5G, middle). The damage is the same from cell to cell, and results in a specific homeostatic shift toward faster metabolism. Intrinsic ageing, by contrast, is a process of largely epigenetic changes, and notably not one of increasing ‘epigenetic noise’ but specific epigenetic modifications designed to reduce the metabolic rate of neighbouring cells (Figure 5G, right). It is an evolved response to damage and could arguably even occur in its absence if non-genetic, environmental or locational differences caused disparity in metabolic rates between cells.

Although we suggest this model is a significant improvement over the ‘complex multifactorial process’, there are aspects of ageing that the two opposing process model does not explain. Cancer and the other age-related diseases are not always counter-regulated; multiple factors and conditions correlate positively with both – the most significant being chronic basal inflammation and metabolic syndrome. In Part 4, we describe how an additional process can explain these inconsistencies to produce a more unifying model of ageing. However, an additional limitation to the two process model is that intrinsic ageing can only apply to multicellular, mitotic tissues. Post-mitotic and single-celled organisms as well as post-mitotic tissues in mitotic multicellular organisms are unlikely to age as a result of intracellular competition to prevent the spread of hyperfunctional cells because such tissues and organisms are relatively invulnerable to this spread. We discuss why selective destruction might still apply to these tissues and organisms in Part 7.

## Part 3: Summary

3.1 | The damage that accumulates with age is thought of as random and not subject to selection. Yet some damage will give cells a competitive advantage, such as pro-growth mutations. These cells will spread and alter homeostasis if not controlled.

We have autonomous mechanisms of cell death and arrest that can remove the threat of mutants with large pro-growth effects. However, such autonomous mechanisms cannot control mutants with growth increases within physiological norms. Such mutations are indistinguishable from changes in environmental growth signalling.

Our tissues have mechanisms for cells to compare metabolic rates between neighbours and then remove mutants which, based on membrane markers, are identified as different.

Notch signalling may reflect a major regulator of cellular consistency.

Even these juxtacrine (between connected cells) mechanisms can fail to prevent the spread of mutants with a selective advantage if wildtype cells lack their own advantage to outcompete them.

3.2 | The spread of faster mutants could result in hyperfunctional diseases such as cancer and fibrosis. The **natural course of tissue-level selection** could be the cause of celerisis, the ageing process responsible for increasing risk of these diseases.

3.3 | A mutant control mechanism which selectively targets faster mutants could prevent the spread of faster cells by giving slower cells their own selective (survival) advantage. We called this process **selective destruction**.

Selective destruction could include an epigenetic mechanism by which slower cells slow the growth of neighbouring faster cells by altering their chromatin in a semi-permanent way.

By balancing the natural course of selection for faster mutants with counterselection for slower mutants, life could maintain homeostasis indefinitely, producing negligible senescence.

However, as growth signals change, so will the relative advantage of faster cells. In some situations they should be unbalanced. Therefore, the body may be overly selective against faster cells, leading to the gradual spread of epigenetic marks which reduce metabolic rate of cells. This **metabolic slowdown** could be the underlying cause of intrinsic ageing and the age-related diseases associated with hypofunction.

## Part 4: Metabolic slowdown causes extrinsic ageing

### 4.1 Metabolic slowdown induces mitochondrial dysfunction

In our model, the primary outcome of selective destruction is the spread of epigenetic marks that slow the anabolic metabolism of cells within proliferative tissues. As cells gain more of these marks their response to growth stimuli decreases either in rate or magnitude or both, reducing the cellular energy demands per unit time.

The main source of bioavailable energy in our bodies is adenosine triphosphate (ATP). Maintaining ATP equilibrium is therefore crucial to continued cellular function, and it is under strict homeostatic control. Despite decreasing basal and stimulated anabolism with ageing^169,170^, ATP levels appear remarkably stable. There is little difference in ATP or total adenosine phosphate levels in ageing mice^171,172^ or ATP in humans from age zero to one hundred^173^. Indeed, Tfam-knockout mice with progressively deteriorating mitochondrial function still maintain ATP levels by increasing mitochondrial mass^174^. As described by Hancock, et al. (2006), “the rate of ATP hydrolysis is matched by the rate of ATP supply without measurable changes in ATP over a wide range of ATP demands (up to very intense short-term near-maximal exercise)”^175,176^.

To maintain ATP equilibrium, any change in ATP usage should be met by the reciprocal change in ATP production. When ATP levels drop, ATP production increases, and when ATP levels rise, ATP production decreases. Notably, as both ATP synthesis and hydrolysis can be described as ‘metabolism’, for simplicity we refer to **metabolism involved in ATP synthesis** as **catabolism** and **metabolising requiring ATP hydrolysis** as **anabolism**.

ATP synthesis is a process of largely mitochondrial catabolism (MC) involving Kreb’s cycle and oxidative phosphorylation (OXPHOS); however, **glycolysis** produces small amounts of ATP outside the mitochondria, as shown in equation 1 for ATP dynamics.

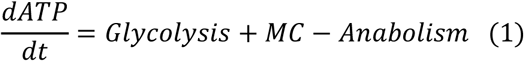

Metabolic slowdown in the context of selective destruction reflects reduced anabolism: cells undergoing less transcription and translation, reducing growth, and slowing proliferation, all reduce requirements for ATP hydrolysis. This is shown in Figure 6A as the first arrow (1↓) in a sequence which will describe the shifts induced by reduced anabolism. As ATP levels rise (2↑), negative feedback systems are activated to reduce ATP synthesis and return ATP levels to equilibrium. Assuming glycolysis is only a small fraction of ATP production, then mitochondrial catabolism (MC) must also decline (Anabolism ≈ MC, from equation 1).

**Figure 6.**
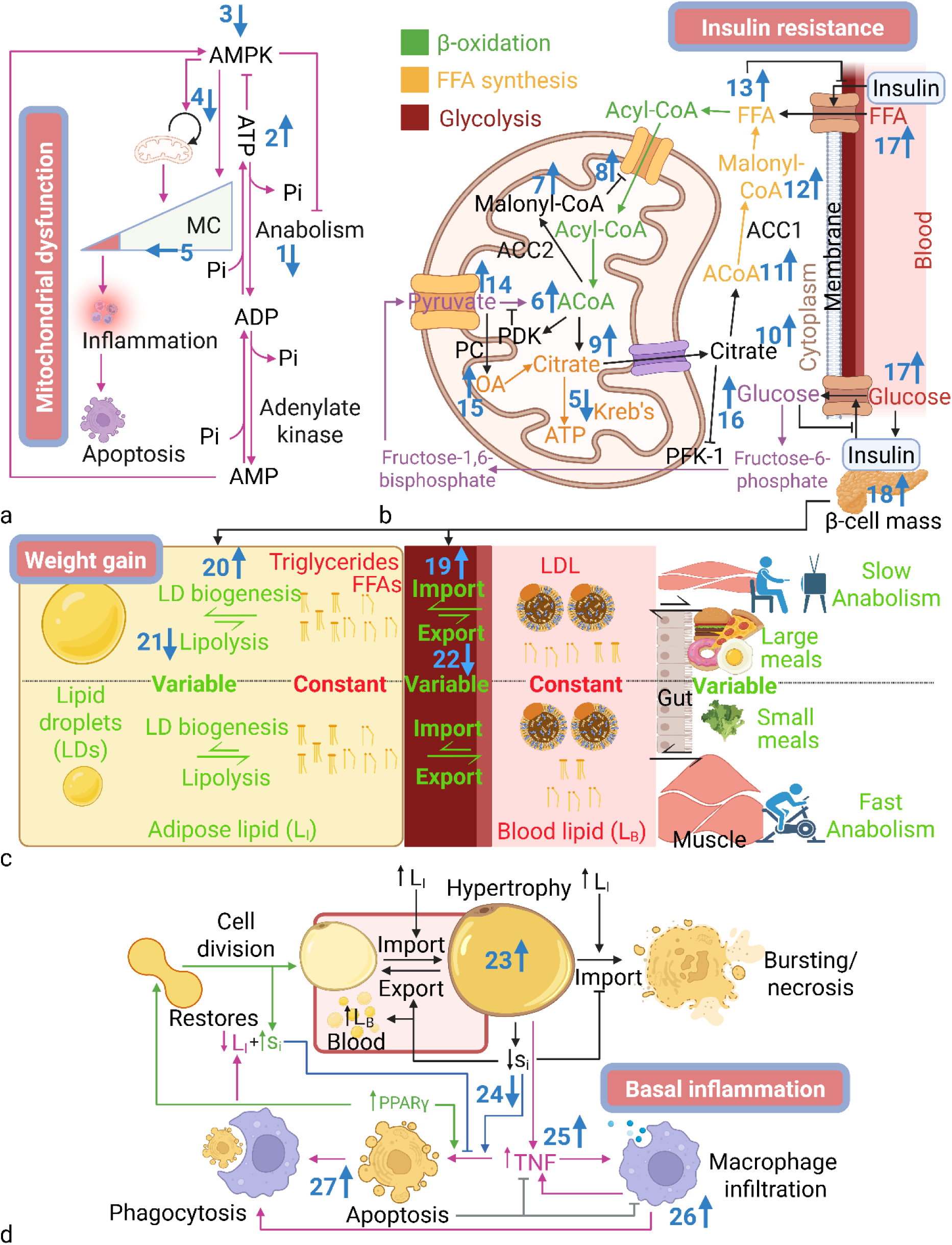
Regulation of energy homeostasis. (A-D) Blue numbers indicate age-related changes and homeostatic effects leading to consequences in blue-bordered boxes. (A) Regulation of ATP dynamics. (B) Model of ACoA dynamics. (C) Regulation of lipid dynamics maintaining homeostasis despite a changing environment. (D) L_I_ homeostasis regulated by inflammation and apoptosis of hypertrophic cells. Image created with BioRender.com. Abbreviations: ACoA, acetyl-coenzyme A; ADP, adenosine diphosphate; AMP, adenosine monophosphate; AMPK, AMP kinase; ATP, adenosine triphosphate; CoA, coenzyme A; FFA, free fatty acid; L_B_, blood lipid; LD, lipid droplet; LDL, low density lipoprotein; L_I_, intracellular lipid; MC, mitochondrial catabolism; OA, oxaloacetate; PC, pyruvate carboxylase; PDK, pyruvate dehydrogenase kinase; PFK, phosphofructokinase; P_i_, inorganic phosphate group; PPAR, peroxisome proliferator-activated receptor; s_i_, insulin sensitivity; TNF, tumour necrosis factor.

Notably, a rigorously maintained equilibrium for ATP is not required for declining anabolism to reduce catabolism. Simple negative feedback – which is part of all biological systems – will result in the same (though weaker) effect. A central regulator of ATP homeostasis is **AMP-activated protein kinase** (**AMPK**) which is activated through an allosteric mechanism by high ratio of AMP:ATP. As such, a key function of AMPK is to restore ATP as levels drop in response to increasing anabolism^177^. Thus, AMPK regulates two main pathways designed to restore ATP levels: firstly, it slows growth and cell cycle by deactivating mammalian Target of rapamycin complex 1 (mTORC1)/mitogen-activated protein kinase (MAPK) pathways, **inhibiting anabolism**^178^, and secondly it activates transcription factor EB (TFEB) and Peroxisome proliferator-activated receptor-γ coactivator (PGC)-1α, triggering mitochondrial biogenesis and OXPHOS, **activating mitochondrial catabolism**^179^ (Figure 6A).

Therefore, as metabolic slowdown induces ATP build-up (high ATP, low AMP), it should deactivate AMPK (3↓) and reduce mitochondrial ATP production through multiple mechanisms, including reduced mitophagy and mitochondrial biogenesis (4↓) and deceleration of the electron transport chain. The predicted result is reduced mitochondrial catabolism (5↓), with decreases in OXPHOS triggering a decrease in NAD^+^:NADH+H^+^ ratio, which then reduces Kreb’s cycle flux. Indeed, although we have focussed on AMPK regulation, mitochondrial metabolism is self-regulating as complex V is sensitive to levels of its substrate, ADP, to complete the final reaction at the mitochondrial membrane^180^. As ATP increases, membranes become hyperpolarised and induce ROS production^181–184^, which could easily be mistaken for the effects of accumulating mitochondrial damage. Indeed, the conventional wisdom is that ROS produced during OXPHOS cause damage accumulation in the mtDNA, reducing mitochondrial function^185^. The reduced catabolism then reduces the cell’s capacity for anabolism as ATP levels decline. However, as described, the picture is complicated by evidence that oxidative damage does not increase and ATP levels do not decline. Contrarily, we suggest that declining anabolism might be the initial cause, necessitating reduced mitochondrial activity to maintain ATP homeostasis. As mitochondrial catabolism slows, the organelles may increasingly activate inflammatory and pro-apoptotic pathways associated with reduced activity. ‘Mitochondrial dysfunction’ may therefore not necessarily reflect damage-induced dysfunction at all, but a homeostatic response to metabolic slowdown.

If the primary inducer of metabolic slowdown is epigenetic modification, one factor that could help distinguish mitochondrial dysfunction as either a homeostatic response or due to mtDNA damage is that Yamanaka (OSKM) factor epigenetic rejuvenation can restore mitochondrial function^186,187^, which is more consistent with epigenetic changes than genetic ones. Notably heteroplasmy is still a confounding factor that requires further investigation. However, in our model, declining catabolism is merely the first compensatory shift in a long chain.

### 4.2 Metabolic slowdown induces insulin resistance (IR)

Just as homeostasis during reduced ATP hydrolysis should cause a compensatory reduction in ATP production, reduced ATP production should stimulate compensatory shifts to prevent build-up of the sugar and lipid fuels no longer burned to produce ATP. The primary fuels entering the mitochondria are pyruvate from glycolysis and acyl-CoA from the **β-oxidation** of free fatty acids (FFAs). Both pyruvate and acyl-CoA are then converted to acetyl-CoA (ACoA), which is then used for mitochondrial catabolism (MC) or converted back into FFAs via **FFA synthesis**, as shown in equation 2.

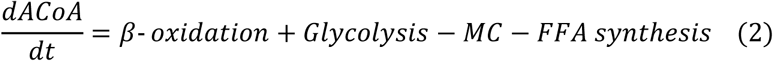

Thus as MC declines (Figure 6B, 5↓) and less ACoA is converted to carbon dioxide by entering Kreb’s cycle, the rise in ACoA levels (6↑) is inhibited by the activation of acetyl-CoA carboxylase enzymes 1 & 2 (ACC1 & 2) in a system of robust negative feedback that converts ACoA back to malonyl-CoA. In non-lipogenic tissues, production of malonyl-CoA by ACC2 within the mitochondria increases (7↑), inhibiting the import of Acyl-CoA (8↑) and thus inhibiting β-oxidation (Figure 6B, Green).

Lipogenic tissues have an additional negative feedback system where the excess citrate (9↑), formed via the condensation of ACoA and oxaloacetate, is pumped out of the mitochondria as part of the citrate-malate shuttle. Cytoplasmic citrate (10↑) is used to generate cytoplasmic ACoA (11↑), which is converted to cytoplasmic malonyl-CoA (12↑), the substrate for FFA synthesis (Figure 6B Yellow). Both reduced β-oxidation and increased FFA synthesis result in increased levels of FFAs (13↑).

Similar negative feedback mechanisms also exist to inhibit glycolysis (Figure 6B Purple). For example, ACoA is an allosteric activator for pyruvate dehydrogenase kinase (PDK), which deactivates pyruvate dehydrogenase, inhibiting the conversion of pyruvate to ACoA, increasing mitochondrial pyruvate (14↑). Excess pyruvate can then be converted to oxaloacetate (OA) by pyruvate carboxylase (15↑), which condenses with ACoA to increase citrate (9↑). As in lipid metabolism, excess citrate is exported via the citrate-malate shuttle. Crucially, cytoplasmic citrate then inhibits phosphofructokinase-1 (PFK-1) which converts fructose-6-phosphate to fructose-1,6-bisphosphate, the rate limiting reaction for glycolysis. Thus, declining mitochondrial catabolism also increases cytoplasmic glucose (16↑).

In non-lipogenic tissues, where excess fuels cannot be easily stored, particularly FFAs quickly become toxic to metabolism^188^, so that equilibrium should be restored by reducing FFA and glucose import into cells. **Import** is controlled primarily by **insulin** through the activation of cluster of differentiation 36 (CD36) for lipids^189^ and GLUT4 for glucose. Non-lipogenic tissues with excess FFAs should therefore reduce their **insulin sensitivity** (**s_i_**), as suggested by clinical studies^190^.

The molecular mechanisms by which this occurs are likely to be tissue dependent, as reviewed elsewhere^191^. For example, excess glucose causes serine phosphorylation of IRS-1/2, reducing PI3K–Akt signalling and the resultant translocation of GLUT4. In muscle, excess cytoplasmic FFAs are converted to ceramides and diacyl-glycerol, which inhibit translocation of GLUT4 and CD36 to the sarcolemmal membrane^192^.

Thus, we observe that in order to prevent the toxic effects of cytoplasmic fuel overload, **metabolic slowdown induces IR**, which is one of the key hallmarks of ageing, highly associated with mortality^193^. Consistently, mice engineered to have lower basal metabolic rate showed propensity toward IR and diabetes, as well as the next product of metabolic slowdown, weight gain^194^.

### 4.3 Metabolic slowdown induces weight gain

Following our chain of causation, as our non-lipogenic tissues become insulin resistant, removing fewer FFAs from the blood, the immediate effect (without compensation) should be increased FFAs and glucose in the blood (17↑). Unlike glucose, FFAs are insoluble and cannot be transported freely through the blood. Some are bound to albumin, but most are transported as esters bound to lipoproteins. For simplicity, the combined **blood lipids** are referred to as **L_B_**.

While levels of L_B_ may fluctuate according to various stimuli, blood glucose is maintained at rigorous equilibrium, usually around 5mM in humans by a system of robust negative feedback^134,195^. Any increase in blood glucose will trigger a corresponding increase in insulin production (18↑) and restoration of blood glucose to equilibrium within two hours.

This would produce a vicious cycle in our non-lipogenic tissues as they become increasingly insulin resistant to remove excess cytoplasmic fuel, inducing increased insulin production to remove excess fuel from the blood, leading to further insulin resistance as the tissues are forced to import the excess fuel again. Thus, we have our lipogenic tissues to prevent this spiral and safely store our excess fuels.

Our two central lipogenic tissues are the liver and adipose, which are usually among the last tissues to become insulin resistant because they can store FFAs in a non-toxic form that does not interfere with metabolism. The fatty acid esters (triglycerides) are packaged up into inert lipid droplets (LDs) coated in cholesterol-rich, hydrophilic membranes^196^. Lipogenic tissues can therefore respond to increased L_B_ and blood glucose with increased import and storage.

The liver is the central storage depot for excess glucose, storing it as a finite quantity of glycogen, limited largely by the soluble nature of the molecule that makes it a ‘heavy’ fuel. Thus, in intense exercise the limited stores of glycogen can be almost completely exhausted. Conversely, if glycogen stores are full, further excess glucose is converted to FFAs in both the liver and adipose via de novo lipogenesis.

By pairing LD biogenesis and import, both stimulated by insulin (Figure 6C 19↑ and 20↑), as well as pairing lipolysis and export which are inhibited by insulin (21↓and 22↓), any change in fuel intake from meals or catabolism of tissues results only in changes in LDs in lipogenic tissues without any increase toxic intracellular FFAs or glucose in any cells, as shown in Figure 6C. Thus, L_B_ and blood glucose (G_B_) homeostasis are maintained as in equation 3 (where import and export refer exclusively to lipogenic tissues).

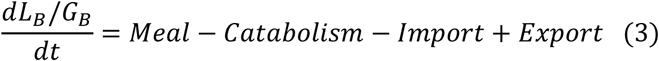

Notably, L_B_ homeostasis is not as robust as G_B_ homeostasis, partly due to the increased complexity of lipid regulation, and the primary response of pancreatic β-cells to G_B_. Therefore, L_B_ takes longer to return to equilibrium, which can also vary according to other factors.

However, as stated above for ATP, robust equilibrium is not required for this cascade. Any level of negative feedback attempting to reduce the amount of L_B_ toward the preferred equilibrium will result in an increase in lipid stored in the adipose tissue, leading to **weight gain with age**.

Thus, the **proximal effects of metabolic slowdown are likely to be weight gain and IR** as we store excess lipids in LDs. However, as metabolic slowdown continues, our lipogenic tissues should induce their own homeostatic mechanisms and induce additional phenotypes of ageing.

### 4.4 Metabolic slowdown induces chronic basal inflammation

There is little evidence for a tightly regulated equilibrium value of intracellular lipid (L_I_)^197^. As fuel availability rises, be it from metabolic slowdown or dietary increase, then assuming other factors remain equal, it causes an increase in storage and a corresponding increase in weight (Figure 6D, 23↑). However, each additional excess calorie does not equal one excess calorie stored, with every rise resulting in an increasingly small fraction being stored^198^. This suggests we do have negative feedback mechanisms to curb increasing L_I_ which become increasingly active as the energy balance becomes increasingly positive.

Likely, part of this mechanism reflects the activation of uncoupling in mitochondria, though its role is still equivocal. At least part of this confusion likely reflects the widespread interpretation of uncoupling studies within the complex narrative of oxidative stress^199^. Uncoupling allows mitochondria to dissipate the proton gradient without generating ATP. Thus fuels can be burned in the mitochondria without producing bioavailable energy, the main emission being heat.

Indeed, in the brown adipose tissue (BAT), expressing high levels of uncoupling protein 1 (UCP1), heat generation appears to be the main function rather than dissipation of the proton gradient: uncoupling is higher under ambiently colder conditions^200^; rises during cold stimulation^201^; and most importantly is blunted by obesity^202^, the latter opposing expectations of a process designed to burn excess lipid. However, while thermogenesis is likely the central function of UCP1 and BAT, most tissues express low levels of uncoupling protein 2 (UCP2), which may function in the event of energy excess. One study in humans found muscle-specific uncoupling protein 3 (UCP3) was upregulated 44% by high fat diet when compared to low fat diet^203^. Another study indicated that increased plasma glucose upregulated expression of UCP2 and UCP3^204^, indicating that UCPs are upregulated post-prandially. However, as weight is still gained, some of the excess must be stored, and additional mechanisms are required to regulate the stored lipid.

Notably, although adipose is a proliferative tissue, with around 10% turnover per year in humans^205^, most studies suggest that the number of adipocytes in adulthood remains relatively fixed (with equal rates of cell death and proliferation)^205–207^. Instead, the adipose mass grows and shrinks mainly by changing cell size^208^. This finite storage capacity combined with metabolic slowdown suggests that without additional homeostatic mechanisms, ageing could saturate the adipose tissue with stored lipid.

As the adipocytes store increasing LDs, they expand, becoming hypertrophic. To avoid bursting, hypertrophic adipocytes can reduce import of L_B_ by reducing insulin sensitivity (s_i_), just as in non-lipogenic tissues (24↓). However, preventing FFA import would increase L_B_, contrary to adipose function. Thus, we predicted an additional mechanism to lower L_I_.

Although the cause of inflammation in obesity is often stated as “unknown”^209^, evidence suggests it plays a central role in the induction of **apoptosis** in hypertrophic adipocytes and their subsequent phagocytosis by macrophages. Hypertrophic adipocytes secrete cytokines such as TNF-α (25↑) which attract resident macrophages (26↑), which then secrete factors promoting apoptosis^210^. This would explain why obesity is associated with an increase in macrophage infiltration^211^, and inflammatory M1 polarisation^212^. Additionally, immune-deficient mice have increased adipose mass^213,214^ and immune reconstitution of immunodeficient mice on high-calorie diets reduces weight gain^214^.

In both lean and obese mice, Cinti, et al.^215^ found that more than 90% of macrophages localized around large or dead adipocytes forming crown-like structures (CLS), suggesting adipocyte removal by apoptosis and ingestion of hypertrophic cells. The same immune cell localisation around hypertrophic or ‘ballooning’ hepatocytes is also found in the liver, the other major organ of lipid storage. Therefore, L_I_ dynamics are shown in equation 4.

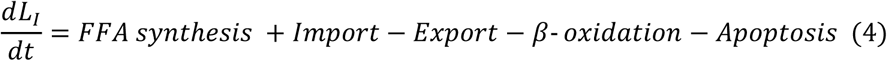

Thus we suggest that as metabolic slowdown increases FFA synthesis and import, and reduces β-oxidation, the increasing L_I_ requires increased apoptosis (27↑) to remove the hypertrophic cells, which requires increasing inflammation.

Intriguingly, one study using the FAT-ATTAC model to induce adipocyte apoptosis found that apoptosis itself recruited M2 anti-inflammatory macrophages, while living adipocytes were required to induce metabolic switching to pro-inflammatory M1 macrophages^216^. Consistently, Haase, et al.^217^ found macrophages mainly proliferate at CLS, where they are inducing apoptosis, but proliferation produces mainly M2 macrophages, presumably providing the negative feedback that inhibits inflammation after the hypertrophic adipocytes are removed (Figure 6D, grey). These results indicate that despite appearances of inflammation sustaining itself in a counterproductive way in obesity, as suggested by damage-centric and inflammaging models, the pro-inflammatory state could be retained mainly by hypertrophic adipocytes as a necessary homeostatic pathway.

Removal of hypertrophic adipocytes has potential to return both L_I_ and insulin sensitivity (s_i_) to equilibrium levels because Peroxisome Proliferator-Activated Receptor gamma (PPARγ) activates both apoptosis and adipocyte differentiation from stem cells^218^ (Figure 6D, green), replacing L_I_-laden, insulin resistant hypertrophic cells with smaller insulin sensitive ones, thus also maintaining constant adipocyte number^205^. Consistently, inflammation-deficient transgenic mice had reduced adipocyte proliferation and fewer total adipocytes than wildtype mice, causing hepatic steatosis and metabolic dysfunction^219^. Therefore, we suggest that **metabolic slowdown induces chronic basal inflammation** to ameliorate the age-related increase in lipid levels.

Notably, macrophage ingestion of the excess L_I_ may not remove the lipid from the body. Macrophages could burn away the lipid in uncoupled mitochondria or in the peroxisomes, but evidence currently suggests that these are not major pathways for lipid removal. Instead, the FFAs and cholesterol are packaged onto high-density lipoprotein (HDL) and targeted to the liver^220^, which is the central organ of nutrient regulation. Excess cholesterol is secreted into the bile to be removed by excretion in a process called reverse cholesterol transport^221^. Despite the vital importance of excess FFA regulation, their fate is currently equivocal. Uncoupling and conversion to cholesterol both occur in the liver, but which process or processes reflect the major pathway(s) and under which conditions requires further research. That we have mechanisms to calculate and remove excess FFAs from the liver is highly likely.

It is worth noting that the systems we have evolved to rid the body of excess calories is good evidence that organisms have not evolved to under-invest in maintenance^74^. Instead of an upregulation of UCP2 and UCP3 after large meals, we might expect instead a boost in cellular maintenance. We suggest that organisms which evolve more efficient energy utilisation, and thus require fewer calories for optimal function, likely gain a fitness advantage that may allow them to outcompete species better able to procure (but less efficiently use) nutrients; however, organisms which cannot afford to invest sufficiently in both somatic maintenance and offspring should be reliably outcompeted by organisms which can procure sufficient nutrition for both.

The evolution of nutrient wastage is good evidence that organisms require processes to maintain energy homeostasis during times of excess even in wild populations.

### 4.5 Metabolic slowdown induces metabolic syndrome

Chronic low-grade inflammation is not without multiple negative implications, as we will discuss in Part 5. However, we suggest it is also a crucial negative feedback system that slows the progression from metabolic slowdown to metabolic syndrome. According to the British Medical Journal (BMJ), metabolic syndrome is defined as ‘a cluster of common abnormalities, including insulin resistance, impaired glucose tolerance, abdominal obesity, reduced high-density lipoprotein (HDL)-cholesterol levels, elevated triglycerides, and hypertension’^222^.

We have already described how IR and elevated triglycerides result directly from metabolic slowdown. The significance of low HDL-cholesterol could reflect the necessity of HDL in removing excess L_I_ from the adipose tissue^220^. While not a direct result of metabolic slowdown, low HDL should reduce ability to clear L_I_ from the adipose and thus exacerbate or predispose to metabolic syndrome, as described below.

The central adipose store for healthy individuals is the subcutaneous adipose tissue, which is designed for long-term storage. The visceral adipose tissue is, by design, more insulin resistant than subcutaneous adipose because its purpose is not long-term storage. It provides a ready supply of lipid to the visceral organs in times of stress where immediate energy is required without having to wait for increased lipid supplies from the more distal subcutaneous adipose. Thus, the primary inducer of visceral lipid storage is not insulin but the stress hormone cortisol, which both induces insulin resistance of subcutaneous adipose and diverts subcutaneous lipid to the viscera for immediate use or storage in the visceral adipose^223^.

As metabolic slowdown causes the subcutaneous adipose to accumulate L_I_, any increase in insulin resistant, hypertrophic adipocytes makes the subcutaneous adipose less capable of removing lipid from the blood. An intermediate rise in L_B_ then instigates its diversion to other adipose tissues for storage. Given that the subcutaneous adipose is increasingly insulin resistant, the sensitivity differential between the subcutaneous and visceral adipose decreases, diverting L_B_ to the viscera. Thus, **metabolic slowdown should induce abdominal obesity** (Figure 7).

**Figure 7.**
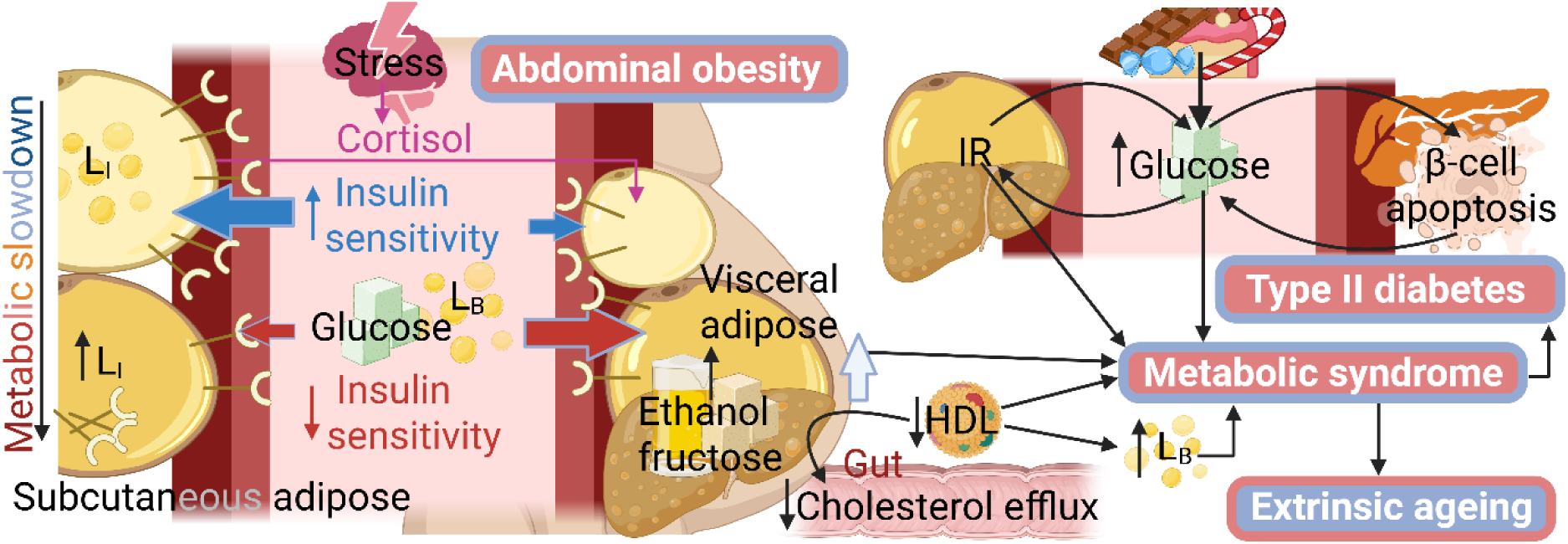
Metabolic slowdown induces metabolic syndrome. As subcutaneous adipose becomes saturated and insulin resistant, glucose and lipids are diverted to the visceral adipose, worsened by fructose and ethanol which are broken down by the liver and frequently converted to visceral fat. As visceral adipose expands, particularly under the influence of a high sugar, high calorie diet, L_B_ and blood glucose can remain elevated for longer inducing glucotoxicity and type II diabetes. Abbreviations: HDL, high density lipoprotein; IR, insulin resistance; L_B_, blood lipid; L_I_, intracellular lipid.

As described above, blood glucose equilibrium is more robust than for lipids. In healthy individuals rises in blood glucose cause increased insulin production and β-cell proliferation ensuring that glucose is absorbed and returns swiftly to homeostasis^134^. The exception is in type II diabetes where chronic elevations of blood glucose shift the β-cell response from proliferation to apoptosis, reflecting the mutant defence mechanism we described in Part 3^132,195^.

Metabolic slowdown and intrinsic ageing alone are not normally sufficient to cause diabetes, as evidenced by the lack of metabolic syndrome and type II diabetes in modern hunter-gatherer and semi-nomadic tribes^224,225^. However, the Western diet produces chronic blood glucose at levels which were supraphysiological when glucotoxicity evolved, turning type II diabetes from a rare disease into the current pandemic^135^.

Type II diabetes is not a disease of ageing in-so-far as its major cause is extrinsic and dietary, but ageing-induced metabolic slowdown predisposes individuals toward reduced glucose clearance rate and increases the risk that a Western diet will induce glucotoxicity and resulting disease. Reduced β-cell proliferation rate with age may also predispose to glucotoxicity^132,195^, and would be a predicted effect of metabolic slowdown.

Hypertension will be discussed in Part 5.1, but aside from low HDL, all the criteria of metabolic syndrome could reflect direct consequences of metabolic slowdown (Figure 7). If we allow that **intrinsic ageing is continually increasing the risk of metabolic syndrome**, it allows us to fully address one of the greatest complexities of ageing.

### 4.6 The three process model: Metabolic slowdown causes extrinsic ageing

Despite ageing being a mainly intrinsic process, it has low heritability, many studies indicating values between 20-30%^226,227^, while factoring in assortative mating suggests it could be lower than 10%^228^. Such results would suggest that the rate of ageing is mainly dependent on environmental factors and behaviour. Yet our two process model predicts that ageing is an intrinsic and irreversible process induced by cell communication and epigenetic marks. It would predict a highly heritable rate that is resistant to environmental influence.

The **progression of metabolic slowdown to metabolic syndrome and associated diseases is** an explanation for the impact of exogenous factors on longevity, which we have called **extrinsic ageing**. Interestingly, exposure to ionising radiation does appear to increase risks of metabolic syndrome but the effects are small and may reflect direct effects on the IGF-1 pathway rather than through accumulated damage^229^. The major extrinsic factors affecting lifespan are not damage inducers but metabolic interventions such as diet and exercise, which are both causally linked to the nutrient accumulation processes that cause metabolic syndrome.

It has been proposed that both CR and exercise influence the ageing process and age-related disease through hormesis^229^. Because exercise generates ROS, it is viewed as a type of ‘stress’ that has lasting benefits via the upregulation of DNA repair and antioxidants^230^, thus protecting the body from additional exercise. Inevitably, these models suggest that all the effects of exercise and CR on energy regulation and the longevity module feed back into reduced oxidation, inflammation, and damage which slows ageing and prevents age-related disease^229^, as we described in Figure 3A.

However, if CR and exercise were affecting intrinsic ageing either through DNA damage accumulation, as in the hormesis model, or via reducing selective destruction, as in the two process model, then the effects should persist even after the regimen ceases – less damage should have accumulated, or selective destruction occurred – reflecting a semi-permanent change.

A crucial study examining the benefits of a dietary restriction (DR) regimen on Drosophila lifespan suggested that the reductions in mortality lasted only while flies remained on DR^231^. Flies which swapped to ad libitum at day 14 or 22 quickly adopted the same mortality profile as the control flies continuing their ad libitum diets. Similarly, flies that swapped from ad libitum to DR showed drops in mortality to the same level as flies on continuous DR, losing all negative effects of their previous ad libitum diet. The effect of diet was only on immediate risk of death, inconsistent with a single intrinsic, irreversible process.

Notably, Mair, et al.^231^ demonstrated that an intrinsic, irreversible process was also occurring, as Drosophila mortality increased in all groups independent of diet or dietary swapping. In humans, exercise has significant positive impact on healthspan, resulting in compression of late-life morbidity^232^. Similarly, in rodent studies, effects are often limited to healthspan and frailty reductions, with some studies also showing mean or median lifespan increases without effects on maximum lifespan^233–236^. A recent longitudinal study comparing different intensity exercise regimens in humans demonstrated that while higher exercise levels were associated with reduced mortality at all ages, the rates of decline in physical fitness showed little difference across intensity groups^237^. Combined, these results suggest that fitness and strength decline at the same rate, independent of exercise level ie. intrinsic ageing is happening independently. Like the dietary restricted Drosophila, where the rise in mortality simply starts later, the high exercise groups appear to have more fitness and strength to lose. Thus we suggest that exercise and DR (in Drosophila) are affecting another process which reversibly impacts short-term mortality, which is likely extrinsic ageing.

Separating intrinsic and extrinsic ageing to produce a third process could explain the low heritability of ageing, because following a Gompertzian curve, at late age when most people die, small changes in behaviour have disproportionately large impact on extrinsic ageing and associated mortality, as indicated in Figure 8A. Interventions affecting extrinsic mortality would be expected to have the most significant effect at late age, as this is when influencing short-term risk is most important. The same interventions started earlier should have little additional longevity benefit and almost none if not maintained, while the mortality reduction gained from healthy behaviour would increase as age increased, having disproportionate effects in late life.

**Figure 8.**
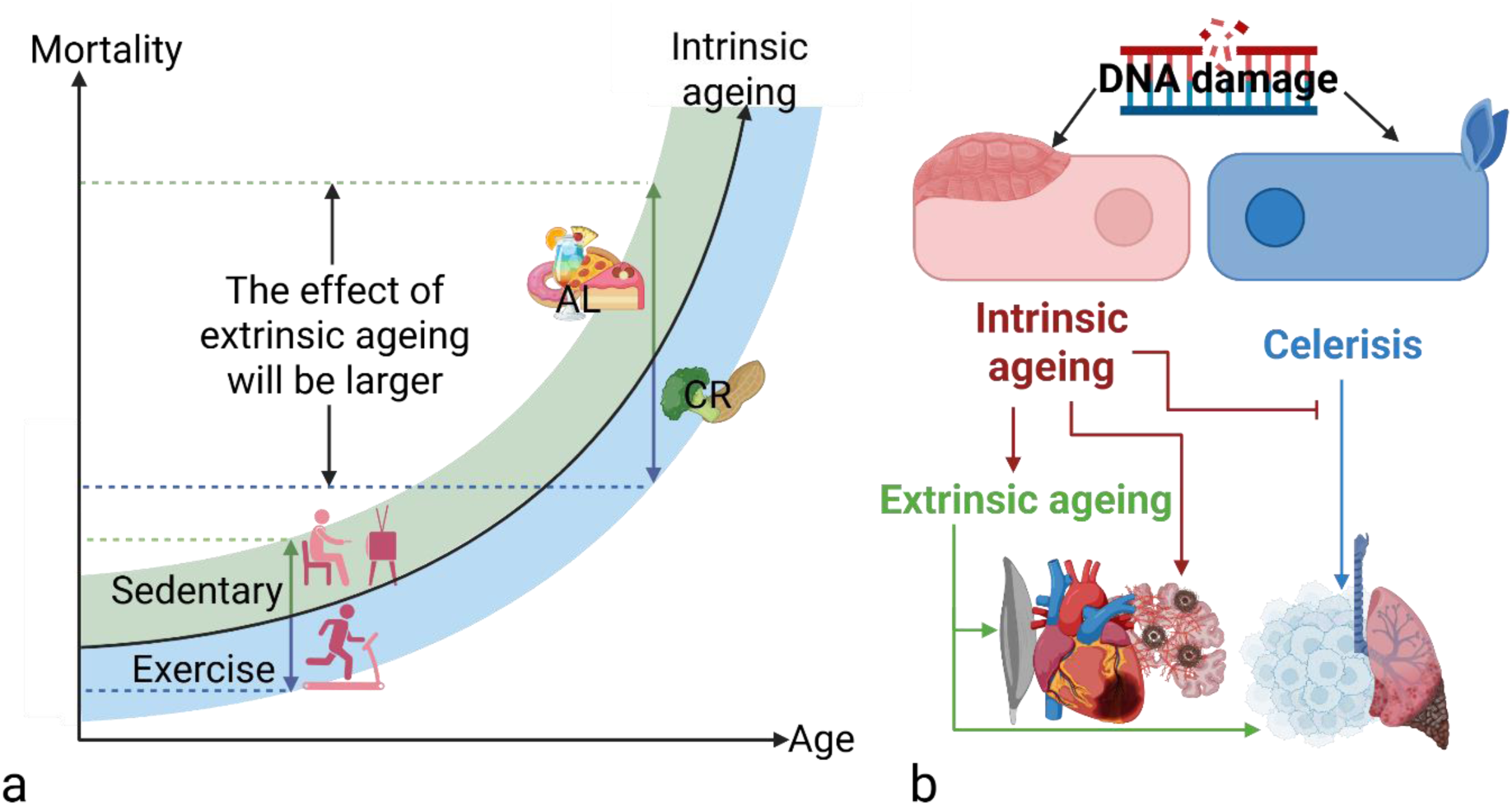
The three process model of ageing. (A) The curve of intrinsic ageing (black) increases mortality with time, relatively unaffected by behaviour. However, extrinsic ageing is highly alterable by good behaviours (blue) and bad behaviours (green), which increasingly affect a person’s mortality as intrinsic ageing accelerates. (B) The three process model. Abbreviations: AL, ad libitum; CR, calorie restriction; DNA, deoxyribose nucleic acid.

The central prediction of the three process model is that age-related decline reflects an underlying epigenetic process which is unidirectional and irreversible (at least by behavioural intervention), resulting in metabolic slowdown, which results in another process increasing risk of nutrient overload and metabolic syndrome (Figure 8B). This process should be reversible by interventions and behaviours which reduce nutrient availability.

In Part 5, we demonstrate how each of the three ageing processes contribute to the age-related diseases, completing the model of ageing from initial cause to ultimate effects.

## Part 4: Summary

4.1 | The epigenetic reduction in anabolism will manifest as reduced rates of transcription, translation, DNA replication, and cell division, reducing ATP utilisation per unit time.

As ATP is kept in strict homeostasis, reduced ATP hydrolysis is compensated for by reduced ATP production. Mitochondrial metabolism therefore drops.

4.2 | Reduced ATP production will reduce the amount of glucose and lipid catabolised in the mitochondria to produce it. These fuels then accumulate in the cell and can become toxic in excess. Homeostatic controls reduce the inflow of fuels by reducing sensitivity to insulin (**inducing insulin resistance**).

4.3 | As our non-lipogenic tissues import less fuels, our lipogenic tissues import the excess to prevent lipid and glucose build-up in the blood. The adipose tissue can store the excess fuels as lipid droplets without toxicity. Thus **inducing weight gain**.

4.4 | Our adipose tissue has its own homeostatic controls where hypertrophic cells secrete inflammatory cytokines to attract macrophages which redistribute the lipid via HDL. As lipid stores increase, and more adipocytes need recycling, this **induces a state of chronic basal inflammation** in the adipose tissue.

4.5 | As the subcutaneous adipose store becomes saturated and insulin resistant, the sensitivity differential between the subcutaneous and visceral adipose diminishes, increasing visceral lipid storage and **omental weight gain**. The inflammatory cytokines produced to recycle the visceral adipocytes can spread to the visceral organs **inducing unwanted chronic inflammation of multiple systems**.

As both adipose stores become saturated, blood glucose spikes take longer to return to equilibrium. When combined with high sugar diets, this **induces type II diabetes**.

Combined, the effects of metabolic slowdown on fuel homeostasis make up the list of symptoms of **metabolic syndrome**.

4.6 | We have termed the induction of metabolic syndrome by metabolic slowdown, **extrinsic ageing**. It is a separate process from intrinsic ageing, and can be modulated easily by behaviour to increase or decrease mortality in the short-term. This is the **three process model** of ageing.

## Part 5: Three ageing processes induce age-related disease, frailty, and death

The exact connections between ageing and age-related diseases remains controversial, with no consensus as to whether ageing is the underlying disease or whether age is merely a single risk factor among many^238^. Attempts to build causal links have focussed on which ageing hallmarks associate with which age-related diseases^97,98^. The three processes described here offer an alternative causal mechanism:

- Diseases which associate mainly with hypofunction and slower anabolism are likely mainly impacted by intrinsic ageing.
- Diseases related to metabolic syndrome could be induced by extrinsic ageing.
- Diseases of hyperfunction and overactivity could be induced by celerisis.

A crucial and surprising observation is how many age-related diseases relate to metabolic syndrome, which in the absence of our three process model would be regarded as a largely extrinsic disease induced by dietary and behavioural factors. In addition to atherosclerosis and Alzheimer’s disease, risk of cancer and fibrosis also increase^239^, which may help explain why gerontologists have retained the idea that cancer is part of a single – if complex – ageing process: a part of the process is indeed the same.

### 5.1 Intrinsic and extrinsic ageing induce atherosclerosis

Outside cancer, the most significant age-related disease in terms of incidence and mortality is cardiovascular disease^240^. While it is widely regarded as a disease of both the heart and blood vessels, it is an arterial disease, caused largely by the build-up of plaques in our artery walls, known as atherosclerosis. The hardening of the artery wall and resultant increase in blood pressure is also a contributing factor. The term ‘cardiovascular disease’ is used because the coronary arteries are frequently sites of the atherosclerotic plaques, thus mainly afflicting the heart as the major organ of dysfunction. The heart suffers a myocardial infarction when the blood vessels providing it with oxygen and nutrients become blocked. Similarly, strokes and pulmonary emboli result from plaque rupture and blockage of the arteries leading to the brain and lungs, respectively.

While we suggest L_I_ induces its detrimental effects mainly through increasing inflammation, rising L_B_ could have multiple effects on health. It is widely observed that one of the proximal events in atherosclerosis is the formation of fatty streaks in the artery wall^241^. The fatty streaks contain cholesterol and its esters within the cholesterol carrier molecules of low density lipoprotein (LDL)^242,243^. Free cholesterol can even recrystalise out of solution, and is frequently demonstrated in atherosclerotic plaques^244^.

While atherosclerosis does not normally develop in mice under laboratory conditions, ApoE knockout mice demonstrate severe hypercholesterolemia and spontaneous atherosclerosis, as do mice lacking the LDL receptor when fed a high fat diet^245^. Thus, rising L_B_ may be a proximate cause of the increases in atherosclerotic plaques with age. Although triglycerides are thought to contribute to atherosclerotic risk independently^246^, cholesterol is thought to be the central lipid component of atherosclerotic plaques^247^. It also induces foam cell formation when macrophages scavenge cholesterol without unloading the cargo sufficiently onto HDL, thus becoming engorged and dysfunctional^248^.

Both FFAs and glucose can be converted to cholesterol through ACoA. Indeed, this may be an important nutrient outlet in times of overnutrition as it has additional advantages over uncoupling. Cholesterol synthesis is a multi-step synthesis pathway that requires oxidising NADPH back to NADP, and uses ATP^249^. Additionally, as cholesterol is a necessary part of the LDs required for excess FFA storage, upregulating cholesterol synthesis is likely a key part of the homeostatic response to metabolic slowdown and the excess of FFAs.

If true, we would predict that blocking cholesterol synthesis would increase risk of metabolic syndrome and diabetes. Statins are 3-Hydroxy-3-methylglutaryl-coenzyme A (HMG-CoA) reductase inhibitors, preventing the initial steps of cholesterol synthesis which convert three acetyl groups attached to Co-enzyme A into mevalonate. This may explain why statins reduce risk of atherosclerosis, by reducing a central component of atherosclerotic plaques^123^, but increase risk of metabolic syndrome and diabetes^250^, and some studies suggest also increase both non-cardiovascular and all-cause mortality^251–253^ by removing an important fuel outlet and exacerbating the cholesterol-independent effects of extrinsic ageing.

We propose that the increased L_I_ and L_B_ levels promoted by extrinsic ageing shifts the equilibrium toward increased cholesterol synthesis, producing more LDL-cholesterol and thus the amount of deposition in fatty streaks during the conditions where they form (Figure 9, left and middle).

**Figure 9.**
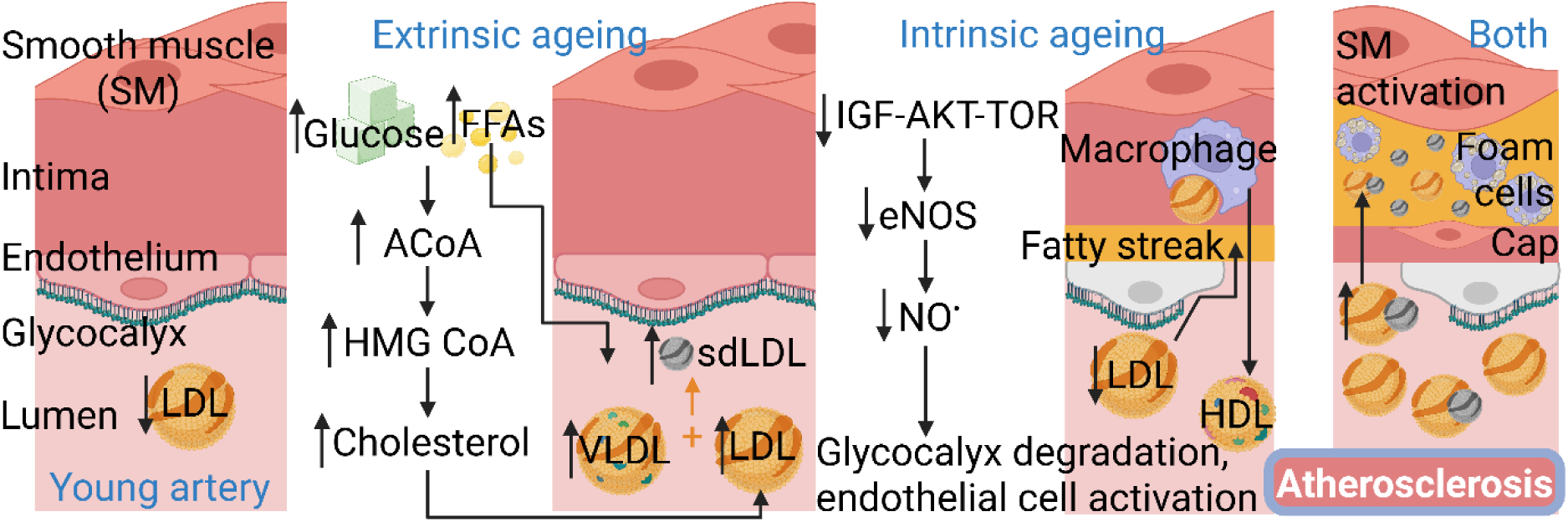
Extrinsic and intrinsic ageing combine to induce atherosclerosis. Extrinsic ageing increases the cholesterol and lipid carriers in the blood, while intrinsic ageing contributes to endothelial dysfunction. When combined, fatty streaks expand into atherosclerotic plaques where macrophages cannot successfully remove the lipid. Abbreviations: ACoA, acetyl-coenzyme A; eNOS, endothelial nitric oxide synthase; FFA, free fatty acid; HDL, high density lipoprotein; HMG CoA, 3-hydroxy-3-methylglutaryl-coenzyme A; IGF, insulin-like growth factor; LDL, low density lipoprotein; NO, nitric oxide; sdLDL, small dense LDL; SM, smooth muscle; TOR, target of rapamycin; VLDL, very low density lipoprotein.

This may be worsened because in the state of hypertriglyceridaemia associated with metabolic syndrome, cholesteryl ester transfer protein (CETP) transfers excess triglycerides from VLDL onto LDL^254^, which has lost the APOE protein required to transfer the triglycerides to tissues.

Instead, hepatic lipase removes the unwanted cargo, producing a smaller, denser version of LDL which does not so easily bind the LDL-receptor. This small, dense LDL (sdLDL) is significantly more atherogenic than normal LDL^255^, further suggesting how metabolic syndrome and extrinsic ageing may contribute to cardiovascular disease.

Although the cholesterol hypothesis is the dominant explanation for atherosclerosis, with a significant evidence backbone^123^, the disease is not simply the product of gradual LDL accumulation in the blood vessel wall which accelerates as levels of LDL increase in the blood.

When the endothelium is intact, regardless of the quantity of LDL in the blood, there is likely very little LDL passage past the vessel’s protective glycocalyx barrier – a thick extracellular matrix of proteoglycans attached to the endothelium. As suggested by the damage-centric hypothesis, plaque build-up could instead be a “response to injury” in the vessel wall^256^.

Indeed, the frequency of atherosclerotic plaques at areas of high blood pressure, such as in the coronary arteries, and the contribution of high blood pressure to plaque formation, suggests endothelial damage is likely a crucial aspect. Other significant risk factors for atherosclerosis include smoking, cocaine, and COVID-19 infection, all of which can induce endothelial damage^257^.

The damage-centric hypothesis suggests that endothelial cell death leads to low-grade inflammation and the resulting “chronic low-level insult” leads to atherosclerosis^242,258^ as predicted by the inflammageing model. However, any disease caused by inflammation could just as easily result from causes related to metabolic slowdown and extrinsic ageing as DNA damage accumulation.

Evidence suggests that inflammation degrades the glycocalyx and increases the porosity of the endothelium allowing immune cell adhesion and then crossing the vessel wall^259,260^, described as endothelial dysfunction. Thus, inflammation would induce the formation of fatty streaks, which would be more likely to become atherosclerotic plaques as LDL levels rise.

However, this model again assumes inflammation as a largely detrimental process where the body responds to injury in many contexts by exacerbating it, unnecessarily inducing disease. By making the endothelium leaky, it causes both the LDL and the macrophages to cross into the vessel wall. Fascinatingly, some major trials suggest the inflammation in our atherosclerotic plaques is a necessary and protective feature. Despite initial enthusiasm for the use of non-steroidal anti-inflammatories (NSAIDs) and other anti-inflammatories against atherosclerosis, multiple trials and studies demonstrated that frequent use of ibuprofen, naproxen, and celecoxib, among others, *increased* rather than decreased risk of atherosclerosis, stroke, and myocardial infarction^261,262^. The inflammation so frequently associated with atherosclerotic plaques was perhaps not causing them but contributing to remodelling and clearance.

Another inflammageing-based narrative may also be inconsistent with the evidence. It has been suggested that LDL itself is not atherogenic because after macrophages intake LDL they downregulate LDL-receptors (LDL-Rs) until the lipids have been processed, thus preventing lipid overload and foam cell formation^263^. Instead it is oxidised LDL, thought to result from inflammageing and increased presence of ROS, which is atherogenic because it upregulates the macrophage LDL-R^263,264^, preventing the negative feedback that stops macrophages absorbing lipid before they become foam cells. However, as with much of the ROS-associated literature, these conclusions engender significant complexity, as overexpression of the macrophage enzyme 15-lipoxygenase which promotes LDL oxidation^265^ actually *protects* rabbits from atherosclerosis^266^. Conversely, this would be the predicted result if LDL-oxidation is instigated by macrophages in response to quantities of LDL-cholesterol that cannot be redistributed without inhibiting LDL-R downregulation. By this hypothesis, foam cell formation and atherosclerosis occur not because of excess LDL oxidation, but because LDL oxidation is insufficient to remove the excess LDL and return L_B_ homeostasis. Inflammation would not be promoting foam cell formation, but preventing foam cell formation by removing extra LDL.

The net effects of inflammation may depend on the conditions, and when associated with autoimmune diseases such as lupus – where vascular dysfunction is a direct effect of a self-reactive immune system^267^ – inflammation likely contributes to disease, but in plaques associated with ageing we suggest that the immune cells could also have a functional role in plaque resolution by removing the cholesterol, degrading the fatty streak as the endothelium repairs.

Of course, inflammation is an umbrella term for multiple processes, some of which are likely detrimental to endothelial cell function, but the consistent presence of pro-inflammatory factors in atherosclerotic plaques, which has been taken as evidence of causation, is potentially a net-protective countermeasure.

Another effect of extrinsic ageing is IR. IGF-1-stimulated PI3K-Akt signalling activates endothelial nitric oxide synthase (eNOS)^268^, which produces nitric oxide (NO), one of the main regulators of endothelial cell functionality and survival^269,270^. It maintains the glycocalyx, inhibits immune cell adhesion and passage, and reduces the leakiness of the artery wall toward LDL^271,272^. IR would be expected to contribute to endothelial dysfunction if it prevented PI3K-Akt signalling. Thus, one narrative is that IR ‘is the etiologic factor responsible for the development of each of the individual cardiometabolic disturbances’^273^. In effect, IR would prevent nutrient uptake^112^, making it a ‘maladaptive response’^274^. If so, then cardiovascular disease could be treated by overcoming IR. However, we suggest here an alternative narrative that cells do not become too insulin resistant to import sufficient nutrients, but become resistant enough to prevent excess import in conditions of nutrient excess. Indeed, others have suggested that IR is a defence mechanism against metabolic stress in patients with type II diabetes^275^.

In such context, the cause of endothelial dysfunction is not IR but intrinsic ageing. It is the reduced PI3K-Akt signalling itself that results from intrinsic ageing which both reduces NO production and increases IR.

Thus, increasing L_B_ is likely the major proximate cause of disease induced by extrinsic ageing, while intrinsic ageing slowly shifts the equilibrium of endothelial cells toward the activation of pathways that induce a leaky barrier prone to immune cell-adhesion. Other factors such as smoking, cocaine, and COVID infection contribute to disease by activating these same pathways, plausibly activating them more strongly with additional intrinsic ageing. Particularly at areas of high pressure, the endothelial barrier may allow lipid carriers and immune cells to cross forming fatty streaks, and when combined with extrinsic ageing these streaks are more likely to become plaques (Figure 9).

Interestingly, for the TCQP, Blagosklonny predicted an opposing mechanism, stating, “atherosclerosis depends on hyperfunctions of local cells [endothelial cells, smooth muscle cells (SMCs), and macrophages] and distant cells (hepatocytes, bone marrow cells, and adipocytes)”^276^. However, we would argue that many of these phenotypes are not true hyperfunction. Endothelial cell ‘activation’ is not activation in the same way as occurs in other cell types, promoting growth, proliferation, and activity. It refers to a pro-apoptotic, pro-inflammatory state^277^ which is inhibited by growth stimuli such as IGF-1^272^, and thus we suggest it may be more reflective of hypofunction from intrinsic ageing. The hypertrophy of distant adipocytes and hepatocytes which “increase levels of […] LDL and procoagulation and pro-inflammation factors”^276^ can be explained by extrinsic ageing, which again results from hypofunction. The activation of immune cells and hyperplasia of SMCs could be true hyperfunction: it is possible that plaque generation is aided by localised celerisis inside a body that is undergoing metabolic slowdown. Most evidence suggests that SMCs promote the stability of the plaque by proliferating and migrating into the core, switching from contractile to synthetic subtype to form the fibrous and later calcified cap which prevents the plaque from rupturing^278^. However, SMCs also contribute to pathogenic plaque enlargement, leading to high blood pressure and angina, potentially also promoting a thrombogenic event and blood vessel occlusion. High rates of cell death and proliferation at sites of high blood pressure could lead to celerisis, and hyperfunctional SMCs could further promote cardiovascular disease.

There are likely many additional factors important in atherosclerosis, and this is not intended to be a complete account. Considering the relevance of aspirin as an anti-coagulant and lipoprotein(a) (LPa) as a clotting factor in atherosclerosis^279^, we suggest that the thrombogenic cascade would be the most crucial addition. The simplified mechanism presented here and those that follow for other age-related diseases are intended to suggest the most crucial factors that connect the process of ageing with eventual disease outcomes. This provides a testable hypothesis that can be rejected or adapted by further experimentation.

### 5.2 Intrinsic and extrinsic ageing induce Alzheimer’s disease

Alzheimer’s is currently thought to reflect the accumulation of some combination of amyloid-β plaques and tau tangles (hereon also referred to as plaques). The hypotheses suggest that amyloid-β peptide and tau phosphoprotein monomers aggregate, first forming oligomers and then plaques^280^. It is thought that both the oligomers and plaques are toxic to neurons, promoting inflammation and cell death^281^; however, additional mechanistic detail is still controversial and split between proponents for amyloid-β and tau as the most significant disease-causing aggregate.

Notably, Gems and McElwee^16^ suggested that protein damage showed more connection to longevity mutants than DNA damage, but suggested that protein damage accumulation could not be the primary cause of ageing as the molecules were insufficiently long-lived. The difference between damage and clearance would have to be minute to explain gradual accumulation in long-lived organisms, and small shifts in the relative rates of damage and clearance (which could be reasonably frequent) would be predicted to result in accelerated ageing and rapid rejuvenation, respectively. Accumulation-based models thus required a macromolecule that lasted roughly the lifespan of the organism, limited to a few long-lived structural proteins such as collagen and elastin in specific joints^282^, and DNA. Thus, while protein damage may play a significant role in ageing, it is more likely a downstream effector responding to homeostatic shifts. An idea put forward in Green Theory is that temporary build-up of ‘molecular rubbish’ could cause an increase in DNA damage, and ageing^16^; however, such mechanisms still struggle to explain why inducing high levels of oxidative damage, usually measured at the protein level, show little or contradictory effects on ageing. Here we suggest an alternative.

The equilibrium levels of amyloid-β should reflect the rate of production, by β-secretase and subsequent γ-secretase cleaving the amyloid precursor protein (APP), in relation to the rates of autophagic and proteasomal degradation. Early onset Alzheimer’s disease results mainly from mutations either in APP itself or in the enzymes and associated proteins which produce amyloid-β from APP^283^. Early onset disease appears to reflect that the amyloid-β produced is more prone to aggregation. Thus, rates of monomer aggregation are likely causative to disease.

One model suggests that elevated rates of monomer aggregation causes gradual plaque accumulation^284^. Like models of ageing by DNA damage accumulation, no equilibrium shifts are required, only continued production of aggregate-prone monomers and the passage of time.

However, we suggest this model is still insufficient because it does not account for negative feedback and homeostasis. All else being equal, increased monomer aggregation should result in increased aggregate degradation. Traditional models have relied on chronic inflammation and associated positive feedback to break the homeostasis, with fibrils and plaques inducing inflammation which creates more plaques, known as the amyloid/neuroinflammation cascade hypothesis. In some renditions, the body supposedly orchestrates some fairly catastrophic cascades, such as amyloid-β plaques activating microglial cells which induce disease by both killing neurons and even producing more amyloid-β^285^. Why evolution would produce such systems requires a mechanism of significant complexity.

Without positive feedback and self-destructive signalling, fibrils and plaques should not gradually accumulate in a homeostatic system purely through the passage of time – only when the equilibrium shifts in favour of persistence. Yet this distinction has significant implications for Alzheimer’s therapies.

Therapies designed specifically to remove plaques work best under the gradual accumulation model: removing the plaques resets the problem because it will take a long time for the plaques to return. As suggested by de Grey, animal tissues treated with alagebrium to break advanced-glycation end product (AGE) crosslinks “become inflexible again so quickly after withdrawal of the drug” because “the underlying cross-links are clearly reforming much more rapidly than they did over the many years that were required for their initial buildup”, which he attributes to the exposure of “highly reactive carbonyl” groups by the action of the drug^286^. This may well be true, but we would also suggest that plaques have not simply been building up prior to drug treatment. Removal of aggregates with alagebrium may also result in a downregulation of aggregate removal pathways, which will slowly reactivate as the plaques return to their equilibrium levels once the regimen has ceased.

If true, we would suggest that neither early onset nor late onset disease can result simply from additional monomer aggregation because homeostasis will reduce the aggregated monomers back to equilibrium via increasing their clearance. Any successful model of disease should explain why increased fibril and plaque production is not counteracted by negative feedback through degradation. The inflammageing model could still predict disease even in the absence of positive feedback, if DNA damage, cell senescence, inflammation, and ROS, raised aggregate production and decreased degradation. This problem was partly solved by the garbage catastrophe theory and the ‘mitochondrial-lysosomal axis theory of cellular aging’ which suggested that increased levels of protein damage would obstruct clearance^287,288^. Although these models are often described as gradual accumulation models, they also reflect feasible homeostatic models predicting equilibrium shifts.

However, the garbage-catastrophe theory does not explain why the equilibrium should shift in favour of plaque accumulation prior to the point of catastrophe. Even in individuals with defective APP and amyloid processing, the mutations do not, themselves, increase the rate of plaque production over time without additional metabolic shifts – the base rate is simply higher than in individuals without the mutations. Assuming at young age the rate of aggregate formation has not already overwhelmed the degradation machinery, then without additional equilibrium shifts it should never do so.

To generate these shifts, inflammageing is reliant on either DNA damage accumulation or positive feedback between protein damage and inflammation. Initially in favour of these ideas, multiple studies demonstrated that use of non-steroidal anti-inflammatories (NSAIDs) for conditions such as arthritis was associated with reduced risk of Alzheimer’s disease. These results prompted multiple clinical trials which all showed NSAIDs provided no benefit to

Alzheimer’s disease patients^289,290^. Inevitably, many concluded that the failure resulted from the drugs not being given early enough, leading to the ADAPT trial in elderly dementia-free patients of the NSAIDs naproxen and celecoxib, which was terminated when results of another trial suggested celecoxib significantly increased risk of cardiovascular complications^261^. However, preliminary results of ADAPT suggested that NSAIDs *increased* risk of Alzheimer’s disease^291^.

Finally, the INTREPAD trial showed no benefit in an even younger group with a family history of Alzheimer’s disease^292^. As stated by the lead researcher, John Breitner, “There was a lot of hope, based on epidemiological data, that these drugs would prove beneficial, but we have learned over the last 10 to 15 years that when you test NSAIDs in trials, they are not beneficial and invariably, you make people sick”^293^.

We suggest that inflammageing, positive feedback, and the garbage catastrophe are unlikely to cause Alzheimer’s disease. Even newer models of the neuroinflammation cascade which suggest that ageing shifts microglial cells from protective to detrimental^285,294^ do not explain why NSAIDs are not beneficial if the inflammatory cells are still the ones perpetuating or inducing most of the plaque production and cell death.

We suggest that while inflammation likely has detrimental effects, just as with atherosclerosis, its consistent association with plaques and disease likely reflects the immune system’s role as a crucial defence mechanism, removing the plaques. Just as macrophages and ROS are blamed for forming foam cells, microglial cells appear only to produce amyloid-β plaques under similar conditions of exhaustion^294^, which likely reflect a combination of intrinsic ageing of the immune cells, reducing their ability to clear the problem molecule, along with an excess of the problem molecule. Chronic inflammation results because the immune system is insufficiently active to correct the problem, not because an overactive immune system is inducing the problem.

If not inflammation, Alzheimer’s is likely caused by another aspect of extrinsic ageing, as evidenced by its association with metabolic syndrome and diabetes, even being referred to as ‘type III diabetes’^295^. However, its association with IR^296^ has often been interpreted as a failure of glucose supply, starving the brain of nutrients and reducing metabolism, potentially inhibited by amyloid-β plaques^297^. This has led to the use of insulin stimulation as therapy^298^. Some studies have shown glucose infusion and insulin stimulation have short-term benefits, but conversely chronic hyperglycaemia and insulin injections are associated with memory deficit and increased cognitive impairments^299,300^.

We would predict the short-term benefits because the additional insulin or glucose is temporarily overcoming the IR induced by intrinsic ageing, increasing brain anabolism and restoring some function. However, according to the three process model, the IR is not an external block preventing neurons from getting enough nutrients, but an internal block protecting them from nutrient overload. We would thus predict that anabolic stimulation therapies should increase IR in brain cells as they protect themselves from the overactive growth signal to restore their preferred equilibrium, also potentially inducing gluco- and lipotoxicity. Thus, these therapies fail not because of the irrelevance of metabolism or metabolic syndrome, but because brain ageing results in IR rather than IR resulting in brain ageing.

However, Alzheimer’s could still be induced or exacerbated by other effects of extrinsic ageing. It is associated with dyslipidaemia^301^ as well as UPR failure and endoplasmic reticulum stress^302^ which can be exacerbated due to higher lipid levels. Particularly in the brain, where absorption of glucose and nutrients is mainly insulin-independent^303^, extrinsic ageing may result in nutrient overload within neurons. This energetic stress can induce the aberrant attachment of glucose and lipid to amino acids forming glycated and lipidated proteins, which can aggregate to form AGEs in ageing neurons^304,305^. While this has been traditionally viewed as the result of ‘stress’, ROS, and inflammation^306^, these types of molecular ‘damage’ could also be described as products of metabolic syndrome – the natural consequence of excess sugars and lipids which result from extrinsic ageing^307^. Although amyloid-β and tau might be the main proteins forming aggregates, others can be bound together in AGEs which colocalise with amyloid and tau in plaques^304^. Thus, extrinsic ageing could contribute to Alzheimer’s development by increasing the probability that amyloid-β and tau monomers and oligomers aggregate with AGEs and other glycated and lipidated proteins into amorphous plaques.

However, as noted above, monomer aggregation is only half of the equation, with negative feedback restoring equilibrium via the upregulation of degradation pathways. Unfortunately, while extrinsic ageing may increase the rate of monomer aggregation and thus plaque production, we suggest that intrinsic ageing is potentially also slowing plaque degradation by decreasing autophagy.

Autophagy is thought to play a central role in ageing and neurodegeneration^308^, and Alzheimer’s disease specifically^309–311^, and reductions in autophagy are known to occur across multiple ageing models^312^. It is worth observing that decreased autophagy is not an immediately intuitive effect of metabolic slowdown. Just as obesity suppresses autophagy by activating pro-growth stimuli which inhibit autophagy through mTOR^313^, reduced mTOR signalling through metabolic slowdown might be predicted to have the opposite effect.

We suggest that metabolic slowdown blocks autophagy despite the reduction in IGF-mTOR axis signalling in part because it slows down total metabolism, reducing both the anabolic and catabolic metabolism in conjunction, maintaining energy equilibrium rather than shifting it toward a catabolic state. As neurons (and other cells) reduce their anabolic rate, this should manifest as a reduction in the rates of protein translation, as is observed in rats and mice^314^, yeast^315^, as well as humans and other model organisms^316,317^. However, as protein synthesis is reduced, cells will take longer to produce sufficient protein to carry out the task the protein is produced for (Figure 10A). The natural result is that cells will delay and deactivate negative feedback systems to accomplish their cellular functions.

**Figure 10.**
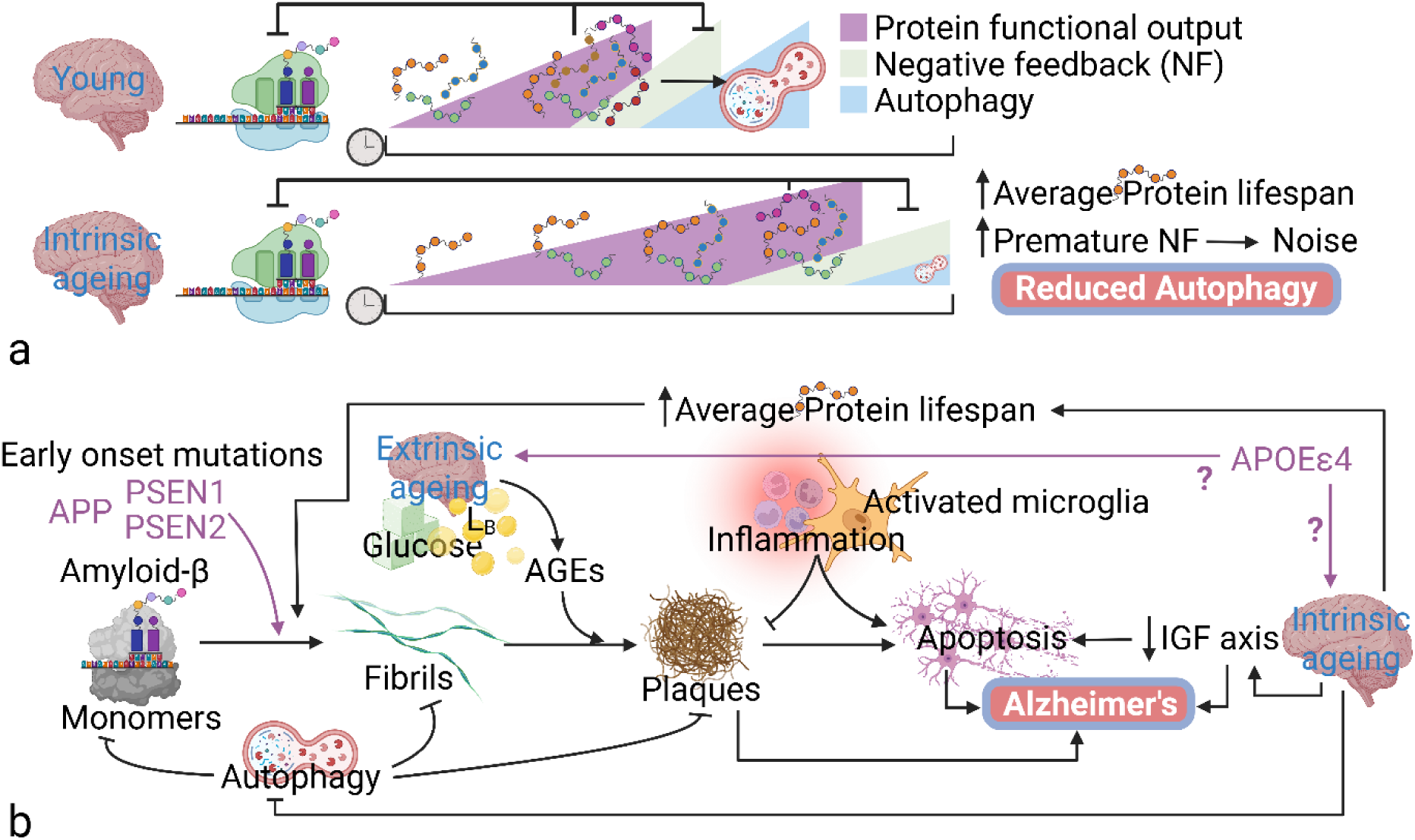
Metabolic slowdown reduces autophagy and induces Alzheimer’s disease. (A) Protein synthesis rate declines due to intrinsic ageing, which increases the time required for the protein to produce its function, and thus induce negative feedback. The chain of reduced anabolism followed by catabolism decreases the amount of autophagy needed per unit time. In older individuals, prolonged low-level protein activity may even induce premature negative feedback and thus noise. (B) A mechanistic model for early and late onset Alzheimer’s disease. Early onset disease is induced partly by mutations which cause increased monomer aggregation. As individuals undergo intrinsic ageing it slows the rate of monomer and aggregate degradation via autophagy, which reduces the cell’s capacity to degrade the monomers and aggregates, shifting equilibrium toward prolonged monomer lifespan and chances of aggregation, further aided by extrinsic ageing and metabolic syndrome. Plaques are mainly degraded by microglia requiring immune involvement and inflammation, which can trigger apoptosis. Alzheimer’s disease is caused by a mixture of factors. The direct effects of metabolic slowdown will reduce the capacity and rate of brain cell function; both plaques and the immune response will induce cell death, reducing the number of functional cells. Image created with BioRender.com. Abbreviations: AGE, advanced glycation end-product; APOE, apolipoprotein E; APP, amyloid precursor protein; L_B_, blood lipid; NF, negative feedback; PSEN, presenilin.

All pathways and processes have their own specific negative feedback systems which inhibit protein activity once its function is complete (and further activity would be deleterious), but all these specific feedback systems share a single general negative feedback system, which is the apparatus of protein degradation. The rate at which all negative feedback systems operate should therefore depend partially on the level of activity of the proteasome and autophagy pathways. The more active these pathways are, the faster negative feedback takes effect; thus, intrinsic ageing and metabolic slowdown should naturally deactivate autophagic and proteasomal activity to compensate for reduced metabolic rate, allowing proteins to carry out their function before degradation, as shown in Figure 10A.

While the garbage catastrophe theory suggests that autophagy becomes impaired with age due to the build-up of aggregates and molecular damage inhibiting the autophagic system, most evidence suggests instead that autophagy is increasingly deactivated^318–320^, and this is at least partially epigenetic^321^. As a result, autophagy can be re-stimulated with pro-longevity and anti-neurodegenerative effects^319^, less consistent if the decline represented genetic or aggregate-induced dysfunction. Thus, we suggest that intrinsic ageing and metabolic slowdown reduce autophagy (see Part 7 for the significance of mitophagy), reducing the removal of aggregate-prone proteins such as amyloid-β and phosphorylated tau. As average protein lifespan increases to carry out their necessary function, so does potential for glycation and lipidation, which encourages plaque formation. As homeostasis shifts in favour of plaque formation, the same shifts are reducing plaque degradation, leading to a higher equilibrium level of plaque.

Notably, the role of plaques in Alzheimer’s disease is increasingly challenged in the literature, in part due to the failure of plaque removal therapies^322^, as well as studies suggesting that cognitive impairment and Alzheimer’s can occur in individuals with low or absent plaque burden^323,324^, while moderate plaque burden can be present in individuals without evidence of disease^325^. Thus, plaques appear neither sufficient nor necessary for disease.

However, the significance of amyloid-β processing mutations in early onset Alzheimer’s suggests that the pathway is of significant relevance to disease, and can induce disease with high penetrance, even if dependent on other factors. A major factor independent of plaques is likely the degree of intrinsic ageing of the neurons, where high levels of anabolism are required for brain function and memory, which are thus directly reduced by metabolic slowdown.

Indeed, hypometabolism is a widely recognised feature of brain ageing and Alzheimer’s disease that does not always correlate well with amyloid-β levels^326^. Consistently, while CR is well known for its beneficial long-term effects on cognition, studies also suggest that it can reduce cognitive flexibility^327^, just as glucose infusion stimulates brain function^299^, likely reflecting shifts in metabolic rate. Decreases in cerebral blood flow^328^ reflecting the atherosclerotic conditions of cerebral blood vessels^329^ may also be key factors reducing metabolic activity in the brain. The hypometabolic state of neurons induced by intrinsic ageing may be the crucial process which reduces cognitive function, which combines with neuronal loss and signal interference from plaques to induce or exacerbate Alzheimer’s disease symptoms. As suggested in Figure 10B, the three process model suggests how ageing could result in plaque formation, Alzheimer’s disease in the absence of plaques, and how plaques could contribute significantly to disease. Of course, we are not suggesting the model is complete, or even wholly accurate, only that it provides a testable null hypothesis that is superior to gradual accumulation-based and existing equilibrium-based models that link ultimate which centralise plaque generation as the major disease-inducing process.

### 5.3 Extrinsic ageing induces inflammageing, autoimmune diseases, fibrosis, and cancer

The immune system declines with age in part through the rapid shrinking of the thymus, reducing T cell maturation with age, and resulting in increased autoimmunity and susceptibility to infections. While the underlying reason for thymic involution being widespread in vertebrates^330^ despite its negative impact on health is still controversial, it increasingly appears as a product of metabolic slowdown and metabolic syndrome, with the build-up of lipid around the shrinking thymus^331^.

As T cells activate B cells, thymic involution results in the accelerated senescence of the adaptive immune response, forcing individuals to rely increasingly on their innate immune systems. The intrinsic ageing of both the adaptive and innate cells will also reduce their capacity (per cell) to fight infection and carry out the other functions of immune cells. As we suggest here, these functions include the clearance of hypertrophic adipocytes for adipose homeostasis, removal of lipid from atherosclerotic plaques, and aggregated proteins from brain plaques. As immune cells age and the adaptive response falters, it forces greater reliance on the innate immune system, which not only induces greater levels of inflammation, but requires greater levels of inflammation to function appropriately. The resultant almost constitutive presence of inflammation at sites of disease, which has been interpreted as evidence of causation, may thus be much closer to a defence mechanism.

Yet, of course, inflammation is not a purely benevolent process. It is a complex network of factors with multiple functions. When inflammation is triggered, some of the multifactor network of co-regulated processes are going to be unwanted and potentially detrimental under any given conditions. We are not going to cover these complexities here. However, there are also likely to be some generally detrimental outcomes of chronic inflammation. For example, the reason visceral adiposity is associated with morbidity and mortality, while subcutaneous adipose tissue is not^332^, may be largely positional. The short-lived inflammatory cytokines produced to remove subcutaneous adipocytes degrade before they reach key organs^333^.

Increased levels of subcutaneous adipose protects against metabolic syndrome^334^, but as the visceral adipose expands and becomes inflamed, the cytokines spread to and inflame the closely situated liver, pancreas, and gut, shifting their homeostasis^335^. Thus, chronic low-grade inflammation may spread from where it is a key component of metabolic health to a destructive force.

One predictable outcome of visceral inflammation that we have already described is fibrosis. While intrinsic ageing reduces the risk that celerisis (the process by which faster mutant fibroblasts spread) will produces a self-sustaining population of hyperfunctional cells, the other key inducer of fibroblast activation is the presence of immune cells and the positive feedback induced between them (Figure 5D-E). Thus, although intrinsic ageing will reduce the risk of hyperactive fibroblasts spreading and reaching the threshold for self-sustaining pathological fibrosis, extrinsic ageing and inflammageing will increase the risk that normal (or even aged, hypofunctioning) fibroblasts reach the threshold due to a continuing activation signal, as shown in Figure 11. Anti-inflammatories are currently the major therapies for fibrotic diseases such as cystic fibrosis and IPF, suggesting that chronic inflammation is likely an underlying cause. Indeed, fibrosis of visceral organs is associated with visceral obesity^336^.

**Figure 11.**
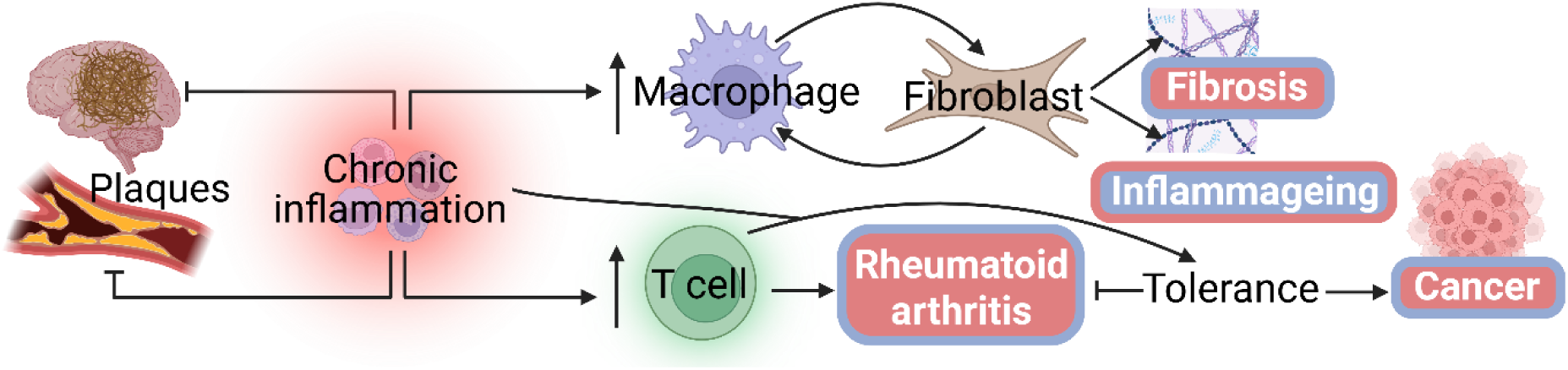
Model of inflammageing and contribution to disease. Image created with BioRender.com.

Another potential effect of increased immune activation in non-infected tissues is autoimmune disease. Auto-reactive T cells would be more likely to enter environments of sufficient inflammation around their self-antigen to trigger activation and a full pathogen response to the body’s own tissue. Rheumatoid arthritis is a common age-related autoimmune disease with incidence increasing up until around age 85^337^, and successful treatment with anti-inflammatories suggests that inflammation is a causal factor in disease. However, many autoimmune diseases, such as type I diabetes, multiple sclerosis, psoriasis, and Crohn’s disease have peak incidences either in youth or younger adults before the age of 40^338^. Notably, several have bimodal incidence with an additional late-onset peak that is more consistent with an age-related disease^338^.

It has been observed that while autoantibodies increase with age, autoimmune diseases were rarer and frequently milder in the elderly^339^. One possibility is that negative feedback is counteracting the increased likelihood of autoimmune activation associated with chronic inflammation, such as through the age-related expansion of regulatory T cells (Tregs) which repress the activity of T cells^340^. This negative feedback promotes immune tolerance at chronically inflamed sites, which could allow dangerous cells to proliferate under these conditions, necessitating that the body makes a trade-off between autoimmunity and cancer^339^. This could explain why inflammageing and other inflammatory diseases such as Chron’s associate with cancer^341^.

Thus, we suggest that intrinsic ageing reduces the risk of cancer, while both celerisis and extrinsic ageing increase the risk of cancer, in the latter case through chronic inflammation and tolerization. This would suggest that sites of age-related cancers should be associated with lower epigenetic age, localised hypermetabolism, extrinsic ageing, and chronic inflammation.

In the liver, hepatocellular carcinoma (HCC) is overwhelming associated with chronic liver disease, either from excess ethanol, in alcohol-related liver disease (ArLD), or fructose and caloric intake in metabolic dysfunction-associated steatotic liver disease (MASLD). As both fructose and ethanol are necessarily metabolised by the liver they produce disproportionate increases in visceral lipid storage^342^. We suggest the hypertrophic or ‘ballooning’ hepatocytes are then removed by macrophages as we described before for the adipocytes in the adipose, producing cycles of cell death and proliferation in a chronically inflamed background.

This model does not rule out a role for inflammation and the associated ROS in the conventionally predicted positive feedback cycles between inflammation and damage that could increase the mutation burden^342^. Unlike ageing, where most evidence suggests increasing DNA damage has no significant effect, the causal evidence for DNA damage and genomic instability in cancer is unequivocal. Whether or not inflammation and ROS significantly impact mutation rates is beyond the scope of this article; however, we suggest that the pro-growth mutations which induce cancer do so most successfully when the following conditions are true:

1. The cell itself has already been mutated knocking out either its ability to autonomously detect its own damage or its ability to kill itself/senesce when damage is detected.
2. The tissue is in a state of rapid proliferation due either to organ growth or reduced cellularity which increases metabolic rate of non-damaged cells and reduces selective destruction.
3. There is a background of chronic inflammation forcing the local immune system to become tolerant to self-antigens which prevent the detection aberrant cells by the immune system.

Loss of these systems allow the cell to escape the autonomous, juxtacrine, and immune surveillance systems. It is worth noting that the RAS titration system developed by Chan, et al.^136^ identified a cluster of RAS expressing cells which induced tumours independent of Notch1 through a non-canonical Notch ligand, Dlk1. While Notch1 tumours were associated with immune cell invasion, Dlk1 tumours were largely absent immune cells. While there are many interpretations of these results, selective destruction and immune surveillance may be more interlinked than we have thus far suggested. Plausibly, it is via cell communication and competition that cells within tissues are forced to express markers identifying them to the immune system.

Given that the three process model suggests that intrinsic ageing is, in part, an anticancer and anti-fibrosis mechanism, which then promotes cancer and fibrosis through extrinsic ageing, it is worth observing that there are differences between the cancers of youth and old age^343,344^.

Paediatric cancers grow faster than adult cancers with higher rates of metastasis^345^. These aggressive cancers might be the norm throughout life in the absence of intrinsic ageing, occurring earlier and in more organs.

Additionally, while cancer and fibrosis are the most observable outcomes of celerisis, selective destruction may also be an important epigenetic tool for maintaining uniformity of metabolism, preventing a mosaic of metabolic rates and outputs across a tissue that make it more difficult to produce an accurate response to stimulus.

### 5.4 Intrinsic ageing induces osteoporosis, sarcopenia, and frailty

In older models, such as Blagosklonny’s TCQP, diseases like osteoporosis were explained as hyperfunctional disorders due to the overactivity of osteoclasts which resorb bone^276^; however, these models did not account for the hypofunction of osteoblasts which produce bone. Others have suggested IR is the key factor, promoting stem cell differentiation to osteoclasts while inhibiting osteoblast differentiation^346,347^ as well as reducing the lifespan of osteoblasts^348^.

Such a shift could be explained by extrinsic ageing, but as we have suggested for other age-related diseases, the IR is not induced upon the cell but by the cell as a consequence of intrinsic ageing. In a healthy, youthful body anabolism is induced whenever the tissue needs additional activity and/or growth by the insulin/IGF-mTOR axis, but inducing both osteoblasts and osteoclasts is counterproductive, so organs may solve this problem by having the activity of anabolic cells repress the activity of catabolic cells, dependent on the central insulin/IGF-mTOR axis for growth and activity. The result is that as anabolic activity decreases it naturally affects anabolic cells more than catabolic ones, shifting the equilibrium toward increased catabolism. In bone, this produces osteoporosis.

It is important to note that in Part 5.2 we stated that anabolism and catabolism are reduced in combination, but this was on a per cell basis. Each cell would maintain its ATP levels by reducing its catabolism to match its anabolism (or vice versa in post-mitotic tissues, see Part 7). However, cell types less responsive to insulin/IGF signalling, or whose metabolism is repressed by insulin/IGF signalling, should be less slowed by intrinsic ageing. Thus, on a per tissue basis there could still be a shift toward catabolism.

Primary sarcopenia is central to age-associated frailty^349^, as is the mass-independent loss of strength, termed dynapenia^350^. Both could result directly from the 2-3% decline in basal metabolic rate in muscle every decade after age 20, and 4% after age 50^169^. Likely, the declining muscle mass with age is a significant contributor to the reduction in fuels used and thus extrinsic ageing^351^.

Loss of muscle and bone are the two central inducers of frailty in the elderly, and both could directly result from slowed metabolism. In many tissues, metabolic rate is tied directly to their specific functional output (i.e. cells increase generalized metabolism and often proliferative rate when required to produce more of a specific protein as this is the most efficient way to regulate output). For example, when β-cells are required to produce insulin, the insulin production and release pathway triggers both elevated metabolism and proliferation. Thus, as suggested by Karin and Alon^132^, decreased cell proliferation rate with age might predispose individuals to type II diabetes as the β-cells are more likely to be overwhelmed.

Additionally, our reduced stress responses including the antioxidant^352^ and DNA damage response^353^ – often touted as evidence of ageing by damage accumulation^354^ – could instead reflect reduced TOR-driven protein translation^355^. If true, we accumulate damage because we age rather than vice versa.

## Part 5: Summary

5.1 | Risk of cardiovascular disease increases from both intrinsic and extrinsic ageing.

Chronic inflammation may have detrimental effects, but is likely a net-benefit in preventing cardiovascular disease.

Intrinsic ageing likely reduces the pro-growth/survival signalling to endothelial cells via nitric oxide generation. Increased endothelial cell dysfunction and death results in the formation of fatty streaks to protect and repair the vessel wall.

Extrinsic ageing increases blood lipid, depositing extra LDL and sdLDL in fatty streaks that result from endothelial dysfunction, overwhelming the macrophages trying to recycle the lipid. As a result, fatty streaks become atherosclerotic plaques.

5.2 | Alzheimer’s is viewed as a disease of gradual plaque accumulation. However, plaque level should reflect an equilibrium between plaque production and degradation, increasing only if production is not compensated for by degradation. Most of the existing models, including inflammageing, do not fully predict why plaque burden should increase with age.

If metabolic slowdown increases the time required for translation to produce sufficient protein to carry out cellular functions, then negative feedback and protein degradation should also be delayed to allow their completion. Protein lifespan should therefore increase, shifting the equilibrium toward increased aggregation, while at the same time reducing the degradation systems responsible for oligomer breakdown.

Extrinsic ageing may contribute to plaque formation by the production of AGEs and other products of nutrient overload that can aggregate with amyloid and tau oligomers.

Disease may be partially dependent on plaques, with increased burden contributing to neuronal death and dysfunction, while immune activation and plaque clearance may induce additional cell death. However, intrinsic ageing may also directly reduce the capacity of neurons to carry out their function, contributing independently to neurodegeneration.

5.3 | Inflammageing will contribute to fibrosis by increasing the positive feedback between immune cells and fibroblasts.

Inflamed tissues increase risk of activating T and B cells into a full autoimmune response, increasing the risk of autoimmune disease. To compensate, chronically inflamed sites could induce immune-tolerance mechanisms which increase risk of transformed cells escaping detection and causing cancer.

5.4 | Catabolic cells are less responsive to insulin and IGF. They are activated by anti-growth signals that suggest there is enough or too much of something. Therefore as our bodies become insulin resistant it shifts metabolism in favour of increased activity of catabolic cells such as osteoclasts compared to osteoblasts, causing bone loss and osteoporosis.

Sarcopenia could result from the intrinsic ageing of muscle cells, reducing the strength of the tissue. These two diseases then increase frailty with age.

## Part 6: Therapeutic interventions

Homeostasis produces a more nuanced interpretation of the context-dependent molecular effects of interventions than either ‘pro-ageing’ or ‘anti-ageing’, but here we try to demonstrate the implications of the three process model for anti-ageing therapies.

### 6.1 Body mass, CR, and the obesity paradox

In 2007, Blagosklonny coined the term ‘protein synthesis paradox’ to reflect that both ageing and anti-ageing interventions reduce protein synthesis^356^, which in 2023 was still suggested as a “substantial barrier to fully understand how protein translation regulates aging”^317^.

The three process model suggests intrinsic ageing results in reduced protein synthesis and the negative effects described above. However, CR also inhibits the IGF/mTOR axis and reduces protein synthesis, which we might predict exacerbates the effects of intrinsic ageing. Indeed, there are several lines of evidence that CR reduces cognitive flexibility^327^, muscle mass^357^, and bone density^358^, mirroring the effects of intrinsic ageing. However, as we discussed in Part 4.6, CR should also have potent rejuvenative effects on extrinsic ageing, reducing nutrient overload and preventing the cascade toward metabolic syndrome^359,360^. As shown by Mair, et al.^231^, DR reduced short-term mortality in flies without affecting long-term risk, consistent with this model. However, when the same lab swapped mice between CR and AL it produced mortality profiles which were still affected by their previous dietary regimen^361^, suggesting that in mammals CR also impacts long-term risk, likely independent of extrinsic ageing.

Partly this may reflect that CR reduces celerisis, inhibiting hyperfunction, cancer, and fibrosis, as the reduced growth rate should reduce the advantage of faster cells (assuming they are more sensitive to growth signals), but it may also reduce intrinsic ageing. According to the three process model, to affect intrinsic ageing, an intervention should reduce the rate of selective destruction. We suggest that by reducing the IGF stimulus and the resultant basal metabolic rate of cells, CR causes a greater reduction in the metabolic rate of more sensitive cells. Thus, the ‘faster’ cells slow more than the slower cells and behave more similarly, allowing them to escape the selective destruction that would otherwise epigenetically reduce their metabolic rate. Thus, while CR may exacerbate the effects of intrinsic ageing by slowing the body’s metabolism, it should also slow the permanent reductions in metabolic rate, so that the rate of intrinsic ageing is reduced (see Figure 12).

**Figure 12.**
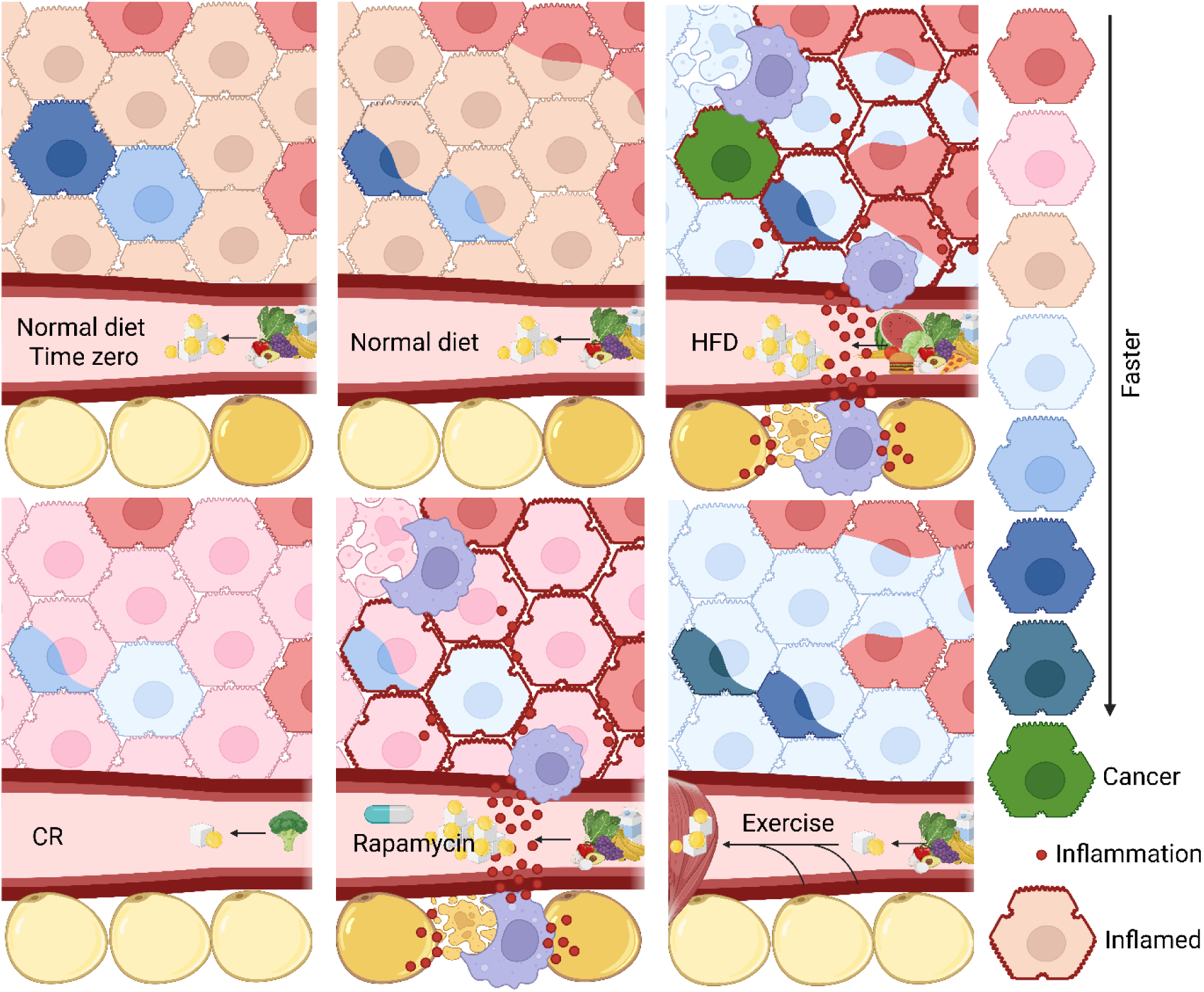
Effects of behaviour and therapies on intrinsic and extrinsic ageing. Top left image shows time zero, allowing comparison of all other images to this time point to see change over time (the mutants present at time zero are there by chance to demonstrate what will happen to them, not an indication of the effects of control diet at time zero). Cells with two colours are undergoing selective destruction, epigenetically reducing their metabolic rate. The colour representing the faster metabolism is their metabolic rate before being slowed by neighbouring cells. When cells change colour (from time zero) to a new full colour it reflects environmentally induced (and temporary) change in metabolic rate. Dead cells have been killed by macrophages (purple) to recycle the lipid. Abbreviations: CR, calorie restriction; HFD, high fat diet.

This distinction may help explain one of the most contentious points^362^ in ageing and health research, the obesity paradox – that overweight and obesity are associated with reduced mortality compared to both normal and underweight individuals. The surprising result was identified for cardiovascular disease^363^, a host of other diseases^364^, and all-cause mortality^365^. These results contrast with other studies demonstrating that overweight and obesity are responsible for significant increases in morbidity and mortality^363,364^.

While obesity may reflect reduced time spent in elevated metabolism (exercise), it is a state of elevated basal metabolic rate (BMR)^366^, including elevated levels of GH, IGF, and mTOR signalling. The effects should therefore be the reverse of CR, exacerbating extrinsic ageing, and stimulating the more sensitive faster cells to an increasingly elevated metabolism that will increase the rate of selective destruction and intrinsic ageing; however, as such, it should also transiently counteract the direct effects of intrinsic ageing and metabolic slowdown by increasing the body’s metabolic rate.

Similarly, GLP-1 inhibitors, reducing appetite, should mimic the effects of CR, reducing anabolism and compounding the effects of intrinsic ageing on muscle loss, as is becoming apparent^367^, but also reducing the rate of intrinsic and extrinsic ageing, with potentially net beneficial effects.

The most common ageing profile for body mass in humans is usually weight gain until around 60-70, followed by a plateau and then weight loss^368^. These trends will result from metabolic shifts that reflect the relative impacts of intrinsic and extrinsic ageing on homeostasis, likely influencing the net impact of weight on mortality. We might predict that the period of weight gain should correspond to the greatest impact of extrinsic ageing, with more rapid weight gain corresponding to accelerated extrinsic ageing and a corresponding impact on mortality. As weight gain tapers and weight loss begins, intrinsic ageing is likely accelerating, with the consequent loss of mainly fat-free mass^369^. Mortality should increasingly reflect frailty rather than nutrient overload and metabolic syndrome. If true, then at this point additional weight may begin to have a net-positive effect, consistent with the obesity paradox. That is, while extrinsic ageing is the dominant factor affecting mortality, higher weight will be detrimental, then as intrinsic ageing overtakes, the elevated stimulus of increased weight may start to produce increasing benefits, eventually outweighing both the detriments of extrinsic ageing and the increased rate of intrinsic ageing, producing short-term metabolic stimulation that counteracts the metabolic slowdown.

Notably, the three process model also highlights differences between CR and intermittent fasting or time-restricted feeding which are designed to have health benefits without radical calorie reduction. By extending the periods without eating, the latter should have beneficial effects mainly on extrinsic ageing, while (assuming zero calorie reduction) this model suggests that the net rate of intrinsic ageing would be unchanged, with the short term effects of diet on intrinsic age and frailty oscillating between fasting and feeding periods. However, not inconsistently, there is some evidence that refeeding increases risk of cancer^370^, as might be predicted if consuming all the calories within a short period advantages faster cells and encourages celerisis.

### 6.2 Rapamycin

Rapamycin is probably the most consistent longevity drug yet documented, enhancing maximum lifespan in mice^371,372^ and flies^373^, including long-lasting effects after the regimen ceases^374^. It has multiple effects on cell and molecular phenotype, including reducing cell division rate, but these are mainly (at least at low dose) thought to result from the inhibition of mTORC1^375,376^, which is a central driver of cell growth and metabolism^377^.

Thus, in some respects, rapamycin is a CR mimetic. By inhibiting the growth pathways, it would be predicted to reduce the rates of both celerisis and intrinsic ageing, while compounding the effects of intrinsic ageing on frailty, just as CR does. However, CR reduces mTOR signalling by reducing the glucose, lipid, and amino acid activators which trigger both insulin/IGF and mTOR directly^378,379^, thus reducing extrinsic ageing, while rapamycin inhibits mTOR without changing the nutrient availability. Thus, we predicted that rapamycin would initially exacerbate extrinsic ageing and metabolic syndrome. Consistently, rapamycin has been shown to induce IR, glucose intolerance, and diabetes, while still being associated with robust increases in lifespan^380,381^. However, by retarding intrinsic ageing and metabolic slowdown the long-term effects on extrinsic ageing will also be beneficial (see Figure 12).

Thus, rapamycin may have the transient, so-called ‘paradoxical’^382^ negative effects of weight gain and raised fasting glucose levels at the start of treatment^383^ because it is accelerating extrinsic ageing. Consistently, Schindler, et al.^384^ found that key factors affecting metabolic impairment from rapamycin included the dose and the age of treatment. Other explanations, such as the mTORC2-specific rapamycin effects^381^ may be significant but do not explain the general paradox of growth signalling and ageing^385,386^. Insulin-like growth factor (IGF) and growth hormone (GH) show robust decreases during lifespan^387,388^, while GH level correlates negatively with cardiovascular risk^389^, and adult supplementation has had short-term beneficial effects, presumably by temporarily reversing the effects of intrinsic ageing with increased anabolism^390^; however, longer-term use in healthy older adults is associated with increased adverse events^391^, and knockouts are associated with increased longevity^392,393^. All these effects fit within the three process model if the short-term benefits of pro-growth factors reflect temporary reversal of metabolic slowdown while worsening its long-term trajectory.

### 6.3 Exercise

Multiple studies and meta-analyses indicate the beneficial effects of exercise on mortality, but the effects are generally limited to healthspan and mean lifespan with no maximum lifespan extension^233–235^, as predicted by the three process model. Exercise, like GH supplementation, is stimulating anabolism. Crucially, skeletal muscle contributes to more than 30% of resting metabolic rate and 80% of whole body glucose uptake^109^, suggesting it is a potent inhibitor of extrinsic ageing and, like obesity, could temporarily reverse the effects of intrinsic ageing with pro-growth stimuli. However, the effects on the rate of intrinsic ageing will be more complex (see Figure 12 for most basic interpretation), and may differ between mitotic and post-mitotic tissues depending on whether selection and counter-selection are mitochondrial or cellular (see Part 7). To answer this question requires further work, but consistent with the lack of maximum lifespan gains, may not always be net positive.

If exercising animals are largely healthier than sedentary ones but experiencing compression of late-life morbidity without maximum lifespan gains, then a negative effect on intrinsic ageing is not implausible. Notably, a recent study in mice which ended the exercise regimen early found healthspan and maximum lifespan gains^394^. Speculatively, acute events triggered by the stresses of exercise in the elderly could help explain why mice lived longer in this study and not those continuing exercise into old age. Consistently, a human study suggested that 2-3% of sudden cardiac arrests in individuals ≥65 years old occurred during or immediately after exercise^395^. However, as the mice assessed by Feng, et al.^394^ showed no median lifespan extension, it would suggest that there are still additional trade-offs at play. Fascinatingly, in a study by Holmstrom, et al.^395^, individuals with fewer cardiovascular risk factors were more likely to die of exercise-induced events. This is compatible with the observation that cardiovascular events of youth are also not associated with the standard cardiovascular risk factors. For example, hypertrophic cardiomyopathy is the most common cause of cardiac mortality in youth, usually precipitated by exercise, and is mechanistically associated with fibroblast activation and fibrosis^396^. Combined, these results suggest that exercise could have detrimental effects by stimulating celerisis and hyperfunction in some contexts. However, its robust association with reduced cancer rates^397^ may be explained by its potent reduction of nutrient overload, extrinsic ageing, and inflammageing.

### 6.4 Metformin

The mechanisms of action for metformin are complex and not well understood^398^; however, it is primarily regarded as an AMPK activator^399^. The central function of AMPK is maintaining ATP levels, thus whether it directly impacts metabolic rate and intrinsic ageing will depend on additional factors and feedbacks regulating the ATP equilibrium. The model we discuss in Part 9 describes the potential effects of metformin (Part 9.2.4.2) on energy homeostasis in more detail.

Studies suggest metformin increases fatty acid oxidation, particularly in brown adipose tissue through futile cycling in thermogenesis/uncoupling^400^, resulting in reduced L_B_ and the lipid droplet content in adipose^401^. Others suggest an increase in futile anaerobic glycolysis to produce lactate, which is then targeted back to the liver for gluconeogenesis^402^, with a net wastage of four ATP. This admittedly non-exhaustive list of metformin’s effects suggest a major role in inducing pathways which burn nutrients without producing ATP, thus reducing the effects of nutrient overload and extrinsic ageing. Consistently, while metformin has beneficial effects on healthspan and mean/median lifespan^399,403^ it does not appear to extend maximum lifespan in mice^404^, flies^405^, or worms^403,406^, suggesting it might not reduce BMR or intrinsic ageing.

### 6.5 Yamanaka (OSKM) factors, parabiosis, and epigenetic rejuvenation

Finally, the Yamanaka factors and parabiosis have both shown robust life extending potential^407,408^, indicating epigenetic markings as an anti-ageing target.

The mechanism underlying parabiosis could theoretically be epigenetic and rely on a kind of selective destruction in blood cells. Crucially, we are not suggesting that selective destruction always results in metabolic slowdown. **Selective destruction is a mutant recognition and correction system**. It gives an advantage to slower cells to balance the natural course of selection with counterselection, but in cases where the slower cell is still more likely to be the mutant, selective destruction may instead induce metabolic acceleration. Thus, parabiosis could reverse the effects of intrinsic ageing by exposure of old, slowed cells to large quantities of young cells. However, studies attempting to address which factors in young blood are responsible for rejuvenation have shown that replacing young blood infusion with a saline infusion produces similar benefits^409^. Potentially, the health benefits of parabiosis reflect the loss of old blood rather than the gain of young blood. If so, a good candidate is the loss of L_B_ in the form of VLDL and LDL, which the body then restores by reducing its adipose stores and activating cholesterol synthesis from L_I_ and glucose. Thus, parabiosis and blood donation would be predicted as beneficial therapies against extrinsic ageing.

According to the three process model, the most effective existing therapy targeting intrinsic ageing would be a cocktail of four factors, Oct4, Sox2, Klf4, and Myc (OSKM), also known as the Yamanaka factors which are used in the generation of induced pluripotent stem cells (iPSCs). While therapies such as CR and rapamycin can reduce the rate of intrinsic ageing by slowing metabolic rate and reducing the rate of selective destruction, only the Yamanaka factors have potential to remove the epigenetic marks induced by selective destruction and thus reverse intrinsic ageing.

The current problem with Yamanaka factor rejuvenation is that it removes all epigenetic marks, thus de-differentiating the cells at the same time as rejuvenating them. Studies are currently working with pulsatile expression designed to trigger rejuvenation without de-differentiation, but if the three process model is correct, we would predict targets should exist that could reverse the epigenetic marks of ageing without affecting cell fate. We would need to identify the pathways of selective destruction and find how they activate the chromatin remodelling factors which induce intrinsic ageing. However, while such therapies might be substantially rejuvenative, they may also trigger celerisis and lead to hyperfunctional disorders. We may have to combine epigenetic rejuvenation with anti-cancer therapies.

## Part 6: Summary

6.1 | Reduced calorie intake should slow extrinsic ageing by reducing nutrient excess, conversely exacerbated by high calorie diets.

CR may also slow intrinsic ageing by reducing the metabolic rate of faster cells more than slower cells, as the former are more sensitive to changes in growth stimuli. Therefore, faster cells are less likely to be noticed as aberrant and epigenetically slowed by selective destruction.

The obesity paradox may result as being overweight exacerbates the detrimental effects of extrinsic ageing as well as accelerating intrinsic ageing, increasing mortality. However, it may also provide short-term pro-growth signals that counteract the metabolic slowdown produced by intrinsic ageing, temporarily boosting cellular function. As mortality from intrinsic ageing increases, additional weight may become increasingly beneficial.

6.2 | Rapamycin should have similar effects to CR, but without reducing nutrient excess. Thus rapamycin may initially exacerbate extrinsic ageing and induce metabolic syndrome.

6.3 | Exercise will reduce the nutrient excess and reduce extrinsic ageing.

6.4 | Metformin does not extend maximum lifespan, limiting its impact to extrinsic ageing.

6.5 | Parabiosis may work primarily by reducing blood lipid carriers, reducing extrinsic ageing.

Yamanaka factors may be the only therapy to offer true rejuvenative potential by removing the epigenetic marks which induce intrinsic ageing. However, epigenetic rejuvenation may increase risk of celerisis.

## Part 7: Additional selection and counterselection pressures

### 7.1 Post-mitotic tissues and organisms

It was initially our intent to discuss selection in post-mitotic tissues in a separate paper; however, ignoring ageing in C elegans, Drosophila, yeast, and even human brains, would have left important mechanisms unexplored in a supposedly ‘unifying’ model. Ageing of post-mitotic tissues and unicellular organisms is unlikely to reflect intercellular competition and selection because these forces are largely or completely absent in such systems.

Post-mitotic cells can gain mutations shifting their metabolic rate, which may present some threat and thus be epigenetically controlled. However, as most DNA damage results from replication, the rate is much reduced, as is the threat from DNA damage, which mainly arises from proliferation if the faster cells are to gain advantage and cause celerisis. The natural course of selection toward faster cells is thus significantly reduced, as is the need for selective destruction to counterbalance it.

However, post-mitotic cells are made increasingly vulnerable to another type of selection, this one intracellular, dependent on the replication of a DNA containing organelle.

Mitochondria exist in a state of ‘relaxed replication’ independent of the cell cycle, which continues in permanently arrested cells. Thus, like dividing cells and reproducing organisms, mitochondria are subject to the forces of selection. In short, we suggest that mitochondrial DNA (mtDNA) damage results in the ageing of post-mitotic and unicellular eukaryotic organisms due to similar forces of selection and counterselection at the level of mitochondria.

#### 7.1.1 Damage to mtDNA may drive ageing of post-mitotic and unicellular systems

The contribution of mtDNA mutations to ageing in post-mitotic human tissues is controversial. Yet multiple studies suggest that brain ageing, cognitive impairment, and Alzheimer’s disease are associated with mtDNA damage^410,411^, the hippocampus of early-stage Alzheimer’s being associated with a significant increase in mtDNA damage^412^. Equally, there is no consensus that mtDNA plays a critical role in the ageing of post-mitotic organisms^413–415^, although some studies suggest it is significant in both Drosophila^416^ and C. elegans^417^, including the loss of dopaminergic neurons in flies^416^. The role of mitochondria and mtDNA damage in yeast replicative ageing is more widely accepted^418–420^. Additionally, mice with mutations that produce an error-prone mitochondrial polymerase also demonstrate shortened lifespan^185^, though again the relationship is not always consistent^421,422^.

The picture is clearly complex and difficult to reconcile with models of straight mtDNA damage accumulation. Antioxidant mutants and ionising radiation can lengthen worm and fly lifespan concomitant with increased mtDNA damage and mutation levels^413–415^. Equally, long-lived daf-2 mutants appear to accumulate more mtDNA mutations with age^414^. Any attempt to associate lifespan with ROS levels or the mitochondrial free radical theory results in a ‘panoply of paradoxes’^423^, and even the studies suggesting mtDNA is relevant to ageing suggest it results from mtDNA replication rather than oxidation^412,424–426^. However, it is possible that the complex relationship between mtDNA damage and ageing is explained by mitochondrial selection.

#### 7.1.2 Selection in mitochondria

Interestingly, the idea of mitochondrial selection is more established than for cells. It was frequently observed that small and large mtDNA deletions (ΔmtDNA) often accumulated within cells, with each cell containing many copies of the same deletion, while different cells could contain different deletions spreading within them.

Multiple specific mtDNA deletions have been observed in humans and model organisms which accumulate with age in post-mitotic tissues. One deletion which spans 4,977 base pairs, preventing complex I, IV, and V function (leaving only complex III intact), is called the ‘common deletion’ because it is frequent both within and between cells in aged tissues^427^. Other common mutations have been identified in humans and rodents^428,429^.

However, it is plausible that the commonality of these specific mutations is the product of the relative rates in which they are produced rather than selection, for example if mtDNA is particularly prone to large deletions. As argued by an influential modelling study, drift alone could explain the number of cells showing accumulation of mtDNA deletions^430^. However, while drift is still considered a potential factor^431,432^, such models do not incorporate selection or suggest what the mutational landscape should look like in its presence. Even if the abundance of deletion mutations can fit with models of drift, mutations which spread by selection should be more numerous.

To explain the absence of the latter suggests that either mutations cannot induce selective advantage, which evidence from oocyte production and zygote formation^433,434^, and mtDNA depletion experiments^435^ all suggests is unlikely, or mitochondria are somehow robustly protected from mutations that might give them advantage while unable to prevent the spread of highly deleterious mutations that spread only by drift. Such defence mechanisms would reflect significant complexity.

As argued persuasively by Kowald and Kirkwood^436^, the near homoplasmy of specific deletions produced by the drift models of Elson, et al.^430^ could only be explained by very low mutation rates, otherwise cells should experience significant heteroplasmy of multiple different mutations. Thus, long human lifespans were required to allow single mutations to spread by drift. In animals with shorter lifespans, such as rodents, higher mutation rates would be required to produce the same level of mtDNA deletions across cells, but this would also make it unlikely that aged cells should be found with single, homoplasmic mutations, inconsistent with data in rats^437^, which also show single deletions spreading within cells^438^. While the frequency of ΔmtDNAs between cells is likely the product of the high rates of specific deletion events, the frequency within cells most likely reflects selection.

Proponents of mitochondrial selection have suggested various mechanisms. Since the initial ideas that deletions would produce smaller plasmids with a replication advantage^435,439–442^, this idea was challenged by de Grey^443^, who suggested that the rate of replication was too quick to be the rate limiting factor^444^, further emphasised in computational models^445^. Thus, de Grey^443^ suggested that the common deletions might undergo selection not because they were smaller, but because the remaining plasmids lacked the capacity to create complexes for functional OXPHOS – it was ‘survival of the slowest’.

As ROS were deemed a natural and unavoidable toxic byproduct of respiration^446^, it was believed that reduced OXPHOS would protect mitochondria from damage, which would thus reduce their rate of degradation. Consistently, studies suggested that the oxidation of the mitochondrial membrane was a significant factor determining which mitochondria were degraded by mitophagy^447–450^.

This notion was again challenged by Kowald and Kirkwood^436^ who observed that mitochondria are not distinct units, but form a fused network where plasmids have only a ‘leaky link’ to the mitochondrial membrane they are attached to. They suggested that fused mitochondria could allow passage of plasmids, proteins, and RNAs between organelles, breaking the link between genotype and membrane phenotype that could allow for selection of ‘mitochondria’ rather than mtDNA.

However, mtDNA do not simply diffuse around the mitochondrial matrix, but are localised to the nucleoid where they are synthesised, with each nucleoid attached to the inner mitochondrial membrane^451^. Each nucleoid contains about 6-10 plasmids which rarely swap between nucleoids^452^, while each nucleoid has a very low diffusion constant or does not move at all^451,453^. Furthermore, nucleoids appear to be accurately positioned within the mitochondria with set distances between them^451,454–457^, suggesting they are highly regulated.

The connection between nucleoids and the protein complexes of the ETC is less clear. Mitochondrial mRNAs often associate together into mitochondrial RNA granules (MRGs). One study suggested that although MRGs were attached to the IMM, they showed a random distribution, in stark contrast to nucleoids. They also rapidly exchanged components, sometimes fusing with each other, consistent with fluid condensates positioned by membrane dynamics^458^. Conversely, the mitochondrial (protein) complexes often associated together as supercomplexes^459,460^, which may^461^ or may not^462^ associate with nucleoids.

The separation of mRNA and proteins from the nucleoids will produce a “leaky link” between the genotype and phenotype as proposed by Kowald and Kirkwood^436^. However, the degree of leakiness will depend on how well the mRNAs and supercomplexes can separate from the nucleoids ultimately responsible for their composition. The IMM forms cristae which envelop away from the outer mitochondrial membrane (OMM), which is where the majority of supercomplexes form^462^. The high protein content of supercomplexes makes their diffusion rate very low^463^, and there was strong co-localisation between complexes and the mitoribosomes^462^. Cristae junctions (CJs) also separate the cristae from the segment of IMM close the OMM (the inner boundary membrane, IBM) forming natural diffusion barriers^464^ which effectively trap the supercomplexes within the same cristae as their nucleoids.

Thus, we suggest that mtDNA will influence the phenotype of the surrounding membrane more than it will that of mitochondrial membranes further away, and selection could occur at the level of mitochondria even in a fused network. In effect, a leaky link is still a link.

Selection at the mitochondrial level through differential degradation would require that fission and mitophagy were selective processes, and consistently dysfunctional mitochondria are targeted for fission from the mitochondrial network^465^ and then degradation by mitophagy^448,450,466,467^. However, as suggested by Kowald and Kirkwood^436^, this still presented a problem for survival of the slowest, because in these studies it was the dysfunctional mitochondria that were being targeted for degradation.

Here, we suggest that this apparent paradox between the targeting of dysfunctional mitochondria for degradation and the escape of dysfunctional mitochondria as suggested by de Grey^443^, is that there is more than one type of dysfunction, and mitochondria may be forced to rely on crude biomarkers to calculate which dysfunction has occurred, just as we described for cells in Part 3.

#### 7.1.3 Selective destruction in mitochondria

Unlike Kreb’s cycle, which is not so much a cycle as a network of major biosynthetic and energy pathways, OXPHOS is a relatively simple linear pathway. As shown in Figure 13A, Complex I accepts the electrons from NADH+H^+^ (the major energy producing product of Kreb’s cycle), pumping the protons across the inner membrane into the intermembrane space. Nuclear encoded complex II accepts the electrons from FADH_2_ (the other major product of Kreb’s cycle), but does not pump protons. Both complexes then pass the electrons to membrane-bound ubiquinone, which passes them to complex III, pumping more protons across the membrane before finally passing the electrons via cytochrome C to complex IV, which pumps more protons while returning the electrons to the matrix to react with oxygen and form water. Complex V then forms a pore in the membrane allowing the protons to flow back into the matrix at the same time as phosphorylating ADP to form ATP. Thus, there are **two major ways that the mitochondria can malfunction**, either through **insufficient flow of electrons and protons into the membrane complexes and intermembrane space** (**depolarisation**), or **too little flow of electrons and protons back into the matrix** (**hyperpolarisation**).

**Figure 13.**
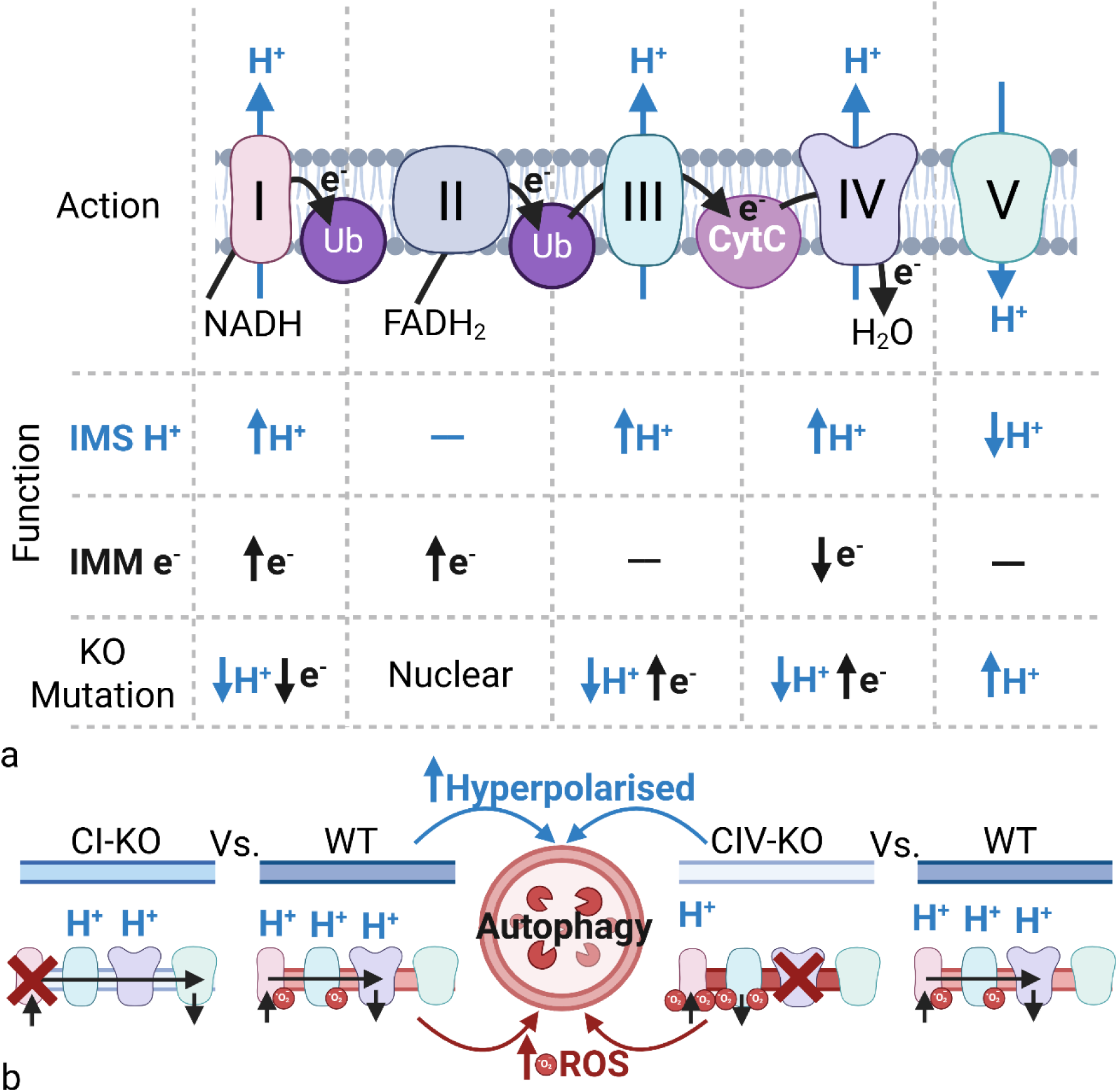
Biomarkers of electron transport chain function and effects of mutation. (A) Table of effects of functional complexes on intermembrane space (IMS) proton levels (polarisation) and electrons present in the ETC on the inner mitochondrial membrane (IMM). Bottom row demonstrates the effects on IMS protons and ETC electrons when the complexes are knocked out. (B) Mutations in complex I (CI-KO) and complex IV (CIV-KO) have opposite effects on mitochondrial oxidation compared to wildtype, which may result in the autophagy of wildtype mitochondria when the network attempts to decipher which mitochondrion is damaged in the presence of complex I mutations. Image created with BioRender.com. Abbreviations: CI-KO, complex I knockout; CIV-KO, complex IV knockout; CytC, cytochrome C; e^-^, electron; FADH_2_, flavin adenine dinucleotide; H^+^, hydrogen ion; KO, knockout; IMM, inner mitochondrial membrane; IMS, intermembrane space; NADH, nicotinamide adenine dinucleotide; ROS, reactive oxygen species; WT, wildtype; Ub, ubiquinone.

Just as a cell must distinguish whether having faster metabolism than its neighbour reflects that it has gained a hyperfunctional mutation or its neighbour has gained a hypofunctional one, mitochondria must calculate whether an increasingly polarised neighbour has a mutation reducing its capacity to return protons and electrons to the matrix, or whether its neighbour is normal, and the difference reflects its own loss of capacity to pump protons into the intermembrane space. Simply removing the ‘most damaged’ is not so easy.

In Figure 13, we indicate how the ETC can be viewed in terms of polarising and depolarising forces. Knockout mutations in different complexes will have different effects on both the negative charge of the membrane and the positive charge of the intermembrane space, both contributing to the polarisation of the mitochondria. Mutations in complex V prevent the flow of protons back to the matrix, causing build-up and hyperpolarisation, whereas mutations in complexes I, III, and IV all lead to depolarisation.

Perhaps predictably, mitochondrial selection appears to favour gradual depolarisation over hyperpolarisation, preventing the unrestrained build-up of charge. While apoptosis is usually associated with the act of depolarisation, it may be initiated by the hyperpolarised state leading to activation of ROS production and membrane pores allowing depolarisation^468,469^.

Additionally, the diffusion of H^+^ away from dysfunctional complexes may make the link even more leaky, necessitating strong selection against potentially hyperpolarised areas of membrane. Thus, during uncertainty as to which part of the network is dysfunctional, mitochondria may favour the selective destruction of the more polarised mitochondria.

Consistently, mitochondrial membrane potential is often observed to decrease with age^470–472^, while a study in mutator mice with accelerated ageing demonstrated reduced content of complexes I, III, and IV, but not V^473^.

However, while this could explain survival of the slowest, it has little to do with ROS as was initially proposed by de Grey^443^. It is plausible that mitochondria have two central biomarkers of dysfunction, the polarity primarily demonstrating the capacity of the ETC to pump protons, while capacity for electron transport is detected by a specific biomarker of electrons forced out of the ETC before reaching complex IV. Superoxide is well known to be generated almost exclusively by complexes I and III^181–184,474,475^, reflecting that it is not an unavoidable, toxic byproduct of mitochondrial metabolism, but intended to oxidise the membrane around dysfunctional complexes as a biomarker of ETC failure. As complex III is required to pass the electrons to cytochrome C which passes the electrons to complex IV, the function of both complexes is required to return the electrons from the ETC to the matrix. Thus, when complex IV is dysfunctional, complex III passes the electrons to oxygen, and when complex III is dysfunctional, complex I passes the electrons to oxygen, both forming superoxide. Superoxide then oxidises the membrane and triggers the degradation of the mitochondrion.

Were this the only problem, mitochondrial selection could plausibly restore function and maintain wildtype dominance with relative ease. Higher ROS would reflect damage, mitochondria could select the more depolarised to select against complex V mutations, and lower ROS levels to select against complex III and IV mutations, ensuring no such mutations were allowed to spread. However, the mutation of complex I results in the reverse. Instead of causing electron build-up, it reduces the capacity of the ETC to accept electrons from NADH+H+, and as a result the ETC will produce fewer ROS than functional mitochondria. Thus, when two mitochondria with differences in membrane oxidation and polarity are competing to induce the fission and degradation of the other, the system has a critical vulnerability that lower ROS oxidation could reflect wildtype status compared to complex III or IV mutation, or complex I mutation compared to wildtype (see Figure 13B). If true, cells have no clear mechanism to select for ‘functional’ mitochondria over ‘damaged’ ones, instead, at least under some conditions, selecting for one type of damage over another.

As elevated ROS levels could result from the knockout of two complexes while decreased ROS levels should be produced by only one, mitochondrial selection might again favour selectively destroying the mitochondria with the highest ROS levels, slowly advantaging complex I mutations over wildtype, and wildtype over complex III and IV mutations.

Notably, we have focussed here on complete knockout mutations. Not all complex I mutations will decrease ROS production: if functionality is altered rather than abrogated, the opposite may be true. Equally, mutations in complex III could specifically reduce its passage of electrons to oxygen, reducing ROS. We have referred to knockout mutations for simplicity, but in reality, any mutation for depolarisation and low ROS should undergo positive selection. To distinguish it from generalized mitochondrial dysfunction or random mtDNA damage accumulation, the process of **selection for low ROS, depolarised mitochondria has been termed mitopenia**.

#### 7.1.4 Mitopenia explains mtDNA mutations and is linked with ageing

If the basic thesis of mitopenia is true, then mitochondria are not selecting for hypofunctional organelles over hyperfunctional ones as with cells. Even the hyperpolarising mutations of complex V do not increase the activity of mitochondria. The rate of mitochondrial biogenesis is regulated by the nucleus, and no changes in the complex activity (or tRNA sequence) will induce a selective ‘hyperfunctional’ advantage for mitochondrial growth as a nuclear oncogenic RAS mutation does for cells. As each mitochondrion has many plasmids, the pro-growth advantage occurs at the level of the mtDNAs. While mutant mtDNAs show little advantage from additional replication origins, and large deletions are unlikely to gain replicative advantage directly from their reduced size^444,445^, a clever study by Kowald and Kirkwood^476^ demonstrated that the coupling of transcriptional output and mtDNA replication could result in additional replication of deletions because they do not produce sufficient product to inhibit transcription which is coupled to DNA replication. Thus, the anti-growth negative feedback is not induced.

A major force of selection for large mitochondrial deletions might therefore still be their growth advantage, reflecting the natural course of selection. However, as we saw with cells, when the natural course of selection is not inhibited by counterselection, it may not reflect an evolutionary incapacity to prevent unidirectional selection as proposed by Nelson and Masel^142^, but the difficulty of balancing selection and counterselection under changing conditions. We suggest that the frequency of mtDNA mutations within cells will reflect three main factors:

- The frequency of the mutation events required to produce the specific genotype (formation advantage).
- The ultimate size of the plasmid and its transcriptional output (growth advantage).
- The level of membrane oxidation and polarisation it produces relative to wildtype and other present mutant mtDNA plasmids (survival advantage).

Notably, multiple mtDNA duplications also appear to spread during ageing^477–479^, which are difficult to explain by either growth advantage alone or a survival advantage based purely on reduced functional output as in survival of the slowest. While the cause of their spread is beyond the scope of this article, we would predict that mtDNA duplications would offer advantage if they produced small shifts toward depolarisation and low ROS by altering the ratio of complexes in the ETC.

The common deletion retains complex III activity, which may contribute to its success. As complex II is nuclear encoded, the ETC can pass electrons from complex II to III, which returns them to the matrix generating superoxide, pumping protons which cannot be returned by complex V. As the accumulating NADH+H^+^ should slow or stall Kreb’s cycle, reducing FADH_2_ generation and complex II activity, the low proton pumping without complex V to return them, plus the high electron leak per electron accepted, may create a depolarised, low oxidised membrane that is close enough to wildtype to be unclear which is the mutant. Under such conditions, the mitochondrial network might choose to degrade the wildtype. As suggested by de Grey^443^, the low ROS levels of the ΔmtDNA may thus contribute to their positive selection and cause survival of the slowest. However, this would not reflect that ROS are toxic byproducts, but a flaw in the selection and counterselection mechanisms, inducing mitopenia. Superoxide is a crucial biomarker regulating mitochondrial selection, without which mitochondria should become vulnerable particularly to the spread of complex III and IV mutations – perhaps explaining in part why antioxidant supplementation can be detrimental to health^10^.

The question of whether large deletions in themselves could be responsible for the age-related degeneration of post-mitotic tissues is still under debate. Studies of ageing rat skeletal muscle showed that ΔmtDNA was at its highest levels of heteroplasmy in regions associated with the atrophy, splitting, and breaking of fibres^480,481^. Early studies suggested ΔmtDNA also accumulated in elderly human skeletal muscle and heart^429,438,480,482–485^, contributing to frailty and arrythmia. In the brain, ΔmtDNA accumulated at higher levels in the substantia nigra, associated with neurodegeneration and parkinsonism^486–488^, and some studies suggest association with Alzheimer’s disease^489^.

Twinkle helicase mutant mice, known as ‘deletor’ mice, accumulate high levels of ΔmtDNA with age, show dysfunctional cells and other abnormalities in the heart^490^, as seen in inherited mitochondrial diseases in humans^491^. However, these same dysfunctions were not observed in control mice even at advanced age, and Twinkle helicase mutant mice did not demonstrate accelerated ageing. Equally, while ΔmtDNA mutations do occur in C elegans, they are likely too rare to contribute to ageing, which may reflect that the worms are too short-lived^492^.

In a fascinating study by Ahlqvist, et al.^493^, ‘mutator’ mice with inactivated exonuclease function of POLγ causing the accumulation of point mutations and complex rearrangements, were compared to the deletor mice with ΔmtDNA accumulation. The results clearly demonstrated that while the deletor mice had respiratory complex deficiency in their post-mitotic tissues, consistent with previous data^494^, this had no effects on their neural stem cells (NSCs) or haematopoietic stem cells (HSCs).

Conversely, the mutator mice showed defects in their NSCs and HSCs inducing more differentiation, as well as a premature ageing phenotype including “progressive hair graying, alopecia, osteoporosis, general wasting, and reduced fertility”. Yet, while the deletor mice had normal lifespan, they showed evidence of mitochondrial dysfunction in neurons not present in mutator mice^493^.

These results suggest that mitochondrial deletions are detrimental to the function of post-mitotic tissue, contributing to disease though not significantly to mortality. However, mitotic cells may utilise cell division as an additional form of selection against ΔmtDNA, potentially by ensuring that all deletions are retained within one daughter cell, as in yeast. This may be largely responsible for the replicative ageing of yeast^495^, though as the process creates a ‘doomed cell’ unloading other damaged or unnecessary molecules from lipids and proteins to other organelles and extra-chromosomal ribosomal circles (ERCs) into it may be an efficient mechanism of removal that further contributes to decline^496^.

This would explain the relative absence of cells with strong heteroplasmy for ΔmtDNA in mitotic tissues, while quiescence may allow such mutations to accumulate. The spread of ΔmtDNA would reflect part of the mitopenia phenotype, but limited to post-mitotic cells and potentially longer lived organisms requiring extended time to spread. Even in long-lived organisms, the effects specifically of deletions may not be a strong contributor to disease in most cases.

In both proliferating and post-mitotic cells the selection and selective destruction of point mutations could contribute more strongly to decline. Even in the shortest lived organisms, such as C elegans, ageing involves profound mitochondrial changes. The fragmentation of the network is accelerated under conditions that shorten lifespan and delayed by conditions which extend lifespan^497^.

Here we suggest that primary cause of these changes is selection against hyperpolarising and oxidising mutations which unavoidably selects for complex I mutations. Fascinatingly, depletion of complex I occurs across a wide variety of tissues and species^498^. A proteomic study assessing mitochondrial genes with decreased representation at older age identified the complex I subunit, NDUFB2, as the most significant, with three of the top five also being complex I subunits, while succinate dehydrogenase complex assembly factor 2 (SDHAF2) of complex II was upregulated^499^, potentially as a compensatory mechanism. Complex I dysfunction is an early change in ageing brains affecting mitochondrial structure and capacity^500^, while the mutator mice with accelerated ageing showed the most pronounced reductions in complex I protein^493^.

Notably, the second most affected complex is frequently complex IV^493,499^, resulting in “COX-deficient cells”, a widely used biomarker of mitochondrial dysfunction, which is not directly predicted by this model, but could reflect a weaker counterselection force against complex I mutations, strengthened under certain conditions such as antioxidant supplementation or FOXO activation.

Mitochondrial selection and mitopenia may therefore be a contributor to ageing and age-related disease across eukaryotic life in both mitotic and post-mitotic tissues. However, we suggest that the mechanisms by which mitopenia causes disease should be through its interaction with intrinsic or extrinsic ageing.

#### 7.1.5 Mitopenia induces intrinsic and extrinsic ageing

An initial prediction might be that as mitopenia makes mitochondria less capable of producing ATP, the levels of ATP would drop. This drop might then precipitate a drop in ATP utilisation to re-establish homeostasis, potentially via an epigenetic mechanism inducing semi-permanent change. In this way, mitopenia would directly induce intrinsic ageing; the catabolic slowdown leading to anabolic slowdown. Consistently, nuclear encoded complex II, whose activity levels are a good biomarker of both OXPHOS and Kreb’s cycle, is a key player in retrograde (nuclear-mitochondrial) signalling and epigenetic changes in the nucleus^102^.

Notably, curtailing anabolic requirements according to energy availability is a poorer evolutionary strategy than ensuring energy availability. As the capacity of the average mitochondrion to produce ATP declines, anabolism could be maintained by additional changes, such as increasing mitochondrial mass. Interestingly, while there is good evidence that mitochondrial mass can increase to compensate for dysfunction^174,501^, ageing appears to associate with the opposite change, with muscles and neurons showing decreasing mitochondrial mass with age^502,503^. If so, mitopenia may directly induce intrinsic ageing.

However, additional factors are likely of significance to explain why homeostasis is shifted rather than maintained by a compensatory change in mitochondrial mass.

Another possible mechanism reflects the interaction between nutritional changes and mitochondrial selection. In situations of nutrient excess, upregulated insulin/IGF signalling increases nutrient uptake, and more ACoA passes into Kreb’s cycle (see Figure 14A).

**Figure 14.**
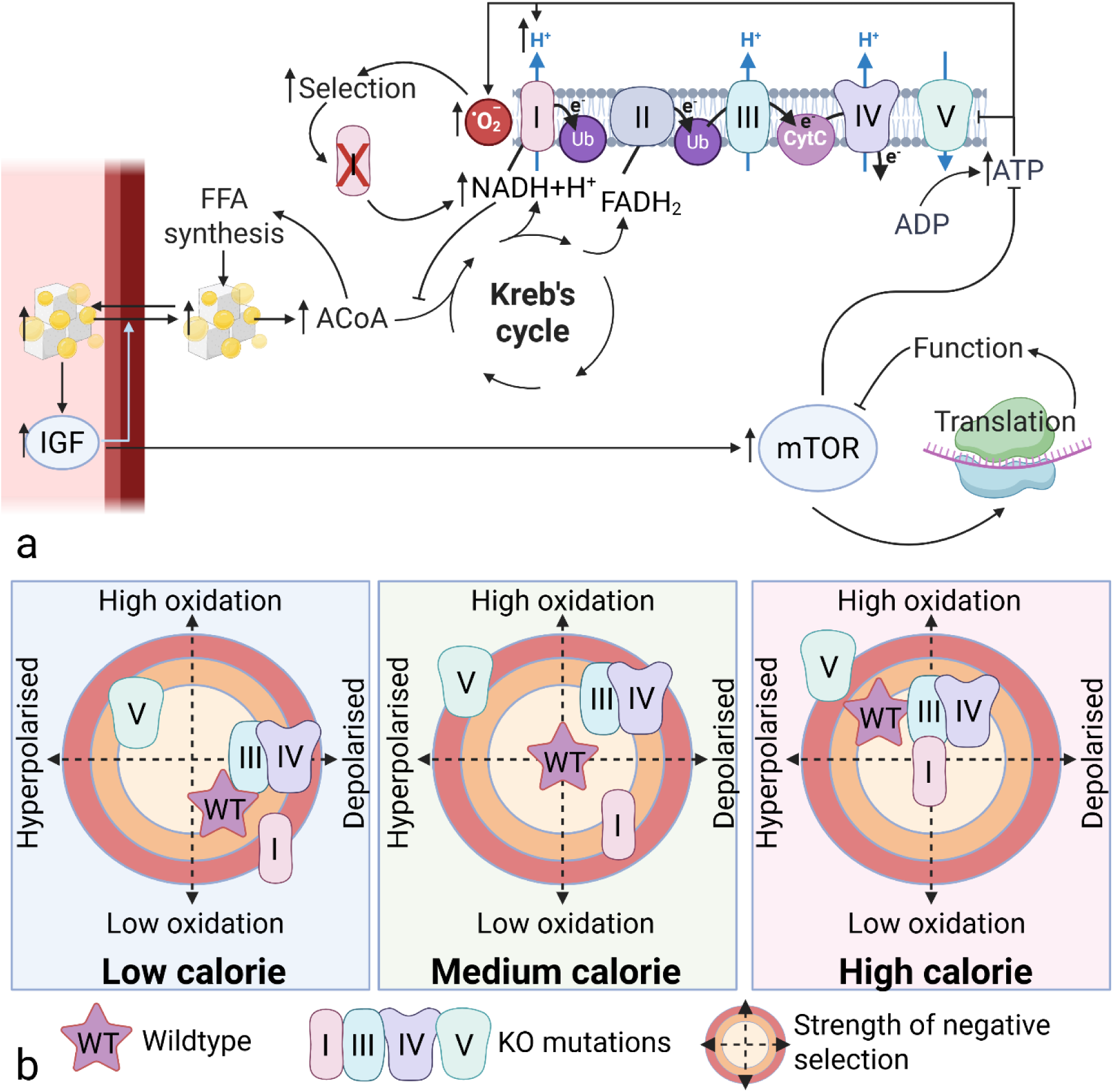
Effect of diet on mitochondrial selection. (A) Nutrient excess increases generation of NADH+H^+^, activating the ETC. However, despite initial activation of mTOR and anabolic stimulation, once functional metabolism is complete, negative feedback should prevent compensation by excessive ATP utilisation in order to speed up electron flow. ATP will therefore increase. Then as both ATP and NADH+H^+^ increasing induces hyperpolarisation and ROS production, it will increase selection for complex I mutations, inducing mitopenia. This blocks NADH+H^+^ oxidation, inhibiting Kreb’s cycle and further increasing nutrient excess. (B) The strength of selection for KO mutations of different individual mitochondrial complexes compared to a combined wildtype under different dietary conditions. Centre of the circle represents optimal positive selection. Moving further away from the centre indicates increasing selection against mitochondria of that genotype. Image created with BioRender.com. Abbreviations: ACoA, acetyl-coenzyme A; ADP, adenosine diphosphate; ATP, adenosine triphosphate; CytC, cytochrome C; e^-^, electron; FADH_2_, flavin adenine dinucleotide; FFA, free fatty acid; H^+^, hydrogen ion; IGF, insulin-like growth factor; KO, knockout; mTOR, mammalian target of rapamycin; NADH, nicotinamide adenine dinucleotide; WT, wildtype; Ub, ubiquinone.

However, while Kreb’s may initially accelerate, the rate of OXPHOS will depend on the level of ATP. While the same increased insulin/IGF signalling stimulating nutrient uptake will also stimulate anabolism, hydrolysing ATP, in the longer term cells do not simply synthesise macromolecules and induce anabolic signalling for the purpose of using ATP. Growth and anabolism are functional processes inhibited upon completion by negative feedback. Thus, we suggest that increased anabolism should not fully compensate for the nutrient-driven supply of ATP. As a result, ATP levels would rise, driven by the difference between increased nutrient supply and the need for anabolic metabolism.

As ATP rises, it will inhibit complex V and AMPK^177,180^, while Kreb’s remains proportionately increased. The higher levels of NADH+H^+^ and FADH_2_ will pass electrons to the ETC causing increased proton pumping. Meanwhile, less complex V activity slows the return of protons to the matrix, both processes polarising the membrane (Figure 14), as demonstrated by the activation of uncoupling proteins^504^.

ROS production is overwhelmingly associated with the hyperpolarised state, rising exponentially from complexes I and III as membrane potential passes 140mV^181–184^. The increased ROS likely reflect a negative feedback system as the ETC attempts to slow its rate of proton pumping by prematurely ejecting electrons and reducing the polarity of the membrane. Consistently, nutrient excess is also associated with high ROS levels^274^.

Just as conditions affecting cell death and proliferation shift the selective advantage between fast and slow cells, conditions affecting the polarity and oxidation of the mitochondrial membrane will shift the relative advantage of the mutant mitochondria degraded by them. The ultimate effects on selection will reflect the mechanisms which induce mitochondrial fission and degradation. Assuming basic threshold effects:

1. Membrane polarisation above a threshold would trigger fission and degradation
2. Membrane oxidation above a threshold would trigger fission and degradation.
3. Membrane polarisation below a threshold would trigger fission and degradation.
4. Membrane oxidation below a threshold would trigger fission and degradation.

In the hyperpolarised, hyper-oxidised membranes of a high calorie diet, complex V mutations are likely to be pushed well over thresholds 1 & 2, causing strong negative selection (see Figure 14B). Equally, complex III and IV mutations, which would reduce polarity, will still be selected against via threshold 2, due to the increased ROS production. However, mutations in complex I which prevent both polarisation and ROS production will be shifted away from thresholds 3 & 4, appearing increasingly functional as wildtype mitochondria are pushed upward toward thresholds 1 & 2. As the NADH+H^+^ cannot pass its electrons to the mutated complex, they can simply diffuse through the matrix to other complexes bound (by a leaky link) to functional nucleoids and exacerbate the disparity.

The excess of nutrients thus accelerates the selection for complex I mutations and mitopenia. Worse, as mitopenia reduces the capacity of mitochondria to accept the electrons from NADH+H^+^, Kreb’s cycle will slow, reducing the mitochondrial capacity to burn nutrients, worsening the nutrient excess which induced the mitopenia. Thus, mitopenia induces extrinsic ageing. This is (ironically in the context of this paper) a vicious cycle, whereby nutrient excess and mitopenia both worsen each other and inducing all the downstream effects of extrinsic ageing discussed in Parts 4 and 5. Consistently, it is now increasingly observed that C elegans ageing, like human ageing, involves significant lipid accumulation^505^.

It is worth noting that the vicious cycle is between mitochondrial selection and nutrient excess, while the short-term effects of these signalling pathways are homeostatic and thus adaptive rather than maladaptive. To reverse the vicious cycle would require either the loss of ATP homeostasis or a shift in selection for more oxidised and more polarised membranes, potentially inducing apoptosis.

While increasing mitochondrial mass would counteract nutrient excess by burning more fuel, this would increase the generation of ATP, which homeostasis would then restore by reducing mitochondrial mass. Only an upregulation of mitochondrial uncoupling should allow cells to counteract the vicious cycle. Consistently, exposure of C elegans to the uncoupling agents carbonylcyanide-3-chlorophenylhydrazone (CCCP) and carbonylcyanide-p-trifluoromethoxyphenylhydrazone (FCCP) both extended lifespan^506,507^, as did the overexpression of zebrafish ucp-2^508^. While overexpression of the worm’s own ucp-4 gene did not, this reflects that it has lost its uncoupling activity, functioning mainly as a succinate transporter^508^. Uncoupling has been widely associated with longevity in other organisms such as Drosophila^509–511^, mice^512^, and chronological lifespan in yeast^513^. In all cases the benefits of uncoupling were ascribed to a reduction in ROS production and damage, as suggested by the ‘uncoupling to survive’ theory of mitochondrial inefficiency^514^.

However, the four process model suggests instead that uncoupling reduces extrinsic ageing. It would predict that under conditions of fasting and DR, the selective pressures will switch in the other direction, advantaging complexes III and IV and disadvantaging complex I mutations as ROS levels are lowered. In these conditions, enforced uncoupling should be highly detrimental, further hiding complex III, IV, and even V mutations. Consistently, in DR experiments uncouplers reverse the longevity gain^515^, which might not be predicted by the ‘uncoupling to survive’ theory. Recent studies have also suggested that C elegans ageing is intricately linked to times of feeding^516^ when the vicious cycle of mitopenia and extrinsic ageing is occurring.

In addition to the direct effects of slowed catabolism on anabolism in the absence of increased mitochondrial mass, there may also be a feedback system between extrinsic and intrinsic ageing that we have thus far ignored.

As the vicious cycle between mitopenia and extrinsic ageing slowly increases nutrient excess, cells prevent toxicity by reducing their sensitivity to nutrient uptake. As described in Part 4, this is controlled by inducing IR. However, IR merely shifts the excess fuel outside of the cell. As adipose stores become saturated, and blood lipid and blood glucose remain higher for longer, it may increasingly overcome the IR that is blocking its entry into cells – particularly in tissues with significant insulin-independent uptake. This could increase IGF/mTOR axis signalling, inducing unwanted fuel uptake and unnecessary anabolism. Speculatively, cells may induce negative feedback to reduce their anabolism again, such as through epigenetic marks which induce metabolic slowdown.

This metabolic slowdown is notably not intrinsic ageing in the same way as epigenetic controls that result from selective destruction. It is potentially a short/medium term response, producing epigenetic marks associated with second generation (reversible) clocks, as described in Part 8.1, but it may reflect another retrograde signal by which the rate of mitochondrial catabolism epigenetically influences the rate of anabolism. It could help explain why post-mitotic organisms like C elegans are rejuvenated by a two chemical epigenetic reprogramming cocktail^517^, if the major underlying mechanism of ageing is mitochondrial

#### 7.1.6 Longevity mutations in mitochondrial metabolism

Interestingly, there are both long-lived and short-lived complex I mutant worms. If this reflects their effects on mitopenia, then worms with mutant complex I are likely born in advanced state of mitopenia, more vulnerable to extrinsic ageing through nutrient excess. If true, these worms are born at the equivalent of old age, which is possibly evidenced by their reduced rates of activity, decreased size, and increased developmental times^518^. Their reduced pharyngeal pumping rate likely protects them from extrinsic ageing in the same way as DR. When they are long-lived it could reflect that the key element inducing further mitopenia has been lost. These worms can experience no selection for complex I dysfunction because complex I is already dysfunctional, so the rate of ageing of complex I mutant worms, and clk-1 mutant worms, which have reduced ubiquinone synthesis (slowing transfer of electrons from complex I), is greatly reduced.

Notably, clk-1 mutant worms live longer when exposed to paraquat or through additional knockout of sod-2, but shorter when it is sod-1, while paraquat does not increase lifespan of isp-1 ETC mutations^519^. While stress is still the dominant narrative, we would suggest that these studies indicate it is not the major factor; its effects are dependent on metabolism, mediated partly by ROS as important signalling molecules.

An comprehensive review by Borkum^102^ looks at many of the mutations in the components of Kreb’s cycle which have effects on longevity. Although this should be an area of extensive future investigation, we suggest that longevity from mitochondrial mutants, or mutants with altered mitochondrial metabolism, could be induced by the following three main mechanisms:

1. Diversion of carbon out of Kreb’s cycle in a way that reduces the efficiency of conversion of calories to ATP.
2. Diversion of carbon away from the key activators of mTOR which stimulate anabolic metabolism and trigger nutrient-driven growth, which may then be repressed epigenetically.
3. Diversion of carbon through pathways that de-necessitate complex I, reducing the rate of selection for increasing mitopenia.

One further condition that may have relevance outside the mechanisms discussed here is effects on glycolytic relative to mitochondrial metabolism, which may have additional feedback loops and selection pressures related to hypoxia and carcinogenesis that are beyond the scope of this article. These aside, we suggest that mitochondrial ageing could largely reflect mitopenia resulting from mitochondrial selection, producing the fourth process of the four process model.

### 7.2 Cell senescence and other consequences of selection

There is one final clarification or addition to the four process model that may contribute significantly to the ageing process. While we addressed the impact of both growth and survival advantage in mitochondria, we focussed mainly on the growth advantage of cells. However, mutations in nuclear DNA can also confer a survival advantage.

Particularly in proliferative tissues, signalling is a balance between growth and death. Growth and survival signals are normally associated and positively reinforcing. While strong growth mutations outside the range of physiological norms can reverse the usual survival signal and induce apoptosis as in the case of several oncogenes^520^ and glucotoxicity^132^, outside these supra-physiological states, most pro-growth mutations also promote survival and inhibit apoptosis. However, there are survival molecules and pathways which are not linked directly with growth, where knockouts or activating mutations directly impact the cell death pathways associated with growth-independent factors such as DNA damage.

While pro-growth, pro-survival mutations result in immediate metabolic shifts according to the ever present growth signals which make the cells subject to selective destruction, conversely survival specific mutations (SSMs) can be silent until stimuli such as DNA damage attempt to activate apoptotic pathways. Such mutations are dangerous because they are relatively unobservable until cells suffer further damage.

As we discussed in Part 3, selective destruction could work through epigenetic slowdown or induction of cell death. Plausibly, a disadvantage to the latter is that it would exacerbate selection for SSMs, which may help explain why selective destruction has evolved to work via epigenetic growth control. Reducing the cell’s sensitivity to growth signals also resensitizes it to apoptotic ones without affecting selection for SSMs.

Despite initial hypotheses that ageing would result in increasing sensitivity to apoptosis as would be consistent with damage accumulation, currently evidence suggests it is tissue-specific^50,521,522^.

While pro-growth, pro-survival mutations can be neutralised by selective destruction, the natural course of tissue-level selection should still be for SSMs. In humans, around 10 billion cells may be created every day to replace those turned over by apoptosis^523^. Stem cells offer some protection from this selection as most of the replicating cells are differentiated cells that are doomed to die as part of the life cycle, such as in the gut and skin where the outermost cells are lost. However, tissues and organs should still reduce sensitivity to apoptosis with age due to the selective advantage of SSMs. Wherever this is not true, it could reflect counterselection.

The point where their apoptosis resistance produces visible changes in signalling, identifying SSMs to the surrounding tissue, is the point where the cell is being instructed to apoptose. Such conditions will include further damage inducing additional mutations in growth and survival signals that could induce transformation and allow the cell to escape tissue controls.

Thus we suggest, in line with previous observations^58,524^, that cell senescence has evolved to reduce the threat of selection for SSMs. In cells that are resistant to apoptosis, permanent arrest likely reflects the next best option. It reduces the risk of further mutation, while the SASP attracts immune cells such as NK cells capable of killing even apoptosis-resistant cells^525^. Both the SASP and Notch signalling may have a further purpose in removing other potentially dangerous cells. The bystander effect of senescent cells^60–62^ and cells hit by high levels of radiation^526^, known as the radiation-induced bystander effect (RIBE), induce senescence and apoptosis of surrounding cells. As the SSMs are hidden under normal metabolism, and neighbouring cells are more likely to be closely related or subject to the same mutagen, when a cell gains additional damage that induces its senescence, it may benefit the tissue to arrest or destroy the local cells, removing the threat that the same SSM persists within them. Senescence and the SASP might then have been co-opted into tissue remodelling programmes^527–529^, particularly as rapidly replicating myofibroblasts that are resistant to death could be potent inducers of fibrosis.

The capacity of senescent cells to spread damage and senescence within tissues is therefore plausibly the result of strong counterselection against SSMs, which may reflect the associated risks of such mutants. The accumulation of senescent cells with age may reflect, in part, the natural course of selection for SSMs, suggesting that targeting these pre-senescent cells may be equally worthwhile.

Other hallmarks and phenotypes of ageing may also reflect the courses of selection predicted by the four process model. Cells get larger with age^530^, which may reflect that such cells are slower to carry out metabolism or proliferate, or have diluted expression of markers and ligands that would cause selection against them.

Similarly, the amount of mtDNA decreases with age as does the distance between nucleoids^531,532^, meaning that the enzymatic complexes of the ETC are more dispersed, contributing to a depolarised, low-oxidised state that will undergo positive selection. While both cell size and mitochondrial nucleoid dispersal could reflect genetic changes, they could plausibly be independent even of epigenetic changes, reflecting selection at other levels.

Maintaining homeostasis is a difficult process. It is the purpose, and indeed speciality of life, by taking in energy to prevent internal entropy, that it does it so successfully, but we suggest that balancing selection and counterselection within changing conditions may be problematic, forcing many species to choose to allow one direction of selection over another, thus causing ageing.

## Part 7: Summary

The three process model applies mainly to mitotic tissues. Post-mitotic cells can differ in metabolic rate, but this should not imbue a selective advantage through increased proliferation, and thus does not threaten tissue homeostasis.

However, post-mitotic tissues have an intracellular organelle with DNA which is competing and subject to selection. We suggest the fourth ageing process may result from the selective destruction of mitochondria.

7.1.1 | The evidence for mtDNA damage in ageing is mixed, but argues against straight damage accumulation models.

7.1.2 | There is good evidence for selection occurring in mitochondria, particularly for large deletions in mtDNA.

7.1.3 | Mitochondria can malfunction in two main ways. Firstly, they can fail to add enough electrons to the ETC and pump enough protons across it; or they can fail to return the electrons and protons to the matrix.

Mitochondrial mutations in complex V, which cannot return protons, cause hyperpolarisation; mutations in complexes I, III, and IV, reduce proton pumping, causing depolarisation. Depending on conditions, wildtype mitochondria might choose to select against hyperpolarised mitochondria more strongly, allowing complex I, III, and IV mutations to spread.

Mitochondrial mutations in complexes III and IV, which cannot return electrons to the matrix, cause high ROS levels and membrane oxidation. Mutations in complex I prevent the ETC accepting electrons, reducing ROS levels and membrane oxidation. By selecting for depolarised, low-oxidised membranes, mitochondria can successfully select against complex III, IV, and V mutations, but over time may **select for complex I mutations over wildtype**, reducing mitochondrial function and causing **mitopenia**.

### 7.1.5 Reduced mitochondrial catabolism and reduced ATP production may reduce anabolism to maintain ATP homeostasis, inducing intrinsic ageing via the retrograde response

Nutrient excess exacerbates mitopenia by increasing the polarisation and ROS production of mitochondria, selecting more strongly against complex V, III, and IV, but also allowing complex I mutations to appear more functional compared to wildtype. The result is a vicious cycle where loss of functional complex I reduces NADH+H^+^ oxidation, increasing nutrient excess and thus inducing extrinsic ageing.

#### 7.2 The increase in senescent cells with age may reflect counterselection against cells which gain survival-specific mutations, preventing them from proliferating and being a threat

## Part 8: Implications of the four process model

In the final part, we will discuss the implications and limitations of the four process model. It is a purely theoretical study which is frequently speculative across multiple disciplines, all of which have intricacies and complexities that we could not possibly hope to address or even understand. Instead of adding to the complexity, the model is designed to simplify ageing to its most significant aspects. Many of the observations will be incomplete and even inaccurate, and if some observations are proved true by experimentation it should not be taken as evidence that the model is true in its entirety. Instead we suggest that the study provides a new mechanistic model of ageing that would allow us to move beyond exclusive reliance on the hallmarks and the complex multifactorial process, back to hypothesis driven research.

### 8.1 Progeria, intrinsic and extrinsic ageing, and the epigenetic clocks

Early ideas of ageing suggested that differences in lifespan reflected the number and type of mutations an individual inherited at conception^40^: organisms were not only ageing at different rates, but theoretically conceived at different starting ages.

Although these ideas may appear outdated, they are implicit to any theory of ageing which ascribes a major role to DNA damage accumulation. If ageing is the result of genetic changes, then organisms should be born at various ages by inheriting pre-damaged DNA. However, which damage would qualify as ‘ageing damage’ is unclear. While individuals born to older parents have higher risks of genetic disease and chromosomal abnormalities such as trisomy 21^533^, even cloned animals created from aged somatic cells produce organisms born at age zero with normal lifespans^534^.

While ageing could still result from genetic noise – i.e. production of mutant cells that behave differently to the starting cells, which we argued was unlikely in Part 1.1, the consistency of organisms born at age zero is inconsistent with the standard paradigm that ageing reflects the accumulation of DNA damage and activation of damage pathways that cause cell senescence and inflammageing.

However, while therapies designed to increase maintenance pathways and reduce damage have largely failed to positively affect lifespan, mutations that increase damage have a more sporadic effect. The progeroid syndromes are perhaps the best evidence for a role of DNA damage in ageing. However, even these are frequently described as ‘segmental’, due to doubts as to whether they reflect true accelerated ageing^535^. Among the most common outcomes are immunodeficiency, neurodegeneration, and cancer, with the latter being the most consistent outcome. In the four process model, cancer is produced by a separate process, suggesting that the progeroid syndromes resulting primarily in cancer would be more accurately interpreted as ‘pro-celeritic’ or ‘pro-carcinogenic’ diseases rather than progeroid per se.

Although neurodegeneration is a common feature of several, the mechanism may not the same as in the age-related neurodegenerative diseases; for example, Cockayne syndrome shows no evidence of amyloid staining or other protein plaques^536^. As nucleotide excision repair affects mtDNA as well as nuclear, it could accelerate mitochondrial damage and selection consistent with age acceleration via the four process model, but the major feature appears to be neuronal cell death^537^. When combined with other outcomes such as developmental delay, dwarfism, cachexia, marrow failure, bone loss, and immunodeficiency, the picture created is one of primary failure in both highly proliferative and post-mitotic tissues, where elevated cell death prevents the maintenance of cellularity, which in the four process model is not a feature of intrinsic ageing.

Hutchinson-Guildford Progeria Syndrome (HGPS) results from a mutated nuclear envelope protein, LMNA, creating highly unstable cells^538^. It does not cause neurodegeneration, consistent with its major effects in a non-mitochondrial pathway. The major progeroid outcome is atherosclerosis which results in death from stroke or myocardial infarction. However, cardiovascular disease in HGPS does not share the same risk factors as normal ageing^539^. Arguably, the pro-fibrotic HGPS plaques from smooth muscle cell loss, filled collagen and proteoglycan^540,541^ do not reflect the same underlying mechanism for fatty plaques we described in Part 5.1. Instead, HGPS may be inducing cell death more strongly than cell ageing, resulting in endothelial dysfunction via the same mechanisms as COVID and smoking. If true, the HGPS plaques would share more mechanistic overlap with the sudden cardiac events associated with youth and exercise – notably associated with reduced age-related cardiovascular risk factors^395^. Highly fibrotic conditions in the lung, liver, and elsewhere are associated with continuous cycles of cell death and proliferation which necessitate a constant scaffold to repopulate the damaged tissue. Thus, we suggest that the progeroid syndromes are not true progerias, instead reflecting an even more fundamental loss of homeostasis than energy levels – the loss of cellularity.

Despite this, these diseases are the main causal evidence for DNA damage accumulation as the underlying cause of ageing. This point is best emphasised by quoting from proponents of the theory. A recent review by Schumacher, et al.^542^ summarises this point in the concluding section entitled, ‘**Is DNA damage the primary cause of ageing?**’, which in paragraph one states, ‘*The strong mechanistic link between DNA damage and ageing, and the role of DNA as the primary template for all cellular functions, suggest DNA damage as a candidate for the primary cause of ageing. However, there are at least three important arguments against this proposition.*’ The bulk of the section then describes evidence against DNA damage accumulation as the major cause of ageing before returning to two points in its favour. Firstly, a meta-analysis of genome-wide association studies identifies maintenance pathways as associated with age-related diseases, which the authors state is ‘*consistent with the observation in both humans and mice that the vast majority of rare genetic progeroid syndromes, in which multiple, bona fide ageing-associated diseases develop early in life, are caused by mutations in DNA repair genes*’. Schumacher, et al.^542^ then conclude that, although the three arguments against damage accumulation are ‘not invalid’, they are ‘unconvincing in view of the sheer complexity of DNA repair processes’. However, without more precise argument it is unclear why complexity should affect the validity of the counterevidence more than the evidence. Consistently, other proponents have accepted that damage has stopped being a major focus of ageing research due to the ‘complexity of the aging process’^543^.

Crucially, we are not suggesting that DNA damage *per se* is not responsible for ageing – only that the mechanism could be updated. As noted by Vijg^17^, while the progeroid syndromes are segmental, defects in DNA maintenance pathways are the only ones that reliably and repeatedly induce premature ageing-like syndromes, which is a significant observation.

While we suggest that the primary outcome of progeria reflects cell death and cell loss, the four process model unequivocally predicts that **increased rates of damage should accelerate ageing**, as accelerated mutation will produce more faster and slower cells. Competition, selection, and counterselection should then result. Notably, the evidence that exogenous damage from radiation beyond the hormetic range accelerates ageing is controversial. Several studies suggest it increases cancer rates^544,545^, and overall mortality but without affecting ageing per se^44,45,544^. However, more recent studies have suggested otherwise, indicating that γ and X-irradiation can accelerate age-related diseases even beyond the expected increases in cancer^546,547^. In such cases, the phenotypes are less extreme than the progeroid syndromes, which may reflect the range of damage where cellularity is maintained, but selection and counterselection between the additional mutants are increased.

When looked at from this angle, the four process model can explain many of the inconsistencies and paradoxes for DNA damage in ageing. Oxidative damage is an important biomarker of mitochondrial function, and reducing it can negatively affect selection, while small increases in damage activate FOXO, shifting metabolic equilibrium to a pro-longevity state and potentially affecting selective destruction or its outcomes, while at the same time preventing much of the extra damage that would induce additional selection and counterselection. However, once levels of damage begin to overwhelm the repair capacities, higher levels of mutations will increase selection and counterselection, but also quantities of cell death, necessitating more proliferation and shifting the advantage to faster cells. The expected results would be celerisis, fibrosis, and cancer with smaller increases in intrinsic ageing that leads to other age-related diseases. As levels of damage increase further, it could begin to induce sufficient cell death that ageing as a homeostatic shift toward slower metabolism becomes less significant than the body’s ability to maintain enough cells to keep organs functioning appropriately. The resultant diseases should be fibrotic but only carcinogenic if the cells are sufficiently stable to allow rapid growth.

While ageing is induced by damage, celerisis is the only ageing process that reflects genetic change. Intrinsic ageing is an epigenetic phenomenon that is wiped clean in the reprogramming of the zygote and early embryo to produce totipotent cells^548^. Thus, animals with mutations in the zygote can be at risk of early cancers, but even cloned animals are born at age zero. Equally, the Yamanaka factors rejuvenate cells for the same reason – both ageing and differentiation are controlled by epigenetics and removed by dedifferentiation. As de Magalhães^549^ suggests, ageing is a ‘software’ problem, not a ‘hardware’ problem.

Consistently, de Magalhães^549^ also argues that cancer is the exception, requiring a ‘cellular balancing act’ that may necessitate reduced rejuvenative capacity which would accelerate ageing. Indeed, the idea that ageing is an anti-cancer mechanism is not novel. Argued most persuasively by Wolf^550^ in the tumour suppressor theory of ageing, there have been hints of pleiotropy between cancer and ageing from studies into telomeres^551^ and p53^552–554^ and trade-offs between regeneration and cancer which could necessitate ageing^555^. The four process model fits neatly with these preceding ideas. However, the mechanistic paradigm utilised by these older models is one of cell loss, cell senescence, and inflammageing^550^, which we suggest may not explain the ageing phenotype.

Alternatively, de Magalhães^549^ argues that ageing is the result of a developmental program, which explains both why the epigenetic clocks begin shortly after conception^556^, and why ‘genes in the vicinity of epigenetic clock methylation sites are often related to growth and development’. However, just as we have argued against the TCQP, ageing is unlikely a continuation of development because growth and development are hyperfunctional processes. As noted by West, et al.^555^ ‘numerous metazoans with developmental programs of continued growth following reproductive maturity show negligible aging’, which was the underlying logic for Bidder’s hypothesis that ‘we die, therefore, as an alternative to becoming giants’^557^. In short, ageing particularly as a phenotype of hypofunction^88^ and an anti-cancer balancing act is more reflective of the cessation of development than its continuation.

Conversely, the Information Theory of Ageing suggests that ageing results from epigenetic changes induced by DNA damage^558–560^, and suggests that the epigenetic changes are random rather than programmed. The four process model would suggest that epigenetic noise is as unlikely to produce the defined phenotypes of intrinsic ageing as genetic noise. As argued by de Magalhães^549^, the capacity of the clocks to so accurately predict chronological and biological age is evidence that a defined process is producing specific epigenetic marks associated with growth and development. However, that process should not be development itself, but a restraining force preventing excessive growth and hyperfunction. We suggest that selective destruction is currently a strong candidate for this specified process. The damage necessitating the program is stochastic as in the Information Theory, but most of this damage is inconsequential. At worst it will increase noise, but in many cases where the damage actually affects function, the cell is simply killed and replaced, as indicated by the stark increases in damage accumulation at non-functional regions vs functional ones^561^. Conversely, damage influencing selection will not only allow the cells to survive, but encourage their spread. The process evolved to prevent this should therefore produce epigenetic marks at the precise points required to counteract this shift – either maintaining metabolism at the wildtype rate or reducing it when the latter is safer.

Thus, we suggest that the four process model combines key aspects of both damage-based and programmatic theories, forming a kind of hybrid that, if accepted, could help unite the field by explaining the evidence consistent and inconsistent with both preceding major theories.

If true, then selective destruction could be producing the epigenetic clocks or aspects of them. However, there are an increasing number of epigenetic clocks, including the first generation clocks developed by Horvath^562^ and Hannum, et al.^563^ designed to measure chronological age (i.e. time since birth) and second generation and beyond clocks designed to measure biological age (i.e. risk of death from age-related causes) with increasing accuracy.

If ageing is a single process, then we would predict that second generation and beyond clocks are better measures of ageing, but if ageing is split into intrinsic and extrinsic ageing, then as suggested by Figure 8A, the increasing relevance of extrinsic ageing in later life would suggest that clocks which better predict mortality are those measuring extrinsic ageing. Intrinsic ageing, as a steady decline in metabolic rate, less affected by behaviour, is more likely to map to chronological age and thus be better measured by first generation clocks. Consistently, Poganik, et al.^564^ recently demonstrated that exposure to various stressful stimuli would transiently increase biological age as measured by second generation clocks, but not the chronological age measured by first generation clocks. The epigenetic effects of extrinsic ageing should be transient – they are responses to short-term stimuli such as stress which might change swiftly and thus need to be removed swiftly. Conversely, first generation clocks would reflect marks induced by selective destruction created to semi-permanently influence a cell’s metabolic rate, which would be useless if the cells could easily remove them.

Consistently, therapies which extend maximum lifespan such as CR and rapamycin slow first generation clocks^565,566^, while exercise and metformin which only extend median lifespan do not^565,567^. Obesity and the high-fat diet, conversely accelerate the Horvath clock^567,568^, as predicted if nutrient overload is having the reverse effect to CR on metabolic rate, while irradiation has no effect^565,569^.

One observation not entirely consistent with the four process model is the initial observation by Horvath^562^ that multiple cancer samples showed significant age acceleration, averaging 36 years ahead of non-cancer samples. We would have predicted that cancer cells might have lower (first generation) epigenetic age as they remove the genetic and epigenetic barriers for growth.

While the inconsistency is worthy of further investigation, a more recent study did at least suggest that the Horvath clock was decelerated in multiple cancers^570^. Fascinatingly, Herzog, et al.^570^ also showed that people who developed cancers had reduced epigenetic ageing across multiple clocks in other non-cancerous tissues, suggesting reduced epigenetic ageing may predispose individuals to cancer, in line with the relationship between celerisis and intrinsic ageing, even if it does not reduce the age of transformed cells themselves. Another study similarly observed a distinct (near significant) trend toward reduced first generation clock epigenetic age as the stage of liver fibrosis advanced^571^, also consistent with celerisis. One important prediction from the four process model is that Notch signalling will be a crucial mediator of first generation clocks, and knockouts may slow their epigenetic ageing. Notably, links between FOXO and Notch could be a particularly enlightening area for further investigation.

### 8.2 Evolution of longevity

A unifying theory of ageing should predict why lifespans differ across the evolutionary tree. Peto’s paradox reflects that larger organisms are longer-lived despite having more cells, which should greatly increase their risk of cancer^572^. The association between increasing size and longevity is so significant that the impact of any other factor on lifespan is often assessed by comparison with organisms of similar size using the longevity quotient (LQ), with an LQ of 1 suggesting the average lifespan for an organism of that size^573^. The LQ was created because size was a proxy measure for metabolic rate, which according to the rate of living theory, first suggested by Rubner^574^ and formalised by Pearl^575^, suggested that all organisms had an ‘inherent vitality’ depleted by their rate of growth, and every organism would expire after roughly the same amount of metabolism.

Pearl’s theory remained dominant for some time, with substantial correlative evidence across the evolutionary tree; however, the observation that birds had significantly higher metabolic rate and lived significantly longer than mammals, perhaps expending 3.5 times the amount of energy per gram of body mass over a lifetime^576^, and the observation that exercise increased survival at the same time as increasing metabolic rate^233^, suggested there was no set limit to lifespan determined by amount of metabolism. Rate of living theory was succeeded by free radical theory, which explained the additional lifespan of birds by proposing that birds could have higher levels of defences against ROS, which were the toxic products of metabolism limiting lifespan. Exercise became hormetic. Metabolism was replaced by damage, which survives as the dominant explanation for ageing despite the mounting counterevidence.

We suggest alternatively, that rate of metabolism determines the relative difference between fast and slow cells, influencing the rate of selective destruction and thus intrinsic ageing. Consistently, lifespan of ectotherms is significantly increased at low temperatures^231,577^, and experiments in endotherms suggest genetically dropping body temperature also increases lifespan^578^.

Less consistently, housing endotherms at higher temperatures also reduces survival, despite lowering metabolic rate^579^, suggesting the effects of temperature lie outside its effects on metabolic rate^580^. Potentially, this reflects differences in anabolism and catabolism: low temperature thermogenic requirements may shift metabolism toward uncoupled mitochondrial metabolism, burning fuels without producing ATP. If cold temperatures activate FOXO, it could have multiple beneficial effects that also impact rates of selective destruction. Although it is an area for further investigation, there may also be additional trade-offs between types of anabolism operating independently of its total rate, such as between RAS stimulated proliferation vs. mTOR stimulated protein synthesis, which may further impact the rates and outcomes of selective destruction, potentially explaining the maximum lifespan extension of the RAS pathway inhibitor, trametinib^581,582^.

Notably, even if large animals live longer by virtue of lower metabolic rates or having more cells – which would require longer to slow, particularly in post-mitotic tissues where selection is intracellular – this would not resolve Peto’s paradox, as larger organisms should still get cancer earlier. Regardless, evidence from dogs suggests it is unlikely. Large dogs die significantly earlier than small dogs, not just due to increased rates of cancer (and celerisis), but also suffer other age-related diseases earlier as well, indicative of increased intrinsic and extrinsic ageing.

Despite this, wolves are both larger and longer-lived than dogs. Potentially, when a mutation initially induces large size, the budding species is more susceptible to both celerisis and intrinsic ageing. Large size is produced by increased IGF-mTOR axis signalling, which would shorten lifespan as it does in dogs. However, in the long-term, natural selection (but not necessarily artificial selection) should respond to the reduced extrinsic mortality of the larger mutant and its offspring which are more resistant to being eaten, cold, and starvation etc. Existing longevity inducing alleles and de novo mutations can now undergo positive selection as they did not (or would not have) in the smaller species, allowing the larger organisms that possess them to outcompete the large organisms which do not.

Various evolutions could reduce the larger organism’s susceptibility to cancer, from increased redundancy in the cell senescence and apoptosis pathways, as in elephants and whales^583,584^, loss of telomerase expression in increasing populations of cells^20^, or more rigorous detection mechanisms against celerisis. However, in the latter case this may only further accelerate intrinsic ageing.

We suggest that to reduce its rate of intrinsic ageing an organism must reduce its rate of epigenetic selective destruction. However, there are several ways this can be achieved without increasing the risk of celerisis. Increased sensitivity to apoptosis could increase longevity by reducing epigenetic slowdown during cell competition, instead relying on cell death and replacement to maintain homeostasis, consistent with recent observations in long-lived bats^585^, although this would increase selection for SSMs. Reduced metabolic and proliferative rate, with increased protection from genetic mistakes and removal of mutant cells, could all reduce the rates of intrinsic ageing. However, all these evolutions could have a single major disadvantage, significantly extending development times.

Indeed, a major mechanism by which development is extended is by reduced IGF signalling. We can see this in multiple organisms, from the inhibition of daf-2 in C. elegans stalling development and causing dauer formation^586^, to dogs where a key difference between dog breeds is in the levels of developmental IGF-1^587^. Bigger, stronger animals and faster development all involve boosted IGF-1 levels. Thus, as a large organism evolves and has its lifespan reduced through increased IGF signalling, then natural selection for longevity should spread the IGF signal over a longer period, increasing lifespan in conjunction with the length of development.

Consistently, wolves have extended developmental period compared to dogs, around 22 months to mean 12.5 average for dogs^588^, and squirrels, which live around eight years compared to the 2-3 years of similarly sized mice and rats, can have developmental periods over a year^589^ compared to 2-4 months for mice and rats. Slowing developmental rate allows slower, more accurate cell and DNA replication followed by a process of more rigorous checking. Additionally, time and energy can be put into removing and replacing imperfect cells rather than simply slowing their metabolism. While these ideas are also consistent with developmental theories like TCQP, the decline in IGF signalling in human adulthood^590^ is more consistent with hypofunction and the four process model.

Maintaining genetic consistency through cell death and replacement is likely most crucial during development, and selective destruction could be described as a mainly developmental program, but it will remain relevant after adult transition. It does not rule out the existence of additional quasi-programmes and antagonistic pleiotropies, including energy trade-offs.

Detrimental mutations can still accumulate through drift if they have little to no effect in wild populations. After intrinsic ageing evolved, producing an intrinsic limit to lifespan, the fitness costs associated with other independent late-acting mutations would be reduced, allowing other antagonistic pleiotropies to evolve with smaller early life benefits, potentially allowing additional kinds of degeneration.

Semelparity could reflect one example. Whether death shortly after mating reflects accelerated wear-and-tear or a death programme has been debated extensively^6,591^. The four process model suggests it could simply reflect accelerated extrinsic ageing without intrinsic ageing. For example, the male antechinus enters a state of frenzy during mating which is usually lethal, and even produces Alzheimer’s like protein plaques in the male brains inducing a type of dementia^592,593^. However, if a male is kept in the lab, isolated from other males under optimal conditions, it can escape its death and live for an additional two years, more than doubling its lifespan^594^. Although the male is sterilised, after the frenzy is over it is otherwise unharmed, capable of living to the next mating.

Potentially, stress hormones such as cortisol are inducing an accelerated bout of extrinsic ageing, which, through the pathways we have suggested in Parts 4.5 and 5.2, could produce protein plaques in the brain, which would then reverse when the stress signalling declines.

Semelparity is unlikely to reflect the necessities of tissue-level selection, suggesting distinct reasons for evolving such as benefits to their kin^77^ or even population stability^78^. It is also possible that additional ageing processes evolved and preceded intrinsic and extrinsic ageing. In primitive multicellular organisms where unselective and selective destruction were likely evolving to control rogue cells in newly formed somas, other control processes could have evolved independent of the IGF/FOXO pathway and the longevity module. In immortal hydra, FOXO is crucial for maintaining stem cell pools, but some species can undergo rapid decline after sexual reproduction by a process called ‘inducible ageing’, which may be dependent on pathways outside the longevity module^595^.

Notably, primitive organisms such as hydra, flatworms, and jellyfish have low levels of tumour formation despite significant regenerative capacity, which is one of the few results we have recognised which is not overtly consistent with the four process model. An organism that lacks intrinsic ageing should demonstrate significant risk of celerisis. We suggest that there could be multiple explanations for the tumour-resistance of simple animals, but perhaps complexity itself may reduce the capacity of tissues to identify and remove mutant cells. In a tissue of one cell type, with shared function and activity, it may be easy to spot cells that are behaving aberrantly, but when organisms have “1,264 separate cell groups, comprising the 400 major cell types across 60 tissues”, as in the human body^596^, it may be increasingly difficult to identify the effects of mutation from positional effects relative to blood vessels and other cell types and differences in differentiation. Thus, negligible senescence may give way to ageing.

### 8.3 Fuel and Energy Model

The significance of the four process model is that we can begin to think about ageing as specific processes allowing us to return to hypothesis driven research aimed at systems level interpretations. As our understanding grows, we can also begin to produce computational models assessing ageing itself rather than ‘themed’ models.

Myriad top-down systems studies have provided the full detail of which components of the various ‘omes’ associate with ageing, and they have successfully reinforced that it is a complex, multifactorial process. While these are excellent studies for identifying biomarkers and therapeutic targets, they are less well designed to advance our understanding of the underlying mechanisms. For this, we need bottom-up systems studies building networks which first simplify the systems and then increase the complexity as our understanding increases.

Our prolonged focus on damage and disease has created a picture of inevitability about ageing as a system of unavoidable positive feedbacks that seem specifically designed to produce disease. Yet such seemingly maladaptive processes would be unlikely to evolve. Life is maintained by negative feedback and the persistence of homeostasis, and factors associated with disease should be assessed in this context. They are more likely to be trade-offs where a dysfunction in one system allows a more important one to retain homeostasis.

In Part 9, we have included a toy model of the basic energy regulation systems and shown how metabolic slowdown leads to extrinsic ageing. The Fuel and Energy Model (FEM) is based on mathematical representations of equations 1-4 from Part 4, and could form the basis for expansion. We suggest that the primary goal of such models should not be to mirror the complex web of known interactants as closely as possible, but produce the simplest possible system that replicates the rough dynamics of the biological system, and crucially maintains homeostasis. Before any new molecule or complexity is added, it should be clear what functionality it produces(by either enhancing homeostasis under the current conditions or responding to additional conditions). Only by learning how we keep ourselves the same can we begin to understand the fundamentals of why we change and eventually fail.

## 8.4 Discussion & conclusion

Biogerontology is arguably in a post-explanatory state. Much of the attention is directed toward omic data gathering, biomarker identification, and translational research. We have come to believe that ageing is too complex, reconciling inconsistent data because it is consistent with complexity.

The four process model is designed to produce a basic mechanistic interpretation of ageing strengthened by an evolutionary framework. Of course, it is not without considerable limitations. Firstly, it covers such a breadth of disciplines that over-simplifications, inaccuracies, and omissions are inevitable. We hope that experts in each field will first try to adapt the model with corrections before concluding its invalidity. Secondly, dividing ageing into four processes de-emphasises the interdependence of the processes. Equally, our attempts to demonstrate that ROS, DNA damage accumulation, and cellular stress are not key factors in the ageing processes has likely unfairly marginalised and discounted their various contributions, just as previous models may have overestimated them.

Lastly, we have suggested that selective destruction is orchestrated by Notch signalling, though our understanding of cell communication and competition is still in its infancy. There could be many contributing pathways, including forces like bioelectric current which could allow accurate comparison across multiple cells^597^, and perhaps others we are yet to imagine.

However, for readers unconvinced by the necessity of selective destruction or the forces of selection in tissue ageing, we would still suggest that splitting ageing into intrinsic and extrinsic ageing adds significant predictive value.

While such theoretical work will doubtless include many errors, it also holds advantages. Viewing organisms as simple functional systems maintaining homeostasis within constraints strips away the complexity to produce a clear chain of events from health to age-related disease. It creates a context which fills the gaps between experimental findings, combining evidence while removing paradoxes. If the four process model proves to have some validity, then our understanding of ageing will have shifted from widespread disagreement about fundamental questions to a position of basic, common knowledge. Ageing would no longer be a complex multifactorial process or a collection of hallmarks, but four defined processes with perhaps more to follow.

## Part 9: Underlying model of metabolic slowdown

To test the predictions of the four process model, we created the Fuel and Energy Model (FEM) based on mathematical representations of equations 1-4 and their respective equations S1-4 defined and justified in the methods according to the evidence below.

The main purposes of the model are:

- Produce a system that maintains homeostasis for key metabolites which includes the major regulatory pathways and molecules as evidenced by literature review.
- Demonstrate that within this system metabolic slowdown induces the key age-related pathologies of mitochondrial dysfunction, weight gain, insulin resistance, and rising inflammation, as well as age-related disease.
- Demonstrate that the model includes all necessary components to assess the mechanism by which anti-ageing therapies may affect morbidity and mortality.

### 9.1 Methods

#### 9.1.1 Model construction

The FEM was constructed using ordinary differential equations (ODEs) based on a non-systematic review of energy homeostasis literature. As with all models, this involved simplifying the biological detail to include only the central effectors of the described processes. A model schematic is shown in Figure 15.

**Figure 15.**
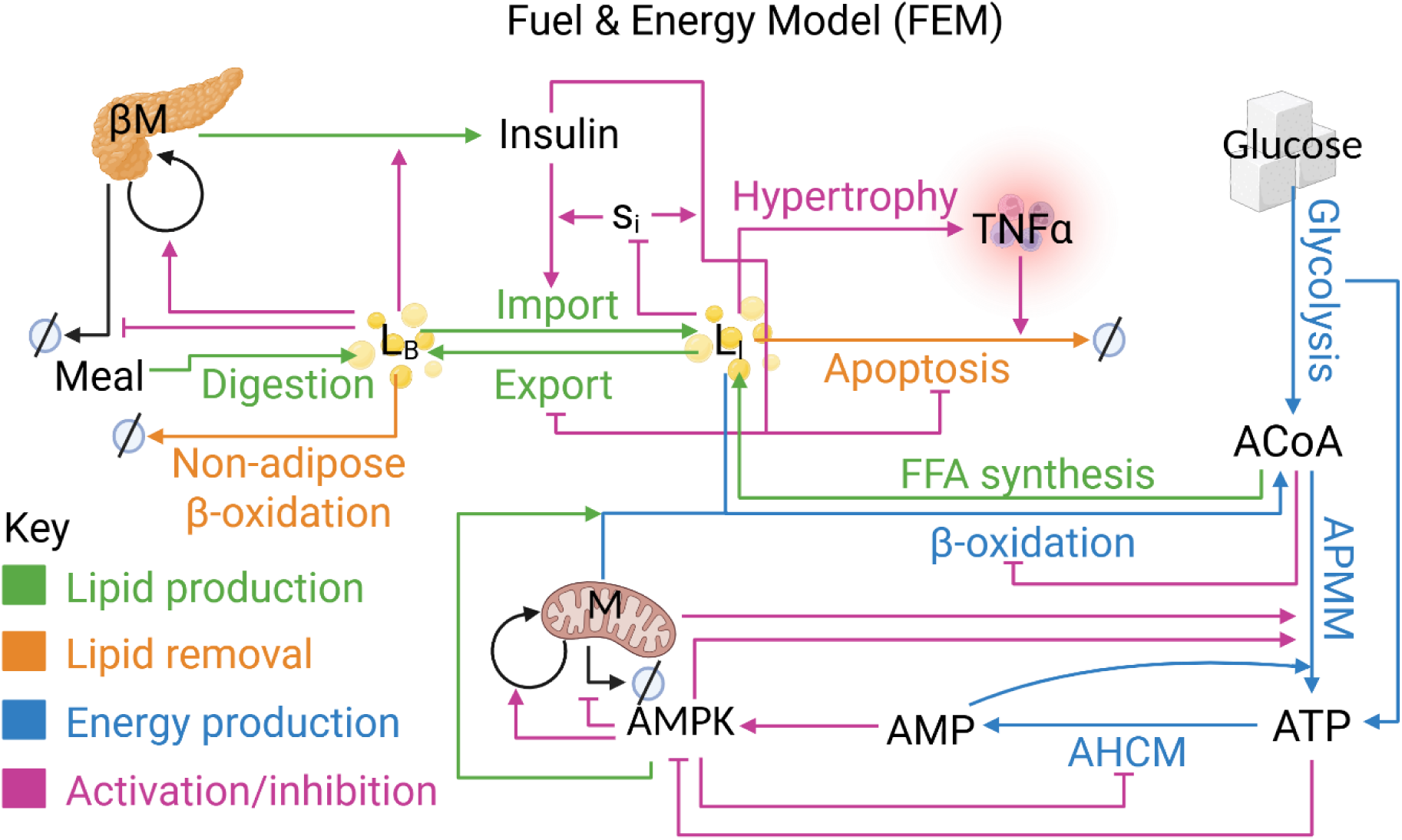
Fuel & Energy Model (FEM) network. L_B_ is increased by digestion of meals and export of stored lipid from the adipose, L_I_ back into to the blood. It is decreased by import into the adipose tissue to form L_I_, and import into non-adipose tissue for β-oxidation. L_B_ inhibits the apoptosis of β-cells (βM) and induces their proliferation while stimulating β-cells to produce insulin. L_I_ is reduced mainly by apoptosis, where increasing L_I_ induces adipocyte hypertrophy which causes secretion of cytokines such as TNF-α which stimulate their apoptosis (and removal of fat by reverse cholesterol transport). Insulin inhibits both apoptosis and the export of L_I_, preventing lipid loss. L_I_ also reduces the insulin sensitivity, s_i_, of the cells storing it, while the effect of insulin on all the processes it regulates (import, export, and apoptosis) is dependent on the insulin sensitivity of the adipose. Lastly, L_I_ is increased by FFA synthesis from acetyl-coenzyme A (ACoA) and reduced by the β-oxidation of FFAs to ACoA, while ACoA inhibits β-oxidation to provide negative feedback. ACoA is additionally formed from glycolysis and converted to ATP in the mitochondria through Kreb’s cycle and oxidative phosphorylation, which we have collectively termed ATP producing mitochondrial metabolism (APMM), though glycolysis can also produce smaller amounts of ATP. APMM is activated by AMP and AMP kinase (AMPK), while AMPK also stimulates increased mitochondrial capacity and prevents its reduction through a mixture of processes (some by increasing mitochondrial mass, such as biogenesis, but others by affecting membrane efficiency, such autophagy, uncoupling, and ROS regulation). AMPK also stimulates the β-oxidation of L_I_ which is dependent on the mitochondrial capacity, as is APMM. Conversely, AMPK inhibits the ATP hydrolysing cellular metabolism (AHCM). Images created with BioRender.com.

##### 9.1.1.1 Main equations

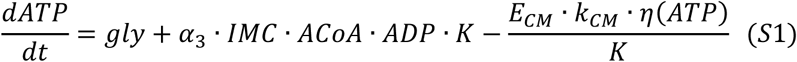

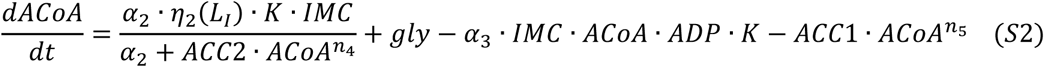

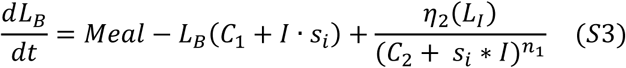

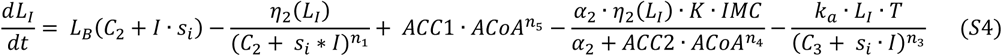

##### 9.1.1.2 Other ODEs

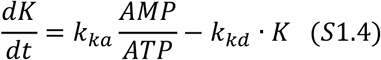

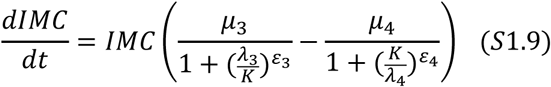

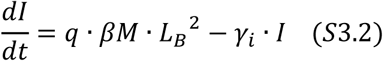

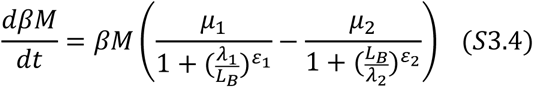

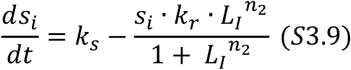

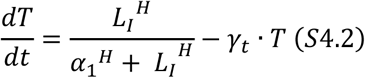

##### 9.1.1.3 Other equations

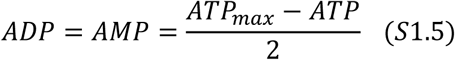

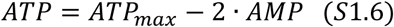

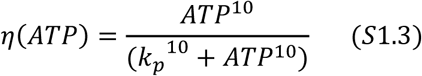

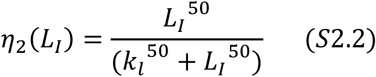

##### 9.1.1.4 Metabolic slowdown

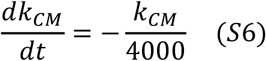

##### 9.1.1.5 ATP dynamics

***Adenosine triphosphate (ATP) dynamics*** are defined by the processes in equation 1 (shown here to aid interpretation) and described in equation S1.

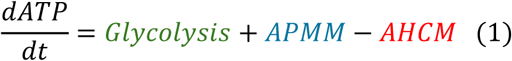

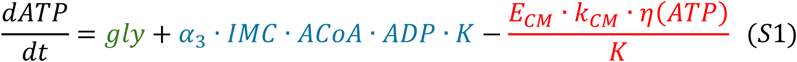

**Glycolysis** is modelled as the parameter **gly**, as in equation S1.1, defining the rate at which ATP is produced by the glycolysis of glucose.

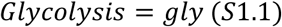

Glycolysis is the production of pyruvate from glucose (and here includes the subsequent step producing acetyl-coenzyme A, ACoA, from pyruvate), which also produces a small amount of ATP. We kept gly at zero for model simulations, choosing to focus entirely on lipid dynamics, so the same parameter could be used for ACoA production (see below).

**ATP hydrolysing cellular metabolism** (**AHCM**) was defined as the non-mitochondrial and mitochondrial metabolism requiring the hydrolysis of **ATP** (i.e. the metabolism using up the bioavailable energy, rather than producing it), as in equation S1.2.

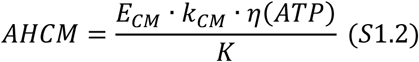

We used a single parameter, **k_CM_**to define the basal metabolic rate. The **E_CM_**parameter defines the tissue’s upregulation of AHCM in response to external cues (the capacity limited by k_CM_, NB: using two separate parameters allowed us to manipulate these aspects in combination). Within this capacity, AHCM can still be regulated by inhibition via AMP-activated protein kinase (AMPK) activity (K, described below), as is well documented^178,598^.

To some extent, AHCM must be dependent on the level of its substrate, ATP. However, the rate of AHCM cannot be allowed to fluctuate according to the energetic availability. It must reflect largely the metabolic needs of the cell. For example, the production of alcohol dehydrogenase to detoxify alcohol should be primarily a reflection of the quantity of alcohol, not ATP. Thus, ATP is maintained well above rate limiting concentrations. We describe this dependence as a switch function **η(ATP),** as in equation S1.3. This allows AHCM to occur at maximal (normal) rate while ATP levels are higher than the (low) threshold determined by the parameter, k_p_, dropping quickly to zero when ATP drops below it. Notably, if ATP is maintained at equilibrium, the levels should never drop below the threshold and η(ATP) = 1 would be an acceptable simplifying substitution.

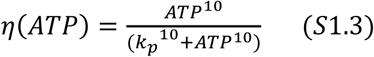

**AMP-activated protein kinase** (**AMPK**) **activity** is described by variable **K**, inhibiting AHCM. K is determined by the ratio of **adenosine monophosphate** (**AMP**):ATP, regulated firstly by upstream kinases and phosphatases phosphorylating/dephosphorylating AMPK’s activation loop, and secondly by phosphorylation-independent, allosteric kinase activation^599^. While ATP and AMP affect both processes, ADP plays a more minor role, mainly protecting against dephosphorylation without contributing to allosteric activation^600–603^. Therefore, we have only considered AMP and ATP in the activation of AMPK. As shown in equation S1.4, AMPK activation is determined by the ratio of AMP:ATP, while the rate of deactivation increases linearly with active AMPK (corresponding to a spontaneous deactivation process with a fixed average rate).

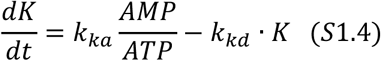

ATP regulation appears to be highly robust; as described by Hancock, et al.^175^, “the rate of ATP hydrolysis is matched by the rate of ATP supply without measurable changes in ATP over a wide range of ATP demands (up to very intense short-term near-maximal exercise)”^176^. Outside the conditions of intense exercise, AMPK can regulate metabolism and respiration to maintain the correct ratio of ATP:ADP/AMP without affecting the size of the adenosine phosphate (AP) pool.

Adenylate kinase reversibly converts 2 ADP to one AMP and one ATP, maintaining equilibrium levels of ADP and AMP under most conditions^604^, which for simplicity we have assumed is a 1:1 ratio so AMP ≈ ADP. This allows us to ignore the least important regulator of K, ADP, as shown in equation S1.5-1.6, where **ATP_max_**is the size of the AP pool, and thus the maximum amount of ATP that could be produced. Thus, the AMP:ATP ratio in equation S1.4 is defined as in equation S1.7.

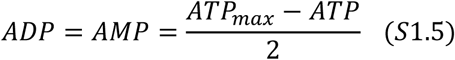

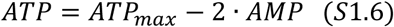

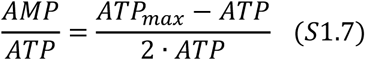

**ATP producing mitochondrial metabolism** (**APMM**) reflects all metabolism that results in net production of ATP within the mitochondria. Here, we have separated this from the processes of glycolysis and β-oxidation. Glycolysis is usually defined by the production of pyruvate (and 2 ATP); however, for simplification of ACoA dynamics (see equation 2), we have defined both processes to include the following step where pyruvate (and acyl-coenzyme A for β-oxidation) are converted to ACoA, which occurs within the mitochondria. APMM thus refers to utilisation of ACoA in Krebs cycle and the subsequent ATP production during oxidative phosphorylation (OXPHOS) from the NADH and FADH_2_ it produces.

The factors governing APMM are complex. For example, Kreb’s cycle can be regulated at multiple points to generate the various intermediates as required for specific metabolic needs other than ATP production. OXPHOS is dependent on the availability of oxygen to accept the electrons at the end of the electron transport chain, forming water by binding the hydrogen ions. For simplicity, we have assumed that external conditions such as the need for Kreb’s intermediates are constant, and oxygen is in plentiful supply. APMM is thus influenced by the availability of its substrates **adenosine diphosphate** (**ADP**, described by equation S1.5) and ACoA, multiplied by the intrinsic mitochondrial capacity (IMC) to convert ACoA to ATP, and the AMPK activity (K) which stimulates APMM as in equation S1.8.

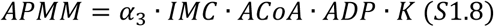

**Intrinsic mitochondrial capacity** (**IMC**) is modelled as a dynamic variable defined by the functionality of the mitochondrial space per unit volume multiplied by the amount of mitochondrial space (i.e. mass · mass-independent functionality). In the main article we mention the complex list of effectors influencing the IMC, but in the FEM we focus on K as the primary regulator of all subsequent factors influencing IMC, which therefore determines the rates of Kreb’s and OXPHOS. Thus, low ATP and high K accelerate Kreb’s and OXPHOS, and high ATP low K slow them^605,606^, and the factors affecting IMC can be condensed into a function of K so that changes in IMC can be defined as in equation S1.9.

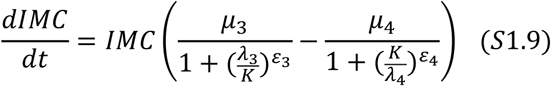

The equation is designed to have dynamic compensation with respect to K, forcing K to an equilibrium value determined by the parameters **λ_3_** and **λ_4_** as described by Karin, et al.^195^ (see Part 9.3 for further discussion). Thus, the ratio of ATP:AMP and total ATP level are maintained at equilibrium despite changes in metabolism (described further in model discussion).

##### 9.1.1.6 ACoA dynamics

***ACoA dynamics*** are defined by the processes in equation 2 (shown here to aid interpretation) and described in equation S2.

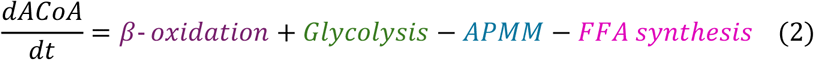

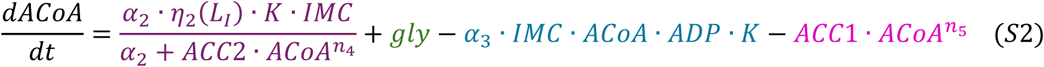

In addition to glycolysis discussed above, ACoA is produced by β-oxidation, and can either be de-acetylated in APMM or converted back into free fatty acids (FFAs) via FFA synthesis.

ACoA is controlled by robust negative feedback. Excess mitochondrial ACoA is converted to malonyl-CoA by acetyl-CoA carboxylase 2 (ACC2), which then allosterically inhibits the transfer of cytoplasmic acyl-CoA through the mitochondrial membrane into the matrix through carnitine palmitoyltransferase 1 (CPT1, forming acyl-carnitine as an intermediate)^607^. This is frequently the rate-limiting step in **β-oxidation**; however, it is tissue dependent^608^. Thus, the main inhibitory factor for β-oxidation was mitochondrial malonyl-CoA, which we define in equation S2.1 as a function of ACoA.

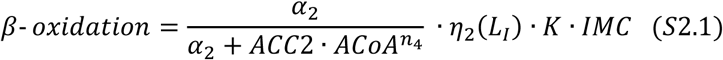

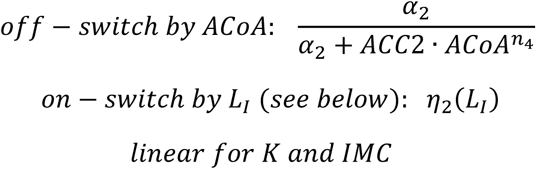

K influences the rate of β-oxidation indirectly by activating peroxisome proliferator activated receptor α (PPARα) and peroxisome proliferator-activated receptor-gamma coactivator 1 (PGC1)^609^, regulating IMC (equation S1.9), but also directly by inhibiting ACCs^610^, and stimulating malonyl CoA decarboxylase (MCD) which degrades malonyl-CoA^611^.

It is also dependent on the availability of L_I_. Just as we described the dependence of AHCM on ATP levels by the function η(ATP), equation S2.2 describes a similar function **η_2_(L_I_)** for the dependence of β-oxidation on L_I_. As most L_I_ is stored as lipid droplets (LDs), while FFAs are kept relatively constant, the rate of β-oxidation should only become dependent on L_I_ once LDs have been exhausted and can no longer replenish the FFAs required for APMM.

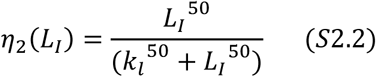

Excess mitochondrial ACoA is also converted to citrate (in Krebs cycle) which is then pumped out of mitochondria by tricarboxylate carriers and converted back to ACoA by citrate lyase.

Cytoplasmic ACoA then forms the substrate for **FFA synthesis**, converted to malonyl-CoA by acetyl-CoA carboxylase 1 (ACC1, the cytoplasmic version of ACC2). Malonyl-CoA is then converted to FFAs by fatty acid synthase (FAS). Thus, we defined FFA synthesis by equation S2.3, where ACC·ACoA is used as a proxy for malonyl-CoA as for β-oxidation, and constants for ACC1 and FAS are combined to form a single constant (ACC1).

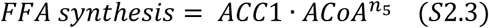

We have assumed FFA synthesis under simulated conditions is low, setting ACC1 to a value of 0.00001 (compared to a value of 2 for ACC2), reflecting that we are initially modelling lipid dynamics in the absence of alternative fuels which would be converted to fats, so FFA synthesis will be downregulated.

##### 9.1.1.7 Lipid dynamics

***Lipid dynamics*** have been simplified to consider only FFAs, triglycerides (three fatty acids bonded to glycerol), and the macromolecules and organelles which bind and carry them. FFAs in **meals** enter the body via the gut, are esterified to form triglycerides, and transferred to the blood attached to lipoproteins (chylomicrons from gut). Equally, FFAs from internal sources are esterified and bound to very-low-density lipoprotein (VLDL) in the liver, which is transported around the blood forming low-density lipoprotein (LDL). Unlike glucose, FFAs are not soluble in water, so only a tiny percentage (<0.01%) are carried free in the plasma with >95% bound to albumin^612^, while more than 90% of total fatty acids in plasma are stored as esters (such as triglycerides), which are mainly transported on lipoproteins. This combined pool of **blood lipids** is referred to as **L_B_**. Similarly, the combined pool of FFAs, triglycerides, and lipid droplets (LDs) are grouped together as **intracellular lipid** (**L_I_**).

###### 9.1.1.7.1 L_B_ dynamics

Changes in L_B_ are defined by the processes in equation 3 (shown here to aid interpretation) and described in equation S3. For reasons described below, both import and export refer exclusively to the adipose tissue (shorted from ‘adipose import’ and ‘adipose export’ for brevity).

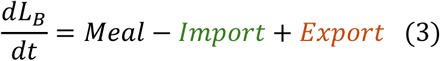

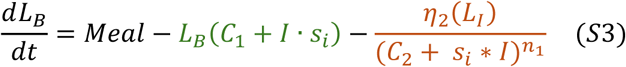

The **import** of L_B_ is controlled primarily by **insulin** (variable **I**) through the activation of cluster of differentiation 36 (CD36), in a similar mechanism to glucose transport via glucose transporter type 4 (GLUT4)^189,613,614^. Insulin also stimulates lipoprotein lipase in the wall of the vascular endothelium, which breaks the ester bonds of triglycerides to release the FFAs for import to adipose tissue^615^. The degree to which insulin stimulates the import of L_B_ is dependent firstly on L_B_ and secondly on the insulin sensitivity, s_i_ (variable described below) of the adipocytes. Thus, import is described as in equation S3.1, with insulin-independent import defined by the parameter **C_1_**.

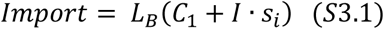

Due to the similarities between L_B_ and blood glucose regulation, we modelled import of L_B_ with a derivation of the blood glucose model produced by Topp, et al.^134^, called the β-cell, insulin, glucose (BIG) model. We adapted a derivation of the BIG model produced by Karin, et al.^195^ for import and insulin dynamics, substituting L_B_ for glucose as in equation S3.2, reflecting that insulin is produced by β-cells in the pancreas, which are stimulated by circulating FFAs both to produce insulin^616,617^ and proliferate^618–620^ similarly to blood glucose. Thus, as in equation S3.2, insulin is produced in an L_B_-dependent manner at rate **q** per β-cell, multiplied by the number of β-cells, or **β-cell mass** (variable **βM**). Insulin then degrades spontaneously at rate **γ_i_**.

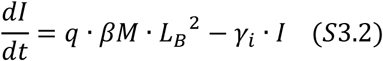

This model was appealing because Karin, et al.^195^ demonstrated that the BIG model had dynamic compensation, meaning that for any value of q or s_i_ the system would adapt to return L_B_ to equilibrium (see model discussion for further detail). This occurs because of the βM expanding and contracting according to the level of L_B_ as in equations S3.3-3.4, based on those used by Karin, et al.^195^. In the FEM, L_B_ stimulates proliferation and inhibits apoptosis so that βM grows and shrinks according to insulin production requirements. It forces L_B_ to an equilibrium value determined by the parameters **λ_1_** and **λ_2_**.

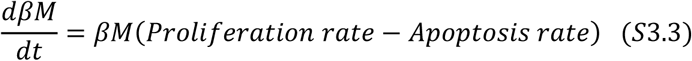

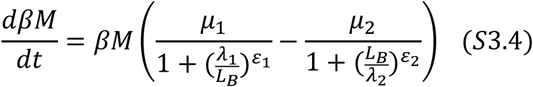

The BIG model did not consider L_I_ **export**, but evidence suggests this is also dependent on insulin. Insulin inhibits lipolysis and export, inhibiting lipolysis in an AKT-independent^621^ and dependent mechanism by dephosphorylating hormone sensitive lipase and preventing de-esterification of intracellular triglycerides in LDs^622,623^. We modelled export with an insulin-dependent and independent component, **C_2_**, as described in equation S3.5.

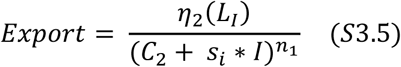

The η_2_(L_I_) function limiting β-oxidation when L_I_ drops below the threshold will also apply to export, as the central function of export (from the adipose) is to replenish L_B_ and ensure the supply of FFAs for *non-adipose* β-oxidation (as well as broader metabolic needs not considered here). Increasing export simply because there is more L_I_ stored would therefore be counterproductive. Notably, as most non-adipose tissues cannot store L_I_ as LDs, excess L_I_ is toxic^188,624,625^, necessitating swift return to equilibrium. The η_2_(L_I_) function may thus not apply to non-adipose export, while import likely adapts much more swiftly with alterations to s_i_. As the functions for non-adipose export and import likely differ from the functions for adipose tissue, L_B_ would more accurately be described by equation S3.6.

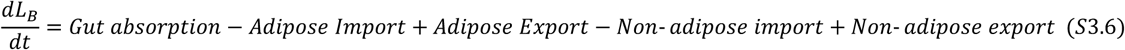

As there is little storage of L_I_ in non-adipose tissues, the level of L_I_ import will roughly equal the level of β-oxidation over time, either because excess L_I_ is exported back into the blood or the insulin sensitivity of the tissues quickly adapts to reduce import to match their metabolic rate. Therefore, net import ≈ β-oxidation. Thus, equation 3.6 could be re-written as in equation S3.7, and **Meal** defined as in equation S3.8, where non-adipose β-oxidation is defined as in equation S3.9.

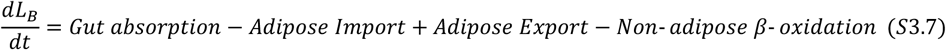

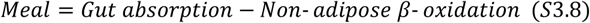

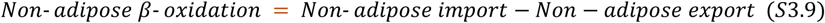

The implications of this simplification are described in the model discussion, but it allowed us to limit the FEM to the adipose and blood rather than including additional tissues. NB: we used the term ‘Meal’ in the main text as this was sufficiently descriptive for the textual analysis of metabolic slowdown before considering that the FEM separates adipose from non-adipose dynamics.

Thus, when we refer to the **insulin sensitivity** (variable **s_i_**) we refer exclusively to the adipose. Although an in-depth review of s_i_ regulation is beyond the scope of the FEM, it is worth noting that because adipose can store L_I_ (as LDs) it cannot regulate s_i_ by the size of the FFA pool as can other tissues. However, at the point of hypertrophy continuing import would lead to bursting and necrosis (uncontrolled cell death). Although this does happen, it appears to reflect mainly stress conditions rather than normal adipose homeostasis^210,626^. Thus, to avoid bursting, adipocytes must regulate s_i_ with respect to their L_I_ as stored in LDs. It is well established that small adipocytes are more insulin sensitive than large ones^627,628^, and adipocyte size correlates with s_i_ independent of fat mass^629^.

One mechanism reducing s_i_ results from the redistribution of cholesterol from the plasma membrane to the LDs^630,631^. Cholesterol forms a major part of lipid rafts called caveolae which act as major docking sites for insulin sensitive transporters such as CD36 and GLUT4^632,633^.

Therefore, cholesterol depletion of the membrane prevents endocytosis and insulin signalling in adipocytes^634^, promoting insulin resistance (IR). Importantly, the main protein stabilising caveolae, caveolin 1 (CAV1), regulates expansion of LDs by its transfer from the plasma membrane to the membrane of the organelles^635,636^, thus as LDs expand, adipocytes become less insulin sensitive. Therefore, s_i_ is reduced by increasing L_I_ as defined by equation S3.10, while production of insulin receptor was modelled to occur at a constant rate defined by parameter, **k_s_**.

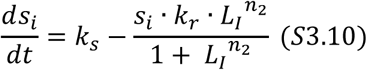

**Plaque formation** and growth was modelled as a function of elevated L_B_. We suggested there would be an **equilibrium point** where the rate of plaque degradation equalled the rate of plaque production, **P_eq_**, and risk from plaques would change as shown in equation S3.11 (NB: risk was reset to 1 at time t = 1900 (after model has reached equilibrium), and could not drop below zero (see supplementary code, lines 61-72).

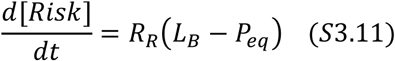

When L_B_ rises above P_eq_, (*L*_*B*_ − *P*_*eq*_ > 0) risk increases as the plaques enlarge and spread (linearly as the distance from equilibrium increases), and when L_B_ drops back below P_eq_ the plaques decrease in size and number as the removal mechanisms begin to work faster than lipid deposition. Notably, P_eq_ is not (necessarily) the same as the equilibrium value for L_B_, and does not have to be constant. We could incorporate ‘wear and tear’ induced endothelial dysfunction as a steadily declining P_eq_, or P_eq_ could be a function of inflammation and/or insulin resistance; see Theofilis, et al.^637^ for recent review.

###### 9.1.1.7.2 L_I_ dynamics

Changes in L_I_ are defined by the processes in equation 4 (shown here to aid interpretation) and described in equation S4.

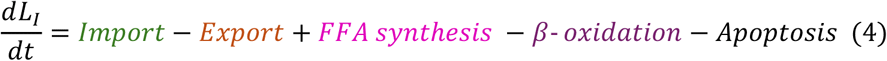

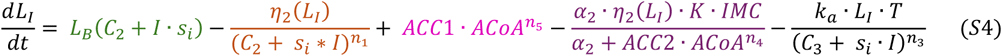

The processes for Import, Export, FFA synthesis and beta-oxidation have all been described above.

**Apoptosis** removes hypertrophic cells via activation of a localised immune response and inflammation. Hypertrophic cells produce inflammatory cytokines which attract immune cells such as macrophages^211^ and cause a phenotypic switch toward the inflammatory M1 polarisation^212^.

Basal inflammation in the adipose tissue is a highly complex process, including an array of activatory and inhibitory cytokines and chemokines, as well as T and B cells in addition to macrophages and monocytes, some of which can also function to inhibit inflammation as well switch between pro and anti-inflammatory states. For the FEM, we modelled inflammation by simplifying it to a single variable, **TNF-α** (**T**), which is thought to be responsible for the extrinsic apoptosis of hypertrophic adipocytes^210,638^ (see Part 9.3 for implications). Hence knockout mouse models with increased inflammation and apoptosis have reduced fat mass^639^.

Infiltrating macrophages localize around large or dead adipocytes forming crown-like structures^215^, suggesting that hypertrophic adipocytes are preferentially targeted for apoptosis, perhaps in part because they are insulin resistant^638,640,641^. Also in obese children, the size of fat cells correlated more strongly with inflammatory markers than either body mass index (BMI) or fat mass^629^. Therefore, we modelled apoptosis as dependent on L_I_, representing the number of hypertrophic cells, and degree of inflammation, T, as in equation S4.1.

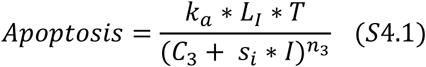

Hypertrophic cells have increased expression of Toll-like receptor 4 (TLR4) and NF-κB expression which not only attract macrophages^642^, but sensitize cells to apoptosis^643,644^, and contribute to IR^645,646^. Importantly, death-receptor-induced apoptosis is inhibited by IGF-1 and insulin in white^638,647^ and brown^648^ adipocytes. Insulin promotes survival in multiple cell types^640,641^, and increasing insulin concentrations in vitro protect adipocytes from TNF-α induced apoptosis^638^. Thus, while both hypertrophic adipocytes and infiltrating macrophages cause adipose inflammation, mainly the insulin resistant hypertrophic adipocytes undergo apoptosis^627–629^. Thus, apoptosis was inhibited by insulin dependent on the degree of s_i_ (and independently of insulin, **C_3_**).

We modelled variable **T** in equation S4.2 to be increased primarily by the number of hypertrophic cells, represented by L_I_, which produced inflammatory T. The rate of T degradation was modelled as spontaneous, similarly to insulin degradation, dependent on the amount of T multiplied by degradation constant, **γ_t_**. Adipocyte-induced inflammation follows a sigmoid switch function reflecting that basal inflammation is a dangerous tool to allow to steadily increase ad infinitum. Beyond a certain point it could initiate a full-blown immune response, consistent with the link between obesity and autoimmunity^649^.

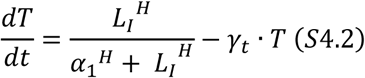

A summary of how the equations were derived is shown in Table 1.

**Table 1.** Rationale for selected equations. Abbreviations, see Table 6.

| Equation | Explanation of key dynamics |
| --- | --- |
| Main equations |  |
| APMM | $\alpha_3 \cdot IMC \cdot ACoA \cdot ADP \cdot K$<br>For simplicity, model assumed linear dynamics based on substrates (ACoA and ADP) dependent firstly on the intrinsic mitochondrial capacity (IMC) to convert these to ATP and secondly the degree to which mitochondria were activated by AMPK (K). |
| AHCM | $\frac{E_{CM} \cdot k_{CM} \cdot \eta(ATP)}{K}$ |
| | Substrate ATP was presumed to be in excess (see $\eta$ equations below). Thus, AHCM was dependent on the basal rate, $k_{CM}$ multiplied by $E_{CM}$, the induced upregulation based on metabolic needs (eg. exercise in muscle). This was inhibited by AMPK. |
| Glycolysis | Model ignored glucose dynamics, so glycolysis was set to zero. |
| β-oxidation | $\frac{\alpha_2 \cdot \eta_2(L_I) \cdot K \cdot IMC}{\alpha_2 + ACC2 \cdot ACoA^{n_4}}$<br>$L_I$ in the adipose is presumed to be in constant excess (see $\eta$ equations below), removing substrate dependence. As β-oxidation is mitochondrial, it was dependent on the IMC and activated by AMPK. The process is inhibited potently by malonyl-CoA, which is produced from ACoA by the enzyme ACC2, hence we used $ACC2 \cdot ACoA^n$. Exponent reflects the potency of inhibition. |
| FFA synthesis | $ACC1 \cdot ACoA^{n_5}$<br>The process is dependent on its substrate ACoA, and the activity of the rate limiting enzyme ACC1. As in β-oxidation, we used $ACoA^n$ to indicate the potent negative feedback of ACoA on its own concentration. |
| Import | $L_B(C_1 + I \cdot s_i)$<br>We replicated the import equation from Karin et al. (2016), replacing glucose with $L_B$. |
| Export | $\frac{\eta_2(L_I)}{(C_2 + s_i \cdot I)^{n_1}}$<br>For export, we used the inverse of the import equation squared to reflect inhibition by insulin. While import was dependent on $L_B$, we considered export would not be dependent on $L_I$ due both to the inefficiency this would produce as a storage mechanism (see text) and that $L_I$ is mainly locked away in LDs while FFAs in the adipose remain constant. Hence, we used a $\eta(L_I)$ for substrate dependence. |
| Apoptosis | $\frac{k_a \cdot L_I \cdot T}{(C_3 + s_i \cdot I)^{n_3}}$<br>The number of hypertrophic cells was modelled as a linear function of $L_I$, and the degree to which this would induce apoptosis depended linearly on the level of inflammation. We used the inverse of the import equation to reflect inhibition by insulin. |
| β-cell mass | $\beta M \left( \frac{\mu_1}{1 + (\frac{\lambda_1}{L_B})^{\varepsilon_1}} - \frac{\mu_2}{1 + (\frac{L_B}{\lambda_2})^{\varepsilon_2}} \right)$<br>We replicated the insulin equation from Karin et al. (2016), replacing glucose with $L_B$. The sigmoid curve produces dynamic compensation, maintaining $L_B$ at homeostasis independent of insulin production and sensitivity (see discussion). |
| Insulin | $q \cdot \beta M \cdot L_B^2 - \gamma_i \cdot I$<br>We replicated the insulin equation from Karin et al. (2016), replacing glucose with $L_B$. The equation reflects basic dynamics of protein production and degradation. |
| AMPK activity | $k_{ka} \frac{AMP}{ATP} - k_{kd} \cdot K$<br>We used the same dynamic equation as for insulin reflecting AMPK production and degradation. AMPK production/activation is dependent on the ration of ATP:AMP. For simplicity, ADP was ignored, and additional factors such as those regulating protein phosphorylation were not included. |
| IMC | $IMC \left( \frac{\mu_3}{1 + \left(\frac{\lambda_3}{K}\right)^{\varepsilon_3}} - \frac{\mu_4}{1 + \left(\frac{K}{\lambda_4}\right)^{\varepsilon_4}} \right)$<br>Because IMC was dependent on mitochondrial mass, we replicated the ODE for β-cell mass replacing $L_B$ with AMPK activity, the primary activator of IMC. However, the goal of using a similar equation was to generate the dynamic compensation for ATP using the same sigmoid curve. |
| $s_i$ | $k_s - \frac{s_i \cdot k_r \cdot L_I^{n_2}}{1 + L_I^{n_2}}$<br>We assumed insulin receptor was produced at a constant rate, $k_s$. Receptor degradation was dependent linearly on the amount of insulin receptor ($s_i$), and $L_I$ in a bounded relationship designed to restrain fluctuations in $s_i$. Exponent was small to produce a smooth change in $s_i$. |
| Inflammation | $\frac{L_I^H}{\alpha_1^H + L_I^H} - \gamma_t \cdot T$<br>As inflammation is a highly complex process, this equation was dramatically simplified to reflect a single inflammatory protein, such as TNF-α. Its degradation dynamics followed insulin and AMPK. Activation was a gradual switch function dependent on $L_I$ that would limit excessive inflammation, preventing an infection response. |
| Other equations |  |
| ATP, ADP, AMP | $ATP = ATP_{max} - 2 \cdot AMP$ $ADP = AMP$<br>We assumed a constant pool of adenosine phosphates ($ATP_{max}$). ATP was therefore equal to $ATP_{max} - (ADP + AMP)$. For simplicity, we assumed ADP and AMP were equal. |
| $\eta_1$ and $\eta_2$ | $\eta(ATP) = \frac{ATP^{10}}{(k_p^{10} + ATP^{10})} \quad , \quad \eta_2(L_I) = \frac{L_I^{50}}{(k_l^{50} + L_I^{50})}$<br>These functions created a switch function to produce substrate independence above a low level of substrate for ATP and $L_I$. |
| $k_{CM}$ | $- \frac{k_{CM}}{4000}$<br>After the model equilibrated, we introduced a gradual decrease in $k_{CM}$ independent of any other variables to reflect metabolic slowdown as a result of selective destruction. |

#### 9.1.2 Model running and storage

The FEM was constructed using R version 3.6.3 with the deSolve package for ordinary differential equations (ODEs). The R code for the model and all data analysis are available on FAIRDOM Hub with the following identifying links: <u>DOI: 10.15490/fairdomhub.1.model.851.1</u> SEEK ID: https://fairdomhub.org/models/851?version=1

Contrary to most biological models, the purpose of the FEM was not to bring several molecular and cellular components of interest together and elucidate the resulting system behaviour. The system behaviour was predicted by the literature evidence, and the model was designed to demonstrate this mathematically. Although further research is required to fit the model with real concentrations and masses of the included components, here we were interested only in broader homeostatic implications of metabolic slowdown as indicated by whether metabolites and components remained at constant rates and concentrations (homeostasis) and if not, the direction of the shift over time.

Values and definitions for static parameters are found in Table 2, and initial variable values and adjusted parameters in Table 3. Rough explanations for the chosen values are provided, but were mainly arbitrary, as were the units (au). We have defined the nature of the units in Table 3; however, as stated, the FEM is supposed to represent broad biological contexts rather than any specific biological conditions. Some values were changed during simulations for aesthetic purposes (ie to speed or slow certain reactions), but none affected the homeostatic implications of the FEM.

**Table 2.**
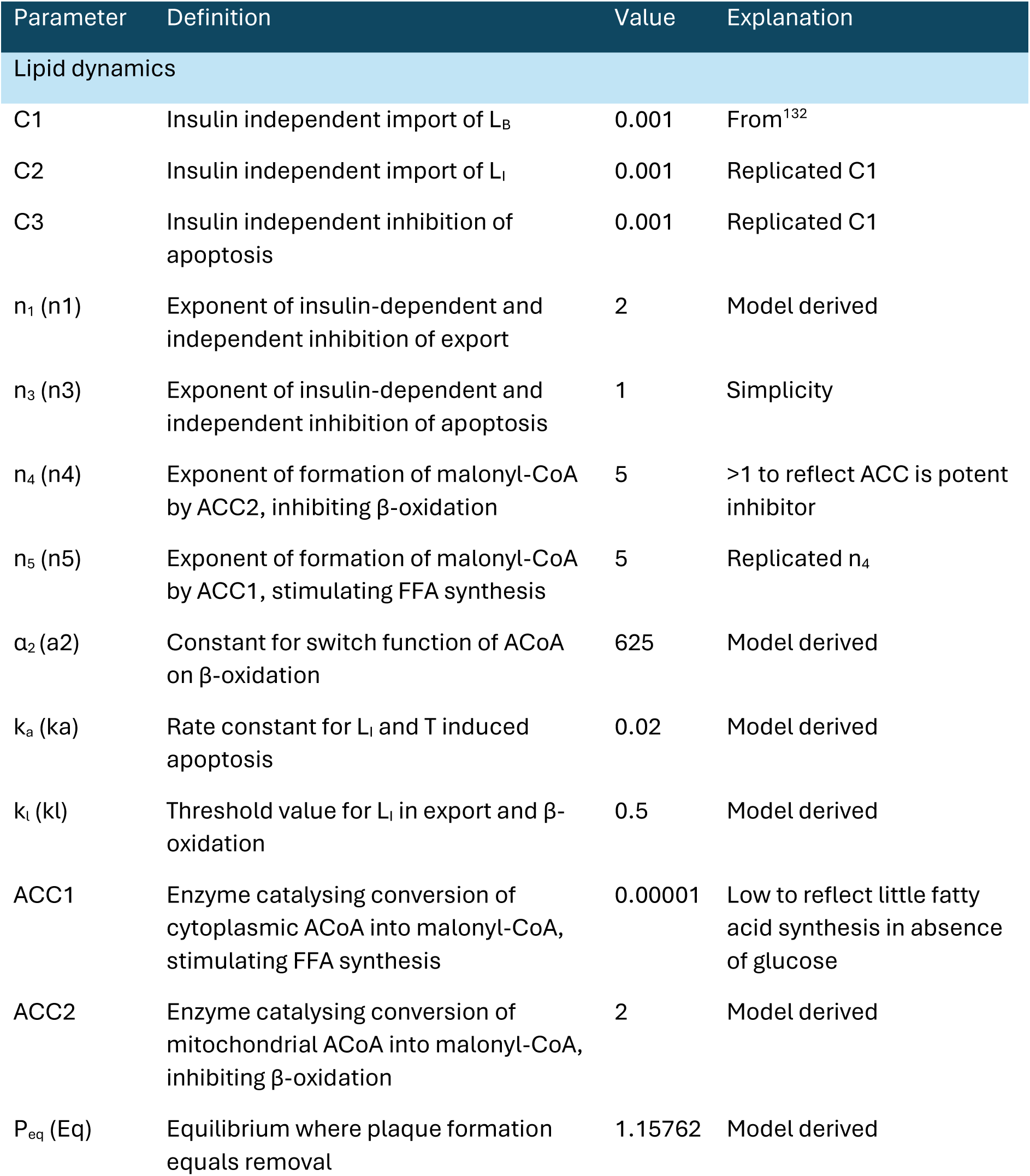
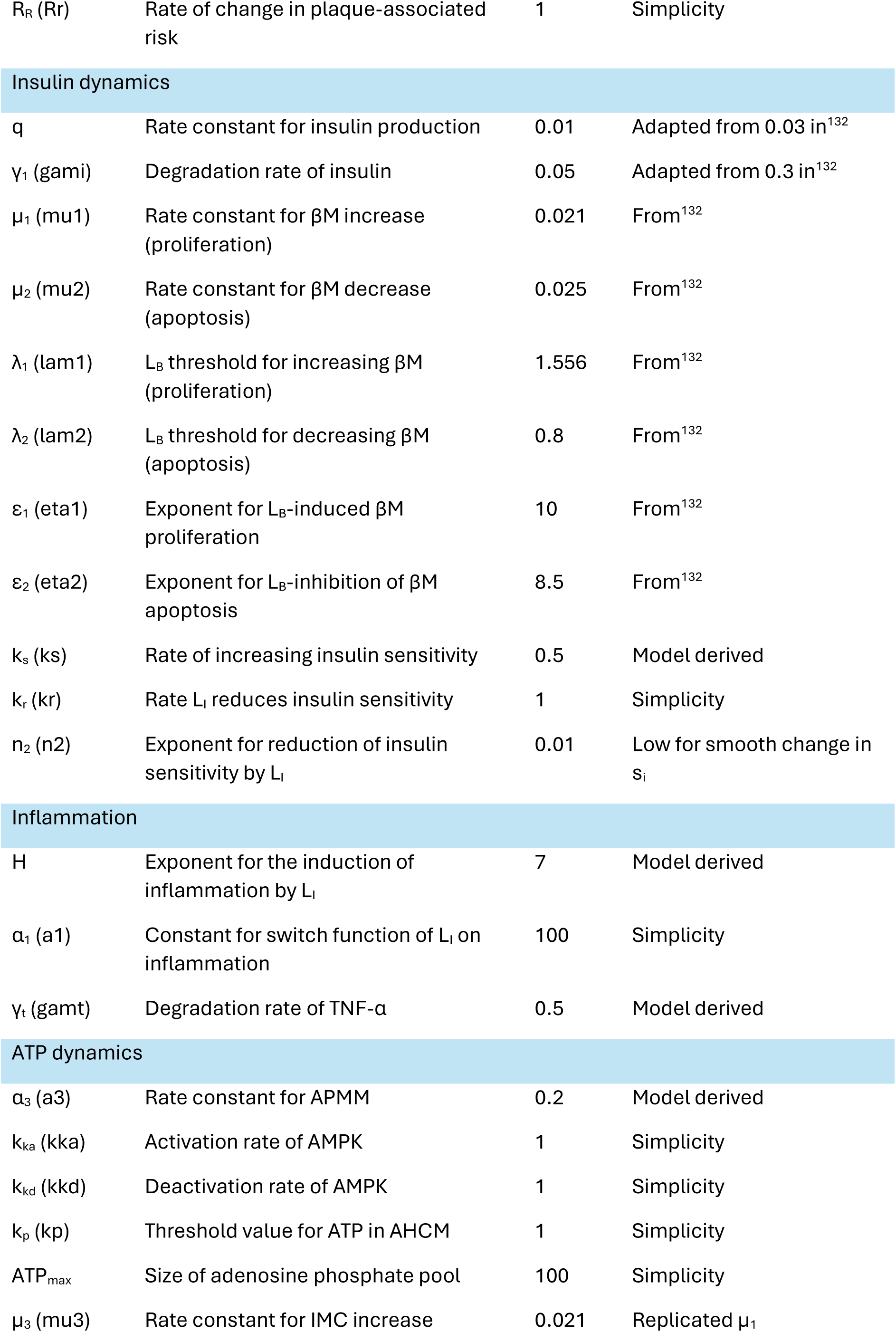
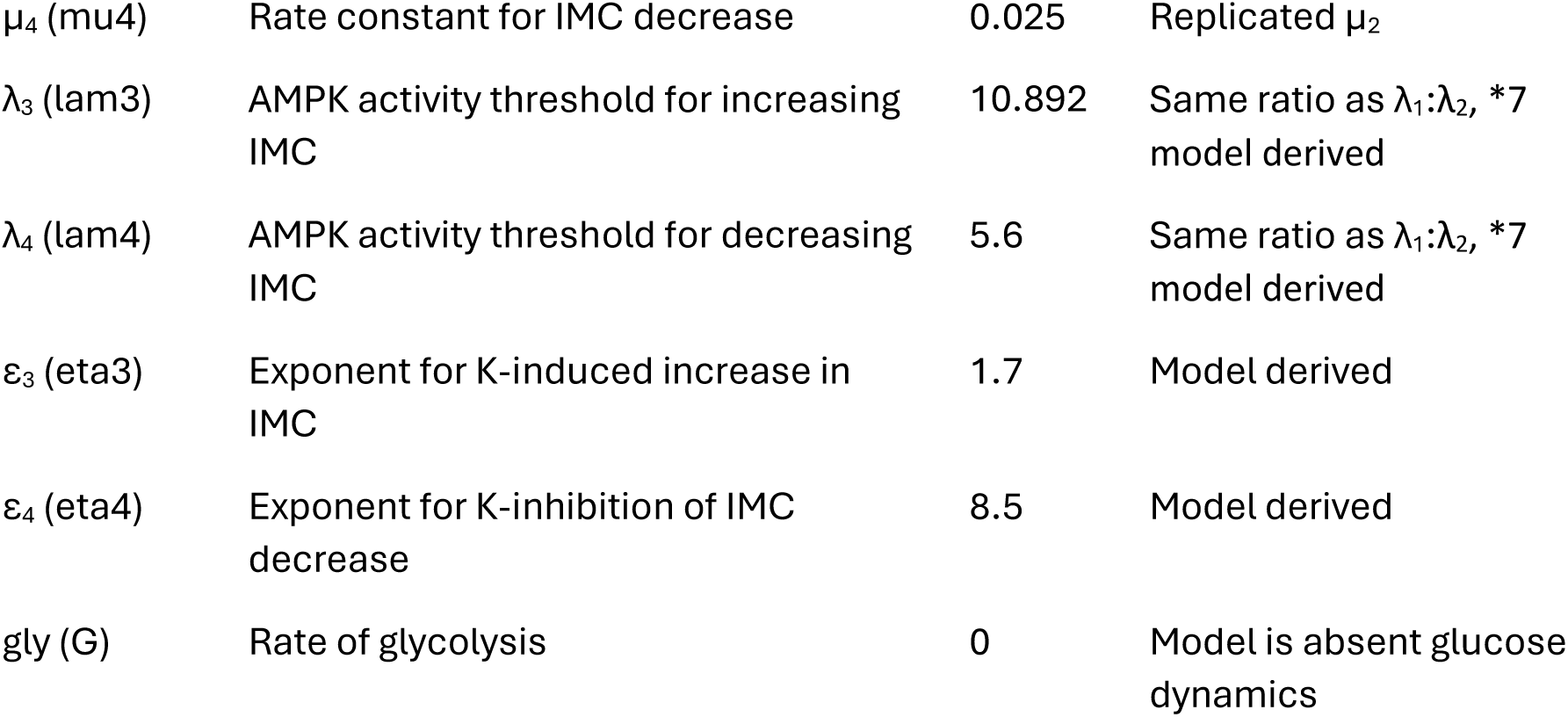
Parameter values for the FEM. Terms in brackets are the parameter names used in the code. For explanations, ‘Model derived’ reflects adoption or adaptation of values based on preceding model runs. Adaptations were for aesthetic purposes and did not alter model behaviour; ‘Simplicity’ reflects retention of starting values selected as multiples of 10 based on crude assessments of relative rate or concentration. Abbreviations, see Table 6.

**Table 3.** Variables and parameters which change during model simulations. Symbol refers to the names used in the code. All units are arbitrary (au). Abbreviations, see Table 6.

| Variable | Symbol | Full name | Initial value | Definition of unit |
| --- | --- | --- | --- | --- |
| ATP | A | ATP | 16.66667 | Concentration per adipocyte (assumed uniformity across tissue) |
| ACoA | Aco | Acetyl-CoA | 5 | Concentration per adipocyte (assumed uniformity across tissue) |
| $\beta M$ | BM | B-cell mass | 1 | Total mass as proxy for cell number |
| I | I | Insulin | 2.3955 | Concentration of insulin in the blood |
| IMC | Ma | Intrinsic mitochondrial capacity | 0.04 | Calculation of mitochondrial mass and efficiency of mitochondrial membrane per cell (see text, assumed tissue uniformity) |
| $L_B$ | LB | Blood lipid | 5 | Combined blood concentration of fatty acids in FFAs, free triglycerides, and in triglycerides bound to lipid carriers |
| $L_i$ | LI | Intracellular lipid | 100 | Average per adipocyte concentration of fatty acids in FFAs, triglycerides, and LDs (will not be uniform) |
| K | K | AMPK activity | 5 | Calculation of AMPK concentration per cell and average activity determined by regulatory protein modifications |
| $s_i$ | si | Insulin sensitivity | 0.9656780 | Sensitivity of average adipocyte reflecting concentration of receptors per length membrane and average receptor activity determined by regulatory protein modifications |
| T | TN | TNF-α | 0 | Concentration of inflammatory cytokines across adipose tissue |

| Parameter | Symbol | Definition | Initial Value and changes |
| --- | --- | --- | --- |
| E<sub>CM</sub> | ECM | Elevation of AHCM | 1 → 1.5 simulates exercise |
| k<sub>CM</sub> | MR | Basal metabolic rate | 1, see equation S6 simulating metabolic slowdown. |
| Meal | m | Meal | 1.2 → 1.201 simulates CR to AL feeding |

##### 9.1.2.1 Parameter estimation

Parameter estimation involved changing each parameter ±1,2,5, and 10% and plotting the effects of changes over time (data not shown). Then to compare effects of each parameter on the variables and processes, we firstly filtered the data to time ≥2000 au (removing the initial equilibration effects) and then used the data for ±10% change in each parameter. We calculated the difference between 10% increase and decrease for each timepoint and took the mean difference across all timepoints for each variable and process, termed X. Then we normalised X values by dividing by the absolute maximum X value for each variable (i.e. the value for the parameter with greatest impact on the variable), as shown in equation S5.

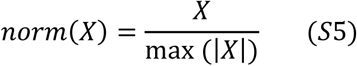

##### 9.1.2.2 Model inputs (ageing and therapies)

Metabolic slowdown was modelled as a continuous decline in k_CM_ over time (from t=2000 au), as defined by equation S6.

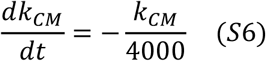

In reality both adipose mass (i.e. L_I_ storage capacity) and βM would be finite, limited by organ size. Notably, the same is true for excreting excess L_B_ via the liver in the bile (reverse cholesterol transport, RCT). To represent all these finite capacities, we added a maximal βM to the FEM (as it already included βM and not liver or adipose mass) as in equation S7.

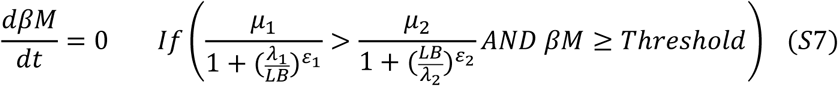

Once βM reached the threshold (7.849) it could no longer increase, preventing the necessary increase in insulin production to sustain L_B_ as AHCM continued to decline.

To model the impact of metformin we altered several parameters, firstly, k_ka_ and k_kd_ which determine the rates of AMPK activation and deactivation respectively. We simulated two doses of metformin, introduced between t = 2500 and t = 5000 au and then between t = 7500 and t = 10000 au, either by increasing k_ka_*1.5 or reducing k_kd_/1.5 to represent increased rate of activation and decreased rate of deactivation, respectively (see Part 9.2.4.2).

We modelled rapamycin as discussed in the main article, as both an initial drop (0.15· k_CM_) in k_CM_, but also a reduction in the rate of k_CM_ decline (0.25 times).

Exercise and calorie restriction were modelled as shown in Table 2.

### 9.2 Results

Initial simulations of the FEM for 2000 arbitrary units of time (au) ensured all variables and processes reached equilibrium before this time. We used the parameter values in Table 1 and starting variable values (and E_CM_ = 1, k_CM_ = 1, and Meal = 1.2) from Table 3.

#### 9.2.1 Model behaviour

##### 9.2.1.1 Parameter estimation at equilibrium

To test whether the model was performing appropriately, we ran a parameter estimation as described in the model running and storage, using a 10% change in parameter value to observe effects on each variable and process. Figure 16 shows the relative impact of changing parameters (±10%) for each variable or process: all values are shown as a fraction of the parameter with the greatest impact (by decreasingly bright colour). Red indicates that increasing the parameter has a positive effect on the variable/process and blue a negative effect.

**Figure 16.**
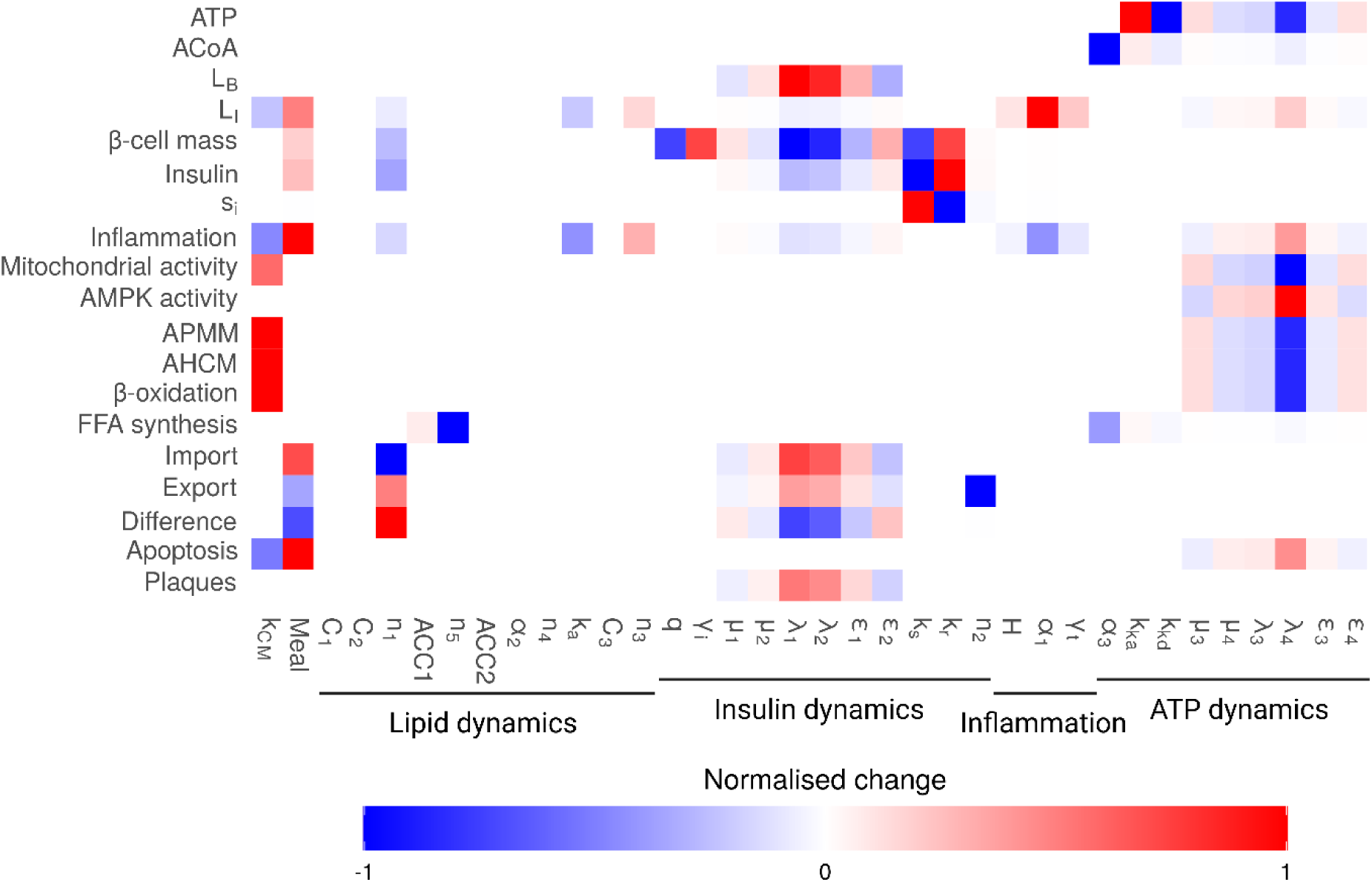
Parameter estimation for FEM post equilibration. Values were normalised as described in the model running and storage section so that each value is relative to other parameters for each individual variable/process. An absolute value of 1 (or −1) represents the parameter that has the greatest effect on that variable or process. Positive values reflect that increases in the parameter increase the variable or process, while negative values reflect that increases in the parameter decrease the variable or process. Abbreviations, see Table 6.

The parameters for ATP dynamics affected most variables and processes, except those which were instead mainly affected by insulin dynamics. Both ATP dynamics and insulin dynamics influenced inflammation (TNF-α) and the parameters for inflammation mainly impacted L_I_.

Six lipid dynamics parameters affected no variables or processes (at the level of 10% change). These included C_1_, C_2_, and C_3_, which reflected the insulin independent components of import, export (inhibition), and apoptosis (inhibition), respectively. This is an interesting observation as these processes can be insulin-independent^650,651^, and the C_1_ parameter for insulin independent import was adopted from the BIG model equation for glucose dynamics used by Karin, et al.^195^. Additionally, the parameters α_2_, ACC2, and n_4_ also had no effect on any variables or processes. All these parameters are part of the β-oxidation equation (S2.1), but have no impact on the FEM because they do not affect the rate of β-oxidation, which is instead determined by the rate of AHCM through k_CM_ and the parameters which affect the equilibrium of ATP (μ_3_, μ_4_, λ_2_, λ_4_, ε_3_, and ε_4_), as shown in Figure 16. We predict as the FEM is expanded with additional variables and processes, such as the inclusion of glucose as an alternative fuel, these parameters may gain relevance.

##### 9.2.1.2 Phase portrait analysis

The model was built to maintain homeostasis of key metabolites (ATP, ACoA, L_B_, and L_I_). To demonstrate this, we first broke the model down into parts which allowed examination of key variables in two-dimensional phase planes.

Firstly, we looked at ATP, which is governed as in equation S1, copied below.

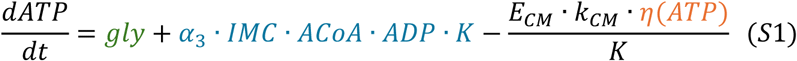

From the parameter estimation in Figure 16, we can see that ATP is not impacted by shifts in k_CM_ while ATP hydrolysis (AHCM, black) is strongly affected. As glycolysis (gly) is zero, changes in AHCM must be balanced by changes in ATP synthesis (APMM, yellow). Within APMM, ADP is a function of ATP and thus not affected; ACoA and AMPK activity are also unaffected by shifts in k_CM_ (Figure 16), leaving IMC as the variable which must shift to maintain ATP equilibrium by balancing changes in AHCM with complimentary shifts in APMM. Thus, fluctuations in ATP are returned to equilibrium by changes in IMC.

To assess the impact of IMC on ATP, we used a subsection of the model containing only the ODEs for ATP and IMC. ACoA was modelled as a constant and AMPK activity modelled directly as a ratio of AMP:ATP. For AHCM, whether we used η(ATP), orange in equation S1, representing substrate independence, or substituted in ATP so the reaction became substrate dependent made no difference to the following analysis (not shown). We plotted the phase portrait including nullclines, indicating a single stable fixed point as shown in Figure 17. ATP adapts quickly toward equilibrium, driven by changes in mitochondrial capacity.

**Figure 17.**
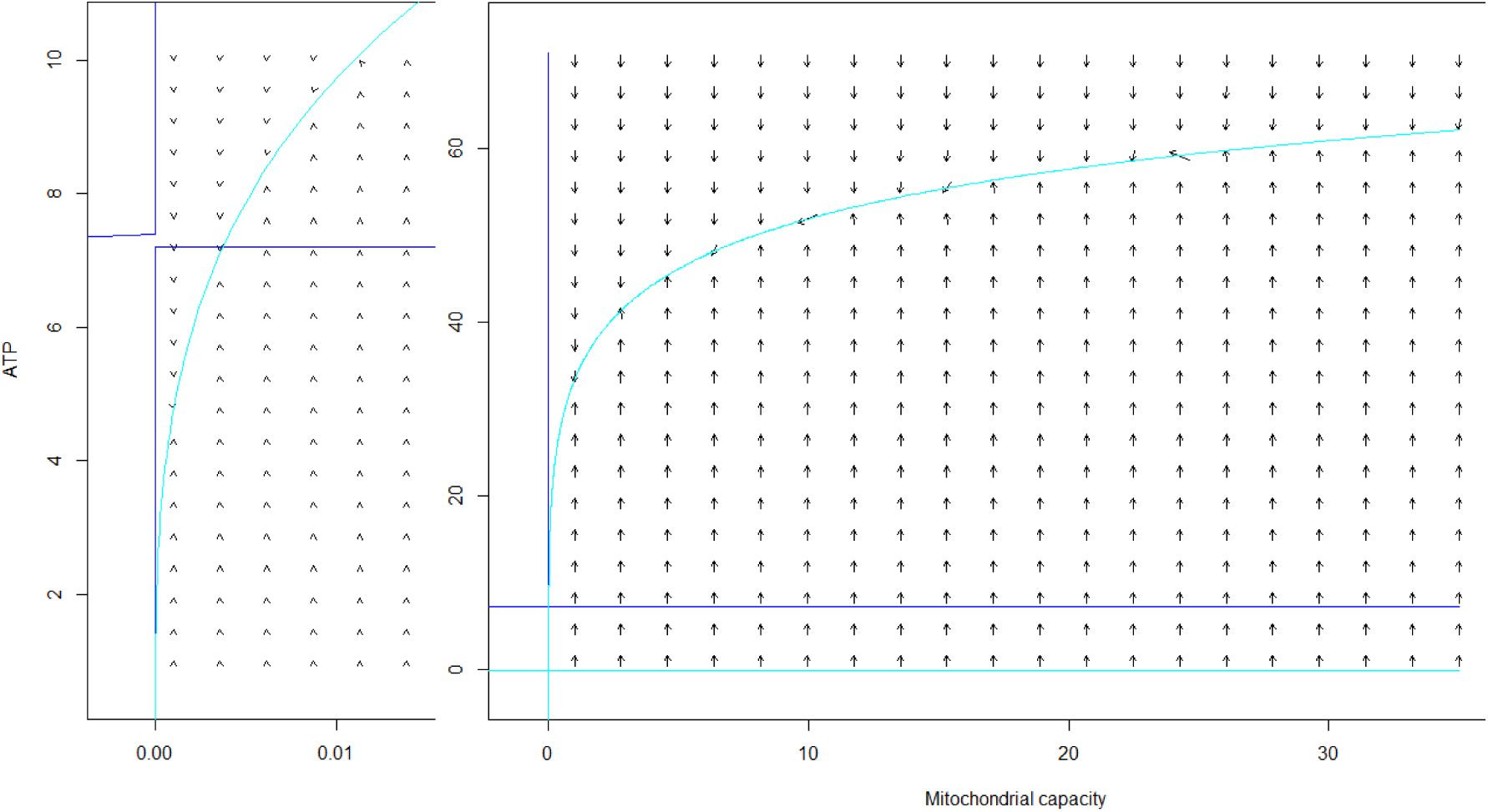
Phase plane analysis of relationship between ATP and IMC from simplified FEM at low (left) and (high) values of IMC (model starting value 0.04) and ATP (model starting value 16.67). x nullclines in blue, y nullclines cyan.

Next, we looked at ACoA which is governed as in equation S2, copied below.

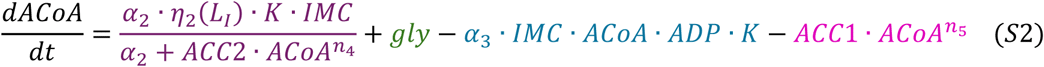

Changes in k_CM_ cause compensatory shifts in IMC, which could therefore impact ACoA. However, IMC affects both β-oxidation which increases ACoA, and APMM which reduces it, converting it to ATP. Although ACoA has strict negative feedback, inhibiting its own formation and inducing removal by fatty acid synthesis, Figure 16 suggests that the only parameters affecting ACoA homeostasis are ATP dynamics. Although the parameters in equation S1.9 for IMC have a minor effect, the primary parameters are α_3_, impacting the rate of APMM, and k_ka_ and k_kd_ which alter the rate of AMPK activation and deactivation respectively. All of these parameters regulate the rate at which ADP is converted to ATP.

Reducing the model to the ODEs for ATP and ACoA, with K modelled again as a straight ratio of ATP:AMP, we produced a phase plane for ATP and ACoA, as shown in Figure 18. Changes in both metabolites resulted in return to a stable fixed point, despite modelling IMC as a constant.

**Figure 18.**
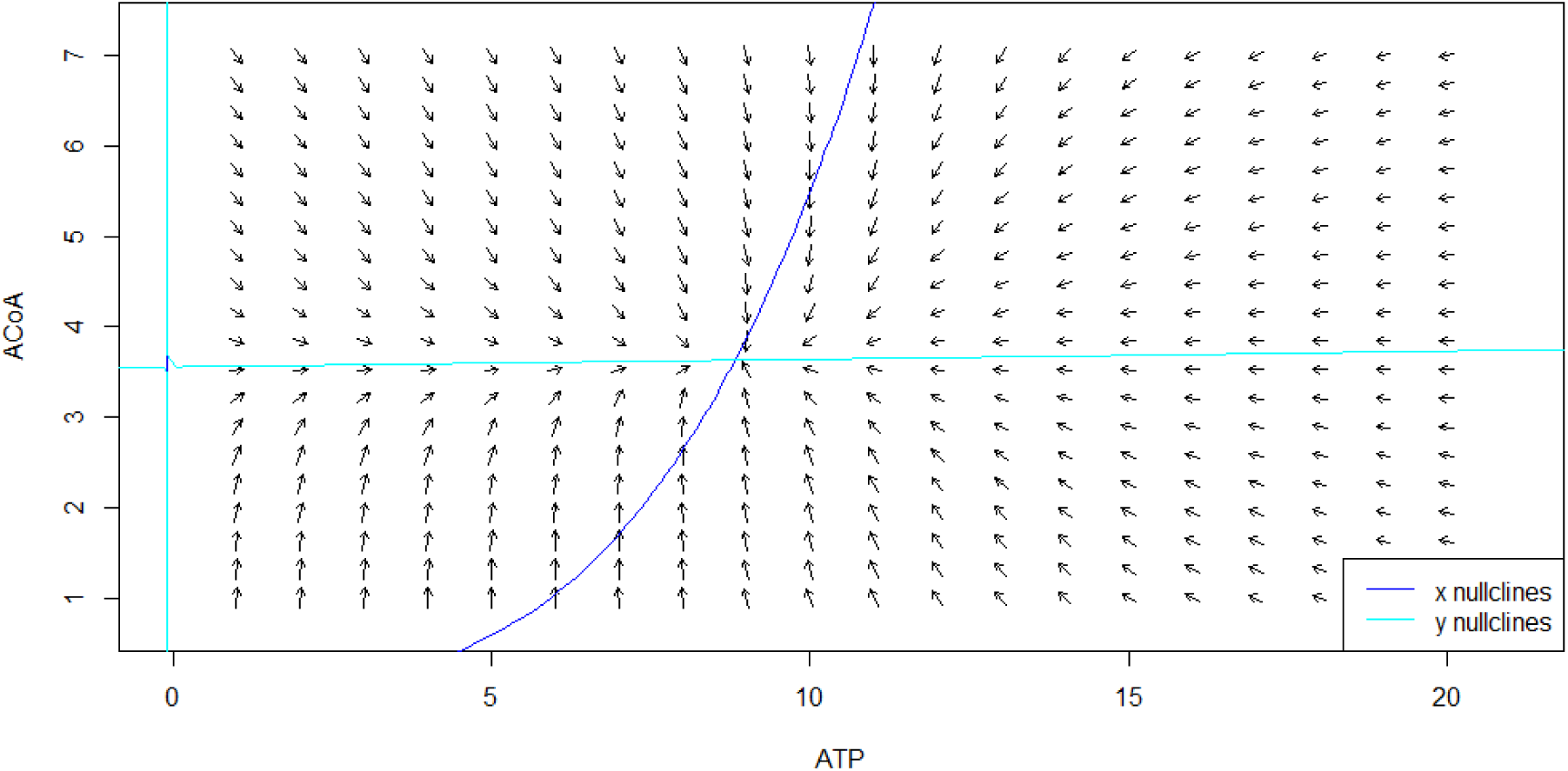
Phase plane analysis of relationship between ATP and ACoA from simplified FEM.

The FEM produces highly stable energy dynamics, where IMC maintains ATP equilibrium, which maintains ACoA equilibrium. The other half of the model concerns fat dynamics: the production of ACoA from FFAs and the conversion of ACoA to FFAs, which can then be transferred between the adipose tissue and blood.

The stability of the L_B_ has already been evidenced by Karin, et al.^195^, whose equations for glucose dynamics we replicated for L_B_, copied here in equation S3.

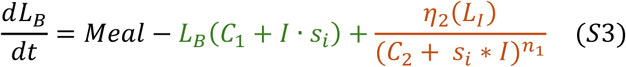

They demonstrated that blood glucose would return to swift equilibrium regardless of meal size, the rate of insulin production, or insulin sensitivity due to increases in insulin production from changes in in β-cell mass. This is consistent with our parameter estimation where L_B_ is impacted exclusively by the parameters affecting changes in β-cell mass (Figure 16). However, Karin, et al.^195^ did not include export in their equations. Phase portrait analysis of a simplified model including only equations S3 for L_B_ and S3.2 for insulin, where L_B_ and insulin were the only variables showed a stable fixed point with and without the export function (Figure 19A-B).

**Figure 19.**
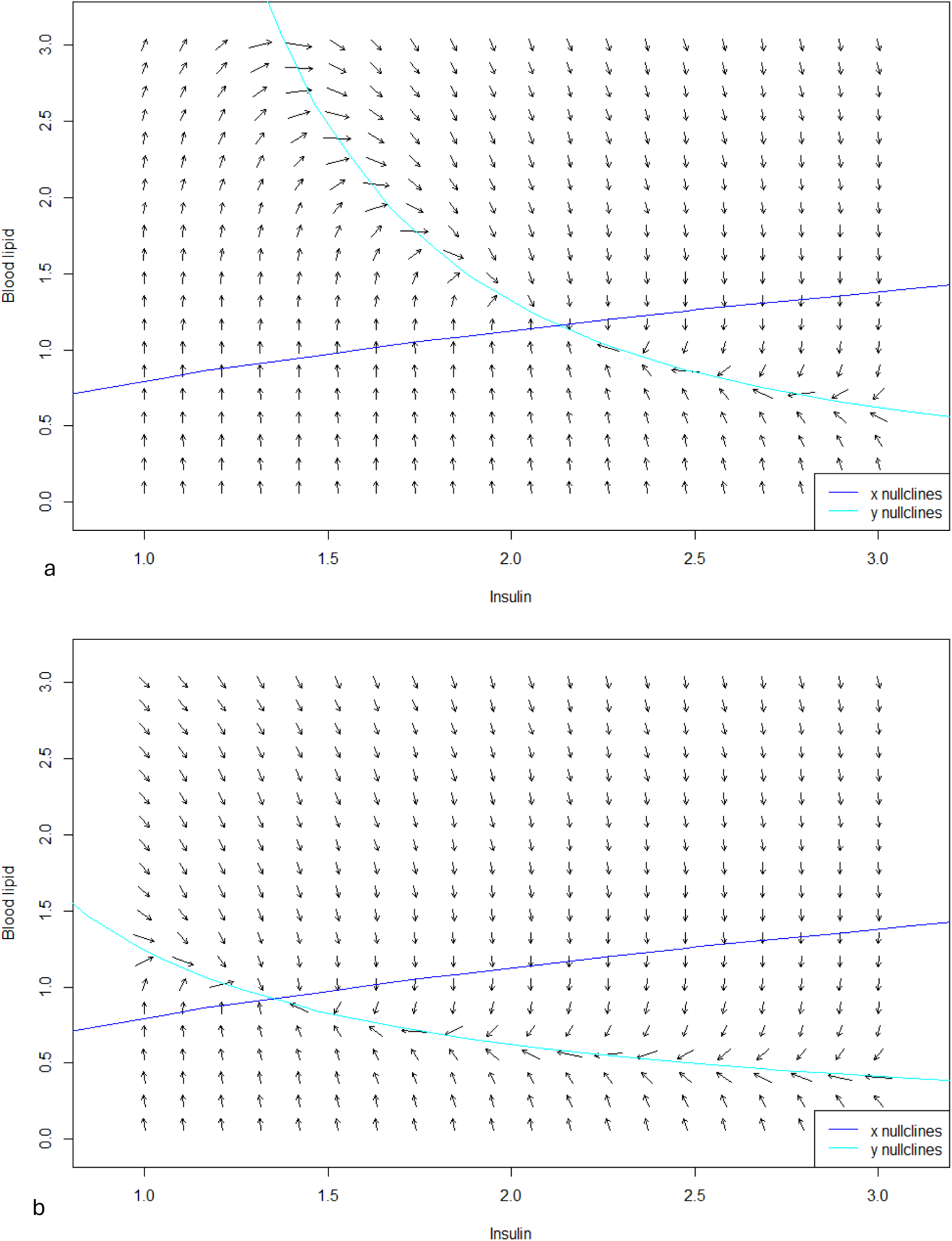
Phase plane analysis of relationship between L_B_ and insulin from simplified FEM (A) with export function. (B) without export function.

L_B_ is swiftly returned to equilibrium by complimentary changes in insulin secretion, driven in the model by changes in β-cell mass, and mass-independent production rate (per cell); in equation S3.2 for insulin this is represented by βM and L_B_^2^, respectively. This is not wholly accurate as the rate of insulin production per cell and likely β-cell mass will both have maximum values that limit the body’s capacity to respond to rising blood lipid levels. The significance a maximum β-cell mass is discussed later.

Prior to the limits of insulin production, of the four metabolite variables included in the FEM, L_I_ is the only one not kept at strict equilibrium. ATP is maintained by IMC, ACoA by ATP, and L_B_ by insulin. As shown in equation S8, the model could be viewed as having two sides, where internal and external factors regulated by processes outside those in the FEM, here called activities (green), impact homeostasis through equations S1-4 to result in increased L_I_.

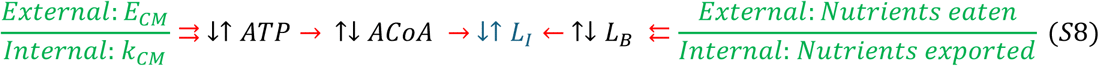

ATP and L_B_ are the metabolites primarily affected by activity changes (for ‘internal: nutrients exported’, see discussion of equations S3.6-S3.8); however, both return to equilibrium after shifting away. ATP is returned by changes in APMM, which alters levels of ACoA until it results in the reciprocal effect on β-oxidation, leading ultimately to changes in L_I_. Thus, as indicated in the parameter estimation in Figure 16, while changes in k_CM_ had no effect on ATP, ACoA, or L_B_, they did affect L_I_. Equally, L_B_ is returned to equilibrium by changes in import and export that, in turn, affect L_I_. Hence the meal value also had no effect on ATP, ACoA, or L_B_, but did affect L_I_.

Indeed, L_I_ is influenced by most of the parameters and processes in the model, as shown in Figure 16. As changes in activity induce downstream changes in IMC and insulin production to keep the other metabolites at equilibrium, as seen equation S4 copied here, the downstream effect is on the rates of β-oxidation (blue), and import and export (green/orange) which impact L_I_. The mechanism responsible for maintaining L_I_ homeostasis is inflammation (black).

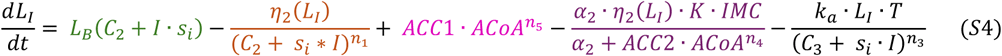

We therefore did phase plane analysis using a simplified model containing only equation S4 for L_I_ and S4.2 for inflammation, where L_I_ and inflammation were the only variables, which produced a stable fixed point at high L_I_, as shown in Figure 20. Between values of 0-200, L_I_ increases gradually, tempered by increasing inflammation. Beyond this, the inflammation response is muted and L_I_ rises more quickly.

**Figure 20.**
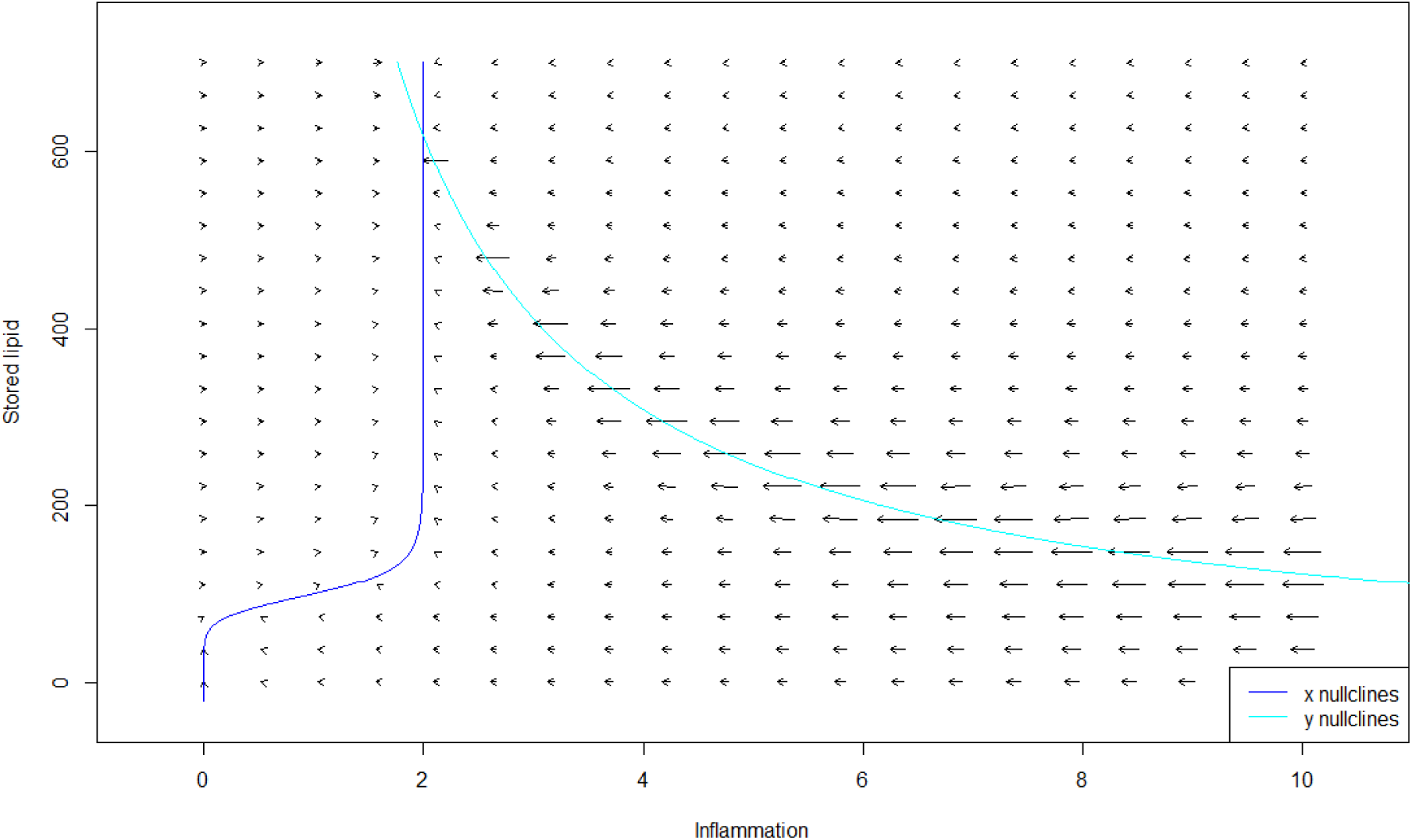
Phase plane analysis of relationship between L_I_ and inflammation from simplified FEM.

Phase plane analysis suggested the model was responding appropriately, and the dynamics would be largely independent of the initial inputs; however, we could not rule out that combining the equations in the FEM would reduce stability and create additional fixed points.

##### 9.2.1.3 Randomisation of initial variables

Therefore, we tested the robustness of homeostasis for 21 runs of the model with random initial value combinations, comparing simulations to the non-randomly chosen values (termed ‘Chosen’), as shown in Table 4. We selected either multiplying or dividing the chosen value by 10 for most variables. For s_i_ we selected random values between 0.01-1, for L_I_ random values between 10-1000, and for inflammation we selected random values from 1-50.

**Table 4.** Initial values generated randomly for 21 FEM runs. Abbreviations, see Table 6.

| Run | I | $\beta M$ | $s_i$ | $L_I$ | $L_B$ | TNFα | K | IMC | ACoA | ATP | $k_{CM}$ | $E_{CM}$ |
| --- | --- | --- | --- | --- | --- | --- | --- | --- | --- | --- | --- | --- |
| Chosen | 2.3955 | 1 | 0.965678 | 100 | 5 | 0 | 5 | 0.04 | 5 | 16.66667 | 1 | 1 |
| 1 | 2.3955 | 1 | 0.396053 | 92 | 5 | 42 | 5 | 0.04 | 5 | 53 | 1 | 1 |
| 2 | 23.955 | 10 | 0.450353 | 405 | 50 | 6 | 0.5 | 0.4 | 5 | 63 | 0.1 | 0.1 |
| 3 | 23.955 | 1 | 0.706795 | 875 | 5 | 42 | 5 | 0.4 | 0.5 | 71 | 0.1 | 0.1 |
| 4 | 23.955 | 0.1 | 0.08342 | 286 | 50 | 42 | 0.5 | 0.04 | 50 | 84 | 1 | 10 |
| 5 | 23.955 | 1 | 0.339973 | 218 | 50 | 38 | 0.5 | 0.04 | 5 | 82 | 1 | 1 |
| 6 | 2.3955 | 10 | 0.681107 | 85 | 5 | 48 | 50 | 0.004 | 50 | 93 | 0.1 | 0.1 |
| 7 | 0.23955 | 10 | 0.317632 | 103 | 0.5 | 16 | 0.5 | 0.04 | 5 | 17 | 10 | 1 |
| 8 | 0.23955 | 1 | 0.831737 | 812 | 50 | 35 | 50 | 0.04 | 0.5 | 97 | 10 | 1 |
| 9 | 23.955 | 10 | 0.215957 | 39 | 0.5 | 36 | 5 | 0.4 | 0.5 | 2 | 0.1 | 0.1 |
| 10 | 0.23955 | 10 | 0.498451 | 226 | 0.5 | 27 | 50 | 0.004 | 50 | 49 | 10 | 10 |
| 11 | 23.955 | 10 | 0.276774 | 955 | 50 | 27 | 0.5 | 0.4 | 50 | 2 | 0.1 | 0.1 |
| 12 | 2.3955 | 10 | 0.192831 | 184 | 5 | 35 | 50 | 0.004 | 0.5 | 13 | 10 | 1 |
| 13 | 23.955 | 10 | 0.950671 | 383 | 0.5 | 23 | 0.5 | 0.4 | 50 | 24 | 0.1 | 0.1 |
| 14 | 23.955 | 10 | 0.322404 | 332 | 0.5 | 3 | 0.5 | 0.4 | 5 | 49 | 0.1 | 0.1 |
| 15 | 2.3955 | 1 | 0.478978 | 124 | 50 | 31 | 0.5 | 0.04 | 0.5 | 67 | 1 | 10 |
| 16 | 23.955 | 0.1 | 0.028965 | 386 | 50 | 48 | 0.5 | 0.4 | 5 | 82 | 1 | 10 |
| 17 | 0.23955 | 1 | 0.547912 | 859 | 50 | 2 | 0.5 | 0.4 | 5 | 2 | 0.1 | 1 |
| 18 | 2.3955 | 1 | 0.644596 | 617 | 0.5 | 47 | 50 | 0.04 | 0.5 | 37 | 1 | 1 |
| 19 | 2.3955 | 0.1 | 0.596667 | 474 | 50 | 23 | 5 | 0.004 | 0.5 | 63 | 10 | 1 |
| 20 | 2.3955 | 1 | 0.322615 | 367 | 0.5 | 3 | 0.5 | 0.04 | 5 | 76 | 1 | 1 |
| 21 | 2.3955 | 1 | 0.891223 | 691 | 50 | 9 | 5 | 0.04 | 5 | 13 | 1 | 1 |

We plotted all runs together and results indicated that multiple runs converged on the same value for ATP (Figure 21A), ACoA (Figure 21B), and L_B_ (Figure 21C). However, while the L_B_ equilibrium was stable, several runs showed ATP did not always converge on the same equilibrium value as with the chosen starting values, potentially not reaching equilibrium at all. We predicted that this may reflect that high values of E_CM_ and k_CM_ would burn more lipid than was replenished from Meal values to sustain the rate of AHCM, depleting L_I_ below the level defined by the η_2_ switch function which would inhibit export and β-oxidation, preventing formation of ACoA and ATP. We removed all runs (white rows in Table 4) during which L_I_ dropped below 1 au, which included all the runs with high E_CM_, k_CM_, or both. No other runs showed the profile of quick ATP depletion (Figure 21D); however, some runs still showed no convergence of ATP to equilibrium. As these runs all had high initial spikes of ATP suggesting they resulted from slower AHCM, we assessed whether they could be explained by the low k_CM_/E_CM_ values. We removed all runs (green rows in Table 4) which had E_CM_ values of 0.1, and found that this did indeed account for all runs still failing to reach equilibrium (Figure 21E, remaining runs are in yellow in Table 4).

**Figure 21.**
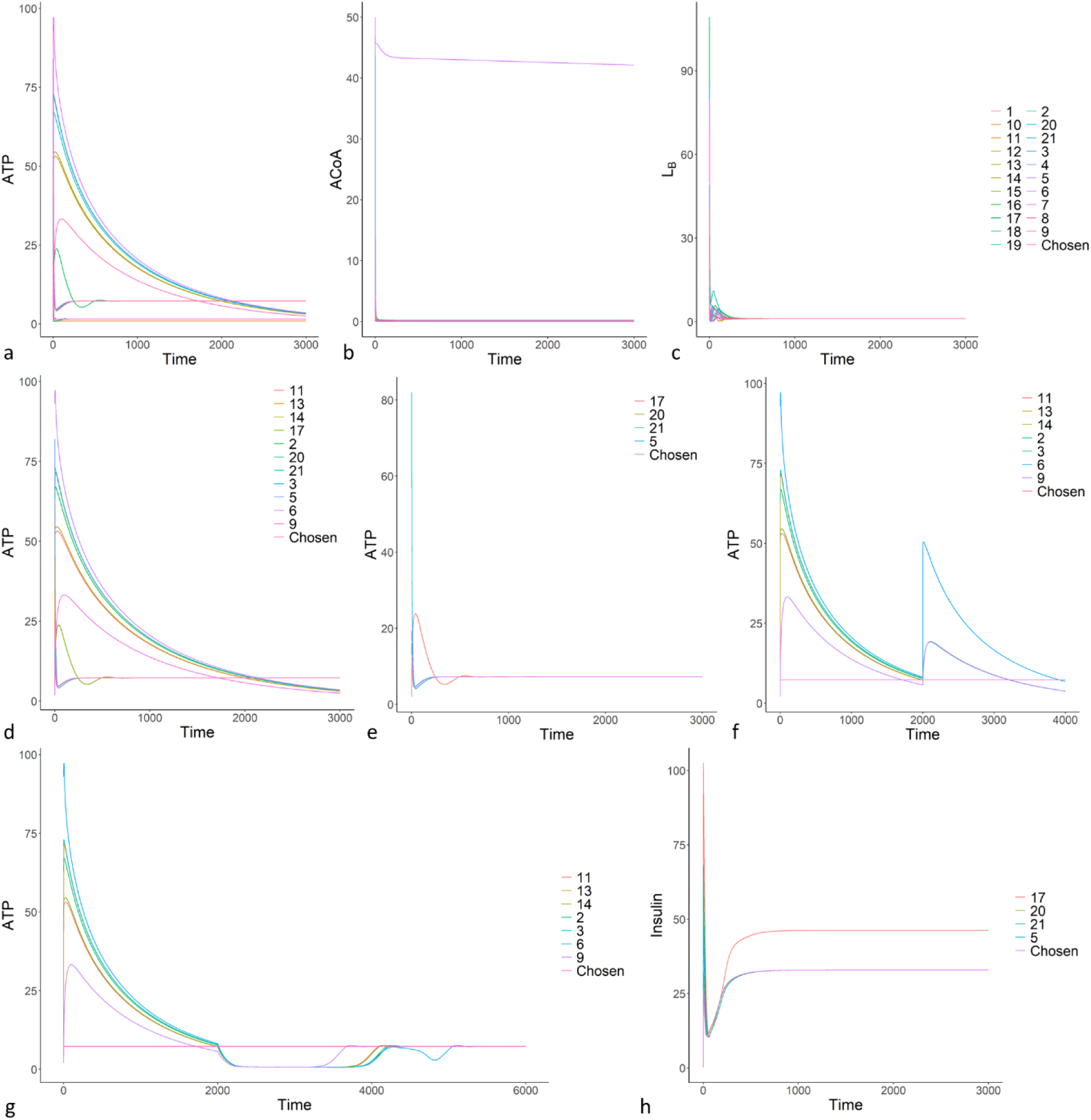
Runs of FEM with random initial variable values. (A-C) All runs including non-random chosen variable values for (A) ATP; (B) ACoA; and (C) L_B_. (D) ATP for runs where L_I_ does not drop below 1 au. (E) ATP for runs where E_CM_ = 1. (F) ATP for runs where E_CM_ < 1, where IMC is artificially restored to 0.1 at time 2000 au. (G) ATP for runs where E_CM_ < 1, where E_CM_ is artificially restored to 1 at time 2000 au. (H) Insulin for runs where E_CM_ = 1. Abbreviations, see Table 6.

To understand why the low AHCM runs failed to reach equilibrium we first increased IMC at time 2000 au. The minimum values of IMC in these runs were critically low, ranging from 10^-16^-10^-22^, but this did not allow ATP to reach equilibrium (Figure 21F), suggesting it was not simply a model equilibration effect. Restoring E_CM_ value to 1 at time 2000 au caused all runs to converge on the same equilibrium as the chosen initial values (Figure 21G). Notably, while the FEM maintains robust homeostasis of key metabolites, this is achieved by other model components shifting to different equilibria, such as insulin (Figure 21H).

These results suggest that, excluding extreme values of fixed variables determined by factors outside the model, the FEM is highly stable, converging on the same equilibrium values for key metabolites independent of the starting values for most variables. We could therefore use the model to show how changes in metabolic rate and meal values would impact fat and energy metabolism and the likely effects on human health.

#### 9.2.2 Stepwise changes in metabolic rate and meal

Initial simulations of the FEM ran for 2000 au, allowing all variables and processes to reach equilibrium long before t=1900 au. Therefore, for all future data we removed t<1900 au so the dynamics could be compared (visually and numerically) to equilibrium values rather than including the equilibration effects.

As shown in Figure 22, we made sure the model responded appropriately to single step changes in k_CM_ (black, defining the basal metabolic rate and intrinsic capacity for AHCM) and meal values (red).

**Figure 22.**
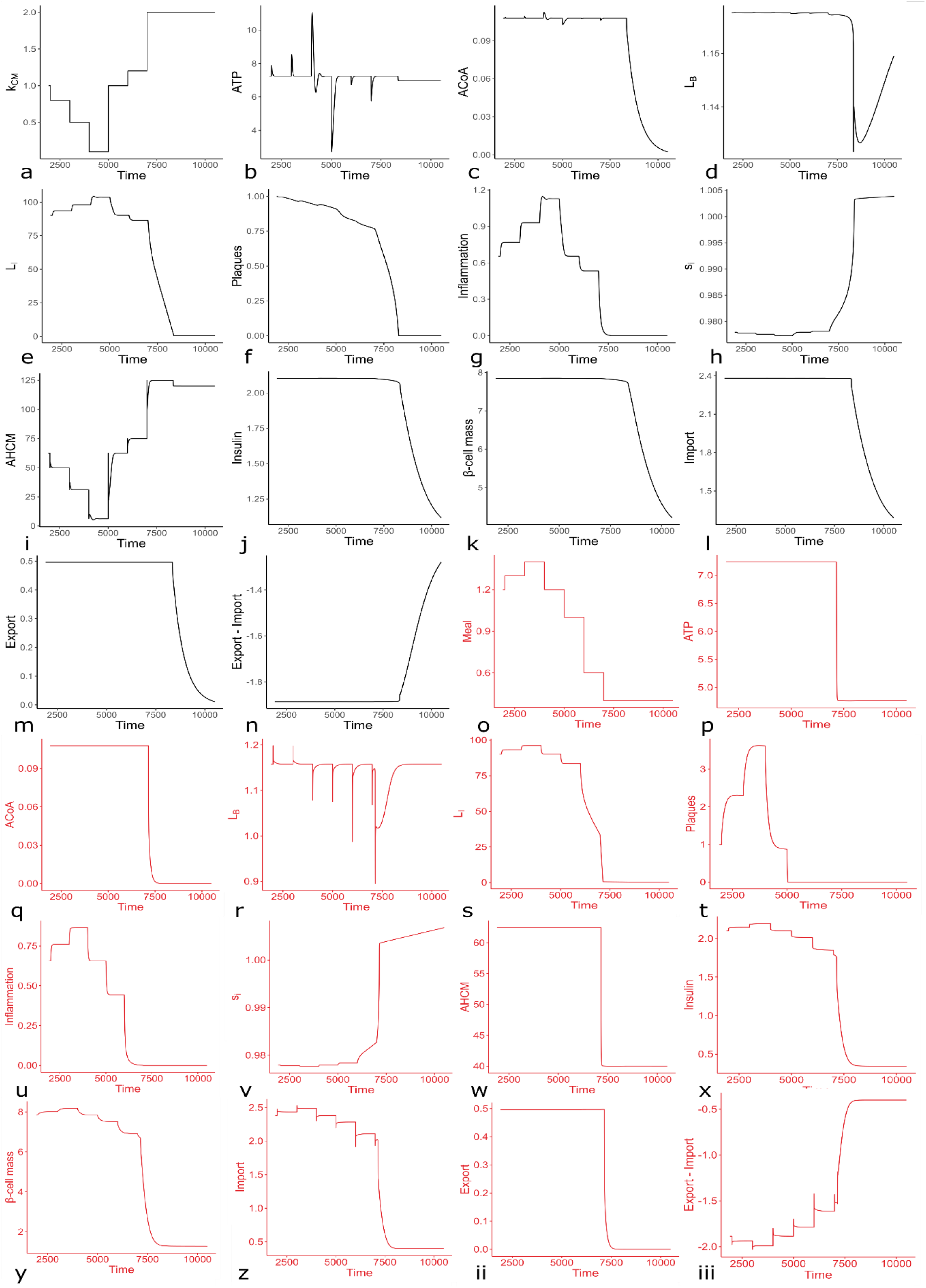
Effects of step changes in k_CM_ (black) and Meal (red) on key parameters. (A-N) Effects of step changes in k_CM_. (A) Input changes in k_CM_. (B) ATP. (C) ACoA. (D) L_B_. (E) L_I_. (F) Plaques. (G) Inflammation. (H) s_i_. (I) AHCM. (J) Insulin. (K) β-cell mass. (L) Import. (M) Export. (N) Export minus import. (O-iii) Effects of step changes in Meal. (O) Input changes in Meal. (P) ATP. (Q) ACoA. (R) L_B_. (S) L_I_. (T) Plaques. (U) Inflammation. (V) s_i_. (W) AHCM. (X) Insulin. (Y) β-cell mass. (Z) Import. (ii) Export. (iii) Export minus import. Abbreviations, see Table 6.

The results indicated that single changes in k_CM_ and Meal caused appropriate changes in all variables and processes as suggested by the literature and described in the model construction. ATP, ACoA, and L_B_ all retained a single equilibrium value despite changes in metabolic rate and meal, suggesting the FEM could robustly maintain these equilibria in response to changes in diet and metabolism. Equilibrium L_I_ shifted as predicted from the data suggesting stored L_I_ (and weight) has no fixed value per individual and shifts according to metabolic and dietary factors^197^.

Importantly, the equilibria for ATP, ACoA, and L_B_ only shifted when either metabolic rate (and thus L_I_ burning) increased above the capacity of the (fixed) Meal value to replenish it, or the Meal value decreased below the necessary (fixed) rate of AHCM to maintain cellular function. Both caused the depletion of L_I_ below the threshold determined by equation S2.2, at which point the model had no remaining stored fat to replenish L_B_ or utilise in metabolism. In reality, this would reflect death by starvation. There are notably multiple mechanisms to supply fuel for the AHCM of crucial tissues as L_I_ (and other readily available fuel) borders exhaustion, which mainly involve the breakdown of less important tissues^652^. As the FEM has not been built to address these pathways, they are not included, and the simulations suggested that all variables and processes responded appropriately at least up to the point of L_I_ exhaustion. We therefore modelled continuous metabolic slowdown.

#### 9.2.3 Gradual AHCM slowdown

Metabolic slowdown was modelled as a continuous decline in k_CM_ over time (from t=2000 au), as defined by equation S6, copied here.

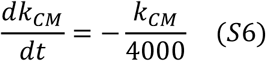

When we steadily reduced the basal AHCM rate (k_CM_, Figure 23A), causing total AHCM to decline (not shown), the FEM maintained ATP (Figure 23B) by reducing intrinsic mitochondrial capacity (not shown), predicting decreasing mitochondrial function, which reduced APMM (Figure 23C). ACoA was maintained by a corresponding decrease in β-oxidation (Figure 23D-E), which shifted L_I_ to a higher equilibrium (Figure 23F), reducing s_i_ (Figure 23G) predicting weight gain and IR. Inflammation and apoptosis increased to retard the increasing L_I_ (Figure 23H-I), predicting chronic inflammation. Notably, the FEM did not predict atherosclerosis as L_B_ was kept in equilibrium by increasing βM and insulin levels (Figure 23J-L). However, the model allowed both adipose and β-cell mass to increase indefinitely.

**Figure 23.**
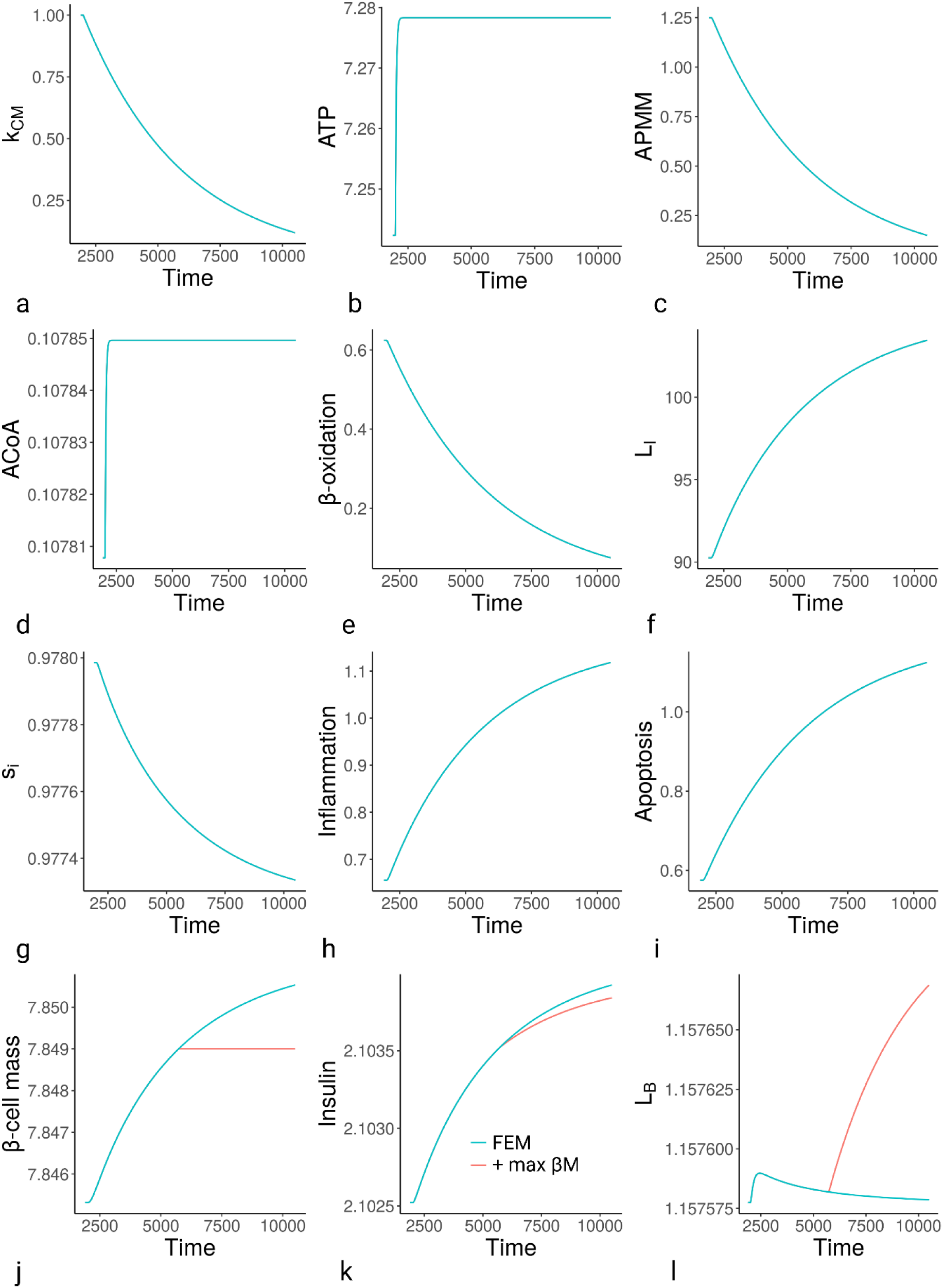
Simulations of the FEM in response to declining k_CM_. (A) k_CM_ input. (B) ATP. (C) APMM. (D) ACoA. (E) β-oxidation. (F) L_I_. (G) s_i_. (H) Inflammation. (I) Apoptosis. (J) β-cell mass. (K) Insulin. (L) L_B_. Simulations start at t=1900 au, and k_CM_ declines according to equation S5 from t=2000 au. All graphs include effects of a maximum βM of 7.849 au as red line; however, it only affects Figures J-L. Abbreviations, see Table 6.

In reality both adipose mass (i.e. L_I_ storage capacity) and βM would be finite, limited by organ size. Notably, the same is true for excreting excess L_B_ via the liver in the bile (reverse cholesterol transport, RCT). To represent all these finite capacities, we added a maximal βM to the FEM (as it already included βM and not liver or adipose mass) as in equation S7.

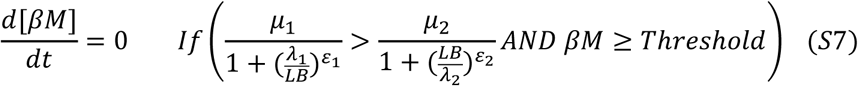

Once βM reached the threshold (7.849, chosen based on previous simulations) it could no longer increase, preventing the necessary increase in insulin production to sustain L_B_ as AHCM continued to decline. When we introduced a maximum βM, reflecting that the pancreatic islets cannot expand forever, once βM plateaued, rising insulin could no longer compensate for additional lipid, and L_B_ began to rise (Figure 23J-L), thus predicting atherosclerosis.

To supplement the results shown in Figure 23J-L, we included an additional scenario in simulations where the declining k_CM_ from t = 2000 was reversed at t = 6250 (after the βM plateau) to show that βM could still decrease from maximal size as required for homeostasis. The results for key variables are shown in Figure 24. ACoA mirrors ATP (not shown).

**Figure 24.**
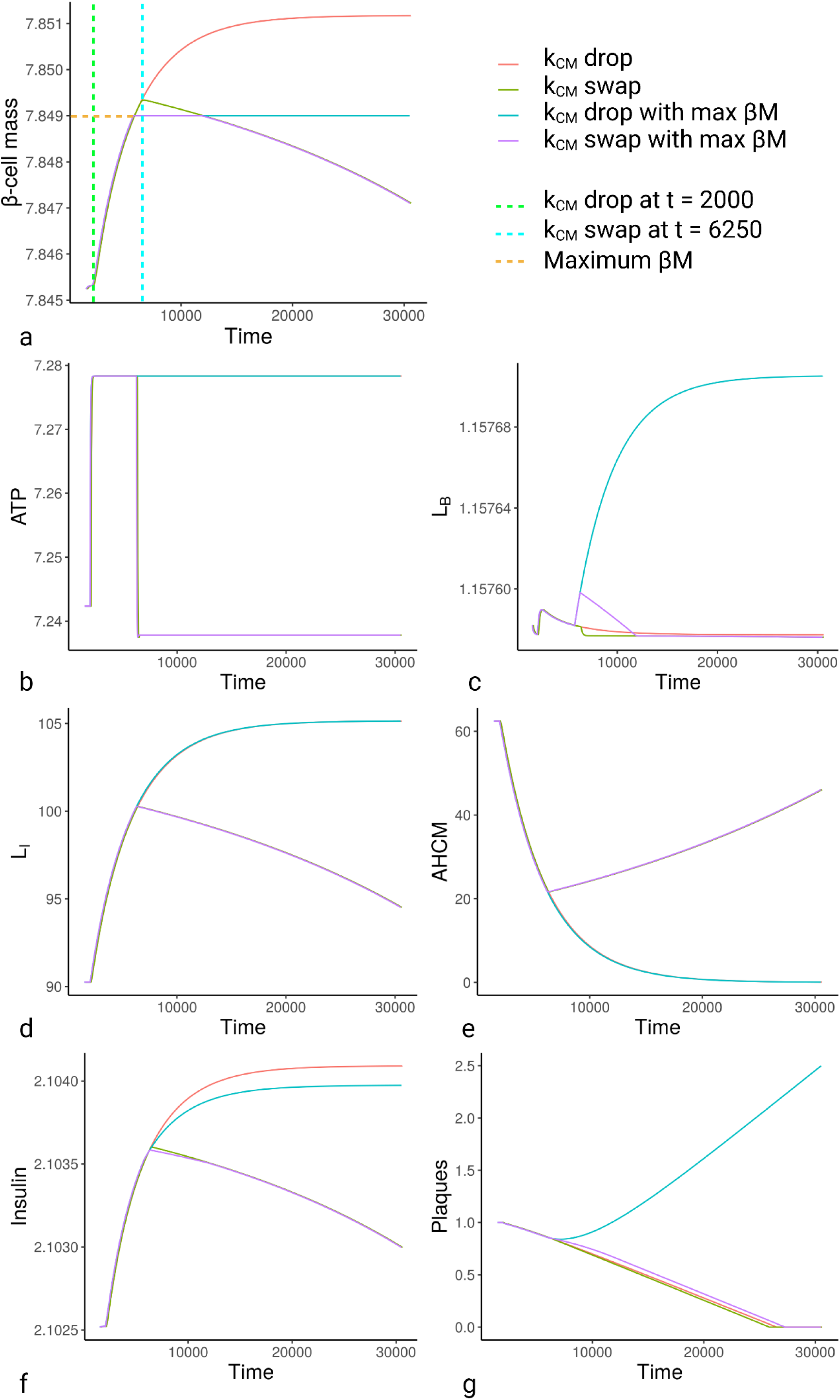
Effect of maximum value of βM on key variables. To demonstrate the effects of a maximum value of βM of 7.849 au, we simulated declining metabolic rate (k_CM_ drop) with and without the max βM. To demonstrate that βM could decrease from this mass as required by energy homeostasis, we also simulated a rise in k_CM_ after the maximum βM had been reached (k_CM_ swap). As some simulations were identical, we added +100 au to the time of k_CM_ drop and k_CM_ swap so they could be distinguihsed from the other lines on the graphs. (A) β-cell mass. (B) ATP. (C) L_B_. (D) L_I_. (E) AHCM. (F) Insulin. (G) Plaques. Abbreviations, see Table 6.

A further parameter estimation of the FEM including a maximum βM demonstrated that the rate of insulin production, q, and the rate of insulin degradation, γ_i_, gained additional significance affecting multiple variables and processes, as did k_s_ and k_r_ determining the rate of change in insulin sensitivity, as shown in Figure 25A (compared to Figure 16). The results were as predicted based on existing work by Karin, et al.^195^ examining the role of dynamic compensation in glucose homeostasis (see discussion). The other equation which provided dynamic compensation was the expansion and contraction of intrinsic mitochondrial capacity. We therefore removed the role of IMC from the β-oxidation equation, as shown in equation S9)

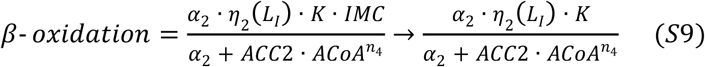

**Figure 25.**
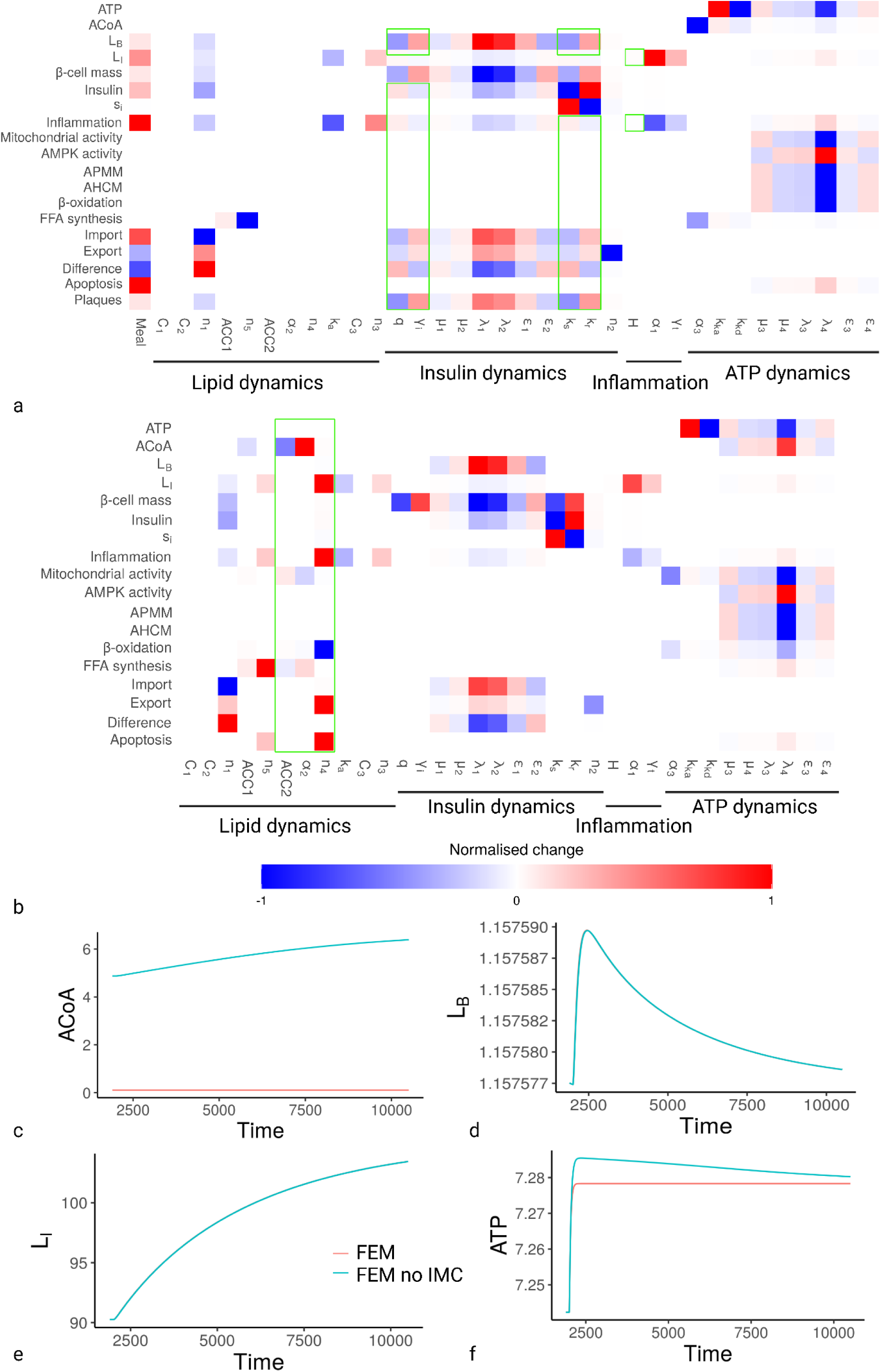
Parameter estimation and variables for modified FEMs. (A) FEM undergoing metabolic slowdown with maximum βM of 7.849 au. (B) FEM without IMC regulation of β-oxidation. (A-B) Values were normalised as described in the model running and storage so that each value is relative to other parameters for each individual variable/process. An absolute value of 1 (or −1) represents the parameter that has the greatest effect on that variable or process. Positive values reflect that increases in the parameter increase the variable or process, while negative values reflect that increases in the parameter decrease the variable or process. Green boxes show differences to parameter estimation under basal conditions. (C-F) Simulations of FEM without IMC regulation of β-oxidation. (C) ACoA. (D) L_B_. (E) L_I_. (F) ATP. Abbreviations, see Table 6.

The parameters α_2_, ACC2, and n_4_ which were part of the β-oxidation equation (S2.1) previously had no effect on any variables or processes. Importantly, these parameters govern how ACoA impacts β-oxidation, and dynamic compensation kept ACoA in equilibrium during metabolic slowdown. As indicated by the results in Figure 25B, all three parameters had multiple impacts on variables and processes when β-oxidation was no longer regulated by IMC, although only n_4_ affected the rate of β-oxidation itself, while ACC2 and α_2_ primarily impacted levels of ACoA, FFA synthesis, and IMC. Removing IMC from the β-oxidation equation demonstrates the robustness of the FEM to maintain the various equilibria: although ACoA levels rise, this does not impact L_B_, or L_I_, and ATP still returns to equilibrium, although more slowly (Figure 25C-F). We further discuss the significance of dynamic compensation in the Model Discussion below.

The results suggested that the FEM could produce multiple ageing phenotypes via a single homeostatic cascade. We therefore tested how the FEM would respond to anti-ageing interventions.

#### 9.2.4 Anti-ageing interventions and metabolic slowdown

##### 9.2.4.1 Diet and Exercise

We modelled calorie restriction (CR) in the FEM as a reduction in the meal value shown in equation S3. As CR did nothing to prevent the underlying metabolic slowdown, not affecting ATP or AHCM (Figure 22P & W), its beneficial effects occurred by reducing L_I_ and chronic inflammation and increasing s_i_ (Figure 22S & U-V), consistent with the literature^359,360^.

To observe how CR would impact mortality, we utilised rising L_B_ as a proxy for atherosclerotic risk. L_B_ began to rise when the βM threshold was reached, occurring earlier for AL than CR (Figure 26A).When we swapped CR to ad libitum (AL), all variables including L_B_ (Figure 26A) returned to the levels of continuous AL, suggesting that CR should not provide lasting health benefits after the regimen ceases. Consistently, Mair, et al.^231^ swapped flies between dietary restriction (DR) and AL and found mortality profiles that matched closely to those produced by the FEM (see their Figure 1B for comparison). These profiles suggested that CR was primarily risk-preventative, affecting only the ultimate inducer of atherosclerosis by reducing L_B_, but not affecting the underlying cause of slow L_B_ accumulation (ie. selective destruction and metabolic slowdown).

**Figure 26.**
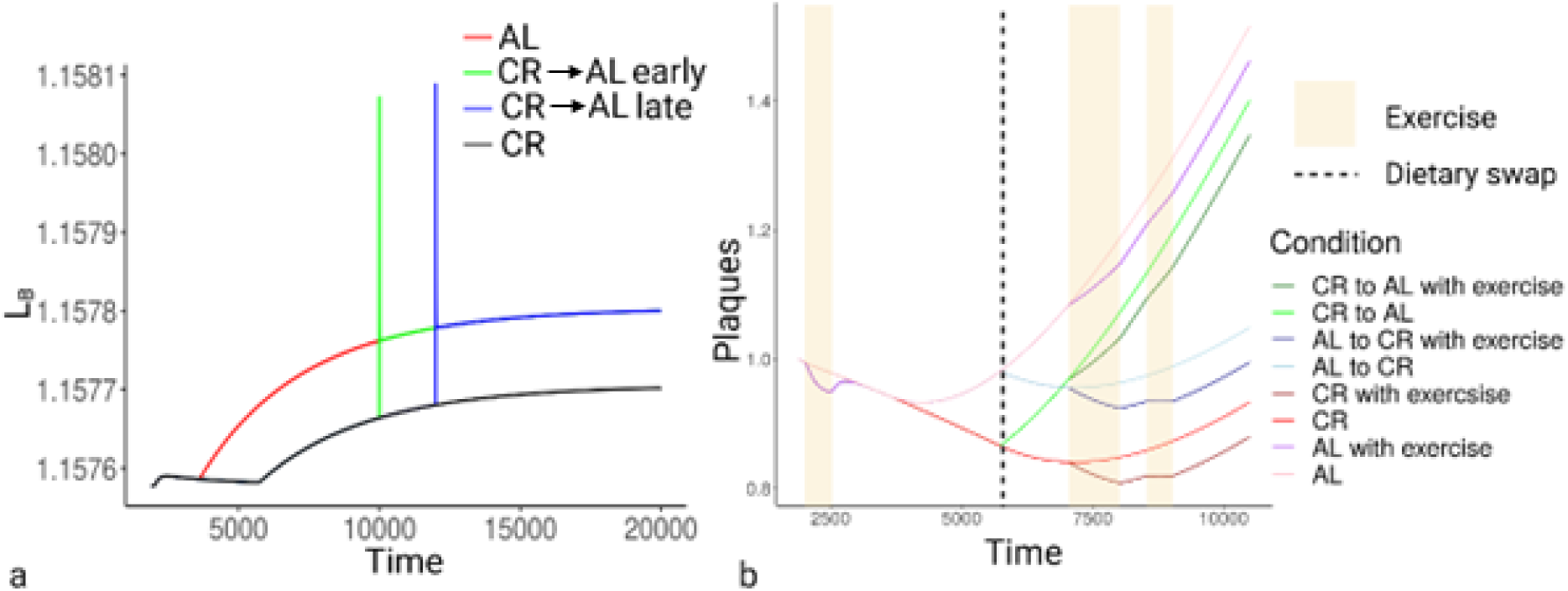
Effects of anti-ageing interventions on energy homeostasis. (A) Effects of reduced Meal (mimicing CR) on L_B_. (B) Combined effects of CR and exercise regimens on mortality as a function of plaque number and size. Abbreviations, see Table 6.

When the same lab tested swapping between CR and AL in mice, they observed some memory effect^361^, suggesting CR may have gero-protective properties in mice, if not flies. Notably, multiple studies and meta-analyses indicate the beneficial effects of exercise on dislipidemia and mortality, and the benefits are not limited to the exact times the individuals are exercising^653,654^, but are limited to healthspan and mean lifespan with no maximum lifespan extension^233–235^. These long-lasting benefits may thus still be primarily risk preventative rather than gero-protective. In asking whether the FEM could also predict this, we considered a more realistic risk assessment (than L_B_ alone) would reflect the levels of atherosclerotic plaques from excess L_B_. While L_B_ can change quickly, plaques are both formed and degraded more slowly, low L_B_ allowing net degradation, with higher L_B_ shifting the parameters so plaque formation is faster than degradation (see equation S3.11). From this we modelled regimens of CR (low meal) and exercise (temporary increases in AHCM), demonstrating that both regimens could induce long-lasting benefits on plaque levels while still being entirely risk preventative (Figure 26B). However, these results do not rule out additional gero-protective effects.

##### 9.2.4.2 Metformin

Metformin is primarily regarded as an AMPK activator, but its mechanisms are not well understood^398^. To model the impact of metformin we altered several parameters, firstly, k_ka_ and k_kd_ which determine the rates of AMPK activation and deactivation respectively. We simulated two doses of metformin, introduced between t = 2500 and t = 5000 au and then between t = 7500 and t = 10000 au, either by increasing k_ka_*1.5 or reducing k_kd_/1.5 to represent increased rate of activation and decreased rate of deactivation, respectively. The results were identical for changes in k_ka_ and k_kd_ (latter not shown). Increasing k_ka_ and its effects on L_B_, plaques, AMPK activity, ATP, inflammation, AHCM, and APMM are shown in Figure 27A-D.

**Figure 27.**
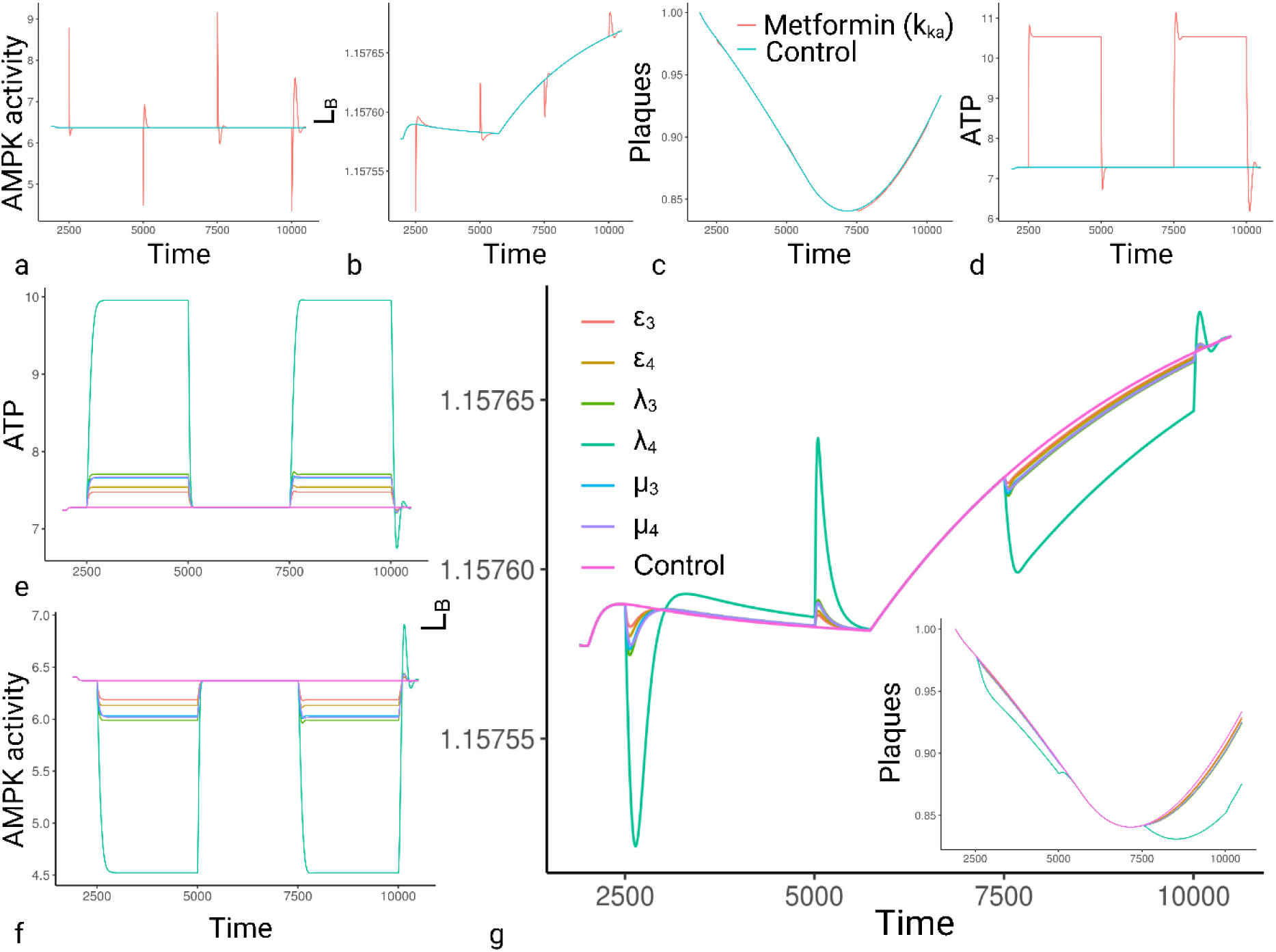
Effects of anti-ageing interventions on FEM variables and processes. (A-M) Treatments were two doses introduced between t = 2500 and t = 5000 au and then between t = 7500 and t = 10000 au. (A-D) Metformin modelled as a change in k_ka_ for AMPK activation (or k_kd_ for deactivation). (A) AMPK activity. (B) L_B_. (C) Plaques. (D) ATP. (E-G) Metformin modelled as a change in parameters affecting AMPK regulation of IMC. (E) ATP. (F) AMPK activity. (G) L_B_ and plaques (bottom right). Abbreviations, see Table 6.

Increasing the rate of AMPK activation (or decreasing rate of deactivation) had little impact on the FEM. Most variables and processes spiked (up or down) at the point k_ka_ (and k_kd_) changed, but quickly returned to control levels, such as L_B_ (Figure 27B), while the number and size of associated plaques showed little change (Figure 27C), despite shifting the equilibrium level of ATP (Figure 27D). Adding a variable for ATP-independent AMPK activation, k_in_ (value 0 in control and 0.5 for metformin), as in equation S10, had the same lack of effect (data not shown).

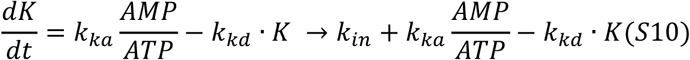

The results reflect that AMPK activity is not primarily affected by the rates of activation and deactivation (dependent or independent of ATP). AMPK activity is determined by the ratio of ATP:AMP, activating and deactivating when levels are out of equilibrium and returning to basal levels when equilibrium returns.

We therefore modelled changes in the parameters μ_3_ and μ_4_, λ_3_ and λ_4_, and ε_3_ and ε_4_, determining how AMPK regulates IMC (altering the impact on IMC for a given activity of AMPK). We multiplied or divided all parameter values by 1.5 so that each change either increased the AMPK-activated IMC or prevented the inhibition, as shown in Table 5. Results for ATP, AMPK activity, L_B_, and plaques are shown in Figure 27E-G.

**Table 5.** IMC parameters and value changes representing metformin. Abbreviations see Table 6.

| Parameter (code version) | FEM | FEM + Metformin |
| --- | --- | --- |
| $\mu_3$ (mu3) | 0.021 | 0.0315 |
| $\mu_4$ (mu4) | 0.025 | 0.01667 |
| $\lambda_3$ (lam3) | 10.892 | 7.26133 |
| $\lambda_4$ (lam4) | 5.6 | 3.7333 |
| $\varepsilon_3$ (eta3) | 1.7 | 1.1333 |
| $\varepsilon_4$ (eta4) | 8.5 | 12.75 |

**Table 6.** Table of abbreviations.

| Abbreviation | Full term | Abbreviation | Full term |
| --- | --- | --- | --- |
| **ACC1** | Acetyl-CoA carboxylase 1 | **k<sub>CM</sub>** | Intrinsic capacity for cellular metabolism |
| **ACC2** | Acetyl-CoA carboxylase 2 | **L<sub>B</sub>** | Blood lipid |
| **ACoA** | Acetyl-coenzyme A | **LD** | Lipid droplet |
| **AHCM** | ATP hydrolysing cellular metabolism | **LDL** | Low density lipoprotein |
| **AL** | Ad libitum | **LDL-R** | LDL-receptor |
| **AMP** | Adenosine monophosphate | **LFEM** | Linear FEM |
| **AMPK** | AMP-activated protein kinase | **L<sub>i</sub>** | Intracellular lipid |
| **APMM** | ATP producing mitochondrial metabolism | **M** | Mitochondria |
| **ATP** | Adenosine triphosphate | **MM** | Mitochondrial mass |
| **au** | Arbitrary units | **MAPK** | Mitogen-activated protein kinase |
| **βM** | β-cell mass | **MD** | Mitochondrial dysfunction |
| **BMI** | Body mass index | **mTOR** | Mammalian target of rapamycin |
| **CD36** | Cluster of differentiation 36 | **mTORC1** | mTOR complex 1 |
| **CDC FEM** | Complete dynamic compensation FEM | **OXPHOS** | Oxidative phosphorylation |
| **CLS** | Crown-like structure | **PGC1α** | Peroxisome proliferator-activated receptor-γ coactivator-1α |
| **CR** | Calorie restriction | **RCT** | Reverse cholesterol transport |
| **DR** | Dietary restriction | **ROS** | Reactive oxygen species |
| **DST** | Disposable soma theory | **SENS** | Strategies for Engineered Negligible Senescence |
| **ECM** | Extracellular matrix | **SDT** | Selective destruction theory |
| **FEM** | Fuel and Energy Model | **s<sub>i</sub>** | Insulin sensitivity |
| **FFA** | Free fatty acid | **TFEB** | Transcription factor EB |
| **GH** | Growth hormone | **TLR4** | Toll-like receptor 4 |
| **IC** | Immune cell | **TNF-α** | Tumour necrosis factor-α |
| **IGF-1** | Insulin-like growth factor-1 | **UC** | Degree of coupling |
| **IMC** | Intrinsic mitochondrial capacity | **UC FEM** | Uncoupling FEM |
| **IR** | Insulin resistance | **UPR** | Unfolded protein response |

Thus, as all parameters for the impact of K on IMC produced similar curves, and λ_4_ produced the greatest effect, we modelled metformin using λ_4_, as shown in Figure 28A-C, predicting transient effects on L_B_ and inflammation during the regimen. However, as indicated in Figure 27G, while effects on L_B_ might be transient, they produced extended reduction of risk from plaque formation (as modelled by equation S3.11).

**Figure 28.**
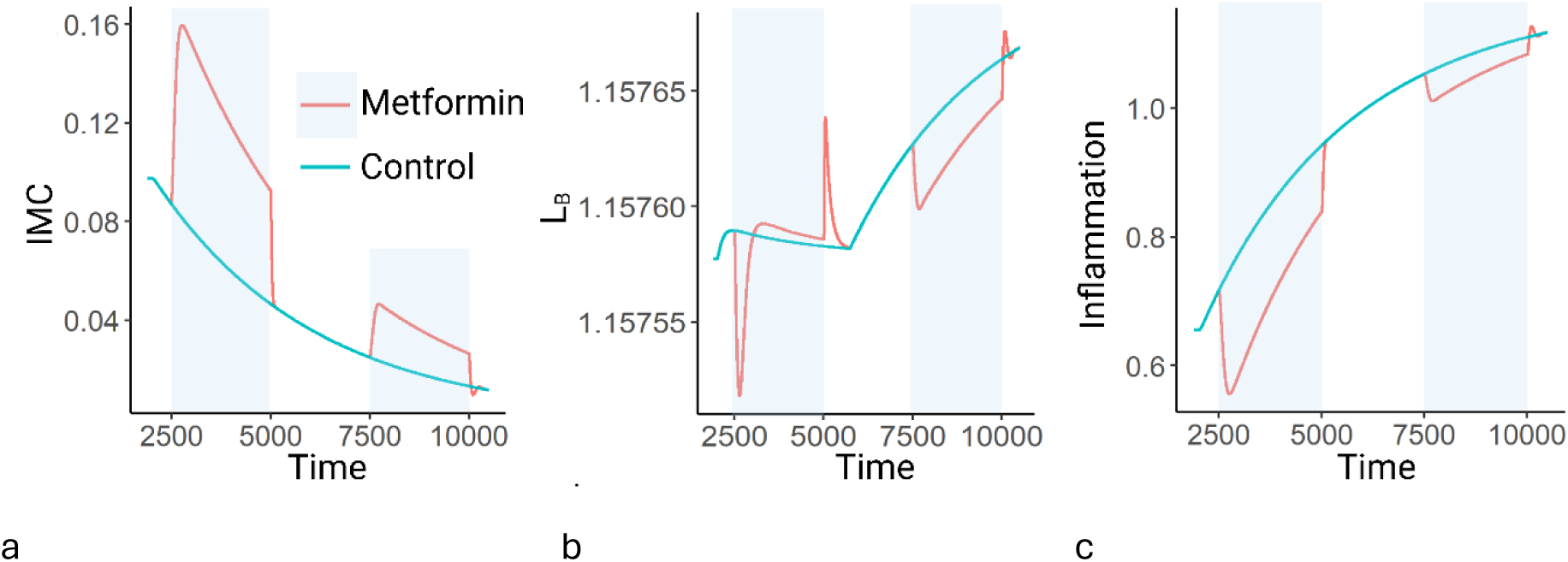
Effects of metformin (as represented by a reduction in λ_4_ on: (A) ATP; (B) L_B_; (C) inflammation. Abbreviations, see Table 6.

Notably, there is little evidence that metformin extends maximum lifespan in mice^404^, flies^405^, or worms^406^. The results suggest that if metformin acts by the mechanisms suggested here that it is not gero-protective, but risk-preventative.

##### 9.2.4.3 Rapamycin

Rapamycin is probably the most consistent longevity drug yet documented, enhancing maximum lifespan in mice^371,372^ and flies^373^, including long-lasting effects after the regimen ceases^374^. It has multiple effects on cell and molecular phenotype, including reducing cell division rate, but these are all (at least at low dose) thought to result from the inhibition of mTORC1^375,376^, which is a central driver of cell growth and metabolism^377^, essentially driving AHCM. Fascinatingly, this is essentially mimicking metabolic slowdown, which the FEM predicts should accelerate ageing, the increase of L_I_, basal inflammation, and atherosclerosis. Notably, rapamycin has been shown to induce IR, glucose intolerance, and diabetes, while still being associated with robust increases in lifespan^380,381^, suggesting it can have the negative effects predicted by the FEM, but may be having other compensatory effects, such as reducing the rate of selective destruction.

By slowing the rate of cell growth/division, rapamycin may reduce either mutation rate or **increase the growth similarity between fast and slow cells** by reducing the rate of fast cell growth toward the level of slower cells (Figure 29A), and therefore reducing the rate of selective destruction. To demonstrate this, we modelled rapamycin to both cause a reduction in AHCM and decelerate the rate of metabolic slowdown (Figure 29B), which would be the primary effect of reduced selective destruction. We modelled rapamycin as both an initial drop (0.15· k_CM_) in k_CM_, but also a reduction in the rate of k_CM_ decline (0.25 times). We chose large effect values to most clearly demonstrate the effects.

**Figure 29.**
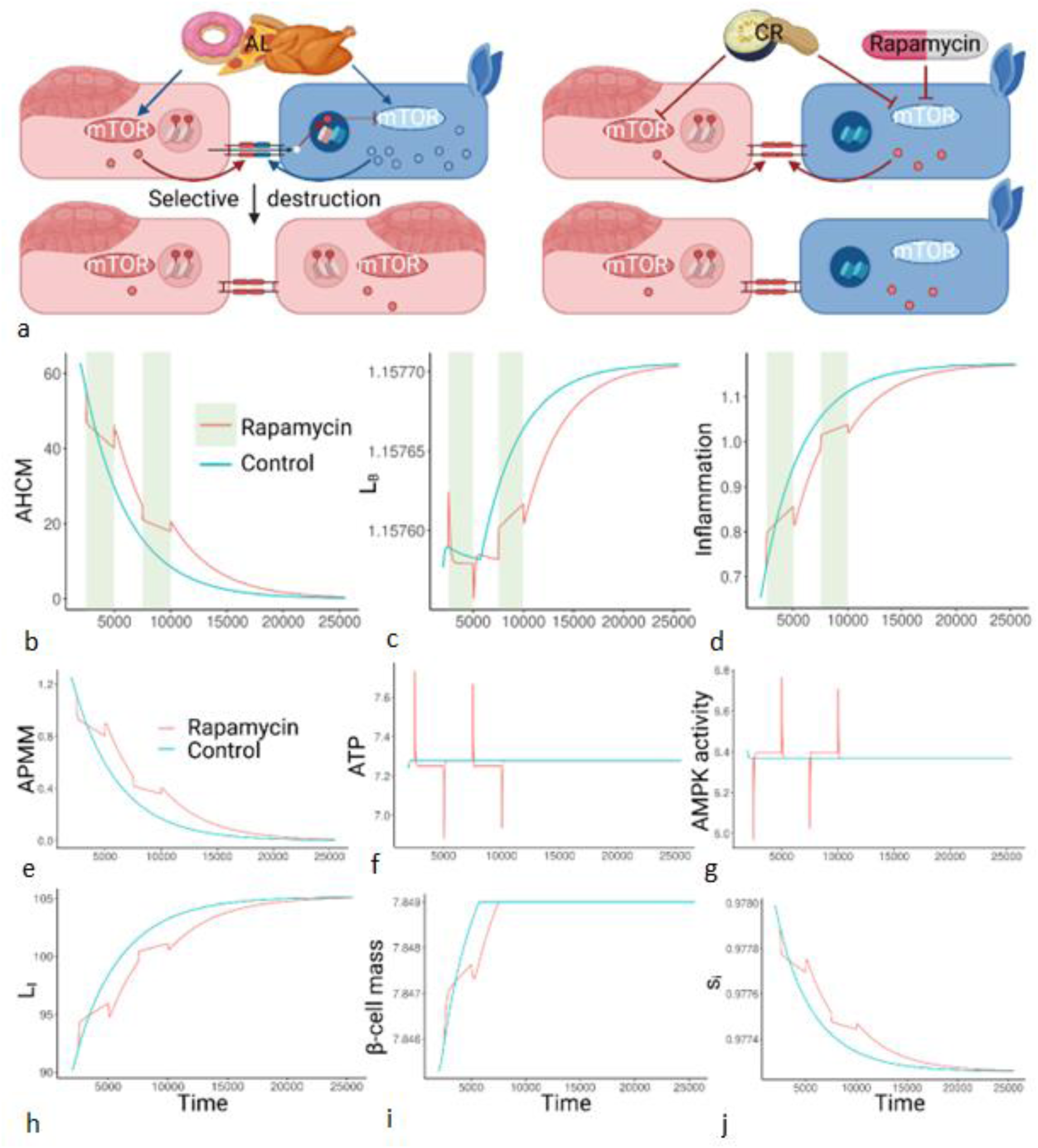
Impact of CR and rapamycin on metabolism. (A) Impact of CR and rapamycin on mTOR signalling and selective destruction (B-J) Effects of rapamycin (represented by changing k_CM_) on: (B) AHCM; (C) L_B_; (D) Inflammation; (E) APMM; (F) ATP; (G) AMPK activity; (H) L_I_; (I) β-cell mass; (J) s_i_. Image created with BioRender.com. Abbreviations, see Table 6.

We compared rapamycin to untreated controls, and consistent with the literature^374^, rapamycin had long lasting beneficial effects on L_B_ and inflammation (Figure 29C-D), as well as many other parameters (Figure 29E-J). However, it also showed that taking rapamycin at the wrong time could cause L_B_ to spike (Figure 29B) and exacerbate inflammation temporarily (Figure 29C), suggesting the drop in AHCM could have detrimental consequences in the short term.

Consistently, rapamycin has transient (paradoxical^382^ negative effects on weight gain and fasting glucose levels at the start of treatment^383^. Schindler, et al.^384^ found that key factors affecting metabolic impairment from rapamycin included the dose and the age of treatment, consistent with the benefit-cost ratio shown in Figure 29B-J.

It should be noted that the FEM was not built to include targets of anti-ageing interventions. The fact that these can be input so easily, and the model responds so realistically, is evidence that the suggested mechanisms are relevant to ageing.

### 9.3 Model discussion

All models are simplifications of reality, and biological models in particular cannot hope to capture the full complexity of the systems they represent. However, here we have tried to demonstrate that homeostatic systems designed to maintain key metabolites at equilibrium will necessarily have knock-on effects leading to increases in either metabolites such as L_I_ or processes such as inflammation (and their molecular constituents). We have not calibrated the model against any specific biological systems, instead validating the model where possible by its consistency with age-related outcomes and interventions. However, in the absence of experimentally derived parameter and variable values this validation is limited to an abstract assessment of model behaviour. We can infer from the model whether components are at equilibrium, increasing, or decreasing, but little more. Fortunately, the purpose of this model was designed to demonstrate which metabolites retain and lose homeostasis under conditions of metabolic slowdown, in the presence or absence of longevity interventions. Further work should include adding organism-specific detail to the model so that it can be properly calibrated to specific systems, but even without these additional steps the FEM indicates that a network designed to maintain equilibrium ATP will have consequences for multiple aspects of fat and energy homeostasis in response to metabolic slowdown.

We suggest that based on the biological evidence available, metabolic slowdown should lead to increasing L_I_, L_B_, basal inflammation, insulin resistance, and mitochondrial dysfunction, likely reflecting the underlying cause of age-related dysfunction and disease. However, the crucial question of whether this outcome is a peculiarity of the model and under what conditions we could expect these outcomes to persist has not been fully addressed. Could organisms with different FEMs escape the consequences of metabolic slowdown, and to what extent need the model be changed to make ageing look radically different?

The central assumption of the model is that the metabolites have systems to maintain equilibrium, ensuring the organism always has the right amount of its key sources of energy to fuel its metabolism/AHCM. To produce this, we borrowed the concept of dynamic compensation.

#### 9.3.1 Metabolite equilibria and dynamic compensation

In the FEM, L_B_, ACoA, and ATP are all maintained at equilibrium despite changes in multiple parameters and variables (see Figure 16). Changing Meal and k_CM_ caused spikes followed by swift return to equilibrium (as long as fuel (L_I_) remained in the adipose). This robustness to changing parameters was termed dynamical compensation by Karin, et al.^195^, who modelled blood glucose. They observed that in the β-cell, insulin, glucose (BIG) model first described by Topp, et al.^134^, changes in blood glucose caused corresponding changes in βM via alterations in the rates of apoptosis and proliferation. By altering the number of insulin producing cells, and the resultant concentration of insulin in the blood, changes in βM could return blood glucose precisely to equilibrium.

As described in the main article, L_B_ is regulated by insulin in a similar way to blood glucose, so we replicated the βM equation^195^ for L_B_ (S3.4). We used the same equation for IMC (S1.9) and its regulation by AMPK, which helped create the robustness of the FEM to maintain L_B_ and ATP at equilibrium despite changes in Meal values or k_CM_ (and most other parameters, as shown in Figure 16). Only the μ, λ, and ε parameters influencing βM and IMC influence the equilibria for L_B_ and ATP respectively.

However, in both cases, the equations S1.9 and S3.4 are unlikely to reflect the real dynamics of L_B_ and ATP as accurately as the equation for βM governing glucose in the BIG model. Glucose is a single, soluble molecule which is carried free in the blood; FFAs include multiple molecules and are not soluble in water, so only a tiny percentage (<0.01%) are carried free in the plasma, with most stored as esters transported on lipoproteins. Thus, while changes in blood glucose are a result of additional molecules entering or leaving the blood and must be counteracted by the opposite change, FFAs can be replenished from sources within the blood such as lipoprotein-bound triglycerides, and different FFAs may be regulated in different ways.

Importantly, the levels of lipoproteins such as LDL also increase with age, as do serum triglycerides, cholesterol, and various markers of L_B_^655,656^, suggesting that it does not have dynamic compensation, at least to the same level as blood glucose which only shows significant changes in the event of diabetes^132,134^. Thus, the regulation of L_B_ is perhaps more similar to the regulation of L_I_, where negative feedback shifts the equilibrium rather than maintains it (Figure 22E).

However, there is also good evidence that at the fundamental level described in the FEM, glucose and L_B_ are regulated in a similar way: glucose increases β-cell survival^657^ and proliferation^658^, as do FFAs at least as far as proliferation^618–620^, although the effect may be more complicated^659^, and some FFAs can inhibit glucose-induced insulin secretion^660^. Whether FFAs protect against β-cell apoptosis is more controversial, with several studies suggesting that FFAs induce β-cell apoptosis^661,662^; however, this (lipo)toxicity effect is also observed for glucose (glucotoxicity), with low levels of glucose protecting β-cells from apoptosis while high levels induce apoptosis^663^, so further investigation is required.

Notably, both glucose and FFAs have been implicated in diabetes and it is also clear that much of the inhibitory effects of FFAs on insulin production reflect prolonged exposure under diabetogenic and obesogenic conditions^662,664^. It is likely that low levels of FFAs protect β-cells from apoptosis, while high levels induce it (just as for glucose), but again it may be complicated by the various types of FFAs. The dynamic compensation for L_B_ may therefore be reduced or removed by factors outside the FEM, or L_B_ may have different effects on insulin production to blood glucose even at the fundamental level, but the question remains of the relevance of dynamic compensation for the conclusions drawn here. We therefore modelled the FEM with constant βM (at different values), as shown in equation S10 and Figure 30A.

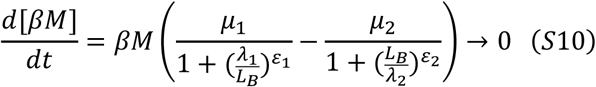

**Figure 30.**
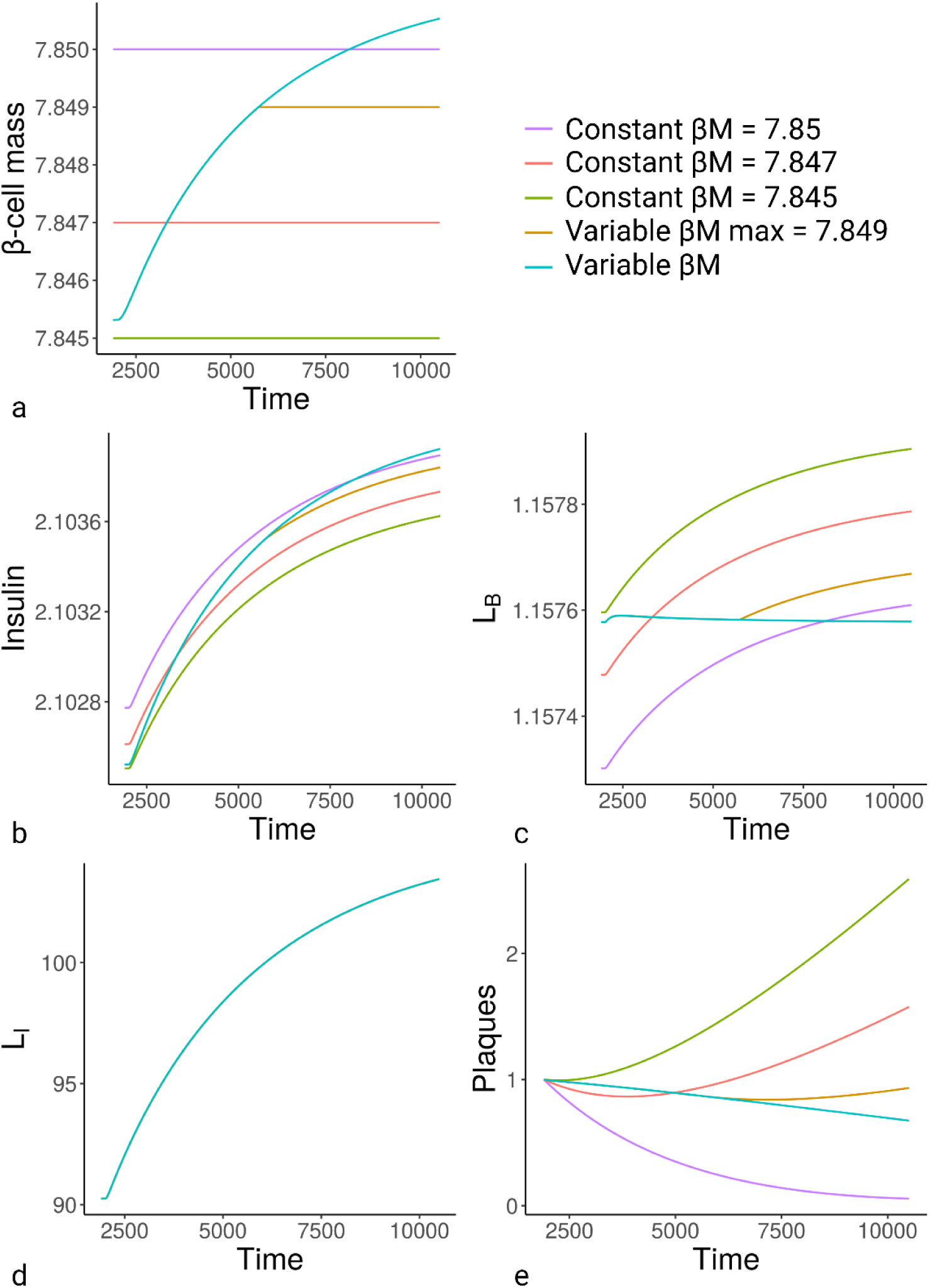
Impact of constant βM on FEM. (A) βM input. (B) Insulin. (C) L_B_. (D) L_I_. (E) Plaques. Abbreviations, see Table 6.

Changes in insulin are shown in Figure 30B. Regardless of the starting value of βM, only a variable βM that adjusted according to insulin requirement allowed dynamic compensation for L_B_ (Figure 30C). However, losing dynamic compensation for L_B_ only accelerated the outcomes predicted by the FEM in response to metabolic slowdown, particularly the rise of L_B_ (Figure 30C) and formation of atherosclerotic plaques (Figure 30E), without affecting L_I_ or basal inflammation (Figure 30D and data not shown). **Thus, the ageing phenotype produced by the FEM is not likely to be a peculiarity of a particular model, but a robust effect of the co-regulation of fuel and energy pathways in response to the continuing unidirectional change of metabolic slowdown**.

Atherosclerosis does appear to occur in young obese individuals^665^, but this may still reflect factors outside the FEM prohibiting the dynamic compensation. Organisms lacking dynamic compensation for L_B_ could expect increasing vascular problems with age – as rising L_B_ would accelerate plaque formation in the blood vessel walls – without much benefit, as L_I_ and its resultant inflammation were largely unaffected (Figure 30D).

Equally, we showed that removal of intrinsic mitochondrial capacity (IMC) from the β-oxidation equation prevented the FEM from maintaining stable levels of ACoA and reduced the stability of ATP (Figure 25A and D), but had no effect on L_I_ or L_B_ (Figure 25B-C). Of course, if IMC does not fluctuate according to ATP and ACoA, then metabolic slowdown will not have detrimental effects on IMC (ie. will not induce mitochondrial dysfunction), but the rest of the age-related phenotypes will happen at the same rate. Essentially, the dynamic compensation for ATP, ACoA, and L_B_ are highly protective behaviours that retard the effects of ageing, and their removal has no positive effects on energy homeostasis that would undermine the conclusions drawn here.

The one case where we did not add dynamic compensation was for L_I_, which could have been done by modelling inflammation as a product of immune cells (IC) and TNF-α using similar equations as βM and insulin, as shown in equations S11 and S12, where IC can expand and contract like βM and IMC (for simplicity we used the existing parameter values for βM in equation S12).

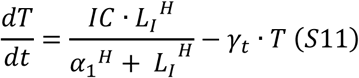

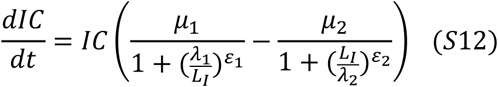

When we altered the FEM with these equations to produce the complete dynamic compensation model (CDC FEM), the results demonstrated that while ATP and ACoA maintained their equilibria (not shown), the equilibrium for L_B_ was lost even in the absence of metabolic slowdown, shown as ‘NegSen’ for negligible senescence in Figure 31. Crucially, while metabolic slowdown (MetSlow) caused the same reduction in AHCM (Figure 31A), β-cell mass (Figure 31B) and insulin (Figure 31C) levels no longer showed the same increase, while insulin sensitivity no longer decreased (Figure 31D). Whereas in the FEM where L_I_ demonstrated a robust rise with age, dependent on metabolic slowdown, in the CDC FEM it was relatively unaffected (Figure 31E), gradually decreasing over time in both NegSen and MetSlow groups.

**Figure 31.**
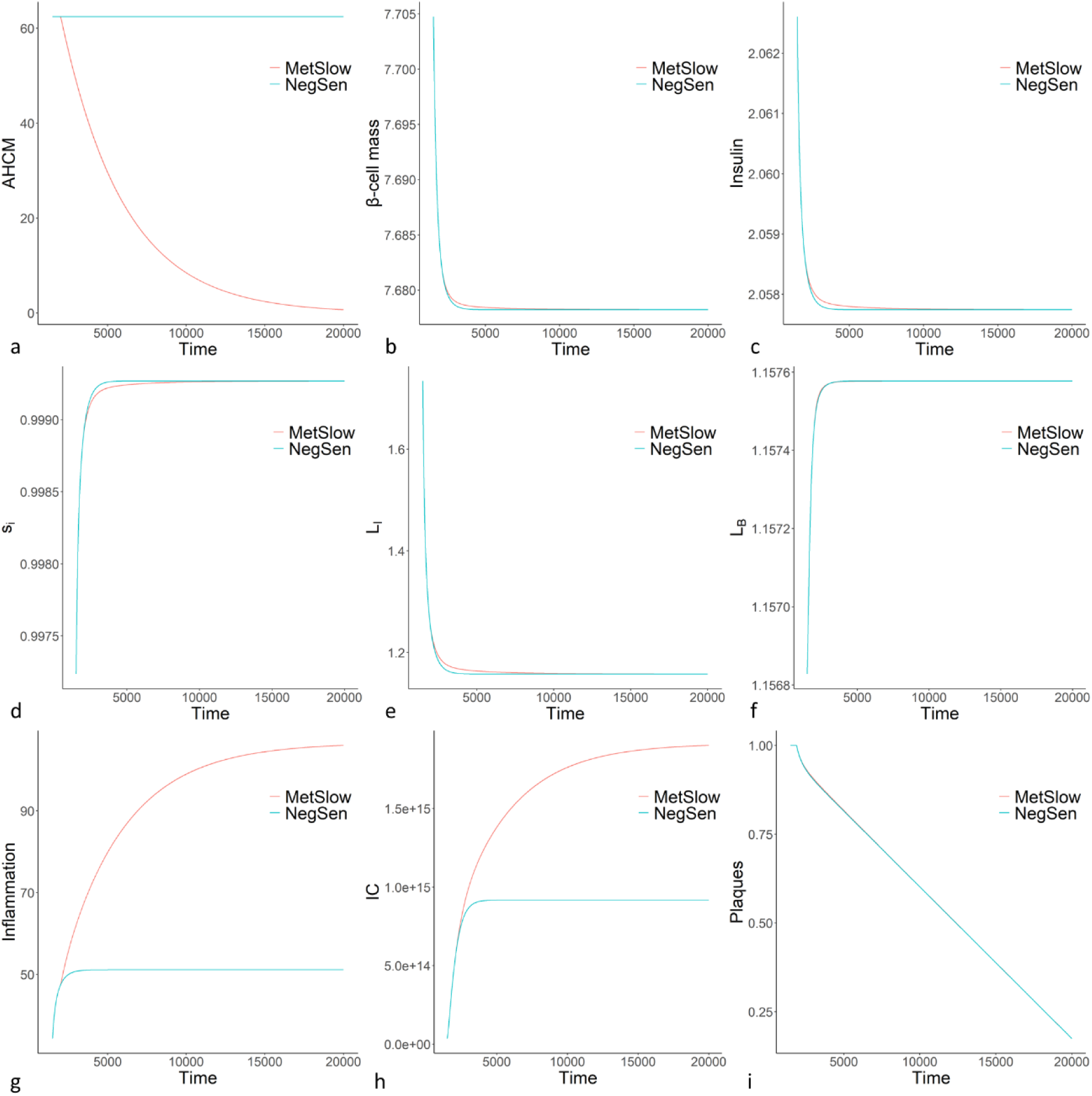
Simulation of the CDC FEM with and without metabolic slowdown (MetSlow and NegSen, respectively). (A) AHCM; (B) β-cell mass; (C) Insulin; (D) insulin sensitivity; (E) L_I_; (F) L_B_; (G) Inflammation; (H) immune cells (I) risk of atherosclerosis. Abbreviations, see Table 6.

Conversely, L_B_ showed a reciprocal gradual rise also independent of metabolic slowdown (Figure 31F). Whereas inflammation (Figure 31G) and inflammatory immune cells (ICs, Figure 31H) demonstrated significant increases dependent on metabolic slowdown, but also rose more gradually without it.

With additional features, the CDC FEM could perhaps return equilibrium to some of the FEM components, but the loss of homeostasis demonstrates the complexity of maintaining equilibrium for every metabolite. By adding an additional control mechanism designed to produce dynamic compensation for L_I_, we have disturbed the equilibrium for L_B_ without gaining it for L_I_. This is good evidence that in the face of a continuous unidirectional change such as metabolic slowdown homeostasis of at least one system must be broken, and L_I_ is surely the least detrimental to bodily function.

While the CDC FEM indicates that weight gain and insulin resistance can be avoided by additional controls for L_I_, the rise in inflammation with age would be substantially higher. Although our current equation for risk, which includes only the level of L_I_ for plaque formation, suggests that risk of death would be lower in the new model (Figure 31), atherosclerosis likely also includes an inflammatory component^666^. Cardiovascular disease would more likely increase than decrease in organisms adopting the CDC FEM to prevent rising body weight.

Indeed, far from producing healthier individuals, the CDC FEM creates organisms that could expect (at least without additional controls) to lose weight over time, even without metabolic slowdown. Their fat reserves would deplete as their adipose tissues became excessively inflamed. Even beyond diseases induced by such changes, these organisms would be at greater risk of energy depletion during times of famine, and would have much more difficulty regulating their key metabolites.

Perhaps this is why there is no evidence for dynamic compensation for L_I_. There is little evidence that the body increases energy expenditure through exercise (see below for heat) to compensate for overfeeding^667^, or for the ‘set-point hypothesis’ that maximum weight is controlled neurologically by the hypothalamus^668^, although it has been suggested that the set-point might be camouflaged by Western diets^197^, specifically refined sugar^669^. Likely, the truth is more complex, reflecting increased fuel burning as L_I_ increases without a set equilibrium, but this additional burning is blunted by Western diets.

Studies examining overfeeding indicate with equal clarity that one excess calorie absorbed does not equal one excess calorie stored^198,670^. In a recent systematic review of 19 studies, Bray and Bouchard^198^ indicated an essentially linear relationship between excess calories in and additional energy stored (energy gain = 3,731 + 0.56 · overfed kcal). At the lower end of excess calories (∼10,000 kcal), 1 excess kcal = 0.93 kcal stored, whereas at higher excesses (∼80,000 kcal), 1 excess kcal = 0.61 kcal stored. While L_I_ is clearly not regulated with the same stringency as ATP and L_B_, these results indicate there are mechanisms to restrain L_I_ increase during positive energy balance, and these mechanisms become increasingly active the higher the excess energy. Notably, it is unlikely to reflect a simple storage incapacity, as extra L_I_ is stored at all intake values.

Another mechanism by which organisms could attempt to thwart the consequences of metabolic slowdown is by regulating L_I_ as is seen in brown adipose tissue (BAT), where mitochondrial uncoupling allows the futile cycling of electrons across the mitochondrial membrane, burning FFAs to produce heat without converting ADP to ATP. Indeed, there is evidence that uncoupling proteins are present in the mitochondria of many cell types, with UCP2 potentially involved in metabolic syndrome, though its significance in regulating weight is unclear ^671^. To consider the effects of uncoupling, we constructed the UC FEM, where uncoupling reduced intrinsic mitochondrial capacity (IMC), which we therefore split into its mass-dependent (MM) and independent parts, the latter now being uncoupling dependent (ignoring other components such as ROS and mitochondrial DNA plasmid quality), determined by the variable UC. MM dynamics could mirror the old IMC in equation S1.9, as shown in equation S13, as the rationale for equation S1.9 was the involvement of mitochondrial mass expanding and contracting in the same way as β-cell mass did in the BIG model.

We modelled UC to mirror equation S3.10 for insulin sensitivity but with L_I_ instead of L_B_, as shown in equation S14, where UC·MM would be substituted for IMC in equation S1 for ATP, as shown in equation S15. However, the APMM in equation S16 would now diverge from the APMM in equation S15 for ATP, as in the latter it reflects OXPHOS (producing ATP) and in the former it is converting ACoA to NADH + H^+^ and FADH_2_, which is Kreb’s cycle. Only the efficiency of OXPHOS is affected by uncoupling, so instead of substituting IMC for MM·UC in equation S2, IMC would be replaced by MM only (which is the same as IMC from before), as shown in equation S16.

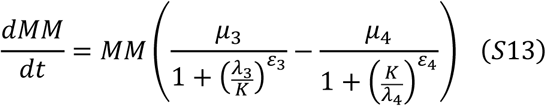

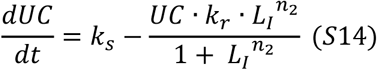

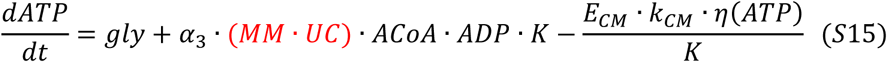

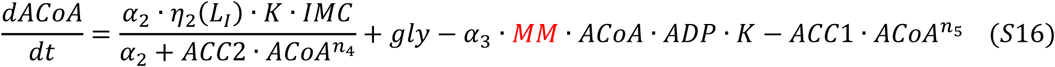

In the UC FEM, the degree of coupling (Figure 32A) determines the amount of productive OXPHOS, which in Figure 32B is the same as total APMM in the FEM. As shown in Figure 32B, the total APMM in the UC FEM exceeds that in the FEM. This ‘total APMM’ refers to the maximal rate of both unproductive and productive OXPHOS, which in the UC FEM is also the rate of Kreb’s cycle. The differences between the rate of Kreb’s and productive OXPHOS reflects the rate that L_I_ is burned to remove excess L_I_ rather than create ATP, the level of which is unaffected by uncoupling (Figure 32C). Uncoupling clearly offers several advantages for L_I_ regulation as it decelerates the rise in L_I_ during metabolic slowdown (Figure 32D), the degree of coupling rising as L_I_ increases (Figure 32A), which slows the increase in chronic inflammation (Figure 32E) and the decline in insulin sensitivity (Figure 32F). However, the UC FEM does not undermine the fundamental conclusions of the FEM as it can only slow but not prevent the phenotypes of weight gain, insulin resistance, and chronic inflammation that induce age-related decline and disease.

**Figure 32.**
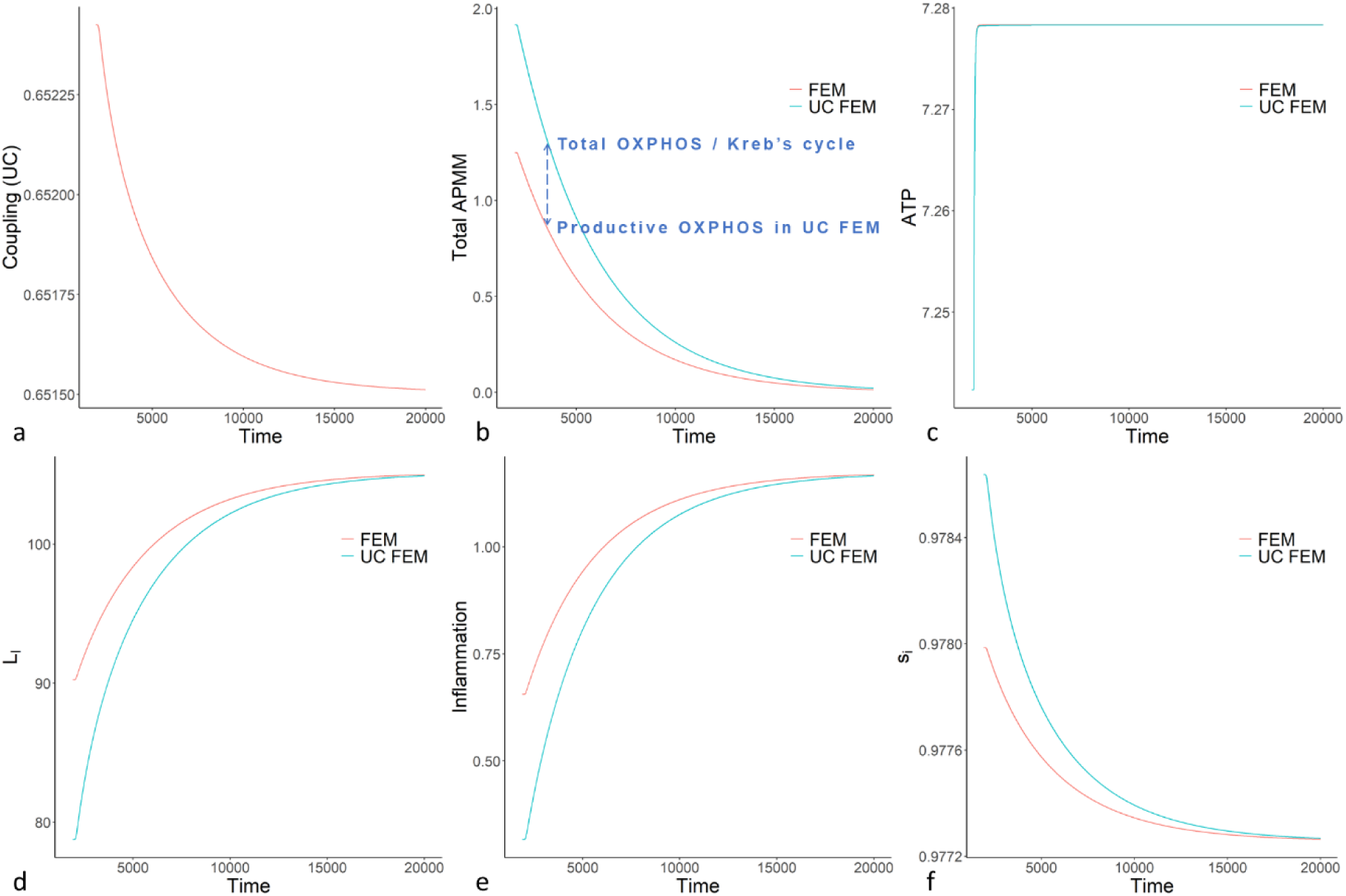
Comparison of key variables between UC FEM and FEM. (A) Fraction of coupled mitochondrial metabolism in UC FEM. (B) Total APMM as defined by the rate of productive and unproductive OXPHOS, also the rate of Kreb’s cycle. In the FEM there is no unproductive OXPHOS, and the rate of OXPHOS is equal to the rate of productive APMM in the UC FEM (as indicated in blue). (C-F) key variables compared between UC FEM and FEM: (C) ATP; (D) L_I_; (E) inflammation; (F) insulin sensitivity. Abbreviations, see Table 6.

Notably, the main purpose of uncoupling appears to be heat generation rather than burning excess calories, as it rises during cold stimulation dependent on the presence of BAT^201^, was higher under ambiently colder conditions^200^, and most importantly was blunted by obesity^202^, which is the opposite to what you would expect if it was a major pathway for the removal of excess fat. Equally, clinical studies examining why excess calories led to smaller gains in energy and fat mass than expected (metabolic inefficiency), found little evidence for an increase in basal metabolic rate (or physical activity) expected by an increase in thermogenesis^672^. Although others suggest “thermogenic activity offers tremendous potential to combat obesity”^673^, increasing in response to high calorie diets to provide obesity resistance^674,675^. Why evolution would not naturally produce this mechanism of slowing the effects of ageing could be a direct trade-off of the kind predicted by disposable soma theory, as any attempt to regulate uncoupling to control L_I_ would lead to increased fat wastage. It is possible that organisms with the fitness of advantage of reduced risk age-related death would have a greater fitness cost in terms of risk of starvation.

Whether or not our bodies naturally regulate uncoupling for the purpose of reducing stored fat, the UC FEM suggests that while uncoupling will not prevent ageing, therapies designed to waste ATP will reduce the effects of metabolic slowdown at all levels, greatly retarding the effects of ageing on mitochondrial function, weight gain, insulin sensitivity and inflammation. As the evidence suggests, the UC FEM predicts that uncouplers should also be highly effective weight loss drugs and effective therapies for metabolic syndrome^671,676^.

The other crucial system the body employs to remove fat is reverse cholesterol transport (RCT), where both cholesterol and FFAs are secreted into the bile and removed via the colon. However, as the UC FEM demonstrates, the body could have multiple such pathways, but unless they have dynamic compensation, they will only delay the effects of metabolic slowdown. The problem arises because one metabolite must be allowed to vary in response metabolic slowdown, as shown in the CDC FEM. If the body is to keep ATP, ACoA, L_B_, and L_I_ at equilibrium, it would require a further metabolite to change. For example, the body could convert the excess FFAs to other lipids, such as cholesterol or ceramides. Indeed, the rise in these problematic lipids with age^677^ may well reflect this shift.

Contrary to L_I_, there is good evidence for a robust ATP equilibrium that does not change with age, as discussed in the model construction. In the FEM, this is maintained by the regulation of IMC by AMPK as in equation S1.9. As AMPK is a function of ATP:AMP ratio, this maintains equilibrium ATP via the same mechanism as the βM equation S3.4 does for L_B_: when ATP decreases, IMC increases to elevate ATP production and restore ATP; and when ATP increases, the IMC decreases reducing ATP production and restoring ATP. However, it should be noted that while the ATP levels remain constant, the size of the adenosine phosphate (AP) pool increases with age (and ratio to AMP decreases)^173^, suggesting that our equations S1.5-1.7 are oversimplifications.

The main oversimplification for ATP dynamics is grouping all the factors affecting intrinsic mitochondrial capacity together into a single variable, intrinsic mitochondrial capacity (IMC). Although the UC FEM splits IMC into mitochondrial mass and the degree of uncoupling, there are multiple other factors which regulate the rate of OXPHOS, Kreb’s, and glycolysis to ensure metabolites remain at equilibrium. Glycolysis and its regulation by hypoxia-inducible factor 1alpha (HIF1α) may be highly significant in ageing and cancer^678^, but is not considered here as we have focussed on lipid dynamics.

#### 9.3.2 Additional model limitations

A key function of the adipose tissue is exporting lipid back into the blood during negative energy balance, allowing non-adipose tissues to maintain ATP production via β-oxidation. In equation S3.5, export is regulated entirely via insulin inhibition, but in reality there are many additional regulating factors. Adipose tissue is one of the few tissues along with the liver that responds to glucagon, stimulating lipolysis in rats^679^ and the release of FFAs and glycerol into the blood^680^ in a mechanism that is inhibited by insulin^681,682^. However, the relationship is likely more complicated in human adipose than rodents^683^, and some studies indicate glucagon may actually reduce blood lipid^684,685^. Clinical studies showed no effect of glucagon on adipose lipolysis in people with and without diabetes^686^, and expression of white adipose tissue (WAT) glucagon receptor did not affect lipolysis or hepatic lipid accumulation in mice^687^. Hence, given the uncertainty, we did not include glucagon, but the implications of a second major regulatory hormone are worth further consideration. If glucose and lipid levels already have dynamic compensation with only insulin, what is the purpose of its direct counter-regulator? Is it merely fine tuning, speed, or redundancy, or are there additional dynamic considerations? There is also evidence for roles of somatostatin and adrenaline in regulating lipolysis and export, but due to the complexities of these regulatory systems, and the likelihood that they regulate export in response to conditions outside the FEM, such as stress, we chose not to include them. However, understanding the significance of glucagon and other hormones in adipose homeostasis may help ameliorate the impact of metabolic slowdown on lipid accumulation.

Another simplification of the FEM regarding L_I_ was described in equations S3.6-3.8. As we reduced the model to include only the adipose and the blood, the model did not respond to metabolic slowdown with compensatory changes to import and export (Figure 22L-N) as it did to changes in Meal value (Figure 22Z-iii). Metabolic slowdown affected only the adipose rate of lipid β-oxidation without impacting additional tissues. The latter would have altered rates of non-adipose import, necessitating compensatory changes in adipose import and export to maintain L_B_. Further development of the model should include a multi-tissue system.

Probably the greatest oversimplification employed in the FEM is the simulation of inflammation as a single cytokine, TNF-α, which is produced only by hypertrophic adipocytes. It would be impossible to include a realistic model of the immune system without it dominating all other components. However, further work should include factors such as hypertrophic adipocytes secreting monocyte chemoattractant protein-1 (MCP-1) and granulocyte colony stimulating factor (G-CSF)^688^, both of which attract immune cells, and the latter mobilizes haematopoietic stem cells for the production and release of neutrophils from the bone marrow^689,690^, expanding the pool of monocytes and macrophages, and enhancing their phagocytic function^691,692^. The incorporation of immune cells and their relationship with adipocytes would provide further understanding of the role of inflammation in adipose homeostasis, but unlike many other studies which interpret inflammation as largely destructive or of unknown function, the FEM suggests a clear function for inflammation in removal of excess fat, insulin sensitization, and the resultant maintenance of lipid homeostasis.

The danger with any model of inflammation is that it spirals out of control, creating a vicious cycle that is purely destructive. This is frequently how inflammation is viewed in gerontology, but inflammation has a series of robust negative feedback mechanisms that largely prevent these vicious cycles. The FEM suggests that rising basal inflammation reflects a mixture of mitochondrial dysfunction and lipid dysfunction, and need not result from damage accumulation or allostatic load^49,693^. However, these causes are not exclusive with each other.

One seemingly vicious cycle we did not include in the FEM is the link between inflammation and insulin resistance, which is so strong that there is great uncertainty as to which is cause and which is effect. A role for IR in inducing inflammation is consistent with inflammation as a mechanism of removal of insulin resistant hypertrophic cells as predicted by the FEM. Thus, evidence such as the inhibition of mTORC2 and resultant IR inducing increased polarisation to inflammatory M1 macrophages through MCP-1^111^ is not surprising. However, that inflammation induces IR is counterintuitive to the mechanisms proposed here.

Macrophage infiltration and activation has been shown to induce insulin resistance in adipose tissue and in vitro adipocytes^694,695^. Equally, knockout of IκB kinase-beta (IKK-β) or c-Jun N-terminal kinase 1 (JNK1) involved in macrophage signalling protected mice from diet-induced insulin resistance^110,696,697^. Importantly, in many of these studies, genetic changes in the macrophages were capable of inducing IR in adipocytes, suggesting the IR was not just a byproduct of obesity which also induced inflammation; the inflammation caused the IR.

It should be noted that many of the conditions demonstrating inflammation induces IR were heavily biased against the insulin-sensitizing effects of inflammation, such as obesity or in vitro conditions with high numbers of immune cells. It is likely that if the macrophages were in a healthy, non-obese tissue where resolution of the excess L_I_ could occur, they would deactivate and exfiltrate, promoting the anti-inflammatory polarisation we have discussed above where dead adipocytes are replaced by new insulin sensitive ones. However, in obesity and in vitro, resolution can be prevented by multiple conditions including immune cells at supraphysiological levels or constitutive activity due to mutation, or the absence of the stem cells to replace insulin resistant adipocytes with new insulin sensitive ones. Under such conditions several factors may raise the level of immune activation so that it is more resembling of an immune response to infection rather than basal inflammation to remove hypertrophic cells.

Crucially, just as the body benefits from reducing ATP production in infected cells, by prohibiting pathogen replication, it also benefits from reducing import of fuel into infected cells, which could instead be absorbed by the immune cells combatting the infection, rather than feeding pathogen infected cells that may aid pathogenic replication. It therefore makes evolutionary sense that higher levels of inflammation consistent with infection – which may occur in obese individuals where the mounting immune response is insufficient to clear the excess L_I_ (and thus deactivate) – induce IR.

#### 9.3.3 Dependency of conclusions on model dynamics

In several equations we employed non-linear dynamics because we believed this was a better reflection of the biology. To demonstrate whether these affected the conclusions drawn, we replaced all the non-linear model functions to produce the linear FEM (LFEM). We did not alter the equations of IMC and βM as we have already addressed the significance of dynamic compensation above. All exponent parameters were removed from the equations, and the sigmoid functions in the equations for s_i_ and T were removed and replaced with the simplified versions shown in equations S17 and S18.

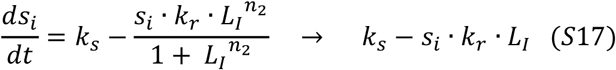

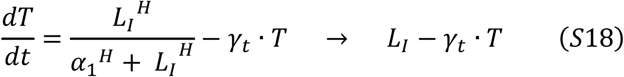

No parameter values needed to be changed for the LFEM to produce the same results as the FEM. Although the metabolites and processes had very different values, their homeostatic profiles remained the same. As AHCM declined (Figure 33A), so did APMM and IMC (Figure 33B and C). ATP remained constant (Figure 33D), L_B_ slowly returned to equilibrium (Figure 33E), maintained by increasing βM (Figure 33F) and insulin (not shown). Insulin sensitivity declined (Figure 33G), and L_I_ slowly rose over time (Figure 33H), as did compensatory inflammation (Figure 33I). The only metabolite that showed significant difference between the LFEM and FEM was ACoA, which slowly declined across the time course (Figure 33I). As it was unclear whether this reflected a slower return to equilibrium, we extended the time course from 20,000 au to 1,000,000 au. However, the results suggested that the LFEM did not maintain ACoA at equilibrium (Figure 33J), levels continually decreasing across the extended timecourse below the level of ACoA when metabolic slowdown began (Figure 33K). We predicted that this reflected the loss of strong negative feedback created by the exponents in equation S2 for ACoA. We therefore readded these exponents (as well as the same ones in equation S4 for L_I_) to the LFEM and replotted ACoA. As predicted, the strong negative feedback restored the ACoA equilibrium (Figure 33L).

**Figure 33.**
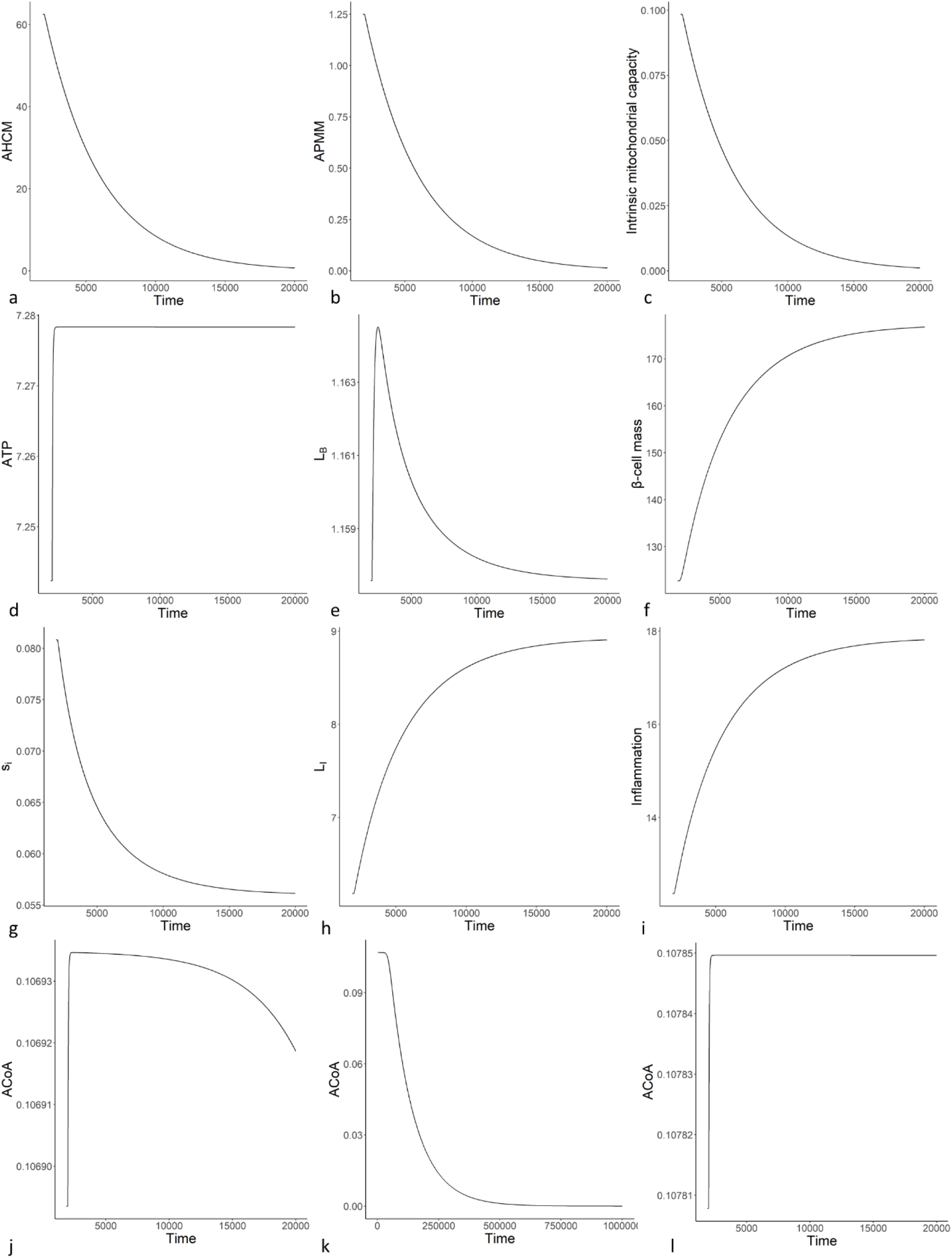
Simulations of key variables in the LFEM demonstrating metabolic slowdown induces the same effects as in the main model (FEM). (A) AHCM; (B) APMM; (C) IMC; (D) ATP; (E) LB; (F) β-cell mass; (G) insulin sensitivity; (H) LI; (I) Inflammation; (J) ACoA; (K) ACoA over extended time course; (L) ACoA from LFEM with FEM equation S2 for ACoA. Abbreviations, see Table 6.

The LFEM suggests the robustness of the FEM to maintain important metabolites at equilibrium, and suggests the results are not a peculiarity of non-linear dynamics or specific formulations of the equations. To invalidate the conclusions drawn from the FEM would require demonstration that the relationship between the processes and molecules is fundamentally different to the formulations created here by (non-systematic) literature review, or the addition of more components capable of reversing the relationships produced here in the context of additional complexity. This is strong evidence that metabolic slowdown alone could be sufficient to explain much of the ageing phenotype, age-related disease, and the central anti-ageing therapies as we understand them. If AHCM declines with age, the resulting metabolic cascade may be unavoidable, resulting in dysfunction, disease, and death.

## Model abbreviations table

## Glossary

In this paper we have invented several terms that are not defined elsewhere. We have also used some standard words in non-standard ways to avoid having to invent further words. All such terms are described below.

**Anabolism** – All processes requiring the use of bioavailable energy to carry out cell functions.

**Antagonistic pleiotropy** – The dual effects inducing both benefit and detriment to organisms. Initially Williams defined it to refer only to genes; however, it is now widely used to refer to any effects from genes, molecules, processes or pathways that are both beneficial and detrimental in different contexts.

**Bioavailable energy** – energy stored in molecules such as ATP which are immediately available for cell functions without first requiring to be broken down by catabolism.

**Catabolism** – all processes which break down sugar, lipid, or protein molecules toward producing acetyl-CoA or ATP.

**Celerisis** – the outcome of the natural course of tissue-level selection which results in the spread of faster cells to cause hyperfunction and associated diseases.

**Cell senescence** – this term has undergone significant concept creep, but here we define it as the state of permanent arrest induced by damage or dysfunction (either autonomously or otherwise).

**Disposable soma theory (DST)** – the theory put forward by Tom Kirkwood that organisms evolve to accumulate damage by reducing energy invested in somatic maintenance. As the energy could be invested in other processes such as growth and reproduction, ageing via damage accumulation would provide a selective advantage over negligible senescence when it produced more (viable and robust) offspring.

**Extrinsic ageing** – The shift in energy homeostasis resulting from metabolic slowdown which induces mitochondrial dysfunction, metabolic syndrome, inflammageing, and the associated diseases.

**Fast/slow cell** – cells with altered metabolic rate. **Hyperfunction** – overactivity with detrimental effects. **Hypofunction** – underactivity with detrimental effects.

**Inflammageing** – any process by which inflammation contributes to the development of age-related disease.

**Inflammageing hypothesis/model** – the specific hypothesis or group of hypotheses by which DNA damage accumulation leads to cycles of positive feedback with inflammation which lead to age-related disease.

**Intrinsic ageing** – the induction of hypofunction and associated processes and diseases that results from selective destruction and metabolic slowdown.

**Juxtacrine** – between neighbouring and connected cells.

**Longevity module** – the collection of energy regulating pathways and processes containing the molecular components which regulate ageing and longevity.

**Metabolic slowdown** – the gradual reduction in metabolic rate of cells and tissues with ageing.

**Mitopenia** – The spread of low ROS, depolarised mitochondria resulting from the process of mitochondrial selection.

**Natural course of selection** – the likely or even unavoidable outcome of selection within tissues or cells that results from mutants which have a selective advantage under normal tissue conditions in the absence of specifically evolved processes to induce counterselection and prevent their spread.

**Selective destruction** – any process of communication which either kills, removes, or otherwise neutralises the advantage of agents that prevents or reverses the natural course of selection.

**TOR-centric quasiprogramme (TCQP)** – The programme proposed by Mikhail Blagosklonny, regulated by mTOR, which defined ageing as the continuation of the developmental process once development was complete, inducing unwanted and detrimental growth, or hyperfunction. The functional programme thus becomes a functionless and even detrimental quasi-programme.

## Author contributions

James Wordsworth developed the theory and model, and wrote the manuscript with input and supervision from Daryl Shanley. Pernille Yde Nielsen helped with coding, modelling, and manuscript development, as well as contributing ideas. Edward Fielder and Sharmilla Chandrasegaran helped with research and contributed to manuscript development.

## Acknowledgements

We would like to thank Professor Tom Kirkwood and Dr Viktor Korolchuk for guidance and input. Images were created using BioRender.com. This work was supported partly by the Novo Nordisk Fonden Challenge Programme: Harnessing the Power of Big Data to Address the Societal Challenge of Aging (grant number NNF17OC0027812) and the NIHR Newcastle Biomedical Research Centre (BRC).

## References

1 Wordsworth, J., H, O. K., Clark, P. & Shanley, D. The damage-independent evolution of ageing by selective destruction. Mech Ageing Dev 207, 111709, doi:10.1016/j.mad.2022.111709 (2022).

2 Wordsworth, J., Nielsen, P. Y., Fielder, E., Chandrasegaran, S. & Shanley, D. Metabolic slowdown as the proximal cause of ageing and death. bioRxiv, 2023.2008.2001.551537, doi:10.1101/2023.08.01.551537 (2023).

3 Gems, D., Okholm, S. & Lemoine, M. Inflated expectations: the strange craze for translational research on aging: Given existing confusion about the basic science of aging, why the high optimism in the private sector about the prospects of developing anti-aging treatments? EMBO Rep 25, 3748–3752, doi:10.1038/s44319-024-00226-2 (2024).

4 Mitteldorf, J. Aging Is Not a Process of Wear and Tear. Rejuvenation Research 13, 322–326, doi:10.1089/rej.2009.0967 (2010).

5 Kirkwood, T. B. L. Evolution of ageing. Nature 270, 301–304, doi:10.1038/270301a0 (1977).

6 Kirkwood, T. B. Understanding the odd science of aging. Cell 120, 437–447, doi:10.1016/j.cell.2005.01.027 (2005).

7 Averill-Bates, D. Reactive oxygen species and cell signaling. Review. Biochim Biophys Acta Mol Cell Res 1871, 119573, doi:10.1016/j.bbamcr.2023.119573 (2024).

8 Sardina, J. L. et al. p22phox-dependent NADPH oxidase activity is required for megakaryocytic differentiation. Cell Death & Differentiation 17, 1842–1854, doi:10.1038/cdd.2010.67 (2010).

9 Guzy, R. D. & Schumacker, P. T. Oxygen sensing by mitochondria at complex III: the paradox of increased reactive oxygen species during hypoxia. Exp Physiol 91, 807–819, doi:10.1113/expphysiol.2006.033506 (2006).

10 Bjelakovic, G., Nikolova, D., Gluud, L. L., Simonetti, R. G. & Gluud, C. Antioxidant supplements for prevention of mortality in healthy participants and patients with various diseases. Cochrane Database Syst Rev 2012, Cd007176, doi:10.1002/14651858.CD007176.pub2 (2012).

11 Halsne, R. et al. Lack of the DNA glycosylases MYH and OGG1 in the cancer prone double mutant mouse does not increase mitochondrial DNA mutagenesis. DNA repair 11, 278–285 (2012).

12 Itsara, L. S. et al. Oxidative stress is not a major contributor to somatic mitochondrial DNA mutations. PLoS genetics 10, e1003974 (2014).

13 Sanchez-Contreras, M. et al. The multi-tissue landscape of somatic mtDNA mutations indicates tissue-specific accumulation and removal in aging. eLife 12, e83395, doi:10.7554/eLife.83395 (2023).

14 Lapointe, J. & Hekimi, S. When a theory of aging ages badly. Cell Mol Life Sci 67, 1–8, doi:10.1007/s00018-009-0138-8 (2010).

15 Gladyshev, V. N. The free radical theory of aging is dead. Long live the damage theory! Antioxid Redox Signal 20, 727–731, doi:10.1089/ars.2013.5228 (2014).

16 Gems, D. & McElwee, J. J. Broad spectrum detoxification: the major longevity assurance process regulated by insulin/IGF-1 signaling? Mechanisms of Ageing and Development 126, 381–387, 10.1016/j.mad.2004.09.001 (2005).

17 Vijg, J. From DNA damage to mutations: All roads lead to aging. Ageing Res Rev 68, 101316, 10.1016/j.arr.2021.101316 (2021).

18 Moore, L. et al. The mutational landscape of human somatic and germline cells. Nature 597, 381–386, doi:10.1038/s41586-021-03822-7 (2021).

19 Olovnikov, A. M. Telomeres, telomerase, and aging: origin of the theory. Exp Gerontol 31, 443–448, doi:10.1016/0531-5565(96)00005-8 (1996).

20 Seluanov, A. et al. Telomerase activity coevolves with body mass not lifespan. Aging cell 6, 45–52 (2007).

21 Hayflick, L. & Moorhead, P. S. The serial cultivation of human diploid cell strains. Exp Cell Res 25, 585–621, doi:10.1016/0014-4827(61)90192-6 (1961).

22 Bodnar, A. G. et al. Extension of life-span by introduction of telomerase into normal human cells. Science 279, 349–352, doi:10.1126/science.279.5349.349 (1998).

23 Vaiserman, A. & Krasnienkov, D. Telomere Length as a Marker of Biological Age: State-of-the-Art, Open Issues, and Future Perspectives. Frontiers in Genetics Volume 11-2020, doi:10.3389/fgene.2020.630186 (2021).

24 Sanders, J. L. & Newman, A. B. Telomere Length in Epidemiology: A Biomarker of Aging, Age-Related Disease, Both, or Neither? Epidemiologic Reviews 35, 112–131, doi:10.1093/epirev/mxs008 (2013).

25 Ye, Q. et al. Telomere length and chronological age across the human lifespan: A systematic review and meta-analysis of 414 study samples including 743,019 individuals. Ageing Res Rev 90, 102031, doi:10.1016/j.arr.2023.102031 (2023).

26 Prowse, K. R. & Greider, C. W. Developmental and tissue-specific regulation of mouse telomerase and telomere length. Proceedings of the National Academy of Sciences 92, 4818–4822, doi:doi:10.1073/pnas.92.11.4818 (1995).

27 Haussmann, M. F. & Mauck, R. A. Telomeres and Longevity: Testing an Evolutionary Hypothesis. Molecular Biology and Evolution 25, 220–228, doi:10.1093/molbev/msm244 (2007).

28 Rudolph, K. L. et al. Longevity, Stress Response, and Cancer in Aging Telomerase-Deficient Mice. Cell 96, 701–712, doi:10.1016/S0092-8674(00)80580-2 (1999).

29 Blasco, M. A. et al. Telomere shortening and tumor formation by mouse cells lacking telomerase RNA. Cell 91, 25–34, doi:10.1016/s0092-8674(01)80006-4 (1997).

30 Kuo, C. L., Pilling, L. C., Kuchel, G. A., Ferrucci, L. & Melzer, D. Telomere length and aging-related outcomes in humans: A Mendelian randomization study in 261,000 older participants. Aging Cell 18, e13017, doi:10.1111/acel.13017 (2019).

31 Allsopp, R. C. et al. Telomere length predicts replicative capacity of human fibroblasts. Proc Natl Acad Sci U S A 89, 10114–10118, doi:10.1073/pnas.89.21.10114 (1992).

32 Toussaint, O., Medrano, E. E. & von Zglinicki, T. Cellular and molecular mechanisms of stress-induced premature senescence (SIPS) of human diploid fibroblasts and melanocytes. Exp Gerontol 35, 927–945, doi:10.1016/s0531-5565(00)00180-7 (2000).

33 Dimri, G. P. et al. A biomarker that identifies senescent human cells in culture and in aging skin in vivo. Proc Natl Acad Sci U S A 92, 9363–9367, doi:10.1073/pnas.92.20.9363 (1995).

34 Baker, D. J. et al. Clearance of p16Ink4a-positive senescent cells delays ageing-associated disorders. Nature 479, 232–236, doi:10.1038/nature10600 (2011).

35 Harrison, D. E. et al. Astaxanthin and meclizine extend lifespan in UM-HET3 male mice; fisetin, SG1002 (hydrogen sulfide donor), dimethyl fumarate, mycophenolic acid, and 4-phenylbutyrate do not significantly affect lifespan in either sex at the doses and schedules used. Geroscience 46, 795–816, doi:10.1007/s11357-023-01011-0 (2024).

36 Wang, B. et al. Intermittent clearance of p21-highly-expressing cells extends lifespan and confers sustained benefits to health and physical function. Cell Metab 36, 1795–1805.e1796, doi:10.1016/j.cmet.2024.07.006 (2024).

37 Raffaele, M. & Vinciguerra, M. The costs and benefits of senotherapeutics for human health. The Lancet Healthy Longevity 3, e67–e77, doi:10.1016/S2666-7568(21)00300-7 (2022).

38 Kowald, A. & Kirkwood, T. B. L. Senolytics and the compression of late-life mortality. Exp Gerontol 155, 111588, doi:10.1016/j.exger.2021.111588 (2021).

39 López-Otín, C., Blasco, M. A., Partridge, L., Serrano, M. & Kroemer, G. The hallmarks of aging. Cell 153, 1194–1217, doi:10.1016/j.cell.2013.05.039 (2013).

40 Szilard, L. On the nature of the aging process. Proceedings of the National Academy of Sciences 45, 30–45 (1959).

41 Bartz, J., Jung, H., Wasiluk, K., Zhang, L. & Dong, X. Progress in Discovering Transcriptional Noise in Aging. Int J Mol Sci 24, doi:10.3390/ijms24043701 (2023).

42 Martinez-Jimenez, C. P. et al. Aging increases cell-to-cell transcriptional variability upon immune stimulation. Science 355, 1433–1436, doi:10.1126/science.aah4115 (2017).

43 Maynard Smith, J. (Nature Publishing Group UK London, 1959).

44 Clark, A. M. & Rubin, M. A. The modification by X-irradiation of the life span of haploids and diploids of the wasp, Habrobracon sp. Radiation Research 15, 244–253 (1961).

45 Morley, A. A. Is ageing the result of dominant and co-dominant mutations? Journal of Theoretical Biology 98, 469–474 (1982).

46 Vijg, J. Somatic mutations and aging: a re-evaluation. Mutation Research/Fundamental and Molecular Mechanisms of Mutagenesis 447, 117–135, 10.1016/S0027-5107(99)00202-X (2000).

47 Giunta, B. et al. Inflammaging as a prodrome to Alzheimer’s disease. Journal of Neuroinflammation 5, 51, doi:10.1186/1742-2094-5-51 (2008).

48 Franceschi, C. et al. Inflamm-aging: an evolutionary perspective on immunosenescence. Annals of the new York Academy of Sciences 908, 244–254 (2000).

49 Franceschi, C. & Campisi, J. Chronic Inflammation (Inflammaging) and Its Potential Contribution to Age-Associated Diseases. The Journals of Gerontology: Series A 69, S4–S9, doi:10.1093/gerona/glu057 (2014).

50 Higami, Y. & Shimokawa, I. Apoptosis in the aging process. Cell and Tissue Research 301, 125–132, doi:10.1007/s004419900156 (2000).

51 Campisi, J. & Warner, H. R. in Advances in Cell Aging and Gerontology Vol. 4 1-16 (Elsevier, 2001).

52 Giangreco, A., Qin, M., Pintar, J. E. & Watt, F. M. Epidermal stem cells are retained in vivo throughout skin aging. Aging Cell 7, 250–259, doi:10.1111/j.1474-9726.2008.00372.x (2008).

53 Pang, W. W. et al. Human bone marrow hematopoietic stem cells are increased in frequency and myeloid-biased with age. Proc Natl Acad Sci U S A 108, 20012–20017, doi:10.1073/pnas.1116110108 (2011).

54 Su, T.-Y. et al. Aging is associated with functional and molecular changes in distinct hematopoietic stem cell subsets. Nature Communications 15, 7966, doi:10.1038/s41467-024-52318-1 (2024).

55 Beerman, I., Maloney, W. J., Weissmann, I. L. & Rossi, D. J. Stem cells and the aging hematopoietic system. Curr Opin Immunol 22, 500–506, doi:10.1016/j.coi.2010.06.007 (2010).

56 Kavathia, N., Jain, A., Walston, J., Beamer, B. A. & Fedarko, N. S. Serum markers of apoptosis decrease with age and cancer stage. Aging (Albany NY) 1, 652–663, doi:10.18632/aging.100069 (2009).

57 Suh, Y. et al. Aging alters the apoptotic response to genotoxic stress. Nature Medicine 8, 3–4, doi:10.1038/nm0102-3 (2002).

58 Salminen, A., Ojala, J. & Kaarniranta, K. Apoptosis and aging: increased resistance to apoptosis enhances the aging process. Cellular and Molecular Life Sciences 68, 1021–1031, doi:10.1007/s00018-010-0597-y (2011).

59 Herndon, L. A. et al. Stochastic and genetic factors influence tissue-specific decline in ageing C. elegans. Nature 419, 808–814, doi:10.1038/nature01135 (2002).

60 Nelson, G. et al. A senescent cell bystander effect: senescence-induced senescence. Aging Cell 11, 345–349, doi:10.1111/j.1474-9726.2012.00795.x (2012).

61 Acosta, J. C. et al. A complex secretory program orchestrated by the inflammasome controls paracrine senescence. Nat Cell Biol 15, 978–990, doi:10.1038/ncb2784 (2013).

62 Hoare, M. et al. NOTCH1 mediates a switch between two distinct secretomes during senescence. Nat Cell Biol 18, 979–992, doi:10.1038/ncb3397 (2016).

63 Kowald, A., Passos, J. F. & Kirkwood, T. B. L. On the evolution of cellular senescence. Aging Cell n/a, e13270, 10.1111/acel.13270 (2020).

64 Van Raamsdonk, J. M. & Hekimi, S. Deletion of the mitochondrial superoxide dismutase sod-2 extends lifespan in Caenorhabditis elegans. PLoS Genet 5, e1000361, doi:10.1371/journal.pgen.1000361 (2009).

65 Van Raamsdonk, J. M. & Hekimi, S. Superoxide dismutase is dispensable for normal animal lifespan. Proc Natl Acad Sci U S A 109, 5785–5790, doi:10.1073/pnas.1116158109 (2012).

66 Calabrese, E. J. & Baldwin, L. A. The effects of gamma rays on longevity. Biogerontology 1, 309–319, doi:10.1023/a:1026510001286 (2000).

67 Kenyon, C., Chang, J., Gensch, E., Rudner, A. & Tabtiang, R. A C. elegans mutant that lives twice as long as wild type. Nature 366, 461–464, doi:10.1038/366461a0 (1993).

68 Murphy, C. T. et al. Genes that act downstream of DAF-16 to influence the lifespan of Caenorhabditis elegans. Nature 424, 277–283, doi:10.1038/nature01789 (2003).

69 Ran, Q. et al. Reduction in glutathione peroxidase 4 increases life span through increased sensitivity to apoptosis. J Gerontol A Biol Sci Med Sci 62, 932–942, doi:10.1093/gerona/62.9.932 (2007).

70 Zhang, Y. et al. Mice deficient in both Mn superoxide dismutase and glutathione peroxidase-1 have increased oxidative damage and a greater incidence of pathology but no reduction in longevity. J Gerontol A Biol Sci Med Sci 64, 1212–1220, doi:10.1093/gerona/glp132 (2009).

71 Doonan, R. et al. Against the oxidative damage theory of aging: superoxide dismutases protect against oxidative stress but have little or no effect on life span in Caenorhabditis elegans. Genes Dev 22, 3236–3241, doi:10.1101/gad.504808 (2008).

72 Shanley, D. P. & Kirkwood, T. B. Calorie restriction and aging: a life-history analysis. Evolution 54, 740–750, doi:10.1111/j.0014-3820.2000.tb00076.x (2000).

73 Speakman, J. R. Why does caloric restriction increase life and healthspan? The ‘clean cupboards’ hypothesis. National Science Review 7, 1153–1156, doi:10.1093/nsr/nwaa078 (2020).

74 Mitteldorf, J. Can experiments on caloric restriction be reconciled with the disposable soma theory for the evolution of senescence? Evolution 55, 1902–1905; discussion 1906, doi:10.1111/j.0014-3820.2001.tb00841.x (2001).

75 Dobrzyński, L., Fornalski, K. W. & Feinendegen, L. E. Cancer Mortality Among People Living in Areas With Various Levels of Natural Background Radiation. Dose Response 13, 1559325815592391, doi:10.1177/1559325815592391 (2015).

76 Blagosklonny, M. V. Aging and immortality: quasi-programmed senescence and its pharmacologic inhibition. Cell Cycle 5, 2087–2102, doi:10.4161/cc.5.18.3288 (2006).

77 Longo, V. D., Mitteldorf, J. & Skulachev, V. P. Programmed and altruistic ageing. Nature Reviews Genetics 6, 866–872, doi:10.1038/nrg1706 (2005).

78 Mitteldorf, J. & Sagan, D. Cracking the Aging Code. (Flatiron Books, 2017).

79 Kirkwood, Thomas B. L. & Melov, S. On the Programmed/Non-Programmed Nature of Ageing within the Life History. Current Biology 21, R701–R707, doi:10.1016/j.cub.2011.07.020 (2011).

80 Blagosklonny, M. V. TOR-driven aging: speeding car without brakes. Cell Cycle 8, 4055–4059, doi:10.4161/cc.8.24.10310 (2009).

81 Williams, G. C. Pleiotropy, Natural Selection, and the Evolution of Senescence. Evolution 11, 398–411, doi:10.2307/2406060 (1957).

82 Blagosklonny, M. V. Revisiting the antagonistic pleiotropy theory of aging: TOR-driven program and quasi-program. Cell Cycle 9, 3171–3176, doi:10.4161/cc.9.16.13120 (2010).

83 Medawar, P. B. An Unsolved Problem of Biology. (H. K. Lewis, 1952).

84 Nussey, D. H., Froy, H., Lemaitre, J.-F., Gaillard, J.-M. & Austad, S. N. Senescence in natural populations of animals: widespread evidence and its implications for bio-gerontology. Ageing Res Rev 12, 214–225, doi:10.1016/j.arr.2012.07.004 (2013).

85 Lemaître, J. F., Moorad, J., Gaillard, J. M., Maklakov, A. A. & Nussey, D. H. A unified framework for evolutionary genetic and physiological theories of aging. PLoS Biol 22, e3002513, doi:10.1371/journal.pbio.3002513 (2024).

86 Korpetayev, A. Adaptive ageing theory of faster adaptation and inconsistency of the conventional selection shadow evolutionary theory of ageing[Preprint]. EcoRxiv, 10.32942/osf.io/2wcrm (2020).

87 Haroon et al. Transcriptomic evidence that insulin signalling pathway regulates the ageing of subterranean termite castes. Scientific Reports 10, 8187, doi:10.1038/s41598-020-64890-9 (2020).

88 Maklakov, A. A. & Chapman, T. Evolution of ageing as a tangle of trade-offs: energy versus function. Proceedings of the Royal Society B: Biological Sciences 286, 20191604, doi:doi:10.1098/rspb.2019.1604 (2019).

89 Katcher, H. L. Towards an evidence-based model of aging. Curr Aging Sci 8, 46–55, doi:10.2174/1874609808666150422110601 (2015).

90 Gems, D. The hyperfunction theory: An emerging paradigm for the biology of aging. Ageing Res Rev 74, 101557, doi:10.1016/j.arr.2021.101557 (2022).

91 Farrell, S., Stubbings, G., Rockwood, K., Mitnitski, A. & Rutenberg, A. The potential for complex computational models of aging. Mechanisms of Ageing and Development 193, 111403, 10.1016/j.mad.2020.111403 (2021).

92 Mc Auley, M. T., et al. Modelling the molecular mechanisms of aging. Biosci Rep 37, BSR20160177, doi:10.1042/BSR20160177 (2017).

93 Karin, O., Agrawal, A., Porat, Z., Krizhanovsky, V. & Alon, U. Senescent cell turnover slows with age providing an explanation for the Gompertz law. Nat Commun 10, 5495, doi:10.1038/s41467-019-13192-4 (2019).

94 Hanahan, D. & Weinberg, R. A. The hallmarks of cancer. cell 100, 57–70 (2000).

95 Gems, D. & de Magalhães, J. P. The hoverfly and the wasp: A critique of the hallmarks of aging as a paradigm. Ageing Res Rev 70, 101407, doi:10.1016/j.arr.2021.101407 (2021).

96 Keshavarz, M., Xie, K., Schaaf, K., Bano, D. & Ehninger, D. Targeting the “hallmarks of aging” to slow aging and treat age-related disease: fact or fiction? Molecular Psychiatry 28, 242–255, doi:10.1038/s41380-022-01680-x (2023).

97 Fraser, H. C. et al. Biological mechanisms of aging predict age-related disease co-occurrence in patients. Aging Cell 21, e13524, doi:10.1111/acel.13524 (2022).

98 Kivimäki, M. et al. Obesity and risk of diseases associated with hallmarks of cellular ageing: a multicohort study. Lancet Healthy Longev 5, e454–e463, doi:10.1016/s2666-7568(24)00087-4 (2024).

99 López-Otín, C., Blasco, M. A., Partridge, L., Serrano, M. & Kroemer, G. Hallmarks of aging: An expanding universe. Cell 186, 243–278, 10.1016/j.cell.2022.11.001 (2023).

100 Cheng, C. L., Gao, T. Q., Wang, Z. & Li, D. D. Role of insulin/insulin-like growth factor 1 signaling pathway in longevity. World J Gastroenterol 11, 1891–1895, doi:10.3748/wjg.v11.i13.1891 (2005).

101 Blagosklonny, M. V. Calorie restriction: Decelerating mTOR-driven aging from cells to organisms (including humans). Cell Cycle 9, 683–688, doi:10.4161/cc.9.4.10766 (2010).

102 Borkum, J. M. The Tricarboxylic Acid Cycle as a Central Regulator of the Rate of Aging: Implications for Metabolic Interventions. Adv Biol (Weinh) 7, e2300095, doi:10.1002/adbi.202300095 (2023).

103 Mammucari, C., Schiaffino, S. & Sandri, M. Downstream of Akt: FoxO3 and mTOR in the regulation of autophagy in skeletal muscle. Autophagy 4, 524–526, doi:10.4161/auto.5905 (2008).

104 Kashtan, N. & Alon, U. Spontaneous evolution of modularity and network motifs. Proceedings of the National Academy of Sciences 102, 13773–13778, doi:doi:10.1073/pnas.0503610102 (2005).

105 Gems, D. & Kern, C. C. Biological constraint, evolutionary spandrels and antagonistic pleiotropy. Ageing Res Rev 101, 102527, 10.1016/j.arr.2024.102527 (2024).

106 Chung, Y. M. et al. FOXO3 signalling links ATM to the p53 apoptotic pathway following DNA damage. Nature Communications 3, 1000, doi:10.1038/ncomms2008 (2012).

107 Tsai, W.-B., Chung, Y. M., Takahashi, Y., Xu, Z. & Hu, M. C. T. Functional interaction between FOXO3a and ATM regulates DNA damage response. Nature Cell Biology 10, 460–467, doi:10.1038/ncb1709 (2008).

108 Cheng, Z. FoxO transcription factors in mitochondrial homeostasis. Biochemical Journal 479, 525–536, doi:10.1042/bcj20210777 (2022).

109 Gross, D. N., van den Heuvel, A. P. J. & Birnbaum, M. J. The role of FoxO in the regulation of metabolism. Oncogene 27, 2320–2336, doi:10.1038/onc.2008.25 (2008).

110 Hirosumi, J. et al. A central role for JNK in obesity and insulin resistance. Nature 420, 333–336, doi:10.1038/nature01137 (2002).

111 Shimobayashi, M. et al. Insulin resistance causes inflammation in adipose tissue. The Journal of Clinical Investigation 128, 1538–1550, doi:10.1172/JCI96139 (2018).

112 Liberale, L., Montecucco, F., Tardif, J.-C., Libby, P. & Camici, G. G. Inflamm-ageing: the role of inflammation in age-dependent cardiovascular disease. European Heart Journal 41, 2974–2982, doi:10.1093/eurheartj/ehz961 (2020).

113 Yamauchi, K. et al. Life-Shortening Effect of Chronic Low-Dose-Rate Irradiation in Calorie-Restricted Mice. Radiat Res 192, 451–455, doi:10.1667/rr15385.1 (2019).

114 Barnett, K. et al. Epidemiology of multimorbidity and implications for health care, research, and medical education: a cross-sectional study. The Lancet 380, 37–43, doi:10.1016/S0140-6736(12)60240-2 (2012).

115 Deelen, J. et al. A meta-analysis of genome-wide association studies identifies multiple longevity genes. Nature Communications 10, 3669, doi:10.1038/s41467-019-11558-2 (2019).

116 Raghavachari, N. The Impact of Apolipoprotein E Genetic Variability in Health and Life Span. J Gerontol A Biol Sci Med Sci 75, 1855–1857, doi:10.1093/gerona/glaa175 (2020).

117 Willcox, B. J. et al. The FoxO3 gene and cause-specific mortality. Aging Cell 15, 617–624, doi:10.1111/acel.12452 (2016).

118 Chen, R. et al. Incidence of Alzheimer’s Disease in Men with Late-Life Hypertension Is Ameliorated by FOXO3 Longevity Genotype. J Alzheimers Dis 95, 79–91, doi:10.3233/jad-230350 (2023).

119 Phillips, M. C. Apolipoprotein E isoforms and lipoprotein metabolism. IUBMB Life 66, 616–623, 10.1002/iub.1314 (2014).

120 Eijkelenboom, A. & Burgering, B. M. FOXOs: signalling integrators for homeostasis maintenance. Nature reviews Molecular cell biology 14, 83–97 (2013).

121 Ni, X. et al. A description of the relationship in healthy longevity and aging-related disease: from gene to protein. Immun Ageing 18, 30, doi:10.1186/s12979-021-00241-0 (2021).

122 Anand, R., Prakash, S. S., Veeramanikandan, R. & Kirubakaran, R. Association between apolipoprotein E genotype and cancer susceptibility: a meta-analysis. J Cancer Res Clin Oncol 140, 1075–1085, doi:10.1007/s00432-014-1634-2 (2014).

123 Ference, B. A. et al. Low-density lipoproteins cause atherosclerotic cardiovascular disease. 1. Evidence from genetic, epidemiologic, and clinical studies. A consensus statement from the European Atherosclerosis Society Consensus Panel. European Heart Journal 38, 2459–2472, doi:10.1093/eurheartj/ehx144 (2017).

124 Benn, M., Tybjærg-Hansen, A., Stender, S., Frikke-Schmidt, R. & Nordestgaard, B. G. Low-density lipoprotein cholesterol and the risk of cancer: a mendelian randomization study. J Natl Cancer Inst 103, 508–519, doi:10.1093/jnci/djr008 (2011).

125 Abu Qubo, A., et al. Idiopathic pulmonary fibrosis and lung cancer: future directions and challenges. Breathe (Sheff) 18, 220147, doi:10.1183/20734735.0147-2022 (2022).

126 Brett, S., Irusen, E. M. & Koegelenberg, C. F. N. Pulmonary scarring and its relation to primary lung cancer. Afr J Thorac Crit Care Med 26, doi:10.7196/AJTCCM.2020.v26i1.050 (2020).

127 Reeves, H. L., Zaki, M. Y. W. & Day, C. P. Hepatocellular Carcinoma in Obesity, Type 2 Diabetes, and NAFLD. Digestive Diseases and Sciences 61, 1234-1245, doi:10.1007/s10620-016-4085-6 (2016).

128 Furuta, M. et al. Whole genome sequencing discriminates hepatocellular carcinoma with intrahepatic metastasis from multi-centric tumors. Journal of Hepatology 66, 363–373, 10.1016/j.jhep.2016.09.021 (2017).

129 Kakiuchi, N. & Ogawa, S. Clonal expansion in non-cancer tissues. Nature Reviews Cancer 21, 239–256, doi:10.1038/s41568-021-00335-3 (2021).

130 Fernández-Medarde, A. & Santos, E. Ras in Cancer and Developmental Diseases. Genes & Cancer 2, 344–358, doi:10.1177/1947601911411084 (2011).

131 Di Micco, R. et al. Oncogene-induced senescence is a DNA damage response triggered by DNA hyper-replication. Nature 444, 638–642, doi:10.1038/nature05327 (2006).

132 Karin, O. & Alon, U. Biphasic response as a mechanism against mutant takeover in tissue homeostasis circuits. Mol Syst Biol 13, 933, doi:10.15252/msb.20177599 (2017).

133 Adler, M. et al. Principles of Cell Circuits for Tissue Repair and Fibrosis. iScience 23, 100841–100841, doi:10.1016/j.isci.2020.100841 (2020).

134 Topp, B., Promislow, K., Devries, G., Miura, R. M. & Finegood, D. T. A Model of β-Cell Mass, Insulin, and Glucose Kinetics: Pathways to Diabetes. Journal of Theoretical Biology 206, 605–619, 10.1006/jtbi.2000.2150 (2000).

135 Ginter, E. & Simko, V. in Diabetes: An Old Disease, a New Insight (ed Shamim I. Ahmad) 42-50 (Springer New York, 2013).

136 Chan, A. S. L. et al. Titration of RAS alters senescent state and influences tumour initiation. Nature 633, 678–685, doi:10.1038/s41586-024-07797-z (2024).

137 Baker, N. E. Emerging mechanisms of cell competition. Nature Reviews Genetics 21, 683–697, doi:10.1038/s41576-020-0262-8 (2020).

138 Hogan, C. et al. Characterization of the interface between normal and transformed epithelial cells. Nature Cell Biology 11, 460–467, doi:10.1038/ncb1853 (2009).

139 Maruyama, T. & Fujita, Y. Cell competition in vertebrates — a key machinery for tissue homeostasis. Current Opinion in Genetics & Development 72, 15–21, 10.1016/j.gde.2021.09.006 (2022).

140 Shi, Q. et al. Notch signaling pathway in cancer: from mechanistic insights to targeted therapies. Signal Transduction and Targeted Therapy 9, 128, doi:10.1038/s41392-024-01828-x (2024).

141 Parry, A. J. et al. NOTCH-mediated non-cell autonomous regulation of chromatin structure during senescence. Nature Communications 9, 1840, doi:10.1038/s41467-018-04283-9 (2018).

142 Nelson, P. G. & Masel, J. Intercellular competition and the inevitability of multicellular aging. Proceedings of the National Academy of Sciences of the United States of America 114, 12982–12987, 10.1073/pnas.1618854114 (2017).

143 Argilés, J. M., Busquets, S., Stemmler, B. & López-Soriano, F. J. Cancer cachexia: understanding the molecular basis. Nature Reviews Cancer 14, 754–762, doi:10.1038/nrc3829 (2014).

144 Rybinski, B., Franco-Barraza, J. & Cukierman, E. The wound healing, chronic fibrosis, and cancer progression triad. Physiol Genomics 46, 223–244, doi:10.1152/physiolgenomics.00158.2013 (2014).

145 Holme, J. A. et al. Lung cancer associated with combustion particles and fine particulate matter (PM2.5) - The roles of polycyclic aromatic hydrocarbons (PAHs) and the aryl hydrocarbon receptor (AhR). Biochemical Pharmacology 216, 115801, 10.1016/j.bcp.2023.115801 (2023).

146 Fan, Y. & Bergmann, A. Apoptosis-induced compensatory proliferation. The Cell is dead. Long live the Cell! Trends Cell Biol 18, 467–473, doi:10.1016/j.tcb.2008.08.001 (2008).

147 Vitellius, C. et al. MASLD-related HCC: Multicenter study comparing patients with and without cirrhosis. JHEP Rep 6, 101160, doi:10.1016/j.jhepr.2024.101160 (2024).

148 Gao, R. N. et al. Incidence and mortality of neuroblastoma in Canada compared with other childhood cancers. Cancer Causes & Control 8, 745–754, doi:10.1023/A:1018483405637 (1997).

149 Yazici, S. et al. Risk Factors for Testicular Cancer: Environment, Genes and Infections-Is It All? Medicina (Kaunas) 59, doi:10.3390/medicina59040724 (2023).

150 Elpek, G. Cellular and molecular mechanisms in the pathogenesis of liver fibrosis: An update. World J Gastroenterol 20, 7260–7276, doi:10.3748/wjg.v20.i23.7260 (2014).

151 Ghio, A. J., Sangani, R. G. & Todd, N. W. The Role of Particle Inhalation in Idiopathic Pulmonary Fibrosis. Int J Mol Sci 26, doi:10.3390/ijms26178736 (2025).

152 Landolt, L., Spagnoli, G. C., Hertig, A., Brocheriou, I. & Marti, H.-P. Fibrosis and cancer: shared features and mechanisms suggest common targeted therapeutic approaches. Nephrology Dialysis Transplantation 37, 1024–1032, doi:10.1093/ndt/gfaa301 (2020).

153 Lederer, D. J. & Martinez, F. J. Idiopathic Pulmonary Fibrosis. New England Journal of Medicine 378, 1811–1823, doi:doi:10.1056/NEJMra1705751 (2018).

154 Nho, R. S., Hergert, P., Kahm, J., Jessurun, J. & Henke, C. Pathological Alteration of FoxO3a Activity Promotes Idiopathic Pulmonary Fibrosis Fibroblast Proliferation on Type I Collagen Matrix. The American Journal of Pathology 179, 2420–2430, 10.1016/j.ajpath.2011.07.020 (2011).

155 Speakman, J. R. & Westerterp, K. R. Associations between energy demands, physical activity, and body composition in adult humans between 18 and 96 y of age. The American journal of clinical nutrition 92, 826–834 (2010).

156 Pontzer, H. et al. Daily energy expenditure through the human life course. Science 373, 808–812, doi:10.1126/science.abe5017 (2021).

157 Krems, C., Lührmann, P. M., Straßburg, A., Hartmann, B. & Neuhäuser-Berthold, M. Lower resting metabolic rate in the elderly may not be entirely due to changes in body composition. European Journal of Clinical Nutrition 59, 255–262, doi:10.1038/sj.ejcn.1602066 (2005).

158 Fukagawa, N. K., Bandini, L. G. & Young, J. B. Effect of age on body composition and resting metabolic rate. Am J Physiol 259, E233–238, doi:10.1152/ajpendo.1990.259.2.E233 (1990).

159 Kitazoe, Y., Kishino, H., Tanisawa, K., Udaka, K. & Tanaka, M. Renormalized basal metabolic rate describes the human aging process and longevity. Aging Cell 18, e12968, 10.1111/acel.12968 (2019).

160 Houtkooper, R. H. et al. The metabolic footprint of aging in mice. Scientific Reports 1, 134, doi:10.1038/srep00134 (2011).

161 Greenberg, J. A. Organ metabolic rates and aging: two hypotheses. Med Hypotheses 52, 15–22, doi:10.1054/mehy.1997.0619 (1999).

162 Even, P. C., Rolland, V., Roseau, S., Bouthegourd, J. C. & Tomé, D. Prediction of basal metabolism from organ size in the rat: relationship to strain, feeding, age, and obesity. Am J Physiol Regul Integr Comp Physiol 280, R1887–1896, doi:10.1152/ajpregu.2001.280.6.R1887 (2001).

163 Miyasaka, K. et al. Physical activity prevented age-related decline in energy metabolism in genetically obese and diabetic rats, but not in control rats. Mech Ageing Dev 124, 183–190, doi:10.1016/s0047-6374(02)00118-5 (2003).

164 Speakman, J. R., van Acker, A. & Harper, E. J. Age-related changes in the metabolism and body composition of three dog breeds and their relationship to life expectancy. Aging Cell 2, 265–275, doi:10.1046/j.1474-9728.2003.00061.x (2003).

165 Tomasetti, C. et al. Cell division rates decrease with age, providing a potential explanation for the age-dependent deceleration in cancer incidence. Proceedings of the National Academy of Sciences 116, 20482–20488, doi:10.1073/pnas.1905722116 (2019).

166 Frenk, S. & Houseley, J. Gene expression hallmarks of cellular ageing. Biogerontology 19, 547–566, doi:10.1007/s10522-018-9750-z (2018).

167 Morrison, A. J. Chromatin-remodeling links metabolic signaling to gene expression. Molecular Metabolism 38, 100973, 10.1016/j.molmet.2020.100973 (2020).

168 Faro, T. A. S. & de Oliveira, E. H. C. Canine transmissible venereal tumor – From general to molecular characteristics: A review. Animal Genetics 54, 82–89, 10.1111/age.13260 (2023).

169 Conley, K. E., Jubrias, S. A. & Esselman, P. C. Oxidative capacity and ageing in human muscle. J Physiol 526 Pt 1, 203–210, doi:10.1111/j.1469-7793.2000.t01-1-00203.x (2000).

170 Ruggiero, C. et al. High Basal Metabolic Rate Is a Risk Factor for Mortality: The Baltimore Longitudinal Study of Aging. The Journals of Gerontology: Series A 63, 698–706, doi:10.1093/gerona/63.7.698 (2008).

171 Gray, C. C. et al. Influence of ageing on functional recovery and guanine nucleotide levels of the heart following cold cardioplegic arrest. European journal of cardio-thoracic surgery 13, 475–480 (1998).

172 Vijg, J. & Dollé, M. E. Large genome rearrangements as a primary cause of aging. Mechanisms of ageing and development 123, 907–915 (2002).

173 Ravera, S. et al. Discrete Changes in Glucose Metabolism Define Aging. Scientific Reports 9, 10347, doi:10.1038/s41598-019-46749-w (2019).

174 Wredenberg, A. et al. Increased mitochondrial mass in mitochondrial myopathy mice. Proc Natl Acad Sci U S A 99, 15066–15071, doi:10.1073/pnas.232591499 (2002).

175 Hancock, C. R., Brault, J. J. & Terjung, R. L. Protecting the cellular energy state during contractions: role of AMP deaminase. Journal of physiology and pharmacology: an official journal of the Polish Physiological Society 57 Suppl 10, 17–29 (2006).

176 Karlsson, J. & Saltin, B. Lactate, ATP, and CP in working muscles during exhaustive exercise in man. J Appl Physiol 29, 596–602, doi:10.1152/jappl.1970.29.5.598 (1970).

177 Meizlish, M. L., Franklin, R. A., Zhou, X. & Medzhitov, R. Tissue Homeostasis and Inflammation. Annual Review of Immunology 39, 557–581, doi:10.1146/annurev-immunol-061020-053734 (2021).

178 Gwinn, D. M. et al. AMPK phosphorylation of raptor mediates a metabolic checkpoint. Molecular cell 30, 214–226 (2008).

179 Settembre, C. et al. TFEB controls cellular lipid metabolism through a starvation-induced autoregulatory loop. Nature cell biology 15, 647–658 (2013).

180 Das, A. M. Regulation of the mitochondrial ATP-synthase in health and disease. Molecular Genetics and Metabolism 79, 71–82, 10.1016/S1096-7192(03)00069-6 (2003).

181 Korshunov, S. S., Skulachev, V. P. & Starkov, A. A. High protonic potential actuates a mechanism of production of reactive oxygen species in mitochondria. FEBS letters 416, 15–18 (1997).

182 Starkov, A. A. & Fiskum, G. Regulation of brain mitochondrial H2O2 production by membrane potential and NAD (P) H redox state. Journal of neurochemistry 86, 1101–1107 (2003).

183 Liu, S.-S. Mitochondrial Q cycle–derived superoxide and chemiosmotic bioenergetics. Annals of the New York Academy of Sciences 1201, 84–95, 10.1111/j.1749-6632.2010.05632.x (2010).

184 Liu, S. S. Cooperation of a “reactive oxygen cycle” with the Q cycle and the proton cycle in the respiratory chain--superoxide generating and cycling mechanisms in mitochondria. J Bioenerg Biomembr 31, 367–376, doi:10.1023/a:1018650103259 (1999).

185 Trifunovic, A. et al. Premature ageing in mice expressing defective mitochondrial DNA polymerase. Nature 429, 417–423 (2004).

186 Suhr, S. T. et al. Mitochondrial Rejuvenation After Induced Pluripotency. PLOS ONE 5, e14095, doi:10.1371/journal.pone.0014095 (2010).

187 Ocampo, A. et al. In Vivo Amelioration of Age-Associated Hallmarks by Partial Reprogramming. Cell 167, 1719–1733.e1712, 10.1016/j.cell.2016.11.052 (2016).

188 Wei, Y., Wang, D., Topczewski, F. & Pagliassotti, M. J. Saturated fatty acids induce endoplasmic reticulum stress and apoptosis independently of ceramide in liver cells. Am J Physiol Endocrinol Metab 291, E275–281, doi:10.1152/ajpendo.00644.2005 (2006).

189 van Oort, M. M. et al. Insulin-induced translocation of CD36 to the plasma membrane is reversible and shows similarity to that of GLUT4. Biochim Biophys Acta 1781, 61–71, doi:10.1016/j.bbalip.2007.11.006 (2008).

190 Fryk, E. et al. Hyperinsulinemia and insulin resistance in the obese may develop as part of a homeostatic response to elevated free fatty acids: A mechanistic case-control and a population-based cohort study. eBioMedicine 65, doi:10.1016/j.ebiom.2021.103264 (2021).

191 Sears, B. & Perry, M. The role of fatty acids in insulin resistance. Lipids Health Dis 14, 121, doi:10.1186/s12944-015-0123-1 (2015).

192 Turner, N., Cooney, G. J., Kraegen, E. W. & Bruce, C. R. Fatty acid metabolism, energy expenditure and insulin resistance in muscle. Journal of Endocrinology 220, T61–T79, doi:10.1530/joe-13-0397 (2014).

193 Zhang, X. et al. Fasting insulin, insulin resistance, and risk of cardiovascular or all-cause mortality in non-diabetic adults: a meta-analysis. Biosci Rep 37, doi:10.1042/bsr20170947 (2017).

194 Maciak, S. et al. Low basal metabolic rate as a risk factor for development of insulin resistance and type 2 diabetes. BMJ Open Diabetes Research & Care 8, e001381, doi:10.1136/bmjdrc-2020-001381 (2020).

195 Karin, O., Swisa, A., Glaser, B., Dor, Y. & Alon, U. Dynamical compensation in physiological circuits. Molecular Systems Biology 12, 886, 10.15252/msb.20167216 (2016).

196 Robenek, H. & Severs, N. J. Lipid droplet growth by fusion: insights from freeze-fracture imaging. J Cell Mol Med 13, 4657–4661, doi:10.1111/j.1582-4934.2009.00950.x (2009).

197 Müller, M. J., Bosy-Westphal, A. & Heymsfield, S. B. Is there evidence for a set point that regulates human body weight? F1000 Med Rep 2, 59, doi:10.3410/m2-59 (2010).

198 Bray, G. A. & Bouchard, C. The biology of human overfeeding: A systematic review. Obesity Reviews 21, e13040, 10.1111/obr.13040 (2020).

199 Hirschenson, J., Melgar-Bermudez, E. & Mailloux, R. J. The Uncoupling Proteins: A Systematic Review on the Mechanism Used in the Prevention of Oxidative Stress. Antioxidants (Basel) 11, doi:10.3390/antiox11020322 (2022).

200 Chen, K. Y. et al. Brown fat activation mediates cold-induced thermogenesis in adult humans in response to a mild decrease in ambient temperature. J Clin Endocrinol Metab 98, E1218–1223, doi:10.1210/jc.2012-4213 (2013).

201 Yoneshiro, T. et al. Brown adipose tissue, whole-body energy expenditure, and thermogenesis in healthy adult men. Obesity (Silver Spring) 19, 13–16, doi:10.1038/oby.2010.105 (2011).

202 Orava, J. et al. Blunted metabolic responses to cold and insulin stimulation in brown adipose tissue of obese humans. Obesity (Silver Spring) 21, 2279–2287, doi:10.1002/oby.20456 (2013).

203 Hesselink, M. K. et al. Increased uncoupling protein 3 content does not affect mitochondrial function in human skeletal muscle in vivo. J Clin Invest 111, 479–486, doi:10.1172/jci16653 (2003).

204 Samec, S., Seydoux, J. & Dulloo, A. G. Post-starvation gene expression of skeletal muscle uncoupling protein 2 and uncoupling protein 3 in response to dietary fat levels and fatty acid composition: a link with insulin resistance. Diabetes 48, 436–441, doi:10.2337/diabetes.48.2.436 (1999).

205 Spalding, K. L. et al. Dynamics of fat cell turnover in humans. Nature 453, 783–787, doi:10.1038/nature06902 (2008).

206 Prins, J. B. & O’Rahilly, S. Regulation of adipose cell number in man. Clin Sci (Lond) 92, 3–11, doi:10.1042/cs0920003 (1997).

207 Kral, J. G., Björntorp, P., Scherstén, T. & Sjöström, L. Body composition and adipose tissue cellularity before and after jejuno-ileostomy in severely obese subjects. Eur J Clin Invest 7, 413–419, doi:10.1111/j.1365-2362.1977.tb01628.x (1977).

208 White, U. & Ravussin, E. Dynamics of adipose tissue turnover in human metabolic health and disease. Diabetologia 62, 17–23, doi:10.1007/s00125-018-4732-x (2019).

209 Karczewski, J. et al. Obesity and inflammation. European Cytokine Network 29, 83–94, doi:10.1684/ecn.2018.0415 (2018).

210 Keuper, M. et al. An inflammatory micro-environment promotes human adipocyte apoptosis. Molecular and Cellular Endocrinology 339, 105–113, 10.1016/j.mce.2011.04.004 (2011).

211 Weisberg, S. P. et al. Obesity is associated with macrophage accumulation in adipose tissue. The Journal of clinical investigation 112, 1796–1808 (2003).

212 Lumeng, C. N., Bodzin, J. L. & Saltiel, A. R. Obesity induces a phenotypic switch in adipose tissue macrophage polarization. The Journal of clinical investigation 117, 175–184 (2007).

213 Stolarczyk, E. et al. Improved Insulin Sensitivity despite Increased Visceral Adiposity in Mice Deficient for the Immune Cell Transcription Factor T-bet. Cell Metabolism 17, 520–533, 10.1016/j.cmet.2013.02.019 (2013).

214 Winer, S. et al. Normalization of obesity-associated insulin resistance through immunotherapy. Nature Medicine 15, 921–929, doi:10.1038/nm.2001 (2009).

215 Cinti, S. et al. Adipocyte death defines macrophage localization and function in adipose tissue of obese mice and humans. J Lipid Res 46, 2347–2355, doi:10.1194/jlr.M500294-JLR200 (2005).

216 Fischer-Posovszky, P., Wang, Q. A., Asterholm, I. W., Rutkowski, J. M. & Scherer, P. E. Targeted deletion of adipocytes by apoptosis leads to adipose tissue recruitment of alternatively activated M2 macrophages. Endocrinology 152, 3074–3081, doi:10.1210/en.2011-1031 (2011).

217 Haase, J. et al. Local proliferation of macrophages in adipose tissue during obesity-induced inflammation. Diabetologia 57, 562–571 (2014).

218 Chawla, A. & Lazar, M. A. Peroxisome proliferator and retinoid signaling pathways co-regulate preadipocyte phenotype and survival. Proc Natl Acad Sci U S A 91, 1786–1790, doi:10.1073/pnas.91.5.1786 (1994).

219 Asterholm, W. I. et al. Adipocyte inflammation is essential for healthy adipose tissue expansion and remodeling. Cell Metab 20, 103–118, doi:10.1016/j.cmet.2014.05.005 (2014).

220 Magalhaes, M. S. et al. Role of Tim4 in the regulation of ABCA1(+) adipose tissue macrophages and post-prandial cholesterol levels. Nat Commun 12, 4434, doi:10.1038/s41467-021-24684-7 (2021).

221 Wang, H. H., Garruti, G., Liu, M., Portincasa, P. & Wang, D. Q. H. Cholesterol and Lipoprotein Metabolism and Atherosclerosis: Recent Advances in Reverse Cholesterol Transport. Annals of Hepatology 16, S27–S42, 10.5604/01.3001.0010.5495 (2017).

222 BMJ. BMJ Best Practice: Metabolic syndrome, https://bestpractice.bmj.com/topics/en-gb/212 (2025).

223 Roberge, C. et al. Adrenocortical dysregulation as a major player in insulin resistance and onset of obesity. American Journal of Physiology-Endocrinology and Metabolism 293, E1465–E1478, doi:10.1152/ajpendo.00516.2007 (2007).

224 Raichlen, D. A. et al. Physical activity patterns and biomarkers of cardiovascular disease risk in hunter-gatherers. Am J Hum Biol 29, doi:10.1002/ajhb.22919 (2017).

225 Swanson, Z. S. et al. The effects of lifestyle change on indicators of cardiometabolic health in semi-nomadic pastoralists. Evol Med Public Health 11, 318–331, doi:10.1093/emph/eoad030 (2023).

226 Mangino, M. Genomics of ageing in twins. Proceedings of the Nutrition Society 73, 526–531, doi:10.1017/S0029665114000640 (2014).

227 Hjelmborg, J. v., et al. Genetic influence on human lifespan and longevity. Human Genetics 119, 312–321, doi:10.1007/s00439-006-0144-y (2006).

228 Ruby, J. G. et al. Estimates of the Heritability of Human Longevity Are Substantially Inflated due to Assortative Mating. Genetics 210, 1109–1124, doi:10.1534/genetics.118.301613 (2018).

229 Huang, R., Miszczyk, J. & Zhou, P.-K. Risk and mechanism of metabolic syndrome associated with radiation exposure. Radiation Medicine and Protection 04, 65–69, doi:doi:10.1016/j.radmp.2023.05.001 (2023).

230 Radak, Z., Chung, H. Y. & Goto, S. Exercise and hormesis: oxidative stress-related adaptation for successful aging. Biogerontology 6, 71–75, doi:10.1007/s10522-004-7386-7 (2005).

231 Mair, W., Goymer, P., Pletcher, S. D. & Partridge, L. Demography of dietary restriction and death in Drosophila. Science 301, 1731–1733, doi:10.1126/science.1086016 (2003).

232 Lazarus, N. R. & Harridge, S. D. R. Declining performance of master athletes: silhouettes of the trajectory of healthy human ageing? J Physiol 595, 2941–2948, doi:10.1113/jp272443 (2017).

233 Holloszy, J. O., Smith, E., Vining, M. & Adams, S. Effect of voluntary exercise on longevity of rats. Journal of applied physiology 59, 826–831 (1985).

234 Pekkanen, J. et al. Reduction of premature mortality by high physical activity: a 20-year follow-up of middle-aged Finnish men. The lancet 329, 1473–1477 (1987).

235 Samorajski, T. et al. Effect of exercise on longevity, body weight, locomotor performance, and passive-avoidance memory of C57BL/6J mice. Neurobiology of aging 6, 17–24 (1985).

236 Bronikowski, A. M. et al. Lifelong voluntary exercise in the mouse prevents age-related alterations in gene expression in the heart. Physiological Genomics 12, 129–138 (2003).

237 Harridge, S. D. R. (Unpublished 2024).

238 Gladyshev, V. N. et al. Disagreement on foundational principles of biological aging. PNAS Nexus 3, doi:10.1093/pnasnexus/pgae499 (2024).

239 Friedman, S. L. Hepatic Fibrosis and Cancer: The Silent Threats of Metabolic Syndrome. Diabetes Metab J 48, 161–169, doi:10.4093/dmj.2023.0240 (2024).

240 Shreya, D., Fish, P. N. & Du, D. Navigating the Future of Elderly Healthcare: A Comprehensive Analysis of Aging Populations and Mortality Trends Using National Inpatient Sample (NIS) Data (2010-2024). Cureus 17, e80442, doi:10.7759/cureus.80442 (2025).

241 McGill, H. C., Jr., et al. Origin of atherosclerosis in childhood and adolescence. Am J Clin Nutr 72, 1307s–1315s, doi:10.1093/ajcn/72.5.1307s (2000).

242 Mora, R., Lupu, F. & Simionescu, N. Prelesional events in atherogenesis: colocalization of apolipoprotein B, unesterified cholesterol and extracellular phospholipid liposomes in the aorta of hyperlipidemic rabbit. Atherosclerosis 67, 143–154 (1987).

243 Duewell, P. et al. NLRP3 inflammasomes are required for atherogenesis and activated by cholesterol crystals. Nature 464, 1357–1361 (2010).

244 Baumer, Y., Mehta, N. N., Dey, A. K., Powell-Wiley, T. M. & Boisvert, W. A. Cholesterol crystals and atherosclerosis. Eur Heart J 41, 2236–2239, doi:10.1093/eurheartj/ehaa505 (2020).

245 Hansson, G. K. Inflammation, Atherosclerosis, and Coronary Artery Disease. New England Journal of Medicine 352, 1685–1695, doi:doi:10.1056/NEJMra043430 (2005).

246 Schuitema, P. C. E. et al. Elevated triglycerides are related to higher residual cardiovascular disease and mortality risk independent of lipid targets and intensity of lipid-lowering therapy in patients with established cardiovascular disease. Atherosclerosis 408, 120411, 10.1016/j.atherosclerosis.2025.120411 (2025).

247 Fernández-Friera, L. et al. Normal LDL-cholesterol levels are associated with subclinical atherosclerosis in the absence of risk factors. Journal of the American College of Cardiology 70, 2979–2991 (2017).

248 Chistiakov, D. A., Melnichenko, A. A., Myasoedova, V. A., Grechko, A. V. & Orekhov, A. N. Mechanisms of foam cell formation in atherosclerosis. Journal of Molecular Medicine 95, 1153–1165, doi:10.1007/s00109-017-1575-8 (2017).

249 Cerqueira, N. M. F. S. A. et al. Cholesterol Biosynthesis: A Mechanistic Overview. Biochemistry 55, 5483–5506, doi:10.1021/acs.biochem.6b00342 (2016).

250 Laakso, M. & Fernandes Silva, L. Statins and risk of type 2 diabetes: mechanism and clinical implications. Front Endocrinol (Lausanne) 14, 1239335, doi:10.3389/fendo.2023.1239335 (2023).

251 Jacobs, D. et al. Report of the Conference on Low Blood Cholesterol: Mortality Associations. Circulation 86, 1046–1060, doi:doi:10.1161/01.CIR.86.3.1046 (1992).

252 DuBroff, R. Cholesterol paradox: a correlate does not a surrogate make. Evidence Based Medicine 22, 15–19, doi:10.1136/ebmed-2016-110602 (2017).

253 Petursson, H., Sigurdsson, J. A., Bengtsson, C., Nilsen, T. I. & Getz, L. Is the use of cholesterol in mortality risk algorithms in clinical guidelines valid? Ten years prospective data from the Norwegian HUNT 2 study. J Eval Clin Pract 18, 159-168, doi:10.1111/j.1365-2753.2011.01767.x (2012).

254 Berneis, K. K. & Krauss, R. M. Metabolic origins and clinical significance of LDL heterogeneity. J Lipid Res 43, 1363–1379 (2002).

255 Rizvi, A. A., Stoian, A. P., Janez, A. & Rizzo, M. Lipoproteins and Cardiovascular Disease: An Update on the Clinical Significance of Atherogenic Small, Dense LDL and New Therapeutical Options. Biomedicines 9, 1579 (2021).

256 Ross, R. The pathogenesis of atherosclerosis—an update. New England journal of medicine 314, 488–500 (1986).

257 Kendrick, M. Assessing cardiovascular disease: looking beyond cholesterol. Current Opinion in Endocrinology, Diabetes and Obesity 29 (2022).

258 VanEpps, J. S. & Vorp, D. A. Mechanopathobiology of Atherogenesis: A Review. Journal of Surgical Research 142, 202–217, 10.1016/j.jss.2006.11.001 (2007).

259 Libby, P. Inflammatory mechanisms: the molecular basis of inflammation and disease. Nutr Rev 65, S140–146, doi:10.1111/j.1753-4887.2007.tb00352.x (2007).

260 Castellon, X. & Bogdanova, V. Chronic Inflammatory Diseases and Endothelial Dysfunction. Aging Dis 7, 81–89, doi:10.14336/ad.2015.0803 (2016).

261 ADAPT Research group, A. Cardiovascular and cerebrovascular events in the randomized, controlled Alzheimer’s Disease Anti-Inflammatory Prevention Trial (ADAPT). PLoS Clin Trials 1, e33, doi:10.1371/journal.pctr.0010033 (2006).

262 Trelle, S. et al. Cardiovascular safety of non-steroidal anti-inflammatory drugs: network meta-analysis. Bmj 342, c7086, doi:10.1136/bmj.c7086 (2011).

263 Yoshida, H. & Kisugi, R. Mechanisms of LDL oxidation. Clinica Chimica Acta 411, 1875–1882, 10.1016/j.cca.2010.08.038 (2010).

264 Yoshida, H., Quehenberger, O., Kondratenko, N., Green, S. & Steinberg, D. Minimally oxidized low-density lipoprotein increases expression of scavenger receptor A, CD36, and macrosialin in resident mouse peritoneal macrophages. Arteriosclerosis, thrombosis, and vascular biology 18, 794-802 (1998).

265 Ylä-Herttuala, S. et al. Colocalization of 15-lipoxygenase mRNA and protein with epitopes of oxidized low density lipoprotein in macrophage-rich areas of atherosclerotic lesions. Proc Natl Acad Sci U S A 87, 6959–6963, doi:10.1073/pnas.87.18.6959 (1990).

266 Shen, J. et al. Macrophage-mediated 15-lipoxygenase expression protects against atherosclerosis development. The Journal of clinical investigation 98, 2201–2208 (1996).

267 Alghareeb, R., Hussain, A., Maheshwari, M. V., Khalid, N. & Patel, P. D. Cardiovascular Complications in Systemic Lupus Erythematosus. Cureus 14, e26671, doi:10.7759/cureus.26671 (2022).

268 Zeng, G. & Quon, M. J. Insulin-stimulated production of nitric oxide is inhibited by wortmannin. Direct measurement in vascular endothelial cells. The Journal of clinical investigation 98, 894–898 (1996).

269 Michell, B. J. et al. The Akt kinase signals directly to endothelial nitric oxide synthase. Curr Biol 9, 845–848, doi:10.1016/s0960-9822(99)80371-6 (1999).

270 Gerber, H.-P. et al. Vascular endothelial growth factor regulates endothelial cell survival through the phosphatidylinositol 3′-kinase/Akt signal transduction pathway: requirement for Flk-1/KDR activation. Journal of Biological Chemistry 273, 30336–30343 (1998).

271 Cyr, A. R., Huckaby, L. V., Shiva, S. S. & Zuckerbraun, B. S. Nitric Oxide and Endothelial Dysfunction. Crit Care Clin 36, 307–321, doi:10.1016/j.ccc.2019.12.009 (2020).

272 Drozd, M. et al. Endothelial insulin-like growth factor-1 signalling regulates vascular barrier function and atherogenesis. Cardiovascular Research 121, 1108–1120, doi:10.1093/cvr/cvaf055 (2025).

273 Di Pino, A. & DeFronzo, R. A. Insulin Resistance and Atherosclerosis: Implications for Insulin-Sensitizing Agents. Endocrine Reviews 40, 1447–1467, doi:10.1210/er.2018-00141 (2019).

274 Anderson, E. J. et al. Mitochondrial H2O2 emission and cellular redox state link excess fat intake to insulin resistance in both rodents and humans. The Journal of Clinical Investigation 119, 573–581, doi:10.1172/JCI37048 (2009).

275 Nolan, C. J., Ruderman, N. B., Kahn, S. E., Pedersen, O. & Prentki, M. Insulin Resistance as a Physiological Defense Against Metabolic Stress: Implications for the Management of Subsets of Type 2 Diabetes. Diabetes 64, 673–686, doi:10.2337/db14-0694 (2015).

276 Blagosklonny, M. V. Prospective Treatment of Age-Related Diseases by Slowing Down Aging. The American Journal of Pathology 181, 1142–1146, 10.1016/j.ajpath.2012.06.024 (2012).

277 Zhang, J. Biomarkers of endothelial activation and dysfunction in cardiovascular diseases. RCM 23, doi:10.31083/j.rcm2302073 (2022).

278 Harman, J. L. & Jørgensen, H. F. The role of smooth muscle cells in plaque stability: Therapeutic targeting potential. Br J Pharmacol 176, 3741–3753, doi:10.1111/bph.14779 (2019).

279 Bhatia, H. S. et al. Aspirin and Cardiovascular Risk in Individuals With Elevated Lipoprotein(a): The Multi-Ethnic Study of Atherosclerosis. J Am Heart Assoc 13, e033562, doi:10.1161/jaha.123.033562 (2024).

280 Engelhardt, E., Resende, E. P. F. & Gomes, K. B. Physiopathological mechanisms underlying Alzheimer’s disease: a narrative review. Dement Neuropsychol 18, e2024VR2001, doi:10.1590/1980-5764-dn-2024-vr01 (2024).

281 Heneka, M. T. et al. Neuroinflammation in Alzheimer disease. Nature Reviews Immunology 25, 321–352, doi:10.1038/s41577-024-01104-7 (2025).

282 Toyama, B. H. & Hetzer, M. W. Protein homeostasis: live long, won’t prosper. Nat Rev Mol Cell Biol 14, 55–61, doi:10.1038/nrm3496 (2013).

283 Dai, M. H., Zheng, H., Zeng, L. D. & Zhang, Y. The genes associated with early-onset Alzheimer’s disease. Oncotarget 9, 15132–15143, doi:10.18632/oncotarget.23738 (2018).

284 Koychev, I., Hofer, M. & Friedman, N. Correlation of Alzheimer Disease Neuropathologic Staging with Amyloid and Tau Scintigraphic Imaging Biomarkers. Journal of Nuclear Medicine 61, 1413–1418, doi:10.2967/jnumed.119.230458 (2020).

285 Streit, W. J., Khoshbouei, H. & Bechmann, I. The Role of Microglia in Sporadic Alzheimer’s Disease. J Alzheimers Dis 79, 961–968, doi:10.3233/jad-201248 (2021).

286 De Grey, A. & Rae, M. Ending aging: the rejuvenation breakthroughs that could reverse human aging in our lifetime. 193 (St. Martin’s Press, 2007).

287 Terman, A. Garbage catastrophe theory of aging: imperfect removal of oxidative damage? Redox Rep 6, 15–26, doi:10.1179/135100001101535996 (2001).

288 Brunk, U. T. & Terman, A. The mitochondrial-lysosomal axis theory of aging: accumulation of damaged mitochondria as a result of imperfect autophagocytosis. Eur J Biochem 269, 1996–2002, doi:10.1046/j.1432-1033.2002.02869.x (2002).

289 Reines, S. A. et al. Rofecoxib: no effect on Alzheimer’s disease in a 1-year, randomized, blinded, controlled study. Neurology 62, 66–71, doi:10.1212/wnl.62.1.66 (2004).

290 Pasqualetti, P. et al. A randomized controlled study on effects of ibuprofen on cognitive progression of Alzheimer’s disease. Aging Clin Exp Res 21, 102–110, doi:10.1007/bf03325217 (2009).

291 Lyketsos, C. G. et al. Naproxen and celecoxib do not prevent AD in early results from a randomized controlled trial. Neurology 68, 1800–1808, doi:10.1212/01.wnl.0000260269.93245.d2 (2007).

292 Meyer, P. F. et al. INTREPAD: A randomized trial of naproxen to slow progress of presymptomatic Alzheimer disease. Neurology 92, e2070–e2080, doi:10.1212/wnl.0000000000007232 (2019).

293 McCaffrey, P. Closing the Book on NSAIDs for Alzheimer’s Prevention, https://www.alzforum.org/news/research-news/closing-book-nsaids-alzheimers-prevention (2019).

294 Hardy, J. & Lewis, P. Evidence suggesting that microglia make amyloid from neuronally expressed APP: a hypothesis. Mol Neurodegener 20, 55, doi:10.1186/s13024-025-00847-8 (2025).

295 Nguyen, T. T., Ta, Q. T. H., Nguyen, T. K. O., Nguyen, T. T. D. & Giau, V. V. Type 3 Diabetes and Its Role Implications in Alzheimer’s Disease. Int J Mol Sci 21, doi:10.3390/ijms21093165 (2020).

296 Craft, S. & Stennis Watson, G. Insulin and neurodegenerative disease: shared and specific mechanisms. The Lancet Neurology 3, 169–178, 10.1016/S1474-4422(04)00681-7 (2004).

297 Lowe, V. J. et al. Association of hypometabolism and amyloid levels in aging, normal subjects. Neurology 82, 1959–1967, doi:10.1212/wnl.0000000000000467 (2014).

298 Nguyen, T. T., Ta, Q. T. H., Nguyen, T. T. D., Le, T. T. & Vo, V. G. Role of Insulin Resistance in the Alzheimer’s Disease Progression. Neurochemical Research 45, 1481–1491, doi:10.1007/s11064-020-03031-0 (2020).

299 Watson, G. S. & Craft, S. Modulation of memory by insulin and glucose: neuropsychological observations in Alzheimer’s disease. Eur J Pharmacol 490, 97–113, doi:10.1016/j.ejphar.2004.02.048 (2004).

300 McMillan, J. M., Mele, B. S., Hogan, D. B. & Leung, A. A. Impact of pharmacological treatment of diabetes mellitus on dementia risk: systematic review and meta-analysis. BMJ Open Diabetes Research & Care 6, e000563, doi:10.1136/bmjdrc-2018-000563 (2018).

301 Di Paolo, G. & Kim, T. W. Linking lipids to Alzheimer’s disease: cholesterol and beyond. Nat Rev Neurosci 12, 284–296, doi:10.1038/nrn3012 (2011).

302 Ajoolabady, A., Lindholm, D., Ren, J. & Pratico, D. ER stress and UPR in Alzheimer’s disease: mechanisms, pathogenesis, treatments. Cell Death & Disease 13, 706, doi:10.1038/s41419-022-05153-5 (2022).

303 Ghasemi, R., Haeri, A., Dargahi, L., Mohamed, Z. & Ahmadiani, A. Insulin in the Brain: Sources, Localization and Functions. Molecular Neurobiology 47, 145–171, doi:10.1007/s12035-012-8339-9 (2013).

304 Sasaki, N. et al. Advanced glycation end products in Alzheimer’s disease and other neurodegenerative diseases. Am J Pathol 153, 1149–1155, doi:10.1016/s0002-9440(10)65659-3 (1998).

305 Fournet, M., Bonté, F. & Desmoulière, A. Glycation Damage: A Possible Hub for Major Pathophysiological Disorders and Aging. Aging Dis 9, 880–900, doi:10.14336/ad.2017.1121 (2018).

306 Moldogazieva, N. T., Mokhosoev, I. M., Mel’nikova, T. I., Porozov, Y. B. & Terentiev, A. A. Oxidative Stress and Advanced Lipoxidation and Glycation End Products (ALEs and AGEs) in Aging and Age-Related Diseases. Oxid Med Cell Longev 2019, 3085756, doi:10.1155/2019/3085756 (2019).

307 Uribarri, J. et al. Elevated Serum Advanced Glycation Endproducts in Obese Indicate Risk for the Metabolic Syndrome: A Link Between Healthy and Unhealthy Obesity? The Journal of Clinical Endocrinology & Metabolism 100, 1957–1966, doi:10.1210/jc.2014-3925 (2015).

308 Cansu, K., Rebecca, A. F., So Jung, P., Mariana, P. & David, C. R. Autophagy in ageing and ageing-related neurodegenerative diseases. Ageing and Neurodegenerative Diseases 1, 2, doi:10.20517/and.2021.05 (2021).

309 Boland, B. et al. Autophagy induction and autophagosome clearance in neurons: relationship to autophagic pathology in Alzheimer’s disease. J Neurosci 28, 6926–6937, doi:10.1523/jneurosci.0800-08.2008 (2008).

310 Tian, Y., Bustos, V., Flajolet, M. & Greengard, P. A small-molecule enhancer of autophagy decreases levels of Abeta and APP-CTF via Atg5-dependent autophagy pathway. Faseb j 25, 1934–1942, doi:10.1096/fj.10-175158 (2011).

311 Vingtdeux, V. et al. Novel synthetic small-molecule activators of AMPK as enhancers of autophagy and amyloid-β peptide degradation. Faseb j 25, 219–231, doi:10.1096/fj.10-167361 (2011).

312 Carroll, B., Hewitt, G. & Korolchuk, V. I. Autophagy and ageing: implications for age-related neurodegenerative diseases. Essays in Biochemistry 55, 119–131, doi:10.1042/bse0550119 (2013).

313 Lavallard, V. J., Meijer, A. J., Codogno, P. & Gual, P. Autophagy, signaling and obesity. Pharmacological Research 66, 513–525, 10.1016/j.phrs.2012.09.003 (2012).

314 Gonskikh, Y. & Polacek, N. Alterations of the translation apparatus during aging and stress response. Mechanisms of Ageing and Development 168, 30–36, 10.1016/j.mad.2017.04.003 (2017).

315 Motizuki, M. & Tsurugi, K. The effect of aging on protein synthesis in the yeast Saccharomyces cerevisiae. Mech Ageing Dev 64, 235–245, doi:10.1016/0047-6374(92)90081-n (1992).

316 Solyga, M., Majumdar, A. & Besse, F. Regulating translation in aging: from global to gene-specific mechanisms. EMBO reports 25, 5265–5276, 10.1038/s44319-024-00315-2 (2024).

317 Kim, H. S. & Pickering, A. M. Protein translation paradox: Implications in translational regulation of aging. Frontiers in Cell and Developmental Biology Volume 11 - 2023, doi:10.3389/fcell.2023.1129281 (2023).

318 Lim, S. H. Y., Hansen, M. & Kumsta, C. Molecular Mechanisms of Autophagy Decline during Aging. Cells 13, doi:10.3390/cells13161364 (2024).

319 Simonsen, A. et al. Promoting basal levels of autophagy in the nervous system enhances longevity and oxidant resistance in adult Drosophila. Autophagy 4, 176–184, doi:10.4161/auto.5269 (2008).

320 Lipinski, M. M. et al. Genome-wide analysis reveals mechanisms modulating autophagy in normal brain aging and in Alzheimer’s disease. Proc Natl Acad Sci U S A 107, 14164–14169, doi:10.1073/pnas.1009485107 (2010).

321 Khalil, H. et al. Aging is associated with hypermethylation of autophagy genes in macrophages. Epigenetics 11, 381–388, doi:10.1080/15592294.2016.1144007 (2016).

322 Zhang, Y., Chen, H., Li, R., Sterling, K. & Song, W. Amyloid β-based therapy for Alzheimer’s disease: challenges, successes and future. Signal Transduction and Targeted Therapy 8, 248, doi:10.1038/s41392-023-01484-7 (2023).

323 Serrano-Pozo, A. et al. Mild to moderate Alzheimer dementia with insufficient neuropathological changes. Ann Neurol 75, 597–601, doi:10.1002/ana.24125 (2014).

324 Rodrigue, K. M., Kennedy, K. M. & Park, D. C. Beta-amyloid deposition and the aging brain. Neuropsychol Rev 19, 436–450, doi:10.1007/s11065-009-9118-x (2009).

325 Katzman, R. et al. Clinical, pathological, and neurochemical changes in dementia: a subgroup with preserved mental status and numerous neocortical plaques. Ann Neurol 23, 138–144, doi:10.1002/ana.410230206 (1988).

326 Jagust, W. Imaging the evolution and pathophysiology of Alzheimer disease. Nature Reviews Neuroscience 19, 687–700, doi:10.1038/s41583-018-0067-3 (2018).

327 O’Leary, J., Georgeaux-Healy, C. & Serpell, L. The impact of continuous calorie restriction and fasting on cognition in adults without eating disorders. Nutr Rev 83, 146–159, doi:10.1093/nutrit/nuad170 (2025).

328 Korte, N., Nortley, R. & Attwell, D. Cerebral blood flow decrease as an early pathological mechanism in Alzheimer’s disease. Acta Neuropathologica 140, 793–810, doi:10.1007/s00401-020-02215-w (2020).

329 Wingo, A. P. et al. Shared proteomic effects of cerebral atherosclerosis and Alzheimer’s disease on the human brain. Nat Neurosci 23, 696–700, doi:10.1038/s41593-020-0635-5 (2020).

330 Tong, Q.-Y. et al. Human Thymic Involution and Aging in Humanized Mice. Frontiers in Immunology Volume 11 - 2020, doi:10.3389/fimmu.2020.01399 (2020).

331 Conway, J., Certo, M., Lord, J. M., Mauro, C. & Duggal, N. A. Understanding the role of host metabolites in the induction of immune senescence: Future strategies for keeping the ageing population healthy. Br J Pharmacol 179, 1808–1824, 10.1111/bph.15671 (2022).

332 Vegiopoulos, A., Rohm, M. & Herzig, S. Adipose tissue: between the extremes. The EMBO Journal 36, 1999–2017, 10.15252/embj.201696206 (2017).

333 Sopasakis, V. R. et al. High Local Concentrations and Effects on Differentiation Implicate Interleukin-6 as a Paracrine Regulator. Obesity Research 12, 454–460, 10.1038/oby.2004.51 (2004).

334 Booth, A. D. et al. Subcutaneous adipose tissue accumulation protects systemic glucose tolerance and muscle metabolism. Adipocyte 7, 261–272, doi:10.1080/21623945.2018.1525252 (2018).

335 Rytka, J. M., Wueest, S., Schoenle, E. J. & Konrad, D. The portal theory supported by venous drainage-selective fat transplantation. Diabetes 60, 56–63, doi:10.2337/db10-0697 (2011).

336 Kolb, H. Obese visceral fat tissue inflammation: from protective to detrimental? BMC Med 20, 494, doi:10.1186/s12916-022-02672-y (2022).

337 Doran, M. F., Pond, G. R., Crowson, C. S., O’Fallon, W. M. & Gabriel, S. E. Trends in incidence and mortality in rheumatoid arthritis in Rochester, Minnesota, over a forty-year period. Arthritis Rheum 46, 625–631, doi:10.1002/art.509 (2002).

338 Conrad, N. et al. Incidence, prevalence, and co-occurrence of autoimmune disorders over time and by age, sex, and socioeconomic status: a population-based cohort study of 22 million individuals in the UK. The Lancet 401, 1878–1890, doi:10.1016/S0140-6736(23)00457-9 (2023).

339 Vadasz, Z., Haj, T., Kessel, A. & Toubi, E. Age-related autoimmunity. BMC Medicine 11, 94, doi:10.1186/1741-7015-11-94 (2013).

340 Raynor, J., Lages, C. S., Shehata, H., Hildeman, D. A. & Chougnet, C. A. Homeostasis and function of regulatory T cells in aging. Curr Opin Immunol 24, 482–487, doi:10.1016/j.coi.2012.04.005 (2012).

341 Laukoetter, M. G. et al. Intestinal Cancer Risk in Crohn’s Disease: A Meta-Analysis. Journal of Gastrointestinal Surgery 15, 576–583, 10.1007/s11605-010-1402-9 (2011).

342 Ding, Z. et al. Hepatocellular carcinoma: pathogenesis, molecular mechanisms, and treatment advances. Front Oncol 15, 1526206, doi:10.3389/fonc.2025.1526206 (2025).

343 Anders, C. K. et al. Young age at diagnosis correlates with worse prognosis and defines a subset of breast cancers with shared patterns of gene expression. J Clin Oncol 26, 3324–3330, doi:10.1200/jco.2007.14.2471 (2008).

344 Chatsirisupachai, K., Lagger, C. & de Magalhães, J. P. Age-associated differences in the cancer molecular landscape. Trends in Cancer 8, 962–971, doi:10.1016/j.trecan.2022.06.007 (2022).

345 Chen, X., Yang, W., Roberts, C. W. M. & Zhang, J. Developmental origins shape the paediatric cancer genome. Nat Rev Cancer 24, 382–398, doi:10.1038/s41568-024-00684-9 (2024).

346 Ferron, M., McKee, M. D., Levine, R. L., Ducy, P. & Karsenty, G. Intermittent injections of osteocalcin improve glucose metabolism and prevent type 2 diabetes in mice. Bone 50, 568–575, doi:10.1016/j.bone.2011.04.017 (2012).

347 Saleem, U., Mosley, T. H., Jr. & Kullo, I. J. Serum osteocalcin is associated with measures of insulin resistance, adipokine levels, and the presence of metabolic syndrome. Arterioscler Thromb Vasc Biol 30, 1474–1478, doi:10.1161/atvbaha.110.204859 (2010).

348 Potikanond, S. et al. Obesity does not aggravate osteoporosis or osteoblastic insulin resistance in orchiectomized rats. J Endocrinol 228, 85–95, doi:10.1530/joe-15-0333 (2016).

349 Fried, L. P. et al. Frailty in older adults: evidence for a phenotype. J Gerontol A Biol Sci Med Sci 56, M146–156, doi:10.1093/gerona/56.3.m146 (2001).

350 Goodpaster, B. H. et al. The Loss of Skeletal Muscle Strength, Mass, and Quality in Older Adults: The Health, Aging and Body Composition Study. The Journals of Gerontology: Series A 61, 1059–1064, doi:10.1093/gerona/61.10.1059 (2006).

351 Srikanthan, P. & Karlamangla, A. S. Relative Muscle Mass Is Inversely Associated with Insulin Resistance and Prediabetes. Findings from The Third National Health and Nutrition Examination Survey. The Journal of Clinical Endocrinology & Metabolism 96, 2898–2903, doi:10.1210/jc.2011-0435 (2011).

352 Kozakiewicz, M., Kornatowski, M., Krzywińska, O. & Kędziora-Kornatowska, K. Changes in the blood antioxidant defense of advanced age people. Clin Interv Aging 14, 763–771, doi:10.2147/cia.S201250 (2019).

353 Moriwaki, S.-I., Ray, S., Tarone, R. E., Kraemer, K. H. & Grossman, L. The effect of donor age on the processing of UV-damaged DNA by cultured human cells: reduced DNA repair capacity and increased DNA mutability. Mutation Research/DNA Repair 364, 117–123 (1996).

354 Vijg, J. & Dollé, M. E. T. Large genome rearrangements as a primary cause of aging. Mechanisms of Ageing and Development 123, 907–915, 10.1016/S0047-6374(02)00028-3 (2002).

355 Goukassian, D. et al. Mechanisms and implications of the age-associated decrease in DNA repair capacity. The FASEB journal 14, 1325–1334 (2000).

356 Blagosklonny, M. V. Paradoxes of aging. Cell cycle 6, 2997–3003 (2007).

357 Anyiam, O. et al. A Systematic Review and Meta-Analysis of the Effect of Caloric Restriction on Skeletal Muscle Mass in Individuals with, and without, Type 2 Diabetes. Nutrients 16, doi:10.3390/nu16193328 (2024).

358 Veronese, N. & Reginster, J.-Y. The effects of calorie restriction, intermittent fasting and vegetarian diets on bone health. Aging Clinical and Experimental Research 31, 753–758, doi:10.1007/s40520-019-01174-x (2019).

359 Larson-Meyer, D. E. et al. Effect of calorie restriction with or without exercise on insulin sensitivity, β-cell function, fat cell size, and ectopic lipid in overweight subjects. Diabetes care 29, 1337–1344 (2006).

360 Kökten, T. et al. Calorie Restriction as a New Treatment of Inflammatory Diseases. Adv Nutr 12, 1558–1570, doi:10.1093/advances/nmaa179 (2021).

361 Hahn, O. et al. A nutritional memory effect counteracts the benefits of dietary restriction in old mice. Nature Metabolism 1, 1059–1073, doi:10.1038/s42255-019-0121-0 (2019).

362 Flegal, K. M. The obesity wars and the education of a researcher: A personal account. Progress in Cardiovascular Diseases 67, 75–79, 10.1016/j.pcad.2021.06.009 (2021).

363 Lavie, C. J., Milani, R. V. & Ventura, H. O. Obesity and cardiovascular disease: risk factor, paradox, and impact of weight loss. J Am Coll Cardiol 53, 1925–1932, doi:10.1016/j.jacc.2008.12.068 (2009).

364 Simati, S., Kokkinos, A., Dalamaga, M. & Argyrakopoulou, G. Obesity Paradox: Fact or Fiction? Curr Obes Rep 12, 75–85, doi:10.1007/s13679-023-00497-1 (2023).

365 Flegal, K. M., Kit, B. K., Orpana, H. & Graubard, B. I. Association of all-cause mortality with overweight and obesity using standard body mass index categories: a systematic review and meta-analysis. Jama 309, 71–82, doi:10.1001/jama.2012.113905 (2013).

366 Oshakbayev, K. et al. Overweight effects on metabolic rate, time perception, diseases, aging, and lifespan: A systematic review with meta-regression analysis. Translational Medicine of Aging 9, 15–24, 10.1016/j.tma.2024.12.001 (2025).

367 Pantazopoulos, D., Gouveri, E., Papazoglou, D. & Papanas, N. GLP-1 receptor agonists and sarcopenia: Weight loss at a cost? A brief narrative review. Diabetes Res Clin Pract 229, 112924, doi:10.1016/j.diabres.2025.112924 (2025).

368 Song, M. et al. Longitudinal Analysis of Genetic Susceptibility and BMI Throughout Adult Life. Diabetes 67, 248–255, doi:10.2337/db17-1156 (2018).

369 Newman, A. B. et al. Weight change and the conservation of lean mass in old age: the Health, Aging and Body Composition Study. Am J Clin Nutr 82, 872–878; quiz 915-876, doi:10.1093/ajcn/82.4.872 (2005).

370 Imada, S. et al. Short-term post-fast refeeding enhances intestinal stemness via polyamines. Nature 633, 895–904, doi:10.1038/s41586-024-07840-z (2024).

371 Harrison, D. E. et al. Rapamycin fed late in life extends lifespan in genetically heterogeneous mice. Nature 460, 392–395, doi:10.1038/nature08221 (2009).

372 Miller, R. A. et al. Rapamycin, but not resveratrol or simvastatin, extends life span of genetically heterogeneous mice. The Journals of Gerontology: Series A 66, 191–201 (2011).

373 Bjedov, I. et al. Mechanisms of life span extension by rapamycin in the fruit fly Drosophila melanogaster. Cell metabolism 11, 35–46 (2010).

374 Juricic, P. et al. Long-lasting geroprotection from brief rapamycin treatment in early adulthood by persistently increased intestinal autophagy. Nature Aging, doi:10.1038/s43587-022-00278-w (2022).

375 Sabatini, D. M., Erdjument-Bromage, H., Lui, M., Tempst, P. & Snyder, S. H. RAFT1: a mammalian protein that binds to FKBP12 in a rapamycin-dependent fashion and is homologous to yeast TORs. Cell 78, 35–43 (1994).

376 Brown, E. J. et al. A mammalian protein targeted by G1-arresting rapamycin–receptor complex. Nature 369, 756–758 (1994).

377 Wullschleger, S., Loewith, R. & Hall, M. N. TOR signaling in growth and metabolism. Cell 124, 471–484 (2006).

378 Condon, K. J. & Sabatini, D. M. Nutrient regulation of mTORC1 at a glance. Journal of Cell Science 132, doi:10.1242/jcs.222570 (2019).

379 Drummond, M. J. et al. Amino acids are necessary for the insulin-induced activation of mTOR/S6K1 signaling and protein synthesis in healthy and insulin resistant human skeletal muscle. Clin Nutr 27, 447–456, doi:10.1016/j.clnu.2008.01.012 (2008).

380 Miller, R. A. et al. Rapamycin-mediated lifespan increase in mice is dose and sex dependent and metabolically distinct from dietary restriction. Aging Cell 13, 468–477, 10.1111/acel.12194 (2014).

381 Lamming, D. W. et al. Rapamycin-Induced Insulin Resistance Is Mediated by mTORC2 Loss and Uncoupled from Longevity. Science 335, 1638, doi:10.1126/science.1215135 (2012).

382 Fang, Y. et al. Duration of rapamycin treatment has differential effects on metabolism in mice. Cell Metab 17, 456–462, doi:10.1016/j.cmet.2013.02.008 (2013).

383 Flynn, J. M. et al. Late-life rapamycin treatment reverses age-related heart dysfunction. Aging Cell 12, 851–862, doi:10.1111/acel.12109 (2013).

384 Schindler, C. E., Partap, U., Patchen, B. K. & Swoap, S. J. Chronic rapamycin treatment causes diabetes in male mice. Am J Physiol Regul Integr Comp Physiol 307, R434–R443, doi:10.1152/ajpregu.00123.2014 (2014).

385 Cohen, E. & Dillin, A. The insulin paradox: aging, proteotoxicity and neurodegeneration. Nature Reviews Neuroscience 9, 759–767, doi:10.1038/nrn2474 (2008).

386 Sharples, A. P. et al. Longevity and skeletal muscle mass: the role of IGF signalling, the sirtuins, dietary restriction and protein intake. Aging Cell 14, 511–523, 10.1111/acel.12342 (2015).

387 Finkelstein, J., Roffwarg, H., Boyar, R., Kream, J. & Hellman, L. Age-related change in the twenty-four hour spontaneous secretion of growth hormone. The Journal of Clinical Endocrinology & Metabolism 35, 665–670 (1972).

388 Zadik, Z., Chalew, A., McCARTER JR, R. J., Meistas, M. & Kowarski, A. A. The influence of age on the 24-hour integrated concentration of growth hormone in normal individuals. The Journal of Clinical Endocrinology & Metabolism 60, 513–516 (1985).

389 Clasey, J. L. et al. Abdominal visceral fat and fasting insulin are important predictors of 24-hour GH release independent of age, gender, and other physiological factors. The Journal of Clinical Endocrinology & Metabolism 86, 3845–3852 (2001).

390 Jørgensen, J. et al. Beneficial effects of growth hormone treatment in GH-deficient adults. The lancet 333, 1221–1225 (1989).

391 Liu, H. et al. Systematic review: the safety and efficacy of growth hormone in the healthy elderly. Ann Intern Med 146, 104–115, doi:10.7326/0003-4819-146-2-200701160-00005 (2007).

392 Brown-Borg, H. M., Borg, K. E., Meliska, C. J. & Bartke, A. Dwarf mice and the ageing process. Nature 384, 33–33 (1996).

393 List, E. O. et al. Endocrine Parameters and Phenotypes of the Growth Hormone Receptor Gene Disrupted (GHR−/−) Mouse. Endocrine Reviews 32, 356–386, doi:10.1210/er.2010-0009 (2011).

394 Feng, M. et al. Early-life exercise extends healthspan but not lifespan in mice. Nature Communications 16, 6328, doi:10.1038/s41467-025-61443-4 (2025).

395 Holmstrom, L. et al. Sudden Cardiac Arrest During Sports Activity in Older Adults. JACC: Clinical Electrophysiology 9, 893–903, doi:doi:10.1016/j.jacep.2022.10.033 (2023).

396 Teekakirikul, P. et al. Cardiac fibrosis in mice with hypertrophic cardiomyopathy is mediated by non-myocyte proliferation and requires Tgf-β. J Clin Invest 120, 3520–3529, doi:10.1172/jci42028 (2010).

397 Brown, J. C., Winters-Stone, K., Lee, A. & Schmitz, K. H. Cancer, Physical Activity, and Exercise. Comprehensive Physiology 2, 2775–2809, 10.1002/j.2040-4603.2012.tb00476.x (2012).

398 Rena, G., Hardie, D. G. & Pearson, E. R. The mechanisms of action of metformin. Diabetologia 60, 1577–1585, doi:10.1007/s00125-017-4342-z (2017).

399 Martin-Montalvo, A. et al. Metformin improves healthspan and lifespan in mice. Nat Commun 4, 2192, doi:10.1038/ncomms3192 (2013).

400 Karise, I., Bargut, T. C., Del Sol, M., Aguila, M. B. & Mandarim-de-Lacerda, C. A. Metformin enhances mitochondrial biogenesis and thermogenesis in brown adipocytes of mice. Biomed Pharmacother 111, 1156–1165, doi:10.1016/j.biopha.2019.01.021 (2019).

401 Geerling, J. J. et al. Metformin lowers plasma triglycerides by promoting VLDL-triglyceride clearance by brown adipose tissue in mice. Diabetes 63, 880–891 (2014).

402 Schommers, P. et al. Metformin causes a futile intestinal-hepatic cycle which increases energy expenditure and slows down development of a type 2 diabetes-like state. Mol Metab 6, 737–747, doi:10.1016/j.molmet.2017.05.002 (2017).

403 Onken, B. & Driscoll, M. Metformin induces a dietary restriction-like state and the oxidative stress response to extend C. elegans Healthspan via AMPK, LKB1, and SKN-1. PLoS One 5, e8758, doi:10.1371/journal.pone.0008758 (2010).

404 Alfaras, I. et al. Health benefits of late-onset metformin treatment every other week in mice. npj Aging and Mechanisms of Disease 3, 16, doi:10.1038/s41514-017-0018-7 (2017).

405 Slack, C., Foley, A. & Partridge, L. Activation of AMPK by the Putative Dietary Restriction Mimetic Metformin Is Insufficient to Extend Lifespan in Drosophila. PLOS ONE 7, e47699, doi:10.1371/journal.pone.0047699 (2012).

406 Cabreiro, F. et al. Metformin Retards Aging in C. elegans by Altering Microbial Folate and Methionine Metabolism. Cell 153, 228–239, 10.1016/j.cell.2013.02.035 (2013).

407 Carlson, B. & Faulkner, J. Muscle transplantation between young and old rats: age of host determines recovery. American Journal of Physiology-Cell Physiology 256, C1262–C1266 (1989).

408 Carlson, B. M., Dedkov, E. I., Borisov, A. B. & Faulkner, J. A. Skeletal muscle regeneration in very old rats. The Journals of Gerontology Series A: Biological Sciences and Medical Sciences 56, B224–B233 (2001).

409 Mehdipour, M. et al. Rejuvenation of three germ layers tissues by exchanging old blood plasma with saline-albumin. Aging (Albany NY) 12, 8790–8819, doi:10.18632/aging.103418 (2020).

410 Coskun, P. E., Beal, M. F. & Wallace, D. C. Alzheimer’s brains harbor somatic mtDNA control-region mutations that suppress mitochondrial transcription and replication. Proc Natl Acad Sci U S A 101, 10726–10731, doi:10.1073/pnas.0403649101 (2004).

411 Swerdlow, R. H., Burns, J. M. & Khan, S. M. The Alzheimer’s disease mitochondrial cascade hypothesis: Progress and perspectives. Biochimica et Biophysica Acta (BBA) - Molecular Basis of Disease 1842, 1219–1231, 10.1016/j.bbadis.2013.09.010 (2014).

412 Hoekstra, J. G., Hipp, M. J., Montine, T. J. & Kennedy, S. R. Mitochondrial DNA mutations increase in early stage Alzheimer disease and are inconsistent with oxidative damage. Ann Neurol 80, 301–306, doi:10.1002/ana.24709 (2016).

413 Kauppila, T. E. S. et al. Mutations of mitochondrial DNA are not major contributors to aging of fruit flies. Proceedings of the National Academy of Sciences 115, E9620–E9629, doi:doi:10.1073/pnas.1721683115 (2018).

414 Ng, L. F., Ng, L. T., van Breugel, M., Halliwell, B. & Gruber, J. Mitochondrial DNA Damage Does Not Determine C. elegans Lifespan. Frontiers in Genetics 10, doi:10.3389/fgene.2019.00311 (2019).

415 Siibak, T. et al. A multi-systemic mitochondrial disorder due to a dominant p.Y955H disease variant in DNA polymerase gamma. Hum Mol Genet 26, 2515-2525, doi:10.1093/hmg/ddx146 (2017).

416 Samstag, C. L. et al. Deleterious mitochondrial DNA point mutations are overrepresented in Drosophila expressing a proofreading-defective DNA polymerase γ. PLoS Genet 14, e1007805, doi:10.1371/journal.pgen.1007805 (2018).

417 Haroon, S. et al. Multiple Molecular Mechanisms Rescue mtDNA Disease in C. elegans. Cell Rep 22, 3115–3125, doi:10.1016/j.celrep.2018.02.099 (2018).

418 Jazwinski, S. M. Yeast replicative life span--the mitochondrial connection. FEMS Yeast Res 5, 119–125, doi:10.1016/j.femsyr.2004.04.005 (2004).

419 Pernice, W. M., Swayne, T. C., Boldogh, I. R. & Pon, L. A. Mitochondrial Tethers and Their Impact on Lifespan in Budding Yeast. Front Cell Dev Biol 5, 120, doi:10.3389/fcell.2017.00120 (2017).

420 Azbarova, A. V. & Knorre, D. A. Role of Mitochondrial DNA in Yeast Replicative Aging. Biochemistry (Moscow) 88, 1997–2006, doi:10.1134/S0006297923120040 (2023).

421 Vermulst, M. et al. Mitochondrial point mutations do not limit the natural lifespan of mice. Nature Genetics 39, 540–543, doi:10.1038/ng1988 (2007).

422 Khrapko, K. & Vijg, J. Mitochondrial DNA mutations and aging: a case closed? Nature Genetics 39, 445–446, doi:10.1038/ng0407-445 (2007).

423 Howes, R. M. The free radical fantasy: a panoply of paradoxes. Ann N Y Acad Sci 1067, 22–26, doi:10.1196/annals.1354.004 (2006).

424 Trifunovic, A. et al. Somatic mtDNA mutations cause aging phenotypes without affecting reactive oxygen species production. Proceedings of the National Academy of Sciences 102, 17993–17998, doi:doi:10.1073/pnas.0508886102 (2005).

425 Zheng, W., Khrapko, K., Coller, H. A., Thilly, W. G. & Copeland, W. C. Origins of human mitochondrial point mutations as DNA polymerase γ-mediated errors. Mutation Research/Fundamental and Molecular Mechanisms of Mutagenesis 599, 11–20 (2006).

426 Khrapko, K. et al. Mitochondrial mutational spectra in human cells and tissues. Proceedings of the National Academy of Sciences 94, 13798–13803, doi:doi:10.1073/pnas.94.25.13798 (1997).

427 Wallace, D. C. et al. Mitochondrial DNA mutations in human degenerative diseases and aging. Biochimica et Biophysica Acta (BBA)-Molecular Basis of Disease 1271, 141–151 (1995).

428 Gadaleta, M. N. et al. Mitochondrial DNA copy number and mitochondrial DNA deletion in adult and senescent rats. Mutat Res 275, 181–193, doi:10.1016/0921-8734(92)90022-h (1992).

429 Brierley, E. J., Johnson, M. A., Lightowlers, R. N., James, O. F. & Turnbull, D. M. Role of mitochondrial DNA mutations in human aging: implications for the central nervous system and muscle. Ann Neurol 43, 217–223, doi:10.1002/ana.410430212 (1998).

430 Elson, J. L., Samuels, D. C., Turnbull, D. M. & Chinnery, P. F. Random intracellular drift explains the clonal expansion of mitochondrial DNA mutations with age. Am J Hum Genet 68, 802–806, doi:10.1086/318801 (2001).

431 Lawless, C., Greaves, L., Reeve, A. K., Turnbull, D. M. & Vincent, A. E. The rise and rise of mitochondrial DNA mutations. Open Biol 10, 200061, doi:10.1098/rsob.200061 (2020).

432 Insalata, F., Hoitzing, H., Aryaman, J. & Jones, N. S. Stochastic survival of the densest and mitochondrial DNA clonal expansion in aging. Proceedings of the National Academy of Sciences 119, e2122073119, doi:doi:10.1073/pnas.2122073119 (2022).

433 Kremer, L. S. et al. The bottleneck for maternal transmission of mtDNA is linked to purifying selection by autophagy. Sci Adv 11, eaea4660, doi:10.1126/sciadv.aea4660 (2025).

434 Sato, K. & Sato, M. Multiple ways to prevent transmission of paternal mitochondrial DNA for maternal inheritance in animals. The Journal of Biochemistry 162, 247–253, doi:10.1093/jb/mvx052 (2017).

435 Diaz, F. et al. Human mitochondrial DNA with large deletions repopulates organelles faster than full-length genomes under relaxed copy number control. Nucleic Acids Res 30, 4626–4633, doi:10.1093/nar/gkf602 (2002).

436 Kowald, A. & Kirkwood, T. B. Evolution of the mitochondrial fusion-fission cycle and its role in aging. Proc Natl Acad Sci U S A 108, 10237–10242, doi:10.1073/pnas.1101604108 (2011).

437 Kowald, A. & Kirkwood, T. B. Mitochondrial mutations and aging: random drift is insufficient to explain the accumulation of mitochondrial deletion mutants in short-lived animals. Aging Cell 12, 728–731, doi:10.1111/acel.12098 (2013).

438 Cao, Z., Wanagat, J., McKiernan, S. H. & Aiken, J. M. Mitochondrial DNA deletion mutations are concomitant with ragged red regions of individual, aged muscle fibers: analysis by laser-capture microdissection. Nucleic Acids Res 29, 4502–4508, doi:10.1093/nar/29.21.4502 (2001).

439 Wallace, D. C. Mitochondrial genetics: a paradigm for aging and degenerative diseases? Science 256, 628–632 (1992).

440 Moraes, C. T., Kenyon, L. & Hao, H. Mechanisms of human mitochondrial DNA maintenance: the determining role of primary sequence and length over function. Mol Biol Cell 10, 3345–3356, doi:10.1091/mbc.10.10.3345 (1999).

441 Hayashi, J. et al. Introduction of disease-related mitochondrial DNA deletions into HeLa cells lacking mitochondrial DNA results in mitochondrial dysfunction. Proceedings of the National Academy of Sciences 88, 10614–10618 (1991).

442 Takai, D., Isobe, K. & Hayashi, J.-I. Transcomplementation between different types of respiration-deficient mitochondria with different pathogenic mutant mitochondrial DNAs. Journal of Biological Chemistry 274, 11199–11202 (1999).

443 de Grey, A. D. A proposed refinement of the mitochondrial free radical theory of aging. Bioessays 19, 161–166, doi:10.1002/bies.950190211 (1997).

444 De Grey, A. D. The mitochondrial free radical theory of aging. (RG Landes Austin, TX, 1999).

445 Kowald, A., Dawson, M. & Kirkwood, T. B. Mitochondrial mutations and ageing: can mitochondrial deletion mutants accumulate via a size based replication advantage? J Theor Biol 340, 111–118, doi:10.1016/j.jtbi.2013.09.009 (2014).

446 Suski, J. M. et al. Relation between mitochondrial membrane potential and ROS formation. Methods Mol Biol 810, 183–205, doi:10.1007/978-1-61779-382-0_12 (2012).

447 Li, X. X., Tsoi, B., Li, Y. F., Kurihara, H. & He, R. R. Cardiolipin and its different properties in mitophagy and apoptosis. J Histochem Cytochem 63, 301–311, doi:10.1369/0022155415574818 (2015).

448 Frank, M. et al. Mitophagy is triggered by mild oxidative stress in a mitochondrial fission dependent manner. Biochim Biophys Acta 1823, 2297–2310, doi:10.1016/j.bbamcr.2012.08.007 (2012).

449 Wang, Y., Nartiss, Y., Steipe, B., McQuibban, G. A. & Kim, P. K. ROS-induced mitochondrial depolarization initiates PARK2/PARKIN-dependent mitochondrial degradation by autophagy. Autophagy 8, 1462–1476, doi:10.4161/auto.21211 (2012).

450 Kim, I. & Lemasters, J. J. Mitophagy selectively degrades individual damaged mitochondria after photoirradiation. Antioxid Redox Signal 14, 1919–1928, doi:10.1089/ars.2010.3768 (2011).

451 Iborra, F. J., Kimura, H. & Cook, P. R. The functional organization of mitochondrial genomes in human cells. BMC biology 2, 1–14 (2004).

452 Gilkerson, R. W., Schon, E. A., Hernandez, E. & Davidson, M. M. Mitochondrial nucleoids maintain genetic autonomy but allow for functional complementation. J Cell Biol 181, 1117–1128 (2008).

453 Sukhorukov, V. M. et al. Determination of protein mobility in mitochondrial membranes of living cells. Biochimica et Biophysica Acta (BBA)-Biomembranes 1798, 2022–2032 (2010).

454 Lewis, S. C., Uchiyama, L. F. & Nunnari, J. ER-mitochondria contacts couple mtDNA synthesis with mitochondrial division in human cells. Science 353, aaf5549 (2016).

455 Margineantu, D. H. et al. Cell cycle dependent morphology changes and associated mitochondrial DNA redistribution in mitochondria of human cell lines. Mitochondrion 1, 425–435 (2002).

456 Legros, F., Malka, F., Frachon, P., Lombès, A. & Rojo, M. Organization and dynamics of human mitochondrial DNA. Journal of cell science 117, 2653–2662 (2004).

457 Kang, D. & Hamasaki, N. Mitochondrial transcription factor A in the maintenance of mitochondrial DNA: overview of its multiple roles. Annals of the New York Academy of Sciences 1042, 101–108 (2005).

458 Rey, T. et al. Mitochondrial RNA granules are fluid condensates positioned by membrane dynamics. Nat Cell Biol 22, 1180–1186, doi:10.1038/s41556-020-00584-8 (2020).

459 Seelert, H. et al. From protons to OXPHOS supercomplexes and Alzheimer’s disease: structure–dynamics–function relationships of energy-transducing membranes. Biochimica Et Biophysica Acta (BBA)-Bioenergetics 1787, 657–671 (2009).

460 Frenzel, M., Rommelspacher, H., Sugawa, M. D. & Dencher, N. A. Ageing alters the supramolecular architecture of OxPhos complexes in rat brain cortex. Experimental gerontology 45, 563–572 (2010).

461 Wang, Y. & Bogenhagen, D. F. Human Mitochondrial DNA Nucleoids Are Linked to Protein Folding Machinery and Metabolic Enzymes at the Mitochondrial Inner Membrane*. Journal of Biological Chemistry 281, 25791–25802, 10.1074/jbc.M604501200 (2006).

462 Zorkau, M., Albus, C. A., Berlinguer-Palmini, R., Chrzanowska-Lightowlers, Z. M. A. & Lightowlers, R. N. High-resolution imaging reveals compartmentalization of mitochondrial protein synthesis in cultured human cells. Proc Natl Acad Sci U S A 118, doi:10.1073/pnas.2008778118 (2021).

463 Wilkens, V., Kohl, W. & Busch, K. Restricted diffusion of OXPHOS complexes in dynamic mitochondria delays their exchange between cristae and engenders a transitory mosaic distribution. Journal of cell science 126, 103–116 (2013).

464 Mannella, C. A. et al. Topology of the mitochondrial inner membrane: dynamics and bioenergetic implications. IUBMB life 52, 93–100 (2001).

465 Twig, G. et al. Fission and selective fusion govern mitochondrial segregation and elimination by autophagy. Embo j 27, 433–446, doi:10.1038/sj.emboj.7601963 (2008).

466 Elmore, S. P., Qian, T., Grissom, S. F. & Lemasters, J. J. The mitochondrial permeability transition initiates autophagy in rat hepatocytes. The FASEB Journal 15, 1–17 (2001).

467 Priault, M. et al. Impairing the bioenergetic status and the biogenesis of mitochondria triggers mitophagy in yeast. Cell Death & Differentiation 12, 1613–1621 (2005).

468 Kadenbach, B., Arnold, S., Lee, I. & Hüttemann, M. The possible role of cytochrome c oxidase in stress-induced apoptosis and degenerative diseases. Biochim Biophys Acta 1655, 400–408, doi:10.1016/j.bbabio.2003.06.005 (2004).

469 Juhaszova, M. et al. Glycogen synthase kinase-3β mediates convergence of protection signaling to inhibit the mitochondrial permeability transition pore. The Journal of Clinical Investigation 113, 1535–1549, doi:10.1172/JCI19906 (2004).

470 Kokoszka, J. E., Coskun, P., Esposito, L. A. & Wallace, D. C. Increased mitochondrial oxidative stress in the Sod2 (+/−) mouse results in the age-related decline of mitochondrial function culminating in increased apoptosis. Proceedings of the National Academy of Sciences 98, 2278–2283, doi:doi:10.1073/pnas.051627098 (2001).

471 LaFrance, R., Brustovetsky, N., Sherburne, C., Delong, D. & Dubinsky, J. M. Age-related changes in regional brain mitochondria from Fischer 344 rats. Aging Cell 4, 139–145, doi:10.1111/j.1474-9726.2005.00156.x (2005).

472 Hagen, T. M. et al. Mitochondrial decay in hepatocytes from old rats: Membrane potential declines, heterogeneity and oxidants increase. Proceedings of the National Academy of Sciences 94, 3064–3069, doi:10.1073/pnas.94.7.3064 (1997).

473 Hiona, A. et al. Mitochondrial DNA Mutations Induce Mitochondrial Dysfunction, Apoptosis and Sarcopenia in Skeletal Muscle of Mitochondrial DNA Mutator Mice. PLOS ONE 5, e11468, doi:10.1371/journal.pone.0011468 (2010).

474 Koopman, W. J. et al. Mammalian mitochondrial complex I: biogenesis, regulation, and reactive oxygen species generation. Antioxid Redox Signal 12, 1431–1470, doi:10.1089/ars.2009.2743 (2010).

475 St-Pierre, J., Buckingham, J. A., Roebuck, S. J. & Brand, M. D. Topology of superoxide production from different sites in the mitochondrial electron transport chain. J Biol Chem 277, 44784–44790, doi:10.1074/jbc.M207217200 (2002).

476 Kowald, A. & Kirkwood, T. B. L. Transcription could be the key to the selection advantage of mitochondrial deletion mutants in aging. Proceedings of the National Academy of Sciences 111, 2972–2977, doi:doi:10.1073/pnas.1314970111 (2014).

477 Bodyak, N. D., Nekhaeva, E., Wei, J. Y. & Khrapko, K. Quantification and sequencing of somatic deleted mtDNA in single cells: evidence for partially duplicated mtDNA in aged human tissues. Hum Mol Genet 10, 17–24, doi:10.1093/hmg/10.1.17 (2001).

478 Kajander, O. A. et al. Human mtDNA sublimons resemble rearranged mitochondrial genoms found in pathological states. Hum Mol Genet 9, 2821–2835, doi:10.1093/hmg/9.19.2821 (2000).

479 Moore, C. A., Gudikote, J. & Van Tuyle, G. C. Mitochondrial DNA rearrangements, including partial duplications, occur in young and old rat tissues. Mutat Res 421, 205–217, doi:10.1016/s0027-5107(98)00169-9 (1998).

480 Herbst, A. et al. Accumulation of mitochondrial DNA deletion mutations in aged muscle fibers: evidence for a causal role in muscle fiber loss. J Gerontol A Biol Sci Med Sci 62, 235–245, doi:10.1093/gerona/62.3.235 (2007).

481 Wanagat, J., Cao, Z., Pathare, P. & Aiken, J. M. Mitochondrial DNA deletion mutations colocalize with segmental electron transport system abnormalities, muscle fiber atrophy, fiber splitting, and oxidative damage in sarcopenia. Faseb j 15, 322–332, doi:10.1096/fj.00-0320com (2001).

482 Müller-Höcker, J. Cytochrome-c-oxidase deficient cardiomyocytes in the human heart--an age-related phenomenon. A histochemical ultracytochemical study. Am J Pathol 134, 1167–1173 (1989).

483 Cortopassi, G. A., Shibata, D., Soong, N. W. & Arnheim, N. A pattern of accumulation of a somatic deletion of mitochondrial DNA in aging human tissues. Proc Natl Acad Sci U S A 89, 7370–7374, doi:10.1073/pnas.89.16.7370 (1992).

484 Bua, E. et al. Mitochondrial DNA–deletion mutations accumulate intracellularly to detrimental levels in aged human skeletal muscle fibers. The American Journal of Human Genetics 79, 469–480 (2006).

485 Mohamed, S. A. et al. Mitochondrial DNA deletions and the aging heart. Experimental gerontology 41, 508–517 (2006).

486 Bender, A. et al. High levels of mitochondrial DNA deletions in substantia nigra neurons in aging and Parkinson disease. Nature genetics 38, 515–517 (2006).

487 Kraytsberg, Y. et al. Mitochondrial DNA deletions are abundant and cause functional impairment in aged human substantia nigra neurons. Nature genetics 38, 518–520 (2006).

488 Ozawa, T. et al. Quantitative determination of deleted mitochondrial DNA relative to normal DNA in parkinsonian striatum by a kinetic PCR analysis. Biochemical and Biophysical Research Communications 172, 483–489, 10.1016/0006-291X(90)90698-M (1990).

489 Phillips, N. R., Simpkins, J. W. & Roby, R. K. Mitochondrial DNA deletions in Alzheimer’s brains: a review. Alzheimers Dement 10, 393–400, doi:10.1016/j.jalz.2013.04.508 (2014).

490 Baris, O. R. et al. Mosaic deficiency in mitochondrial oxidative metabolism promotes cardiac arrhythmia during aging. Cell metabolism 21, 667–677 (2015).

491 Bates, M. G. et al. Cardiac involvement in mitochondrial DNA disease: clinical spectrum, diagnosis, and management. European heart journal 33, 3023–3033 (2012).

492 Lakshmanan, L. N. et al. Clonal expansion of mitochondrial DNA deletions is a private mechanism of aging in long-lived animals. Aging Cell 17, e12814, 10.1111/acel.12814 (2018).

493 Ahlqvist, Kati J. et al. Somatic Progenitor Cell Vulnerability to Mitochondrial DNA Mutagenesis Underlies Progeroid Phenotypes in Polg Mutator Mice. Cell Metabolism 15, 100–109, 10.1016/j.cmet.2011.11.012 (2012).

494 Tyynismaa, H. et al. Mutant mitochondrial helicase Twinkle causes multiple mtDNA deletions and a late-onset mitochondrial disease in mice. Proceedings of the National Academy of Sciences 102, 17687–17692 (2005).

495 McFaline-Figueroa, J. R. et al. Mitochondrial quality control during inheritance is associated with lifespan and mother-daughter age asymmetry in budding yeast. Aging Cell 10, 885–895, doi:10.1111/j.1474-9726.2011.00731.x (2011).

496 Aguilaniu, H., Gustafsson, L., Rigoulet, M. & Nyström, T. Asymmetric inheritance of oxidatively damaged proteins during cytokinesis. Science 299, 1751–1753, doi:10.1126/science.1080418 (2003).

497 Regmi, S., Rolland, S. G. & Conradt, B. Age-dependent changes in mitochondrial morphology and volume are not predictors of lifespan. Aging (Albany NY) 6, 10.18632/aging.100639 (2014).

498 Navarro, A. & Boveris, A. The mitochondrial energy transduction system and the aging process. American Journal of Physiology-Cell Physiology 292, C670–C686, doi:10.1152/ajpcell.00213.2006 (2007).

499 Ubaida-Mohien, C. et al. Discovery proteomics in aging human skeletal muscle finds change in spliceosome, immunity, proteostasis and mitochondria. eLife 8, e49874, doi:10.7554/eLife.49874 (2019).

500 Thomsen, K. et al. Initial brain aging: heterogeneity of mitochondrial size is associated with decline in complex I-linked respiration in cortex and hippocampus. Neurobiol Aging 61, 215–224, doi:10.1016/j.neurobiolaging.2017.08.004 (2018).

501 Bratic, I. & Trifunovic, A. Mitochondrial energy metabolism and ageing. Biochimica et Biophysica Acta (BBA) - Bioenergetics 1797, 961–967, 10.1016/j.bbabio.2010.01.004 (2010).

502 Regmi, S. G., Rolland, S. G. & Conradt, B. Age-dependent changes in mitochondrial morphology and volume are not predictors of lifespan. Aging (Albany NY) 6, 118–130, doi:10.18632/aging.100639 (2014).

503 Crane, J. D., Devries, M. C., Safdar, A., Hamadeh, M. J. & Tarnopolsky, M. A. The Effect of Aging on Human Skeletal Muscle Mitochondrial and Intramyocellular Lipid Ultrastructure. The Journals of Gerontology: Series A 65A, 119–128, doi:10.1093/gerona/glp179 (2009).

504 Cole, M. A. et al. A high fat diet increases mitochondrial fatty acid oxidation and uncoupling to decrease efficiency in rat heart. Basic Res Cardiol 106, 447–457, doi:10.1007/s00395-011-0156-1 (2011).

505 Qin, C., Liu, Z., Duan, D. & Wu, L. Genome-wide RNAi analysis in adult Caenorhabditis elegans unveils genes controlling age-related fat accumulation. Mech Ageing Dev 226, 112089, doi:10.1016/j.mad.2025.112089 (2025).

506 Morcos, M. et al. Glyoxalase-1 prevents mitochondrial protein modification and enhances lifespan in Caenorhabditis elegans. Aging Cell 7, 260–269, 10.1111/j.1474-9726.2008.00371.x (2008).

507 Lemire, B. D., Behrendt, M., DeCorby, A. & Gášková, D. C. elegans longevity pathways converge to decrease mitochondrial membrane potential. Mechanisms of Ageing and Development 130, 461–465, 10.1016/j.mad.2009.05.001 (2009).

508 Sagi, D. & Kim, S. K. An Engineering Approach to Extending Lifespan in C. elegans. PLOS Genetics 8, e1002780, doi:10.1371/journal.pgen.1002780 (2012).

509 Padalko, V. I. Uncoupler of oxidative phosphorylation prolongs the lifespan of Drosophila. Biochemistry (Mosc) 70, 986–989, doi:10.1007/s10541-005-0213-1 (2005).

510 Ulgherait, M. et al. Circadian regulation of mitochondrial uncoupling and lifespan. Nat Commun 11, 1927, doi:10.1038/s41467-020-15617-x (2020).

511 Fridell, Y.-W. C., Sánchez-Blanco, A., Silvia, B. A. & Helfand, S. L. Targeted expression of the human uncoupling protein 2 (hUCP2) to adult neurons extends life span in the fly. Cell metabolism 1, 145–152 (2005).

512 Andrews, Z. B. & Horvath, T. L. Uncoupling protein-2 regulates lifespan in mice. Am J Physiol Endocrinol Metab 296, E621–627, doi:10.1152/ajpendo.90903.2008 (2009).

513 Padalko, V. “Mild” mitochondrial uncoupling as potentially effective intervention to slow aging. Oxid Antioxid Med Sci 3, 27–42, doi:10.5455/oams.161213.rv.012 (2014).

514 Brand, M. D. Uncoupling to survive? The role of mitochondrial inefficiency in ageing. Exp Gerontol 35, 811–820, doi:10.1016/s0531-5565(00)00135-2 (2000).

515 Berry, B. J. et al. Preservation of mitochondrial membrane potential is necessary for lifespan extension from dietary restriction. Geroscience 45, 1573–1581, doi:10.1007/s11357-023-00766-w (2023).

516 Nonninger, T. J. et al. A TFEB–TGFβ axis systemically regulates diapause, stem cell resilience and protects against a senescence-like state. Nature Aging 5, 1340–1357, doi:10.1038/s43587-025-00911-4 (2025).

517 Schoenfeldt, L. et al. Chemical reprogramming ameliorates cellular hallmarks of aging and extends lifespan. EMBO Mol Med 17, 2071–2094, 10.1038/s44321-025-00265-9 (2025).

518 Rea, S. L., Ventura, N. & Johnson, T. E. Relationship Between Mitochondrial Electron Transport Chain Dysfunction, Development, and Life Extension in Caenorhabditis elegans. PLOS Biology 5, e259, doi:10.1371/journal.pbio.0050259 (2007).

519 Schaar, C. E. et al. Mitochondrial and cytoplasmic ROS have opposing effects on lifespan. PLoS Genet 11, e1004972, doi:10.1371/journal.pgen.1004972 (2015).

520 Chi, S. et al. Oncogenic Ras triggers cell suicide through the activation of a caspase-independent cell death program in human cancer cells. Oncogene 18, 2281–2290, doi:10.1038/sj.onc.1202538 (1999).

521 Gupta, S. Molecular mechanisms of apoptosis in the cells of the immune system in human aging. Immunol Rev 205, 114–129, doi:10.1111/j.0105-2896.2005.00261.x (2005).

522 Tower, J. Programmed cell death in aging. Ageing Res Rev 23, 90–100, doi:10.1016/j.arr.2015.04.002 (2015).

523 Renehan, A. G., Booth, C. & Potten, C. S. What is apoptosis, and why is it important? Bmj 322, 1536–1538, doi:10.1136/bmj.322.7301.1536 (2001).

524 Wang, W., Wang, D. & Li, H. Initiation of premature senescence by Bcl-2 in hypoxic condition. Int J Clin Exp Pathol 7, 2446–2453 (2014).

525 Iannello, A., Thompson, T. W., Ardolino, M., Lowe, S. W. & Raulet, D. H. p53-dependent chemokine production by senescent tumor cells supports NKG2D-dependent tumor elimination by natural killer cells. J Exp Med 210, 2057–2069, doi:10.1084/jem.20130783 (2013).

526 Nagasawa, H. & Little, J. B. Induction of sister chromatid exchanges by extremely low doses of alpha-particles. Cancer Res 52, 6394–6396 (1992).

527 Gems, D. & Kern, C. C. Is “cellular senescence” a misnomer? GeroScience 44, 2461–2469, doi:10.1007/s11357-022-00652-x (2022).

528 de Magalhães, J. P. Cellular senescence in normal physiology. Science 384, 1300–1301, doi:doi:10.1126/science.adj7050 (2024).

529 Krizhanovsky, V. et al. Senescence of activated stellate cells limits liver fibrosis. Cell 134, 657–667, doi:10.1016/j.cell.2008.06.049 (2008).

530 Lengefeld, J. et al. Cell size is a determinant of stem cell potential during aging. Sci Adv 7, eabk0271, doi:10.1126/sciadv.abk0271 (2021).

531 Welle, S. et al. Reduced amount of mitochondrial DNA in aged human muscle. Journal of Applied Physiology 94, 1479–1484, doi:10.1152/japplphysiol.01061.2002 (2003).

532 Barazzoni, R., Short, K. R. & Nair, K. S. Effects of aging on mitochondrial DNA copy number and cytochrome c oxidase gene expression in rat skeletal muscle, liver, and heart. J Biol Chem 275, 3343–3347, doi:10.1074/jbc.275.5.3343 (2000).

533 Snijders, R. J., Sundberg, K., Holzgreve, W., Henry, G. & Nicolaides, K. H. Maternal age-and gestation-specific risk for trisomy 21. Ultrasound Obstet Gynecol 13, 167–170, doi:10.1046/j.1469-0705.1999.13030167.x (1999).

534 Sinclair, K. D. et al. Healthy ageing of cloned sheep. Nature Communications 7, 12359, doi:10.1038/ncomms12359 (2016).

535 Miller, R. A. ‘Accelerated aging’: a primrose path to insight? Aging Cell 3, 47–51, 10.1111/j.1474-9728.2004.00081.x (2004).

536 Woody, R. C., Harding, B. N., Brumback, R. A. & Leech, R. W. Absence of beta-amyloid immunoreactivity in mesial temporal lobe in Cockayne’s syndrome. J Child Neurol 6, 32–34, doi:10.1177/088307389100600106 (1991).

537 Laposa, R. R., Huang, E. J. & Cleaver, J. E. Increased apoptosis, p53 up-regulation, and cerebellar neuronal degeneration in repair-deficient Cockayne syndrome mice. Proc Natl Acad Sci U S A 104, 1389–1394, doi:10.1073/pnas.0610619104 (2007).

538 Kubben, N. & Misteli, T. Shared molecular and cellular mechanisms of premature ageing and ageing-associated diseases. Nat Rev Mol Cell Biol 18, 595–609, doi:10.1038/nrm.2017.68 (2017).

539 Gordon, L. B., Harten, I. A., Patti, M. E. & Lichtenstein, A. H. Reduced adiponectin and HDL cholesterol without elevated C-reactive protein: clues to the biology of premature atherosclerosis in Hutchinson-Gilford Progeria Syndrome. J Pediatr 146, 336–341, doi:10.1016/j.jpeds.2004.10.064 (2005).

540 Hamczyk, M. R. & Andrés, V. Vascular smooth muscle cell loss underpins the accelerated atherosclerosis in Hutchinson-Gilford progeria syndrome. Nucleus 10, 28–34, doi:10.1080/19491034.2019.1589359 (2019).

541 Olive, M. et al. Cardiovascular pathology in Hutchinson-Gilford progeria: correlation with the vascular pathology of aging. Arterioscler Thromb Vasc Biol 30, 2301–2309, doi:10.1161/atvbaha.110.209460 (2010).

542 Schumacher, B., Pothof, J., Vijg, J. & Hoeijmakers, J. H. J. The central role of DNA damage in the ageing process. Nature 592, 695–703, doi:10.1038/s41586-021-03307-7 (2021).

543 Gladyshev, V. N. et al. Molecular Damage in Aging. Nat Aging 1, 1096–1106, doi:10.1038/s43587-021-00150-3 (2021).

544 Alexander, P. & Connell, D. I. Shortening of the life span of mice by irradiation with X-rays and treatment with radiomimetic chemicals. Radiation Research 12, 38–48 (1960).

545 Ali, Y. F., Cucinotta, F. A., Ning-Ang, L. & Zhou, G. Cancer Risk of Low Dose Ionizing Radiation. Frontiers in Physics Volume 8 - 2020, doi:10.3389/fphy.2020.00234 (2020).

546 Richardson, R. B. Ionizing radiation and aging: rejuvenating an old idea. Aging (Albany NY) 1, 887–902, doi:10.18632/aging.100081 (2009).

547 Yakunchikova, E. A. et al. Model of Accelerated Aging in CB6F2 Mice Induced by Ionizing Radiation. Bull Exp Biol Med 177, 363–367, doi:10.1007/s10517-024-06190-0 (2024).

548 Morgan, H. D., Santos, F., Green, K., Dean, W. & Reik, W. Epigenetic reprogramming in mammals. Human molecular genetics 14, R47–R58 (2005).

549 de Magalhães, J. P. Ageing as a software design flaw. Genome Biology 24, 51, doi:10.1186/s13059-023-02888-y (2023).

550 Wolf, A. M. The tumor suppression theory of aging. Mech Ageing Dev 200, 111583, doi:10.1016/j.mad.2021.111583 (2021).

551 Shay, J. W. Role of Telomeres and Telomerase in Aging and Cancer. Cancer Discov 6, 584–593, doi:10.1158/2159-8290.Cd-16-0062 (2016).

552 Tyner, S. D. et al. p53 mutant mice that display early ageing-associated phenotypes. Nature 415, 45–53, doi:10.1038/415045a (2002).

553 Kirkwood, T. B. L. p53 and ageing: too much of a good thing? BioEssays 24, 577–579, 10.1002/bies.10111 (2002).

554 Rodier, F., Campisi, J. & Bhaumik, D. Two faces of p53: aging and tumor suppression. Nucleic Acids Res 35, 7475–7484, doi:10.1093/nar/gkm744 (2007).

555 West, M. D. et al. Toward a Unified Theory of Aging and Regeneration. Regenerative Medicine 14, 867–886, doi:10.2217/rme-2019-0062 (2019).

556 Kerepesi, C., Zhang, B., Lee, S. G., Trapp, A. & Gladyshev, V. N. Epigenetic clocks reveal a rejuvenation event during embryogenesis followed by aging. Sci Adv 7, doi:10.1126/sciadv.abg6082 (2021).

557 Bidder, G. P. Growth and death. (éditeur non identifié, 1925).

558 Sinclair, D. A. & LaPlante, M. D. Lifespan: Why we age—And why we Don’t have to. (Atria books, 2019).

559 Yang, J.-H. et al. Loss of epigenetic information as a cause of mammalian aging. Cell 186, 305–326.e327, doi:10.1016/j.cell.2022.12.027 (2023).

560 Lu, Y. R., Tian, X. & Sinclair, D. A. The Information Theory of Aging. Nature Aging 3, 1486–1499, doi:10.1038/s43587-023-00527-6 (2023).

561 Poetsch, A. R., Boulton, S. J. & Luscombe, N. M. Genomic landscape of oxidative DNA damage and repair reveals regioselective protection from mutagenesis. Genome Biology 19, 215, doi:10.1186/s13059-018-1582-2 (2018).

562 Horvath, S. DNA methylation age of human tissues and cell types. Genome biology 14, 1–20, 10.1186/gb-2013-14-10-r115 (2013).

563 Hannum, G. et al. Genome-wide methylation profiles reveal quantitative views of human aging rates. Molecular cell 49, 359–367 (2013).

564 Poganik, J. R. et al. Biological age is increased by stress and restored upon recovery. Cell Metabolism 35, 807–820.e805, 10.1016/j.cmet.2023.03.015 (2023).

565 Kabacik, S. et al. The relationship between epigenetic age and the hallmarks of aging in human cells. Nature Aging 2, 484–493, doi:10.1038/s43587-022-00220-0 (2022).

566 Thompson, M. J. et al. A multi-tissue full lifespan epigenetic clock for mice. Aging (Albany NY) 10, 2832 (2018).

567 Horvath, S. et al. Obesity accelerates epigenetic aging of human liver. Proceedings of the National Academy of Sciences 111, 15538–15543, doi:doi:10.1073/pnas.1412759111 (2014).

568 Stubbs, T. M. et al. Multi-tissue DNA methylation age predictor in mouse. Genome Biology 18, 68, doi:10.1186/s13059-017-1203-5 (2017).

569 Lowe, D., Horvath, S. & Raj, K. Epigenetic clock analyses of cellular senescence and ageing. Oncotarget 7, 8524–8531 (2016).

570 Herzog, C. M. S. et al. Functionally enriched epigenetic clocks reveal tissue-specific discordant aging patterns in individuals with cancer. Communications Medicine 5, 98, doi:10.1038/s43856-025-00739-4 (2025).

571 Wang, H. et al. Association between advanced fibrosis and epigenetic age acceleration among individuals with MASLD. J Gastroenterol 60, 306–314, doi:10.1007/s00535-024-02181-0 (2025).

572 Peto, R., Roe, F. J., Lee, P. N., Levy, L. & Clack, J. Cancer and ageing in mice and men. Br J Cancer 32, 411–426, doi:10.1038/bjc.1975.242 (1975).

573 Austad, S. N. & Fischer, K. E. Mammalian aging, metabolism, and ecology: evidence from the bats and marsupials. J Gerontol 46, B47–53, doi:10.1093/geronj/46.2.b47 (1991).

574 Rubner, M. Das Problem det Lebensdaur und seiner beziehunger zum Wachstum und Ernarnhung. (1908).

575 Pearl, R. The rate of living: being an account of some experimental studies on the biology of life duration. (AA Knopf, 1928).

576 Speakman, J. R. Body size, energy metabolism and lifespan. Journal of Experimental Biology 208, 1717–1730, doi:10.1242/jeb.01556 (2005).

577 Miquel, J., Lundgren, P. R., Bensch, K. G. & Atlan, H. Effects of temperature on the life span, vitality and fine structure of Drosophila melanogaster. Mech Ageing Dev 5, 347–370, doi:10.1016/0047-6374(76)90034-8 (1976).

578 Conti, B. et al. Transgenic mice with a reduced core body temperature have an increased life span. Science 314, 825–828, doi:10.1126/science.1132191 (2006).

579 Zhao, Z. et al. Body temperature is a more important modulator of lifespan than metabolic rate in two small mammals. Nat Metab 4, 320–326, doi:10.1038/s42255-022-00545-5 (2022).

580 Stark, G., Pincheira-Donoso, D. & Meiri, S. No evidence for the ‘rate-of-living’ theory across the tetrapod tree of life. Global Ecology and Biogeography 29, 857–884, 10.1111/geb.13069 (2020).

581 Gkioni, L. et al. The geroprotectors trametinib and rapamycin combine additively to extend mouse healthspan and lifespan. Nature Aging 5, 1249–1265, doi:10.1038/s43587-025-00876-4 (2025).

582 Ureña, E. et al. Trametinib ameliorates aging-associated gut pathology in *Drosophila* females by reducing Pol III activity in intestinal stem cells. Proceedings of the National Academy of Sciences 121, e2311313121, doi:doi:10.1073/pnas.2311313121 (2024).

583 Sulak, M. et al. TP53 copy number expansion is associated with the evolution of increased body size and an enhanced DNA damage response in elephants. eLife 5, e11994, doi:10.7554/eLife.11994 (2016).

584 Tollis, M. et al. Return to the Sea, Get Huge, Beat Cancer: An Analysis of Cetacean Genomes Including an Assembly for the Humpback Whale (Megaptera novaeangliae). Molecular Biology and Evolution 36, 1746–1763, doi:10.1093/molbev/msz099 (2019).

585 Athar, F. et al. Limited cell-autonomous anticancer mechanisms in long-lived bats. Nature Communications 16, 4125, doi:10.1038/s41467-025-59403-z (2025).

586 Riddle, D. L., Swanson, M. M. & Albert, P. S. Interacting genes in nematode dauer larva formation. Nature 290, 668–671, doi:10.1038/290668a0 (1981).

587 Sutter, N. B. et al. A single IGF1 allele is a major determinant of small size in dogs. Science 316, 112–115, doi:10.1126/science.1137045 (2007).

588 Geiger, M., Gendron, K., Willmitzer, F. & Sánchez-Villagra, M. R. Unaltered sequence of dental, skeletal, and sexual maturity in domestic dogs compared to the wolf. Zoological Lett 2, 16, doi:10.1186/s40851-016-0055-2 (2016).

589 Strauss, A., Mascher, E., Palme, R. & Millesi, E. Sexually mature and immature yearling male European ground squirrels: a comparison of behavioral and physiological parameters. Horm Behav 52, 646–652, doi:10.1016/j.yhbeh.2007.08.003 (2007).

590 Gong, Z. et al. Reductions in serum IGF-1 during aging impair health span. Aging Cell 13, 408–418, doi:10.1111/acel.12188 (2014).

591 Kern, C. C. & Gems, D. Semelparous Death as one Element of Iteroparous Aging Gone Large. Front Genet 13, 880343, doi:10.3389/fgene.2022.880343 (2022).

592 McAllan, B. M. Dasyurid marsupials as models for the physiology of ageing in humans. Australian Journal of Zoology 54, 159–172, doi:10.1071/zo05073 (2006).

593 Naylor, R., Richardson, S. J. & McAllan, B. M. Boom and bust: a review of the physiology of the marsupial genus Antechinus. J Comp Physiol B 178, 545–562, doi:10.1007/s00360-007-0250-8 (2008).

594 Finch, C. E. Longevity, Senescence and the Genome. 95 (University of Chicago Press, 1990).

595 Tökölyi, J. et al. Canonical Wnt/β-catenin signalling regulates inducible aging and regeneration loss in hydra. bioRxiv, 2025.2004.2001.646560, doi:10.1101/2025.04.01.646560 (2025).

596 Hatton, I. A. et al. The human cell count and size distribution. Proc Natl Acad Sci U S A 120, e2303077120, doi:10.1073/pnas.2303077120 (2023).

597 Silver, B. B. & Nelson, C. M. The Bioelectric Code: Reprogramming Cancer and Aging From the Interface of Mechanical and Chemical Microenvironments. Front Cell Dev Biol 6, 21, doi:10.3389/fcell.2018.00021 (2018).

598 Herzig, S. & Shaw, R. J. AMPK: guardian of metabolism and mitochondrial homeostasis. Nature reviews. Molecular cell biology 19, 121–135, doi:10.1038/nrm.2017.95 (2018).

599 Yan, Y., Zhou, X. E., Xu, H. E. & Melcher, K. Structure and Physiological Regulation of AMPK. International Journal of Molecular Sciences 19, 3534 (2018).

600 Xiao, B. et al. Structure of mammalian AMPK and its regulation by ADP. Nature 472, 230–233, doi:10.1038/nature09932 (2011).

601 Gowans, G. J., Hawley, S. A., Ross, F. A. & Hardie, D. G. AMP is a true physiological regulator of AMP-activated protein kinase by both allosteric activation and enhancing net phosphorylation. Cell Metab 18, 556–566, doi:10.1016/j.cmet.2013.08.019 (2013).

602 Oakhill, J. S. et al. AMPK is a direct adenylate charge-regulated protein kinase. Science 332, 1433–1435, doi:10.1126/science.1200094 (2011).

603 Carling, D., Clarke, P. R., Zammit, V. A. & Hardie, D. G. Purification and characterization of the AMP-activated protein kinase. Copurification of acetyl-CoA carboxylase kinase and 3-hydroxy-3-methylglutaryl-CoA reductase kinase activities. Eur J Biochem 186, 129-136, doi:10.1111/j.1432-1033.1989.tb15186.x (1989).

604 Lawson, J. W. & Veech, R. L. Effects of pH and free Mg2+ on the Keq of the creatine kinase reaction and other phosphate hydrolyses and phosphate transfer reactions. J Biol Chem 254, 6528–6537 (1979).

605 Victor, V. M. et al. Effects of metformin on mitochondrial function of leukocytes from polycystic ovary syndrome patients with insulin resistance. European Journal of Endocrinology 173, 683–691 (2015).

606 Wang, Y. et al. Metformin Improves Mitochondrial Respiratory Activity through Activation of AMPK. Cell Rep 29, 1511–1523.e1515, doi:10.1016/j.celrep.2019.09.070 (2019).

607 McGarry, J. D., Mannaerts, G. P. & Foster, D. W. A possible role for malonyl-CoA in the regulation of hepatic fatty acid oxidation and ketogenesis. J Clin Invest 60, 265–270, doi:10.1172/jci108764 (1977).

608 Eaton, S. Control of mitochondrial β-oxidation flux. Progress in lipid research 41, 197–239 (2002).

609 Lee, W. J. et al. AMPK activation increases fatty acid oxidation in skeletal muscle by activating PPARα and PGC-1. Biochemical and biophysical research communications 340, 291–295 (2006).

610 Merrill, G. F., Kurth, E. J., Hardie, D. & Winder, W. AICA riboside increases AMP-activated protein kinase, fatty acid oxidation, and glucose uptake in rat muscle. American Journal of Physiology-Endocrinology And Metabolism 273, E1107–E1112 (1997).

611 Saha, A. K. et al. Activation of malonyl-CoA decarboxylase in rat skeletal muscle by contraction and the AMP-activated protein kinase activator 5-aminoimidazole-4-carboxamide-1-β-D-ribofuranoside. Journal of Biological Chemistry 275, 24279–24283 (2000).

612 Fredrickson, S. D. & Gordon, R. S. Transport of Fatty Acids. Physiological Reviews 38, 585–630, doi:10.1152/physrev.1958.38.4.585 (1958).

613 Glatz, J. F., Luiken, J. J. & Bonen, A. Membrane fatty acid transporters as regulators of lipid metabolism: implications for metabolic disease. Physiol Rev 90, 367–417, doi:10.1152/physrev.00003.2009 (2010).

614 Park, Y. M. CD36, a scavenger receptor implicated in atherosclerosis. Exp Mol Med 46, e99, doi:10.1038/emm.2014.38 (2014).

615 Sadur, C. N. & Eckel, R. H. Insulin stimulation of adipose tissue lipoprotein lipase. Use of the euglycemic clamp technique. J Clin Invest 69, 1119–1125, doi:10.1172/jci110547 (1982).

616 Boden, G. Role of fatty acids in the pathogenesis of insulin resistance and NIDDM. Diabetes 46, 3–10 (1997).

617 Bollheimer, L. C., Skelly, R. H., Chester, M. W., McGarry, J. D. & Rhodes, C. J. Chronic exposure to free fatty acid reduces pancreatic beta cell insulin content by increasing basal insulin secretion that is not compensated for by a corresponding increase in proinsulin biosynthesis translation. The Journal of Clinical Investigation 101, 1094–1101, doi:10.1172/JCI420 (1998).

618 McNelis, J. C. et al. GPR43 potentiates β-cell function in obesity. Diabetes 64, 3203–3217 (2015).

619 Villa, S. R. et al. Loss of free fatty acid receptor 2 leads to impaired islet mass and beta cell survival. Scientific reports 6, 1–9 (2016).

620 Fuller, M. et al. The short-chain fatty acid receptor, FFA2, contributes to gestational glucose homeostasis. American Journal of Physiology-Endocrinology and Metabolism 309, E840-E851 (2015).

621 Choi, S. M. et al. Insulin regulates adipocyte lipolysis via an Akt-independent signaling pathway. Mol Cell Biol 30, 5009–5020, doi:10.1128/mcb.00797-10 (2010).

622 Degerman, E. et al. Phosphorylation and activation of hormone-sensitive adipocyte phosphodiesterase type 3B. Methods 14, 43–53, doi:10.1006/meth.1997.0564 (1998).

623 Kitamura, T. et al. Insulin-induced phosphorylation and activation of cyclic nucleotide phosphodiesterase 3B by the serine-threonine kinase Akt. Molecular and cellular biology 19, 6286–6296 (1999).

624 Cunha, D. A. et al. Initiation and execution of lipotoxic ER stress in pancreatic beta-cells. J Cell Sci 121, 2308–2318, doi:10.1242/jcs.026062 (2008).

625 Katsoulieris, E., Mabley, J. G., Samai, M., Green, I. C. & Chatterjee, P. K. alpha-Linolenic acid protects renal cells against palmitic acid lipotoxicity via inhibition of endoplasmic reticulum stress. Eur J Pharmacol 623, 107–112, doi:10.1016/j.ejphar.2009.09.015 (2009).

626 Weigert, A., Jennewein, C. & Brüne, B. The liaison between apoptotic cells and macrophages--the end programs the beginning. Biol Chem 390, 379–390, doi:10.1515/bc.2009.048 (2009).

627 Haczeyni, F. et al. Exercise improves adipose function and inflammation and ameliorates fatty liver disease in obese diabetic mice. Obesity (Silver Spring) 23, 1845–1855, doi:10.1002/oby.21170 (2015).

628 Roberts, R. et al. Markers of de novo lipogenesis in adipose tissue: associations with small adipocytes and insulin sensitivity in humans. Diabetologia 52, 882–890, doi:10.1007/s00125-009-1300-4 (2009).

629 Maffeis, C. et al. Fat Cell Size, Insulin Sensitivity, and Inflammation in Obese Children. The Journal of Pediatrics 151, 647–652, 10.1016/j.jpeds.2007.04.053 (2007).

630 Soccio, R. E. & Breslow, J. L. Intracellular cholesterol transport. Arterioscler Thromb Vasc Biol 24, 1150–1160, doi:10.1161/01.ATV.0000131264.66417.d5 (2004).

631 Haczeyni, F., Bell-Anderson, K. S. & Farrell, G. C. Causes and mechanisms of adipocyte enlargement and adipose expansion. Obesity Reviews 19, 406–420, 10.1111/obr.12646 (2018).

632 Ring, A., Le Lay, S., Pohl, J., Verkade, P. & Stremmel, W. Caveolin-1 is required for fatty acid translocase (FAT/CD36) localization and function at the plasma membrane of mouse embryonic fibroblasts. Biochim Biophys Acta 1761, 416–423, doi:10.1016/j.bbalip.2006.03.016 (2006).

633 Gerbod-Giannone, M. C. et al. Involvement of caveolin-1 and CD36 in native LDL endocytosis by endothelial cells. Biochim Biophys Acta Gen Subj 1863, 830–838, doi:10.1016/j.bbagen.2019.01.005 (2019).

634 Shigematsu, S., Watson, R. T., Khan, A. H. & Pessin, J. E. The adipocyte plasma membrane caveolin functional/structural organization is necessary for the efficient endocytosis of GLUT4. J Biol Chem 278, 10683–10690, doi:10.1074/jbc.M208563200 (2003).

635 Robenek, H. et al. Topography of lipid droplet-associated proteins: insights from freeze-fracture replica immunogold labeling. Journal of Lipids 2011 (2011).

636 Le Lay, S. et al. Cholesterol-induced caveolin targeting to lipid droplets in adipocytes: a role for caveolar endocytosis. Traffic 7, 549–561 (2006).

637 Theofilis, P. et al. Inflammatory Mechanisms Contributing to Endothelial Dysfunction. Biomedicines 9, doi:10.3390/biomedicines9070781 (2021).

638 Qian, H. et al. TNFα Induces and Insulin Inhibits Caspase 3-Dependent Adipocyte Apoptosis. Biochemical and Biophysical Research Communications 284, 1176–1183, 10.1006/bbrc.2001.5100 (2001).

639 Gao, Z. et al. P65 inactivation in adipocytes and macrophages attenuates adipose inflammatory response in lean but not in obese mice. Am J Physiol Endocrinol Metab 308, E496–505, doi:10.1152/ajpendo.00532.2014 (2015).

640 Díaz, B., Pimentel, B., de Pablo, F. & de La Rosa, E. J. Apoptotic cell death of proliferating neuroepithelial cells in the embryonic retina is prevented by insulin. Eur J Neurosci 11, 1624–1632, doi:10.1046/j.1460-9568.1999.00577.x (1999).

641 Tanaka, M., Sawada, M., Yoshida, S., Hanaoka, F. & Marunouchi, T. Insulin prevents apoptosis of external granular layer neurons in rat cerebellar slice cultures. Neurosci Lett 199, 37–40, doi:10.1016/0304-3940(95)12009-s (1995).

642 Schaeffler, A. et al. Fatty acid-induced induction of Toll-like receptor-4/nuclear factor-kappaB pathway in adipocytes links nutritional signalling with innate immunity. Immunology 126, 233–245, doi:10.1111/j.1365-2567.2008.02892.x (2009).

643 Pagliassotti, M., Wang, D. & Wei, Y. (Wiley Online Library, 2006).

644 Zhang, Y. & Bliska, J. B. Role of Toll-like receptor signaling in the apoptotic response of macrophages to Yersinia infection. Infect Immun 71, 1513–1519, doi:10.1128/iai.71.3.1513-1519.2003 (2003).

645 Gregor, M. F. & Hotamisligil, G. S. Thematic review series: Adipocyte Biology. Adipocyte stress: the endoplasmic reticulum and metabolic disease. J Lipid Res 48, 1905–1914, 10.1194/jlr.R700007-JLR200 (2007).

646 Shi, H. et al. TLR4 links innate immunity and fatty acid-induced insulin resistance. J Clin Invest 116, 3015–3025, doi:10.1172/jci28898 (2006).

647 Fischer-Posovszky, P., Tornqvist, H., Debatin, K. M. & Wabitsch, M. Inhibition of death-receptor mediated apoptosis in human adipocytes by the insulin-like growth factor I (IGF-I)/IGF-I receptor autocrine circuit. Endocrinology 145, 1849–1859, doi:10.1210/en.2003-0985 (2004).

648 Tseng, Y. H., Ueki, K., Kriauciunas, K. M. & Kahn, C. R. Differential roles of insulin receptor substrates in the anti-apoptotic function of insulin-like growth factor-1 and insulin. J Biol Chem 277, 31601–31611, doi:10.1074/jbc.M202932200 (2002).

649 Matarese, G. The link between obesity and autoimmunity. Science 379, 1298–1300, doi:doi:10.1126/science.ade0113 (2023).

650 Digel, M. et al. FATP4 contributes as an enzyme to the basal and insulin-mediated fatty acid uptake of C2C12 muscle cells. American Journal of Physiology-Endocrinology and Metabolism 301, E785–E796, doi:10.1152/ajpendo.00079.2011 (2011).

651 Williams, C. M. Lipid metabolism in women. Proceedings of the Nutrition Society 63, 153–160, doi:10.1079/PNS2003314 (2004).

652 Finn, P. F. & Dice, J. F. Proteolytic and lipolytic responses to starvation. Nutrition 22, 830–844, 10.1016/j.nut.2006.04.008 (2006).

653 Pedersen, B. K. & Saltin, B. Evidence for prescribing exercise as therapy in chronic disease. Scandinavian journal of medicine & science in sports 16, 3–63 (2006).

654 Kokkinos, P. F., Faselis, C., Myers, J., Panagiotakos, D. & Doumas, M. Interactive effects of fitness and statin treatment on mortality risk in veterans with dyslipidaemia: a cohort study. The Lancet 381, 394–399 (2013).

655 Kuzuya, M., Ando, F., Iguchi, A. & Shimokata, H. Changes in serum lipid levels during a 10 year period in a large Japanese population: a cross-sectional and longitudinal study. Atherosclerosis 163, 313–320 (2002).

656 Upmeier, E. et al. Longitudinal changes in serum lipids in older people The Turku Elderly Study 1991–2006. Age and Ageing 40, 280–283, doi:10.1093/ageing/afq180 (2011).

657 Hoorens, A., Van de Casteele, M., Klöppel, G. & Pipeleers, D. Glucose promotes survival of rat pancreatic beta cells by activating synthesis of proteins which suppress a constitutive apoptotic program. J Clin Invest 98, 1568–1574, doi:10.1172/jci118950 (1996).

658 Swenne, I. The role of glucose in the in vitro regulation of cell cycle kinetics and proliferation of fetal pancreatic B-cells. Diabetes 31, 754–760, doi:10.2337/diab.31.9.754 (1982).

659 Liu, J.-L., Segovia, I., Yuan, X.-L. & Gao, Z.-h. Controversial Roles of Gut Microbiota-Derived Short-Chain Fatty Acids (SCFAs) on Pancreatic β-Cell Growth and Insulin Secretion. International Journal of Molecular Sciences 21, 910 (2020).

660 Tuo, Y. et al. Long-term in vitro treatment of INS-1 rat pancreatic β-cells by unsaturated free fatty acids protects cells against gluco- and lipotoxicities via activation of GPR40 receptors. Clin Exp Pharmacol Physiol 39, 423–428, doi:10.1111/j.1440-1681.2012.05691.x (2012).

661 Shimabukuro, M., Zhou, Y. T., Levi, M. & Unger, R. H. Fatty acid-induced beta cell apoptosis: a link between obesity and diabetes. Proc Natl Acad Sci U S A 95, 2498–2502, doi:10.1073/pnas.95.5.2498 (1998).

662 Kristinsson, H., Smith, D. M., Bergsten, P. & Sargsyan, E. FFAR1 is involved in both the acute and chronic effects of palmitate on insulin secretion. Endocrinology 154, 4078–4088, doi:10.1210/en.2013-1352 (2013).

663 Efanova, I. B. et al. Glucose and tolbutamide induce apoptosis in pancreatic beta-cells. A process dependent on intracellular Ca2+ concentration. J Biol Chem 273, 33501-33507, doi:10.1074/jbc.273.50.33501 (1998).

664 Oh, Y. S., Bae, G. D., Baek, D. J., Park, E. Y. & Jun, H. S. Fatty Acid-Induced Lipotoxicity in Pancreatic Beta-Cells During Development of Type 2 Diabetes. Front Endocrinol (Lausanne) 9, 384, doi:10.3389/fendo.2018.00384 (2018).

665 McGill, H. C., Jr., et al. Obesity accelerates the progression of coronary atherosclerosis in young men. Circulation 105, 2712–2718, doi:10.1161/01.cir.0000018121.67607.ce (2002).

666 Libby, P. Inflammation in Atherosclerosis—No Longer a Theory. Clinical Chemistry 67, 131–142, doi:10.1093/clinchem/hvaa275 (2020).

667 Westerterp, K. R. Physical activity, food intake, and body weight regulation: insights from doubly labeled water studies. Nutrition reviews 68, 148–154 (2010).

668 Farias, M. M., Cuevas, A. M. & Rodriguez, F. Set-point theory and obesity. Metab Syndr Relat Disord 9, 85–89, doi:10.1089/met.2010.0090 (2011).

669 Pressler, M. et al. Dietary Transitions and Health Outcomes in Four Populations - Systematic Review. Front Nutr 9, 748305, doi:10.3389/fnut.2022.748305 (2022).

670 Webb, P. & Annis, J. F. Adaptation to overeating in lean and overweight men and women. Hum Nutr Clin Nutr 37, 117–131 (1983).

671 Fisler, J. S. & Warden, C. H. Uncoupling proteins, dietary fat and the metabolic syndrome. Nutrition & Metabolism 3, 38, doi:10.1186/1743-7075-3-38 (2006).

672 Joosen, A. M. C. P., Bakker, A. H. F. & Westerterp, K. R. Metabolic efficiency and energy expenditure during short-term overfeeding. Physiology & Behavior 85, 593–597, 10.1016/j.physbeh.2005.06.006 (2005).

673 Kazak, L. et al. Genetic Depletion of Adipocyte Creatine Metabolism Inhibits Diet-Induced Thermogenesis and Drives Obesity. Cell Metab 26, 660–671.e663, doi:10.1016/j.cmet.2017.08.009 (2017).

674 Bachman, E. S. et al. βAR signaling required for diet-induced thermogenesis and obesity resistance. Science 297, 843–845 (2002).

675 Rothwell, N. J. & Stock, M. J. A role for brown adipose tissue in diet-induced thermogenesis. Nature 281, 31–35 (1979).

676 Goedeke, L. & Shulman, G. I. Therapeutic potential of mitochondrial uncouplers for the treatment of metabolic associated fatty liver disease and NASH. Molecular Metabolism 46, 101178, 10.1016/j.molmet.2021.101178 (2021).

677 Tippetts, T. S., Holland, W. L. & Summers, S. A. Cholesterol - the devil you know; ceramide - the devil you don’t. Trends Pharmacol Sci 42, 1082–1095, doi:10.1016/j.tips.2021.10.001 (2021).

678 Yeo, E.-J. Hypoxia and aging. Experimental & Molecular Medicine 51, 1–15, doi:10.1038/s12276-019-0233-3 (2019).

679 Perea, A., Clemente, F., Martinell, J., Villanueva-Peñacarrillo, M. L. & Valverde, I. Physiological effect of glucagon in human isolated adipocytes. Hormone and metabolic research 27, 372–375 (1995).

680 Hagen, J. H. Effect of glucagon on the metabolism of adipose tissue. J Biol Chem 236, 1023–1027 (1961).

681 Prigge, W. F. & Grande, F. Effects of glucagon, epinephrine and insulin on in vitro lipolysis of adipose tissue from mammals and birds. Comparative Biochemistry and Physiology Part B: Comparative Biochemistry 39, 69–82, 10.1016/0305-0491(71)90254-9 (1971).

682 Rodbell, M. & Jones, A. B. Metabolism of isolated fat cells. 3. The similar inhibitory action of phospholipase C (Clostridium perfringens alpha toxin) and of insulin on lipolysis stimulated by lipolytic hormones and theophylline. J Biol Chem 241, 140–142 (1966).

683 Galsgaard, K. D., Pedersen, J., Knop, F. K., Holst, J. J. & Wewer Albrechtsen, N. J. Glucagon Receptor Signaling and Lipid Metabolism. Front Physiol 1, 413, doi:10.3389/fphys.2019.00413 (2019).

684 Paloyan, E. & Harper, P. V., Jr. Glucagon as a regulating factor of plasma lipids. Metabolism 10, 315–323 (1961).

685 Habegger, K. M. et al. The metabolic actions of glucagon revisited. Nat Rev Endocrinol 6, 689–697, doi:10.1038/nrendo.2010.187 (2010).

686 Jensen, M. D., Heiling, V. J. & Miles, J. M. Effects of glucagon on free fatty acid metabolism in humans. J Clin Endocrinol Metab 72, 308–315, doi:10.1210/jcem-72-2-308 (1991).

687 Vasileva, A., Marx, T., Beaudry, J. L. & Stern, J. H. Glucagon Receptor Signaling at White Adipose Tissue Does Not Regulate Lipolysis. Am J Physiol Endocrinol Metab, doi:10.1152/ajpendo.00078.2022 (2022).

688 Skurk, T., Alberti-Huber, C., Herder, C. & Hauner, H. Relationship between adipocyte size and adipokine expression and secretion. The Journal of Clinical Endocrinology & Metabolism 92, 1023–1033 (2007).

689 Roberts, A. W. et al. Genetic influences determining progenitor cell mobilization and leukocytosis induced by granulocyte colony-stimulating factor. Blood, The Journal of the American Society of Hematology 89, 2736–2744 (1997).

690 Lévesque, J. P., Hendy, J., Takamatsu, Y., Simmons, P. J. & Bendall, L. J. Disruption of the CXCR4/CXCL12 chemotactic interaction during hematopoietic stem cell mobilization induced by GCSF or cyclophosphamide. J Clin Invest 111, 187–196, doi:10.1172/jci15994 (2003).

691 Fattorossi, A. et al. Effects of granulocyte-colony-stimulating factor and granulocyte/macrophage-colony-stimulating factor administration on T cell proliferation and phagocyte cell-surface molecules during hematopoietic reconstitution after autologous peripheral blood progenitor cell transplantation. Cancer Immunology, Immunotherapy 49, 641–648 (2001).

692 Bermudez, L., Petrofsky, M. & Stevens, P. Treatment with recombinant granulocyte colony-stimulating factor (FilgrastinTM) stimulates neutrophils and tissue macrophages and induces an effective non-specific response against Mycobacterium avium in mice. Immunology 94, 297–303 (1998).

693 McEwen, B. S. Stress, adaptation, and disease. Allostasis and allostatic load. Ann N Y Acad Sci 840, 33–44, doi:10.1111/j.1749-6632.1998.tb09546.x (1998).

694 Hirasaka, K. et al. Deficiency of Cbl-b gene enhances infiltration and activation of macrophages in adipose tissue and causes peripheral insulin resistance in mice. Diabetes 56, 2511–2522 (2007).

695 Kamei, N. et al. Overexpression of monocyte chemoattractant protein-1 in adipose tissues causes macrophage recruitment and insulin resistance. Journal of biological chemistry 281, 26602–26614 (2006).

696 Arkan, M. C. et al. IKK-β links inflammation to obesity-induced insulin resistance. Nature medicine 11, 191–198 (2005).

697 Solinas, G. et al. JNK1 in hematopoietically derived cells contributes to diet-induced inflammation and insulin resistance without affecting obesity. Cell metabolism 6, 386–397 (2007).

